# *f*-statistics estimation and admixture graph construction with Pool-Seq or allele count data using the R package *poolfstat*

**DOI:** 10.1101/2021.05.28.445945

**Authors:** Mathieu Gautier, Renaud Vitalis, Laurence Flori, Arnaud Estoup

## Abstract

By capturing various patterns of the structuring of genetic variation across populations, *f* -statistics have proved highly effective for the inference of demographic history. Such statistics are defined as covariance of SNP allele frequency differences among sets of populations without requiring haplotype information and are hence particularly relevant for the analysis of pooled sequencing (Pool-Seq) data. We here propose a reinterpretation of the *F* (and *D*) parameters in terms of probability of gene identity and derive from this unified definition unbiased estimators for both Pool-Seq and standard allele count data obtained from individual genotypes. We implemented these estimators in a new version of the R package `poolfstat`, which now includes a wide range of inference methods: (i) three-population test of admixture; (ii) four-population test of treeness; (iii) *F*_4_-ratio estimation of admixture rates; and (iv) fitting, visualization and (semi-automatic) construction of admixture graphs. A comprehensive evaluation of the methods implemented in `poolfstat` on both simulated Pool-Seq (with various sequencing coverages and error rates) and allele count data confirmed the accuracy of these approaches, even for the most cost-effective Pool-Seq design involving low sequencing coverages. We further analyzed a real Pool-Seq data made of 14 populations of the invasive species *Drosophila suzukii* which allowed refining both the demographic history of native populations and the invasion routes followed by this emblematic pest. Our new package `poolfstat` provides the community with a user-friendly and efficient all-in-one tool to unravel complex population genetic histories from large-size Pool-Seq or allele count SNP data.

## 1 Introduction

In their seminal paper, Reich *et al* (2009) introduced a new population genetics framework to decipher the history of Indian human populations. This inference approach relied on a set of so-called *f* -statistics that are aimed at capturing various patterns of the structuring of genetic diversity across-population based on Single Nucleotide Polymorphism (SNP) assayed on a genome-wide scale (see also Patterson *et al*, 2012). The parameters underlying these statistics and denoted *F* following Patterson *et al* (2012) are defined as covariances in allele frequency difference among sets of two (*F*_2_), three (*F*_3_) or four (*F*_4_) populations and were demonstrated to be highly informative about populations demographic history when modeled as admixture graphs, i.e., population trees possibly including admixture events (Patterson *et al*, 2012). Hence, formal tests of admixture, called *three–population* test, between a target population and two source population surrogates can be derived from estimates of *F*_3_. Conversely, via the so-called *four–population* test, estimating *F*_4_ among quadruplets of populations allows to test for their treeness, i.e., if their joint history can be modeled as a simple (unrooted) bifurcating tree. Under certain restrictive assumptions about the underlying phylogeny, accurate estimates of the relative contributions of the ancestral sources of an admixed population may be obtained from ratios of *F*_4_ involving some of its related sampled populations. A normalized version of the *F*_4_ parameter, called Patterson’s *D*, was also developed by Green *et al* (2010) and has become quite popular to characterize introgression in phylogenies of closely related species (Durand *et al*, 2011). Finally, *f* -statistics can directly be used to fit admixture graphs (i.e., estimate branch lengths and/or admixture proportions) and to rigorously assess their support (Patterson *et al*, 2012; Lipson *et al*, 2013; Lipson, 2020).

A critical advantage of *F* and *D* parameters is that they only depend on population allele frequencies and their estimation does not require haplotype information. The non-independence of neighboring SNPs (Linkage Disequilibrium or LD) can be accurately accounted for with block-jackknife statistical techniques (Patterson *et al*, 2012; Kunsch, 1989; Reich *et al*, 2009; Busing *et al*, 1999) when computing standard errors of the estimated *f* -statistics which are noticeably required for the derivation of formal tests of admixture or treeness and also to assess the residuals of fitted admixture graphs. These characteristics make the *f* -statistics based inference framework particularly attractive for the analysis of Pool-Seq data that result from the massive sequencing of pools of individual DNA and have become quite popular, most particularly in non-model species (Schlötterer *et al*, 2014). Indeed, although LD information is generally lost in Pool-Seq experiments (but see Long *et al*, 2011; or Feder *et al*, 2012), they lead to accurate and cost-effective assessment of allele frequencies across populations on a whole genome basis (Gautier *et al*, 2013; Schlötterer *et al*, 2014). If the derivation of unbiased estimates of allele frequencies from Pool-Seq data is straightforward, estimation of more elaborated population genetics parameters characterizing the structuring of genetic diversity within or across populations is more challenging (Gautier *et al*, 2013; Ferretti *et al*, 2013; Hivert *et al*, 2018). As the individual origin of the sequencing reads is not identifiable within pools, it is not possible to assess whether reads are identical because they are sequenced copies of the same individual chromosome or because they are copies of different chromosomes carrying the same allele. The resulting additional level of variation thus needs to be accounted for in the estimation which, in contrast to the nucleotide diversities (heterozygosities) or the well-known *F*_*ST*_ differentiation measure (Ferretti *et al*, 2013; Hivert *et al*, 2018), has to our knowledge not been investigated for the estimation of *F* and *D* parameters (but see Leblois *et al*, 2018; Collin *et al*, 2021).

In the present paper, we first propose a (re)interpretation of the different *F* and *D* parameters in terms of probability of identity in state (IIS or AIS for Alike-In-State) of pairs of genes sampled either within the same population (*Q*_1_) or between two different populations (*Q*_2_), extending results we introduced in some earlier studies (Hivert *et al*, 2018; Leblois *et al*, 2018; Collin *et al*, 2021). This unified definition simplified the derivation of the unbiased estimators for both allele-count and Pool-Seq read count data, that we implemented in a new version of our R package `poolfstat` (Hivert *et al*, 2018) together with methods that rely on the estimated f– statistics for historical and demographic inference. These methods include i) three-population test of admixture; ii) four-population test of treeness; iii) *F*_4_-ratio estimation of admixture proportion; and iv) fitting, visualization and (semi-automatic) construction of admixture graphs. For completion, we briefly present the underlying methods as implemented in the package. We then carried out a comprehensive evaluation of the whole package on both simulated allele count and Pool-Seq read count data, considering for the latter various sequencing coverages and the presence or not of sequencing errors. Finally, we illustrate the power and limitations of `poolfstat` by analyzing real Pool-Seq data available from a previous study (Olazcuaga *et al*, 2020) for 14 populations of the invasive species *Drosophila suzukii*. This example illustrates how *f* -statistics based inference and admixture graph construction may confirm previous inferences and provide new insights into both the history of populations from the native area and the invasion routes followed by an emblematic invasive species. We provide as Supplementary Materials, a first vignette (Supplementary Vignette V1) designed as a detailed hands-on manual to outline the main functionalities of `poolfstat` and a second vignette (Supplementary Vignette V2) detailing the analysis of the *D. suzukii* data to make it fully reproducible.

## 2 Material and Methods

### 2.1 Definition, estimation and *f* −statistics based inference methods

#### 2.1.1 A unified definition of *F*_2_, *F*_3_ and *F*_4_ parameter and their scaled version *F*_*ST*_, 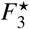 and *D* in terms of *Q*_1_ and *Q*_2_ probabilities

Let *p*_*A*_,*p*_*B*_, *p*_*C*_ and *p*_*D*_ the allele frequency of an arbitrarily chosen allele at a random SNP segregating in populations *A, B, C* and *D* respectively. The parameters *F*_2_, *F*_3_ and *F*_4_ were originally defined in terms of covariance in allele frequencies difference among different sets of populations as follows (Reich *et al*, 2009; Patterson *et al*, 2012):

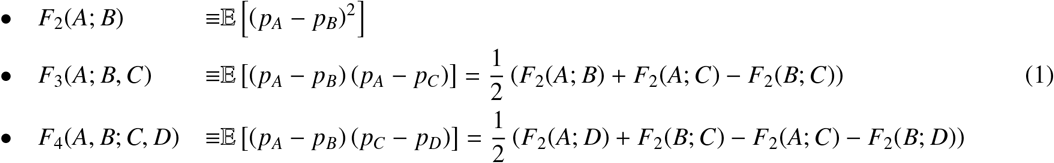

In total, with *n* populations, there are 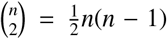 possible *F*_2_; 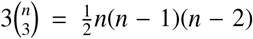 possible *F*_3_; and 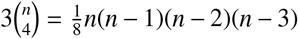 possible *F*_4_. Note that these values exclude the alternative equivalent configurations that result from the permutation of populations within pairs (since *F*_2_(*A*; *B*) = *F*_2_(*B*; *A*); *F*_3_(*A*; *B, C*) = *F*_3_(*A*; *C, B*) and *F*_4_(*A, B*; *C, D*) = *F*_4_(*B, A*; *D, C*) = −*F*_4_(*B, A*; *C, D*) …). Due to the linear dependency of all these parameters (eq. 1), the 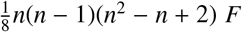 parameters actually span a vector space of dimension 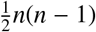 the basis of which may be specified by the set of all the 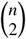 possible *F*_2_ or, given a reference population *i* (randomly chosen among the *n* ones) the set of all the *n* − 1 *F*_2_ of the form *F*_2_(*i*; *j*) (with *j* ≠ *i*) and all the 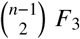 of the form *F*_3_(*i*; *j, k*) (with *j* ≠ *i*; *k* ≠ *i* and *j* ≠ *k*) (Patterson *et al*, 2012; Lipson, 2020). As mentioned by Patterson *et al* (2012), it is important to notice that these definitions are invariant in the choice of the reference SNP allele since:

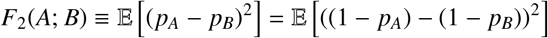

It directly follows from this property that:

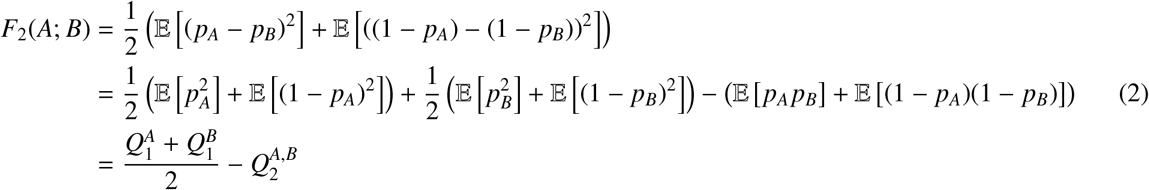

where 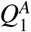 (resp. 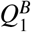) is actually the probability of sampling two genes (or alleles) identical in state (IIS) within population *A* (resp. *B*) and 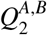 is the probability of sampling two IIS genes from *A* and *B*. It directly follows from equations 1 and 2 that:

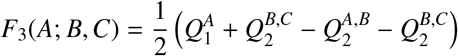

and,

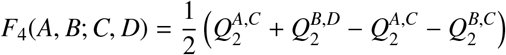

The *Q*_1_ and *Q*_2_ probabilities, and hence the *F*_2_, *F*_3_ and *F*_4_ parameters depend on both demographic parameters (i.e., population sizes, divergence times and other historical events) and marker polymorphism (i.e., their mutation rates and ascertainment process). For instance, under a simple pure-drift model with no mutation, if *p*_*r*_ denotes the allele frequency of the ancestral population *R* of two isolated populations *A* and *B* then 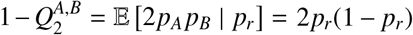 which is the heterozygosity in *R*. Similarly, 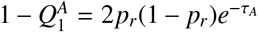 (resp., 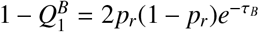) where *τ*_*A*_ (resp. *τ*_*B*_) is the divergence time separating *R* and *A* (resp. *B*) on a diffusion timescale (i.e., in drift units of 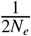 where *N*_*e*_ is the effective population along the branch). As a consequence, the resulting estimates of *F*_2_, *F*_3_ and *F*_4_ strongly depend on the underlying set of genetic markers and may not be compared across different datasets, even from the same populations. Various scaling procedure may actually helps in reducing this dependence. Scaling the *F*_2_ with respect to the across population heterozygosity 1 − *Q*_2_ leads to the standard definition of pairwise-population *F*_*ST*_ in terms of IIS probabilities (Rousset, 2007; Hivert *et al*, 2018) which is also concordant with its original definition as the numerator of *F*_*ST*_ (Reich *et al*, 2009; Peter, 2016):

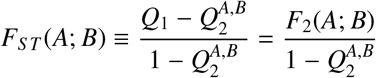

where 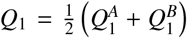 is the overall probability of sample two IIS genes within the same population (i.e., averaged over populations *A* and *B*). Similarly, the scaled versions of the *F*_3_ and *F*_4_ statistics named 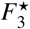 and *D* respectively (Patterson *et al*, 2012; Green *et al*, 2010; Durand *et al*, 2011), can be expressed as: 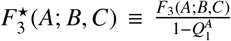 and 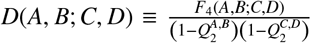. To sum up, expressions of the *F* and *D* parameters as a function of *Q*_1_ and *Q*_2_ probability are finally defined as follows:

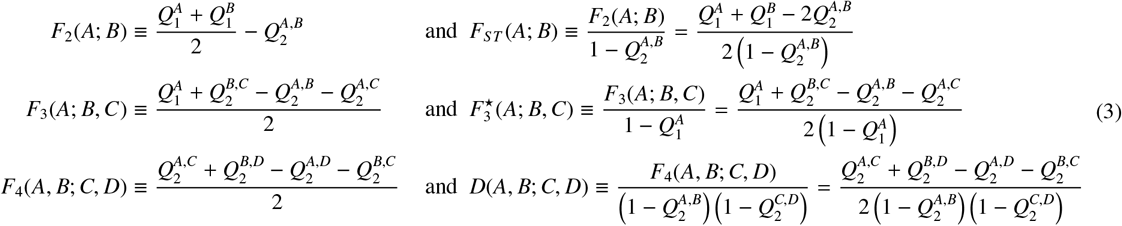

#### 2.1.2 Unbiased parameter estimators from Pool-Seq read count and standard allele count data

Let *y*_*ij*_ be the allele count for an arbitrarily chosen reference allele and *n*_*ij*_ the total number of sampled alleles (e.g., twice the number of genotyped individuals for a diploid species) at SNP *i* in population *j*. For Pool–Seq read count data, the *y*_*ij*_’s are not observed and for a given pool *j*, it is assumed that *n*_*ij*_ = *n*_*j*_ (the haploid sample size) for each and every SNP. We thus similarly defined *r*_*ij*_ as the read counts for the reference allele and *c*_*ij*_ the overall coverage observed at SNP *i* in population *j*.

If allele count data are directly observed, unbiased estimators of the IIS probability within population 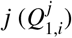 and between a pair of populations *j* and 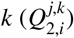 for a given SNP *i* are:

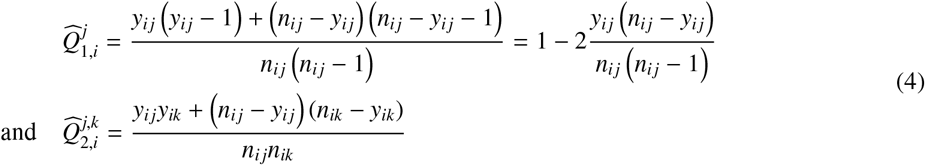

For Pool–Seq read count, unbiased estimators of *Q*_1,*i*_ and *Q*_2,*i*_ are similarly defined as (Hivert *et al*, 2018, eqns A37 and A40):

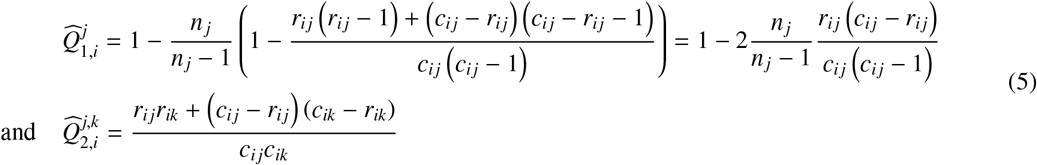

Genome-wide estimates of all the parameters defined in eq. 3 above are then simply obtained from these unbiased estimators of IIS probabilities over all the *I* SNPs as:

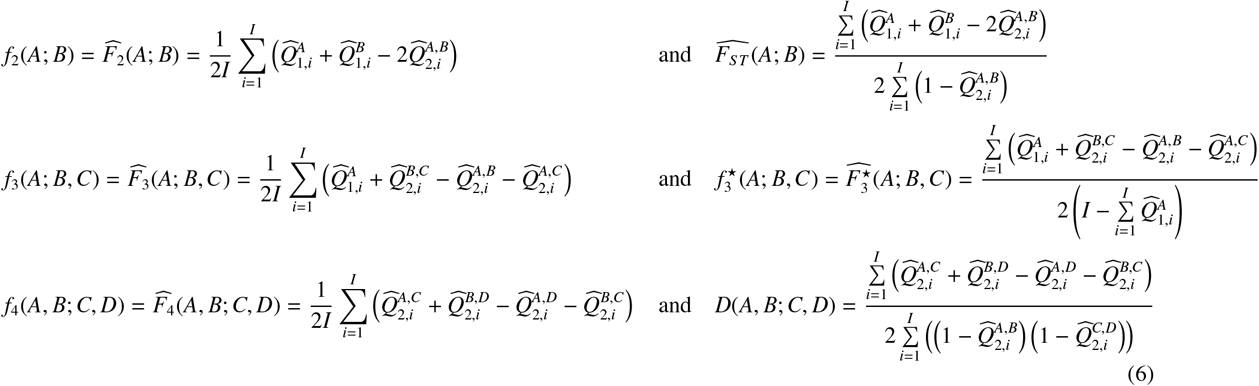

Similarly, the within-population heterozygosity 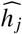 for each population is simply estimated as:

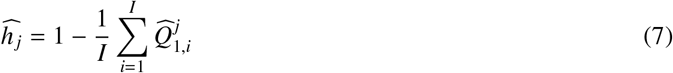

Importantly, for the three scaled parameters *F*_*ST*_, 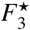 and *D*, multi-locus estimators consist of ratios of the numerator and denominator averages and not average of ratios (see e.g., Rousset, 2007; Patterson *et al*, 2012; Bhatia *et al*, 2013; Weir & Goudet, 2017; Hivert *et al*, 2018). Hence, for pairwise *F*_*ST*_, the above estimator is similar to the one described in Rousset (2007) for allele count data and identical to the alternative PID estimator described in Hivert *et al* (2018) for Pool-Seq read count data (so-called “Identity” method of the `computeFST` function from the `poolfstat` package).

#### 2.1.3 Block-Jackknife estimation of standard errors

Following Reich *et al* (2009), standard-errors of genome-wide estimates of the different statistics are computed using block-jackknife (Kunsch, 1989; Busing *et al*, 1999) which consists of dividing the genome into contiguous chunks of a predefined number of SNPs and then removing each block in turn to quantify the variability of the estimator. For a given parameter *F*, if *n*_*b*_ blocks are available and 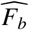 is the estimated statistics when removing all SNPs belonging to block *b*, the standard error 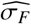 of the genome-wide estimator 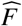 is computed as:

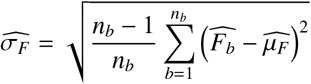

where 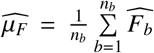, which may be slightly different than the estimator obtained with all the *I* markers since the latter may include SNPs that are not eligible for block-jackknife sampling (e.g., those at the chromosome or scaffolds boundaries). Finally, block-jackknife sampling may also be used to obtain estimates of the error covariance between two estimates 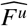 and 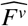 as:

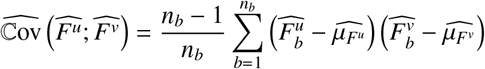

For convenience, we here chose to specify the same number of SNPs for each block instead of a block size in genetic distance (Patterson *et al*, 2012; Reich *et al*, 2009). We therefore do not recourse to a weighted block-jackknife (Busing *et al*, 1999). In practice, this has little impact providing the distribution of markers is homogeneous along the genome and the amount of missing data is negligible.

#### 2.1.4 Admixture Graph fitting

The approach implemented in the new version of `poolfstat` to fit admixture graphs from *f* −statistics is directly inspired from the one proposed by Patterson *et al* (2012) and implemented in the `qpGraph` software (see also Lipson, 2020). Briefly, let 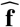 the vector (of length 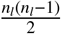 where *n*_*l*_ is the number of graph leaves) of the estimated *f*_2_ and *f*_3_ statistics forming the basis of all the *f* −statistics (see above). Similarly, let **g**(**e**; **a**) = **X**(**a**) × **e** the vector of their expected values given the graph edge lengths vector **e** and an incidence matrix **X**(**a**), which summarize the structure of the graph given the vector **a** of proportions of all admixture events (for a tree-topology, **X**(**a**) only consists of 0 or 1). In `poolfstat`, **X**(**a**) is derived using simple operations from another *n*_*l*_ by *n*_*e*_ matrix (where *n*_*e*_ is the number of graph edges) that specifies the weights of each edge along all the paths connecting the graph leaves to the root. It should be noticed that an admixture event is modeled as an instantaneous mixing of two populations *S* _1_ and *S* _2_ into a population *S* directly ancestral to a child population *A*. An admixture event may thus be specified by i) one admixture rate *α* quantifying the relative *S* _1_ and *S* _2_ ancestry proportions (*α* and 1−*α*) in population *S* ; and ii) three edge lengths *e*_*S*↔*A*_ for the branch connecting *S* and *A* and 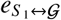 and 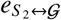 for the branches connecting the two source populations to the rest of the graph 𝒢. Yet, these three edge lengths are not identifiable and can only be estimated jointly in a single compound parameter (Pickrell & Pritchard, 2012; Patterson *et al*, 2012; Lipson, 2020): 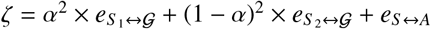. Following Lipson (2020), this identifiability issue is solved by setting 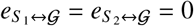 (i.e., nullifying the edges connecting the two source populations to the graph). Although it has no impact on the interpretation of the graph, this may overestimate the length of *e*_*S*↔*A*_ (i.e, the divergence between the admixed population *A* and its direct ancestor *S*). Proceeding this way differs from the choice made by Pickrell & Pritchard (2012) in the `Treemix` package which consists, following our notations, of setting 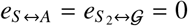 if 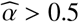 and 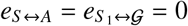 otherwise.

We finally define **Q** as the 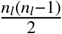 by 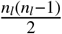 covariance matrix of the basis *f* −statistics estimated by block-jackknife. Graph fitting consists of finding the graph parameter values (**ê** and **â**) that minimize a cost (score of the model) defined as:

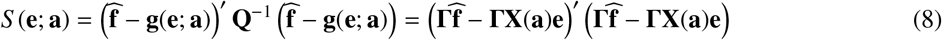

where **Γ** results from the Cholesky decomposition of **Q**^−1^ (i.e., **Q**^−1^ = **Γ**′ **Γ**). Given admixture rates **a**, *S* (**e**; **a**) is quadratic in the edge lengths **e** (Patterson *et al*, 2012) leading us to rely on the Lawson-Hanson non-negative linear least squares algorithm implemented in the R package `nnls` (Lawson & Hanson, 1995)) to estimate the vector **ê** that minimizes *S* (**e**; **a**) (subject to the constraint of positive edge lengths). Full minimization of *S* (**e**; **a**) is thus reduced to the identification of the admixture rates **a** which is performed using the L-BFGS-B algorithm implemented in the `optim` function of the R package `stats` (Nocedal & Wright, 1999).

#### 2.1.5 Confidence Intervals and model fit assessment

Assume 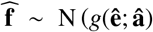, i.e., the vector of the basis *f* −statistics follows a multivariate normal distribution centered on the vector *g*(**ê**; **â**) specified by the fitted admixture graph parameters and the estimated error covariance matrix *Q*. The optimized score *S* (**ê**; **â**) then verifies *S* (**ê**; **â**) = −2*log*(*L*) − *K* where *L* is the likelihood of the fitted graph and *K* = *n* log(2*π*) + log(|**Q**|). This makes it straightforward to compute a *BIC* (Bayesian Information Criterion) for the fitted graph from the optimized score as:

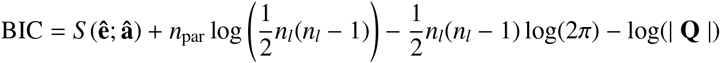

*BIC* may then be useful to compare different fitted admixture graph topologies. When comparing two graphs 𝒢_1_ and 𝒢_2_ with BIC equal to *BIC*_1_ and *BIC*_2_ respectively, we have Δ_12_ = *BIC*_2_ − *BIC*_1_ ≃ 2 log (*BF*_12_) where *BF*_12_ is the Bayes Factor associated to the comparison of the graphs 𝒢_1_ and 𝒢_2_ (Kass & Raftery, 1995, eq. 9). We may further rely on the modified Jeffreys’ rule proposed by Kass & Raftery (1995) to assess to which extent the data support either the 𝒢_1_ or 𝒢_2_ graphs, with Δ_12_ > 6 (respectively Δ_12_ > 10) providing “strong” (respectively “very strong”) evidence in favor of 𝒢_1_ (Supplementary Vignette V1).

Moreover, the likelihood interpretation of the optimized score *S* (**ê**; **â**) = −2*log*(*L*) − *K* allows constructing confidence intervals (CI) for the fitted parameters of a given graph (i.e., elements of the **e** and **a** vectors) using the following uni-dimensional procedure. For a given parameter *ν* (either a edge length or an admixture rate), the difference *S* _*ν*_(*x*)−*S* (**ê**; **â**) (where *S* _*ν*_(*x*) is the score when *ν* = *x* and all the other parameters are set to their best fitted values) can be interpreted as a likelihood-ratio test statistics following a *χ*^2^ distribution with one degree of freedom. Lower and upper boundaries *ν*_min_ and *ν*_max_ of the 95% CI (such *S* _*ν*_(*x*) − *S* (**ê**; **â**) < 3.84 for all *ν*_min_ < *x* < *ν*_max_) may then simply be computed using a bisection method, as implemented in `poolfstat`.

Finally, a straightforward (but highly informative and recommended) approach to assess the fit of an admixture graph is to evaluate to which extent the *f* −statistics derived from the fitted admixture graph parameters (*g*(**ê**; **â**)) depart from the estimated ones (Patterson *et al*, 2012; Lipson, 2020). This can be summarized via a Z-score of residuals computed as 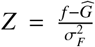 where 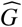 is a given fitted *f* −statistics; *f* is its corresponding estimated values; and 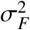 the block-jackknife standard error. The presence of outlying Z-scores for at least one *f* -statistics (e.g., | *Z* |> 1.96 at a 95% significance threshold) may suggest poor model fit while also providing insights into the leaves or graph edges that are the most problematic (Lipson, 2020).

#### 2.1.6 Scaling of branch lengths in drift units

Admixture graph fitting results in estimated edge lengths on the same scale as *F*_2_ which limits their interpretation, because they depend both on the overall level of SNP polymorphism and on their distance to the root (Patterson *et al*, 2012). Lipson *et al* (2013) proposed an empirical approach to rescale edge lengths on a diffusion timescale using estimates of overall marker heterozygosities within (i.e., 1 − *Q*_1_) or across (i.e., 1 − *Q*_2_) populations. The argument echoes the aforementioned interpretation of pairwise *F*_*ST*_ as a scaled *F*_2_. If *p*_*C*_ and *p*_*P*_ are the reference allele frequencies in a child node *C* and its direct parent node *P* and their divergence time (on a diffusion timescale) is 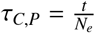 (where *t* is the branch length in generations), then conditional on *p*_*C*_, 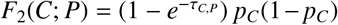 and 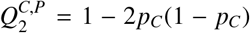 leading to 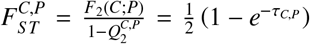 (i.e., 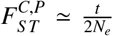 when *τ*_*C,P*_ *≪* 1). Hence, the estimated graph edges length 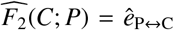 are scaled in units of drift by a factor equal to 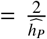 where 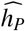 is the estimated heterozygosity (i.e., 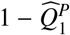) in the (parent) node *P*. Rearranging equation 2 and using 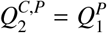 (conditional on *p*_*P*_) shows that *h*_*P*_ = *F*_2_(*C*; *P*) + *h*_*C*_, where 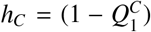 is the heterozygosity of the child node *C*. Hence, all the node heterozygosities can be inferred iteratively from the leaves to the root along the admixture graph using the leave heterozygosities (directly estimated from the data) and the fitted edge lengths (Lipson *et al*, 2013).

#### 2.1.7 Admixture Graph construction

Comprehensive exploration of the space of possible admixture graphs rapidly becomes impossible even for a moderate number of populations. We implemented in `poolfstat` different heuristics to facilitate admixture graph construction based on a supervised approach (see Supplementary Vignette V1 for details). First the `add.leaf` function allows exploring all the possible connections of a new population to an existing admixture graph. If *n*_*e*_ is the number of edges of the admixture graphs, *n*_*e*_ + 1 possible graphs connecting the new leaf with a non-admixed edges (i.e., including a new rooting with the candidate leaf as an outgroup) and 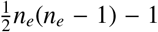 connecting the new leaf with a two-way admixture event are then tested. Note that an admixture between the two root edges is excluded from the exploration since it results in a singular model. More generally, the different possible graphs are always checked for singularity by empirically verifying that the rank of the model incidence matrix **X**(**a**) is equal to the number of edges to fit. The different fitted graph can then be ranked according to their *BIC*, the graph with the lowest *BIC* having the strongest support.

The `graph.builder` function allows a larger exploration of the graph space by successively adding several leaves in a given order to an existing admixture graph. At each step of the process, a heap stores the best resulting graph together with some intermediary sub-optimal graphs based on their *BIC*. After initializing the heap with some graph (or a list of graphs), the `add.leaf` function is called to evaluate, for each candidate leaf in turn, all its possible connections (with non-admixed or admixed edges) to all the graphs stored in the heap. Among the obtained graphs, the one with the lowest *BIC* together with those with a *BIC* within a given Δ_*BIC*_ (Δ_*BIC*_ = 6 by default) are included in a newly generated heap. If the resulting heap contains more than a predefined number of graphs 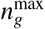 (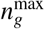 by default), only the 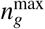 graphs with the lowest *BIC* are finally kept in the heap of graphs to be used for the addition of the next leaf. Although helpful, such heuristic should be used cautiously and we recommend to only try adding a small number of populations (i.e., ≤ 5) to an existing graph. One also needs to evaluate different orders of population inclusion (Supplementary Vignettes V1 and V2).

It is also critical to start these supervised procedures with graphs that are representative of the whole history of the populations under study and not too unbalanced with respect to the candidate leaves. In particular, starting with a small tree of closely related populations which are distantly related to the candidate leaves must be avoided. When prior knowledge about the history of the investigated population is limited (which is usually the case), Lipson *et al* (2013) proposed to start admixture graph construction with a scaffold tree of populations displaying no evidence of admixture. As in the absence of admixture, *F*_2_ statistics are expected to be additive along the paths of the (binary) population tree, its unrooted topology and branch lengths may simply be inferred with a neighbor-joining algorithm. In `poolfstat`, we implemented two functions that allow i) identifying candidate sets of unadmixed populations among all the genotyped ones (`find.tree.popset`); and ii) building rooted neighbor-joining tree (`rooted.njtree.builder`). Briefly, `find.tree.popset` implements a procedure consisting of i) discarding all the populations showing at least one significant three-population test (i.e., displaying a negative *F*_3_ Z-score lower than −1.65 by default) among all the possible ones; and ii) identifying via a greedy algorithm the largest sets of populations for which all the possible quadruplets pass the four-population test of treeness (i.e., with an absolute *F*_4_ Z-score lower than 1.96 by default). The `rooted.njtree.builder` function builds a scaffold tree from a candidate set of (presumably) unadmixed populations using the `nj` function from the `ape` package (Paradis *et al*, 2004) and then compare the consistency of population heterozygosities between the partitions of the tree to root it (Lipson *et al*, 2013). Note that this latter procedure may be sensitive to long-branch attraction and should thus be used carefully when including highly divergent populations.

### 2.2 Overview of the new poolfstat package

Tables 1 and 2 describe the main objects and functions implemented in our new version (v2.0.0) of the R package `poolfstat` publicly available from the CRAN repository (https://cran.r-project.org/web/packages/poolfstat/index.html). In-depth analyses of two Pool-Seq and allele count simulated datasets (see below) are described for illustration purposes in the package vignette provided as Supplementary Vignette V1. Detailed documentation page of the different objects and functions can also be directly accessed from an R terminal with `poolfstat` loaded using the `help` function (or the ? operator).

**Table 1.** Description of the main S4 objects of the poolfstat package. ^†^Object already existing in the first `poolfstat` version.

| S4 object | Description |
| --- | --- |
| `countdata` | Standard allele count data (i.e., obtained from individual genotyping or sequencing data) |
| `pooldata`<sup>†</sup> | Pool-Seq read count data |
| `pairwisefst` | Store pairwise $F_{ST}$ estimates. This object is generated by the `compute.pairwiseFST` function. Estimates can be conveniently visualized with the `heatmap` or `plot` functions, the latter interfacing the `plot.fstats` function of `poolfstat`. |
| `fstats` | Store $F_2$, pairwise $F_{ST}$, $F_3$, $F_3^*$, $F_4$ and $D$ estimation results. This object is generated by the `compute.fstats` function. Estimates can be conveniently visualized with the `heatmap` or `plot` functions, the latter interfacing the `plot.fstats` function of `poolfstat`. |
| `graph.params` | Represent a population tree or an admixture graph and its parameter. This object is generated by the `generate.graph.params` function. The graph can be visualized with the `plot` function that interfaces the `grViz` function from the `DiagrammeR` package (Iannone, 2020). |
| `fitted.graph` | Represent a population tree or an admixture graph and its underlying fitted parameters as obtained from the `fit.graph` or other fitting functions. The graph can be visualized with the `plot` function that interfaces the `grViz` function from the `DiagrammeR` package (Iannone, 2020). |

**Table 2.**
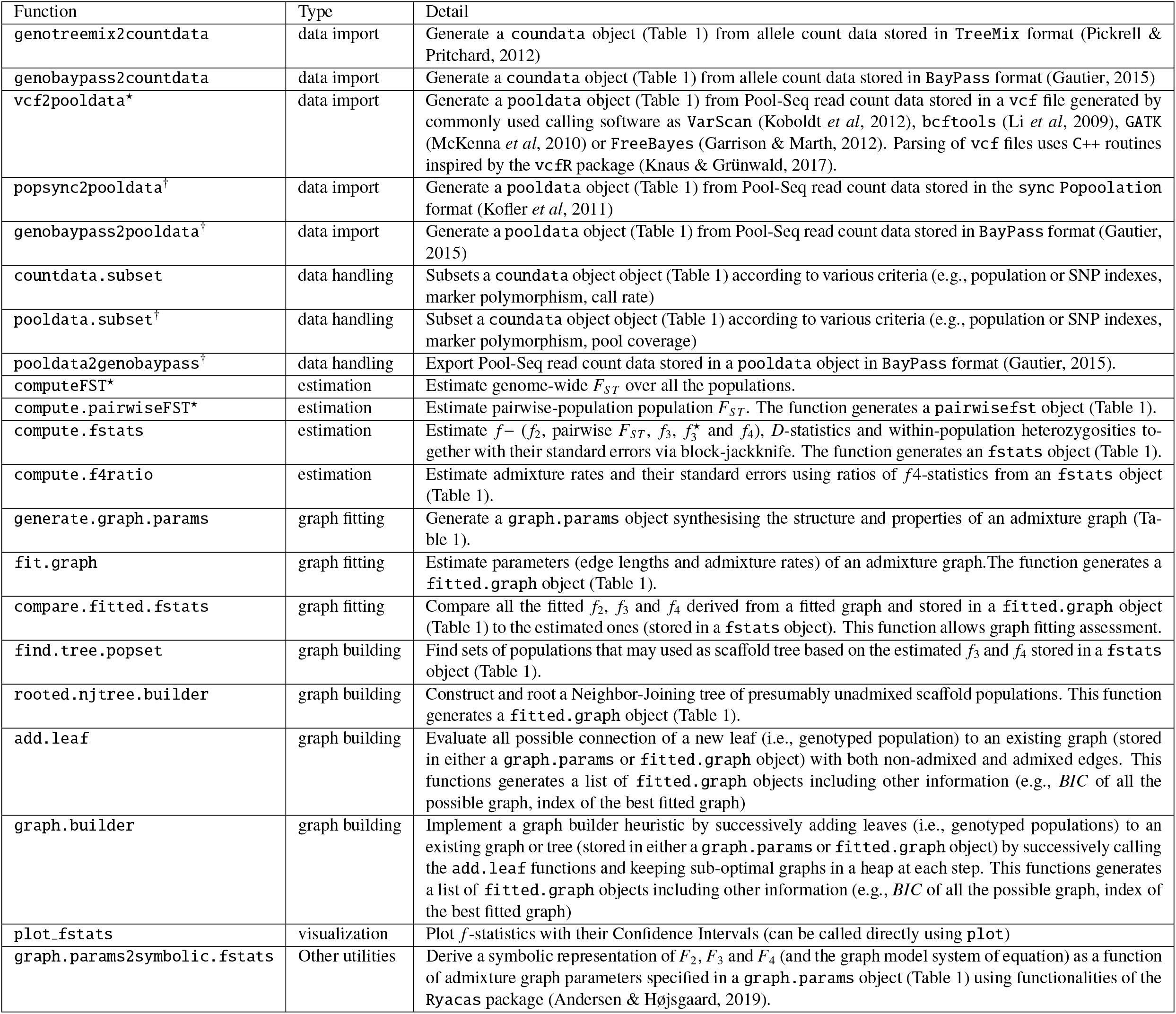
Description of the main poolfstat functions. ^†^ and ^⋆^ objects existing or significantly improved since the first `poolfstat` version, respectively.

The package includes several functions to parse allele count (e.g., `genotreemix2countdata`) or Pool-Seq (e.g., `vcf2pooldata`) input data stored in various formats commonly used in population genomics studies (Table 2). These functions allow to clearly distinguish these two different types of data by producing objects of either the so-called `countdata` (for allele count) or `pooldata` (for Pool-Seq data) classes (Table 1). This step is critical to further rely on the appropriate unbiased estimators for the *F* and *D* parameters. Some functions allow to perform subsequent manipulation of the input data, for instance to only consider some of the populations or to remove SNPs according to various criteria (Table 2).

The three functions `computeFST`, `compute.pairwiseFST` and `compute.fstats` implement the unbiased estimators of the different *f* −, *D*− and within-population heterozygosities (based on allele IIS probabilities within and between pairs of populations) together with block-jackknife estimation of their standard errors. Importantly, these three functions automatically detect the appropriate estimators given the type of data (either allele or Pool-Seq read counts) according to the input object class (either `countdata` or `pooldata`). For the estimation of *F*_*ST*_, the `computeFST` and `compute.pairwiseFST` also implement (by default) estimators based on an Analysis of Variance framework that correspond to those developed by Weir (1996) for allele count data and by Hivert *et al* (2018) for Pool-Seq data.

The `fit.graph` function implements the approach described above to estimate the parameters (i.e., edge lengths and admixture rates) of an admixture graph that is stored in a `graph.params` object (Table 1). Such objects can be generated with the `generate.graph.params` function (Table 2) to include the target basis *f* −statistics and the error covariance matrix (denoted above 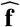 and **Q**, respectively) estimated with `compute.fstats` (stored in an `fstats` object) and to specify the topology and the parameters of the admixture graph. Note that the `graph.params2symbolic.fstats` function allows exploring in details the properties of an admixture graph specified by a `graph.params` object by deriving a symbolic representation of all the *F*_2_, *F*_3_, *F*_4_ and the model equations (see above) by internally relying on the `Ryacas` package for symbolic computation (Andersen & Højsgaard, 2019). The `fit.graph` function then produced an object of class `fitted.graph` that includes the estimated edge lengths (in *F*_2_ and also optionally in drift units) and admixture proportions together with (optionally) their 95% CI. For model fit assessment purposes, `fitted.graph` objects also include the *BIC* and Z-score of the residuals of the fitted basis *f* −statistics. Such a comparison can (and should) be generalized to all the *f*_2_, *f*_3_ and *f*_4_ statistics (not just the ones forming the basis) using `compare.fitted.fstats` jointly applied to a `fitted.graph` and a `fstats` objects. Notice that we developed for comparison purposes a function named `graph.params2qpGraphFiles` to export admixture graph specification and their underlying estimated basis *f* −statistics (both stored in a `graph.params` object) into `qpGraph` format (Patterson *et al*, 2012), allowing independent fitting based on the same estimated statistics to be carried out with this later program.

The `poolfstat` package includes several functions to assist construction of admixture graphs. As mentioned in the previous section, the `find.tree.popset` and `rooted.njtree.builder` functions allow to identify and build rooted tree(s) of scaffold of (presumably) unadmixed populations that may be used as starting graph(s). Besides, the `add.leaf` and `graph.builder` functions implement the above described heuristic to extent an existing graph (or tree) by adding one or several leaves (i.e., genotyped populations). These functions generate a list of `fitted.graph` objects together with other information that may be helpful for graph comparison (e.g., *BIC* of all the graphs or index of the best fitted graph).

Finally, as detailed and exemplified in the Supplementary Vignette V1, fitted graphs (stored in `fitted.graph` objects) and non-fitted graphs (stored in `graph.params` objects) can be directly and conveniently plotted with the `plot` function which internally interfaces the `grViz` function from the `DiagrammeR` package (Iannone, 2020).

### 2.3 Data analyses

#### 2.3.1 Simulation study

Genetic data for a total of 150 diploid individuals belonging to six different populations (n=25 individuals per populations) related by the demographic scenario depicted in Figure 1 were simulated using the `msprime` coalescent simulator (Kelleher *et al*, 2016) with the following command:

**Figure 1.**
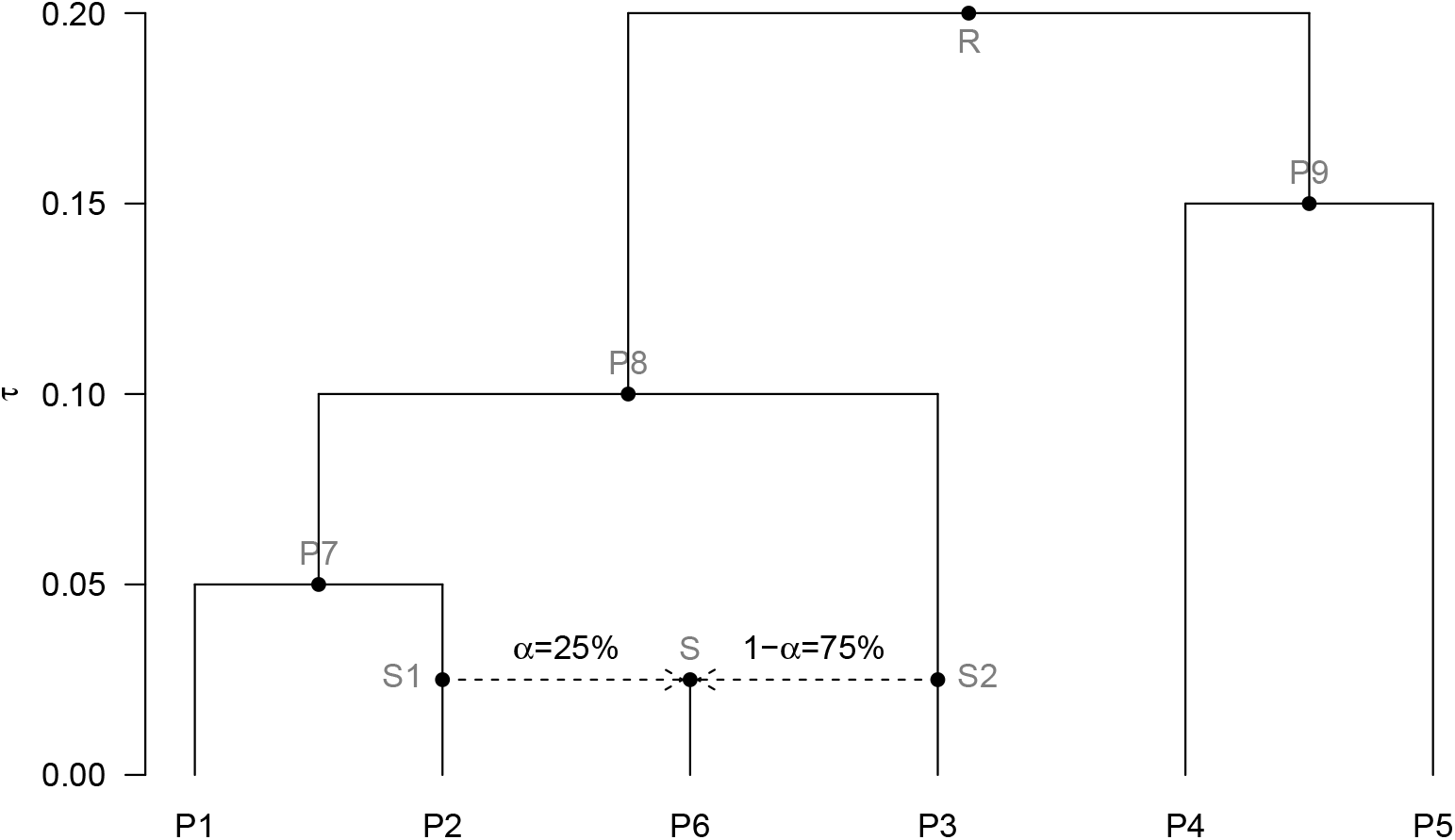
Simulated scenario relating six sampled populations. The population *P*6 derived from a population *S* which is admixed between two ancestral sources (*S* 1 and *S* 2) directly related to populations *P*2 and *P*3 and contributing to *α* = 25% and 1 − *α* = 75% of its genome, respectively. The branch lengths are in a diffusion timescale i.e., with 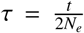 under a pure-drift model of divergence (where *t* is the number of non-overlapping generations and *N*_*e*_ the average diploid effective population sizes along the branch). The names of the internal node populations (not sampled) are represented in grey.

~~~
mspms 300 20 -t 4000 -I 6 50 50 50 50 50 50 0 -r 4000 100000000 -p 8 -es 0.0125 6 0.25
-ej 0.0125 6 2 -ej 0.0125 7 3 -ej 0.025 2 1 -ej 0.05 3 1 -ej 0.075 5 4 -ej 0.1 4 1
~~~

Each genome thus consisted of 20 independent chromosomes of *L* = 100 Mb assuming a scaled chromosome-wide recombination rate of *ρ* = 4*LN*_*e*_*r* = 4, 000 as expected for instance in a population of constant diploid effective size of *N*_*e*_ = 10^3^ when the per-base and per-generation recombination rate is *r* = 10^−8^ (i.e., one cM per Mb). The scaled chromosome-wide mutation rate was set to *θ* = 4*LN*_*e*_*µ* = 4, 000 which is also the expected nucleotide diversity in a population with *N*_*e*_ = 10^3^ at mutation-drift equilibrium when the per-base mutation rate is *µ* = 10^−8^. A total of 250 independent genotyping datasets were simulated and each was subsequently processed to generate 32 different types of datasets corresponding to:

- Two standard allele count datasets (namely AC_*m*≥1%_ and AC_*m*≥5%_) obtained by simple counting of the simulated individual (haploid) genotypes for each population (i.e., assuming Hardy-Weinberg equilibrium within population) and removing SNPs with a Minor Allele Frequency (MAF) computed over all the individuals lower than 1% (for AC_*m*>1%_ datasets) or 5% (for AC_*m*>5%_ datasets)
- Thirty Pool-Seq datasets (coded as 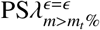) for i) five different average sequencing coverages *λ* (equal to 30, 50, 75, 100 or 200 reads; a 30X Pool-Seq coverage representing a lower limit for population genomics studies); ii) two different MAF thresholds *m*_*t*_ of 1% and 5% (MAF being estimated on the read counts over all the pools); and iii) three different sequencing error rates *E* of 0 (no error), 1‰ and 2.5‰ the two latter being representative of Illumina sequencers (Glenn, 2011).

Pool-Seq datasets were simulated from the AC_*m*≥1%_ allele count datasets following a procedure similar to that described in Hivert *et al* (2018). Briefly, the vector ***r***_*ij*_ = {*r*_*ijk*_} of read counts at SNP position *i* in population *j* for the nucleotide *k* (where by convention *k* = 1 and *k* = 2 for the derived and ancestral alleles respectively) was sampled from a Multinomial distribution parameterized as:

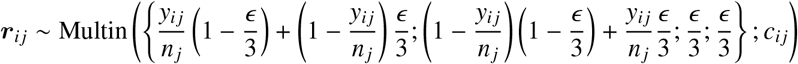

where *y*_*ij*_ is the derived allele count for SNP *i* in population *j* (from the corresponding AC_*m*>1%_ dataset); *n*_*j*_ is the haploid sample size of population *j* (here *n*_*j*_ = 50 for all *j*); and 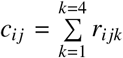 is the overall read coverage. To introduce variation in read coverages across pools and SNPs, each *c*_*ij*_ was sampled from a Poisson distribution with a parameter *λ* (the target Pool-Seq mean coverage). When *ϵ* = 0, only reads for the derived (*k* = 1) or ancestral (*k* = 2) alleles can be generated and the above Multinomial sampling actually reduces to a Binomial sampling following 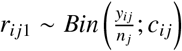 (and *r*_*ij*2_ = *c*_*ij*_ − *r*_*ij*1_). However, when *ϵ* > 0, sequencing errors might lead to non-null read counts for the two other alleles leading to tri- or tetra-allelic SNPs. Morevover, sequencing errors may also introduce spurious additional variation by generating false SNPs at monomorphic sites. To account for the latter, read count vectors ***r***_*i′ j*_ for all the 2 × 10^9^ − *I* monomorphic positions *i′* (where *I* is the number of SNPs observed in the considered AC_*m*>1%_ dataset) were sampled as 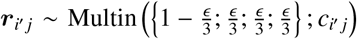 with coverages *c*_*i′ j*_ sampled from a Poisson distribution (as *c*_*ij*_ for polymorphic positions). Yet, as usually done with empirical datasets, we applied a minimum read count filtering step consisting of disregarding all the alleles with less than 2 observed reads (over all the populations). Only bi-allelic SNPs passing the overall MAF threshold *m*_*f*_ were finally retained in the final 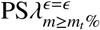 datasets.

Analyses of the simulated data were carried out with `poolfstat` (Supplementary Vignette V1). Briefly, each `msms` simulated dataset was converted into an AC_*m*≥1%_ dataset in `TreeMix` format (Pickrell & Pritchard, 2012) further imported into a `countdata` object with `genotreemix2countdata` (Tables 1 and 2) and used to generate each corresponding AC_*m*≥5%_ dataset using `countdata.subset`. To improve computational efficiency, the different 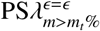 Pool-Seq datasets were generated from the AC_*m*≥1%_ `countdata` objects in the form of `pooldata` object using custom functions (not included in the package) coded in C++ and integrated within `R` using `Rcpp` (Eddelbuettel, 2013). In addition, to evaluate the impact of the (bad) practice consisting of analyzing Pool-Seq data as if they were allele count data (i.e., overlooking the sampling of reads from individual genes of the pool), we also created “fake” `countdata` objects from the different `pooldata` objects. We then used default options (unless otherwise stated) of i) `computeFST` to estimate genome-wide *F*_*ST*_ over all the populations; ii) `compute.fstats` to estimate all the *f* − and *D*− statistics; iii) `compute.f4ratio` to estimate admixture proportions; and iv) `fit.graph` to estimate the admixture graph parameters (Table 2). As the number of SNPs was variable across the different simulated datasets, we adjusted the number of successive SNPs defining a block for block-jackknife estimation of standard errors by dividing the total number of available SNPs by 500. This thus resulted on average in 490 blocks of 4.1 Mb over the genome for all the analyzed datasets (the simulated genomes consisting of 20 chromosomes). Note that the parameter estimates were always taken as the block-jackknife mean values rather than estimates over all SNPs (i.e., including those in the chromosome ends). In practice, the differences between the two are insignificant (e.g., Supplementary Vignette V1).

For validation purposes, we also analyzed the 250 AC_*m*≥1%_ datasets with programs from the `AdmixTools` suite (Patterson *et al*, 2012) after conversion to the appropriate input format using custom `awk` scripts. More specifically, we ran `qpfstats` (v. 200) to estimate the 15 basis *f* −statistics, i.e., taking *P*1 as the reference population, the five *f*_2_ of the form (P1,Px) and the ten *f*_3_ of the form (P1;Px,Py) (where *x* = 2, …, 6 and *y* = 3, …, 6 with *y* ≠ *x*) and their corresponding error covariance matrix. Default options were considered except for the disabling of the scaling of estimated values (using option `-l 1`) to facilitate their comparison with `poolfstat` estimates. We also ran with default options `qp3Pop` (v. 650) to estimate 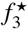 for all the 60 possible triplet configurations and `qpDstat` (v. 970) to estimate the *D*−statistics for all the 45 possible quadruplet configurations together with their associated *Z*−scores. By default, these three programs define blocks of 5 cM to implement the (weighted) block-jackknife procedure. As we here converted the simulated SNP positions from Mb to cM assuming one cM per Mb (see above), the sizes of the 400 blocks was thus about 20% than for `poolfstat` analyses.

#### 2.3.2 Analysis of a real *Drosophila suzukii* Pool-Seq data

The spotted wing drosophila, *Drosophila suzukii*, represents an attractive model to study biological invasion and hence recent historical and demographic history. Native to South East Asia, this pest species was first observed outside its native range in Hawaii in 1980, and later rapidly invaded America and Europe simultaneously between 2008 and 2013 (Fraimout *et al*, 2017). Using DNA sequences and microsatellite markers, Adrion *et al* (2014) and Fraimout *et al* (2017) deciphered the routes taken by *D. suzukii* during its worldwide invasion. Both studies showed that America and European populations globally represent separate invasion routes with different native source populations. Olazcuaga *et al* (2020) recently generated Pool-Seq genomic data from 22 worldwide population samples to detect genetic variants associated with the historical status (i.e. invasive versus native) of the sampled populations. We here focused our illustration on 14 Pool-Seq data from this study (with 50 to 100 diploïd individuals per pool) for populations representative of the Asian native area (six populations), Hawaii (one population) and the invaded continental America (seven populations), where the species was first observed in 2008 on the Western coast of the USA (around Watsonville, CA; Figure 2A). Beside native populations, we have restricted our analysis to the American continent because the invasion of this area is characterized by multiple admixture events between different source populations (Fraimout *et al*, 2017), which makes it an appealing situation to evaluate the power and the limitation of `poolfstat` analyzes. Moreover, 13 of our 14 population samples consist of individuals originating from the same sites (albeit sometimes collected at different dates for some pools; Table 2 in Supplementary vignette V2) as those genotyped at 25 microsatellite markers and analyzed with an Approximate Bayesian Computation Random Forest (ABC-RF) approach to infer the routes of invasion on a worldwide scale by Fraimout *et al* (2017).

**Figure 2.**
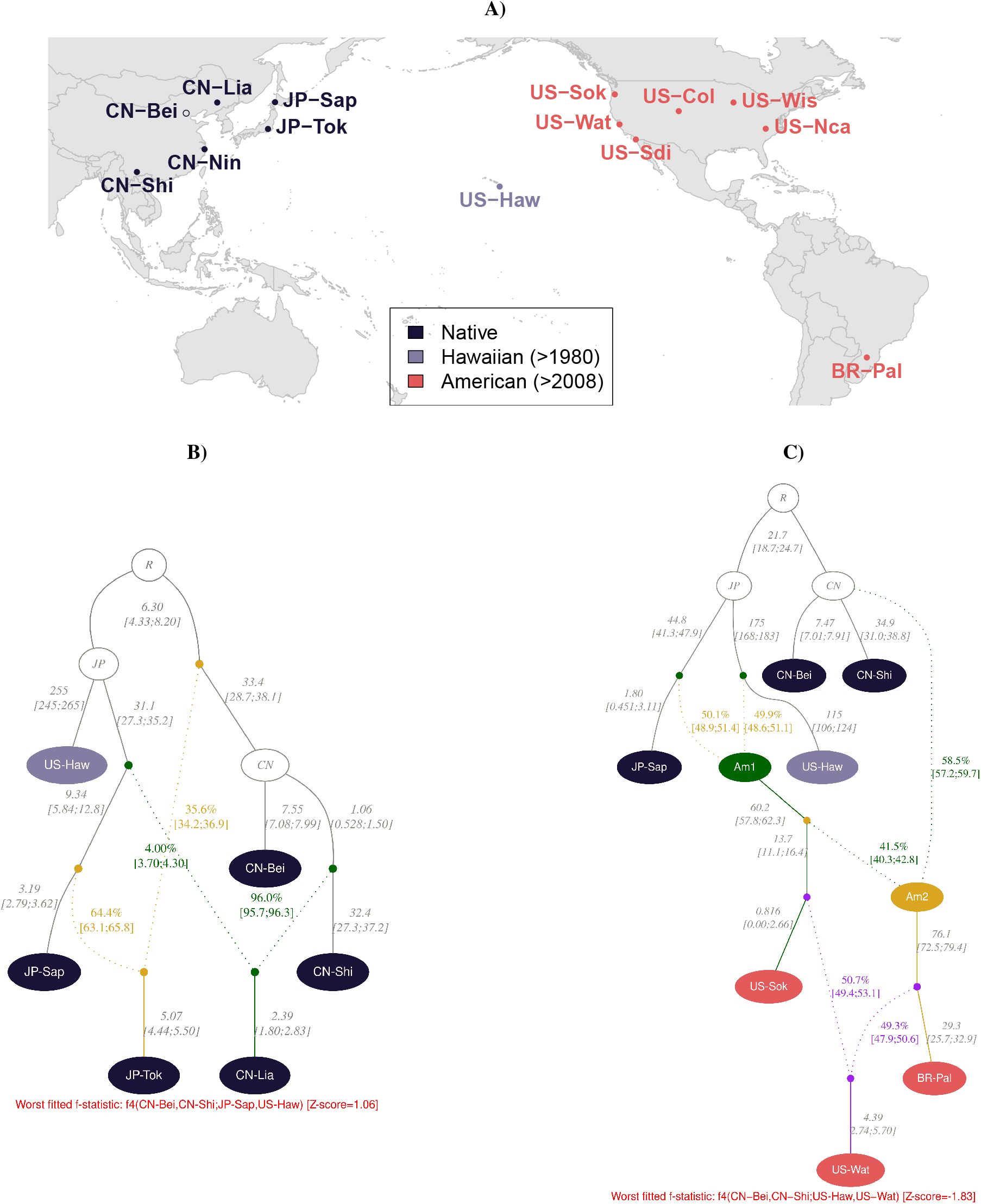
Historical and demographic inferences about native and invasive *Drosophila suzukii* populations from Pool-Seq data based on *f* − statistics. (A) Geographic location of the 14 population samples (Olazcuaga *et al*, 2020). Names are colored according to their area of origin. The (invasive) Hawaiian population, which was considered as intermediate between the Asian native and the continental America invasive area, was first observed in 1980, i.e. ca. 300 generations before the invasion of the American continent assuming 10 generations per year. Solid points indicate the 13 population sampling sites in common with Fraimout *et al* (2017). B) Best fitting admixture graph connecting five populations of the native areas and the Hawaiian population with two inferred admixture events. C) Best fitting admixture graph connecting three invasive populations from continental America with populations from the native area (and Hawaii). In B) and C), estimates of branch lengths (×10^3^, in drift units of 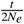) and admixture rates (and their 95% CI into bracket) are indicated next to the corresponding edges. The worst fitted *f*-statistics is written in red for each of the two graphs.

To allow for complete reproduction (and exploration) of our analyses, all the command lines used to analyze the *D. suzukii* Pool-Seq dataset are described in the Supplementary vignette V2. Briefly, we combined the 14 (`bam`) files, obtained by Olazcuaga *et al* (2020) after aligning the 14 Pool-Seq data onto the latest near-chromosome scale *D. suzukii* assembly (Paris *et al*, 2020), into an `mpileup` file using `SAMtools` 1.9 with options `-q 20 -Q20` (Li *et al*, 2009). Variant calling was then performed using `VarScan mpileup2snp` v2.3.4 Koboldt *et al* (2012) run with options `--min-coverage 10 --min-avg-qual 25 --min-var-freq 0.005 --p-value 0.5` (i.e., with very loose criteria). After discarding positions mapping to non-autosomal contigs (Paris *et al*, 2020), the resulting `vcf` file was parsed with the *vcf2pooldata* function of `poolfstat` with default options except for i) the overall MAF threshold (computed from read counts) that was set to 5%; and ii) the minimal read coverage for each pool that was set to 50. The resulting `pooldata` object was further filtered with `pooldata.subset` to discard i) all positions with a coverage higher than the 99th coverage percentile within at least one pool; and ii) discard all SNPs with MAF<5% over all the populations from the native area to favor ancestral SNPs. The final dataset then consisted of read counts for 1,588,569 bi-allelic SNPs with a median read coverage varying from 64 (US-Sok) to 95 (CN-Bei and US-Haw) among the 14 pools (Table 2 in the Supplementary Vignette V2). We defined blocks of 10,000 consecutive SNPs for block-jackknife estimation of standard errors leadint to a total of 145 blocks of 698 kb on average (varying from 414 kb to 2.03 Mb). Hence, most analyses actually relied on 1,450,000 SNPs that mapped to the 15 largest contigs of the assembly (totaling 116 Mb). In other words, SNPs mapping to the smallest (and less reliable) contigs were discarded in addition to the few ones mapping to the end of the 15 retained contigs.

## 3 Results

### 3.1 Evaluation of poolfstat on simulated data

Historical and demographic inference based on *f* − and *D*− statistics has already been extensively evaluated in previous studies (e.g., Patterson *et al*, 2012; Lipson *et al*, 2013; Peter, 2016). Therefore, the purpose of our simulation study was essentially threefold: i) to validate the estimators implemented in `poolfstat` by comparing, for allele count data, with those obtained with the reference `AdmixTools` suite (Patterson *et al*, 2012); ii) to evaluate the performance of the estimators for Pool-Seq data as a function of read coverage and sequencing errors; and iii) to provide example datasets with known ground truth for illustration purposes.

#### 3.1.1 Description of the simulated datasets

We simulated 250 genetic datasets for six populations (named *P*1 to *P*6) each consisting of 25 diploid individuals and that were historically related by the admixture graph represented in Figure 1 (Material and Methods). Each of these datasets was further used as template to generate 2 allele count datasets (applying 1% or 5% threshold on the overall MAF for AC_*m*>1%_ and AC_*m*>5%_ datasets respectively) and to simulate 30 Pool-Seq datasets with five different mean read coverages (*λ* ∈ {30; 50; 75; 100; 200}); three sequencing error rates (*ϵ* ∈ {0; 10^−3^; 2.5 × 10^−3^}) and two MAF (computed over all read counts) thresholds (referred to as 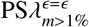and 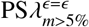for 1% and 5% MAF thresholds, respectively). This thus lead to a total of 8,000 simulated datasets. The average number of available SNPs and false SNPs (for 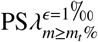and 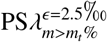 datasets) is given in Table S1 for each of the 32 different types of datasets and represented as a function of the mean coverages *λ* and MAF thresholds in Figure S1.

Overall, 471,919 SNPs and 240,369 SNPs were available on average for allele count datasets at the 1% (AC_*m*>1%_) and 5% (AC_*m*>5%_) MAF thresholds respectively consistent with the L-shaped distribution of allele frequencies (Figure S2A). As expected from binomial sampling (Figure S2B), for Pool-Seq datasets generated with no sequencing error, the number of SNPs remained always lower than the AC_*m*>1%_ datasets at the 1% MAF threshold although increasing with coverages from 13.8% for 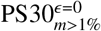 to 2.01% for PS200*m* > 1%^*ϵ*=0^ datasets (see Table S1 legend for details). Conversely, at the 5% MAF threshold, the number of SNPs was slightly higher than the AC_*m*>5%_ datasets (from 2.58% for 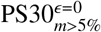to 1.51% for 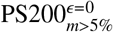) which is related to i) the shape of the allele frequency spectrum (stochastic variation in read sampling leading to include more SNPs with 0.01 < *MAF* < 0.05 than exclude SNPs with *MAF* > 0.05 from the simulated genotying data because they are more numerous); and ii) variation in the simulated read coverages that explains the decreasing trend with *λ*.

With sequencing errors, our filtering steps proved efficient to remove false SNPs except at the 1% MAF threshold when *ϵ* = 2.5‰ or when *ϵ* = 1‰ at the lowest coverage (*λ* = 30 and *λ* = 50). These configurations displayed substantial to very high proportions of false SNPs (up to 93.8% for 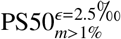) although decreasing with coverage (Figure S1B). A 5% MAF threshold always resulted in the complete removal of all the false SNPs for all the investigated scenarios (Table S1). Note that for the highest coverages, sequencing errors lead to a relative reduction of the number of SNPs due to the generation of spurious tri- or tetra-allelic SNPs from the simulated bi-allelic SNPs (compare e.g., 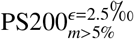 and 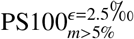 on Figure S1A).

#### 3.1.2 Comparison of poolfstat and Admixtools estimates for allele count data

We first analyzed the 250 simulated AC_*m*>1%_ datasets to estimate with both `poolfstat` and `Admixtools` programs i) the 15 basis *f* −statistics (taking *P*1 as the reference population) consisting of five *f*_2_ and ten *f*_3_ (Figure 3A) and their corresponding error covariance matrix (Figure 3B); ii) the 60 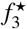 (Figure 3C) and their associated Z-scores (Figure 3D); and iii) the 45 *D*−statistics (Figure 3E) and their associated Z-scores (Figure 3F). The estimates were all found in almost perfect agreement between the two implementations with Mean Absolute Differences (MAD) negligible when compared to the range of variation of the underlying values. For *f* − and *D*− statistics, slight differences were mostly due to the plotted `poolfstat` estimates corresponded to block-jackknife means (i.e., excluding SNPs outside blocks as those from chromosome ends). Using `poolfstat` estimates based on all the SNPs indeed resulted in almost null MAD (MAD’ in Figure 3A, C and E), up to rounding errors due to lower decimal precision in the printed output of the `Admixtools` programs. Note that the differences in block-jackknife implementation among the two programs (Material and Methods) had very minor impact on the estimation of error variance and covariance of the estimates (Figures 3B). Accordingly, the MAD computed on Z-scores remained very small (although inflated for higher values) and Z-score based decision for the underlying three-population admixture (Figures 3D) or four-population treeness tests (Figures 3F) were highly consistent (with a proportion *β* = 97.7% and *β* = 98.0% respectively of Z-scores significant with the two programs among the ones significant with at least one program).

**Figure 3.**
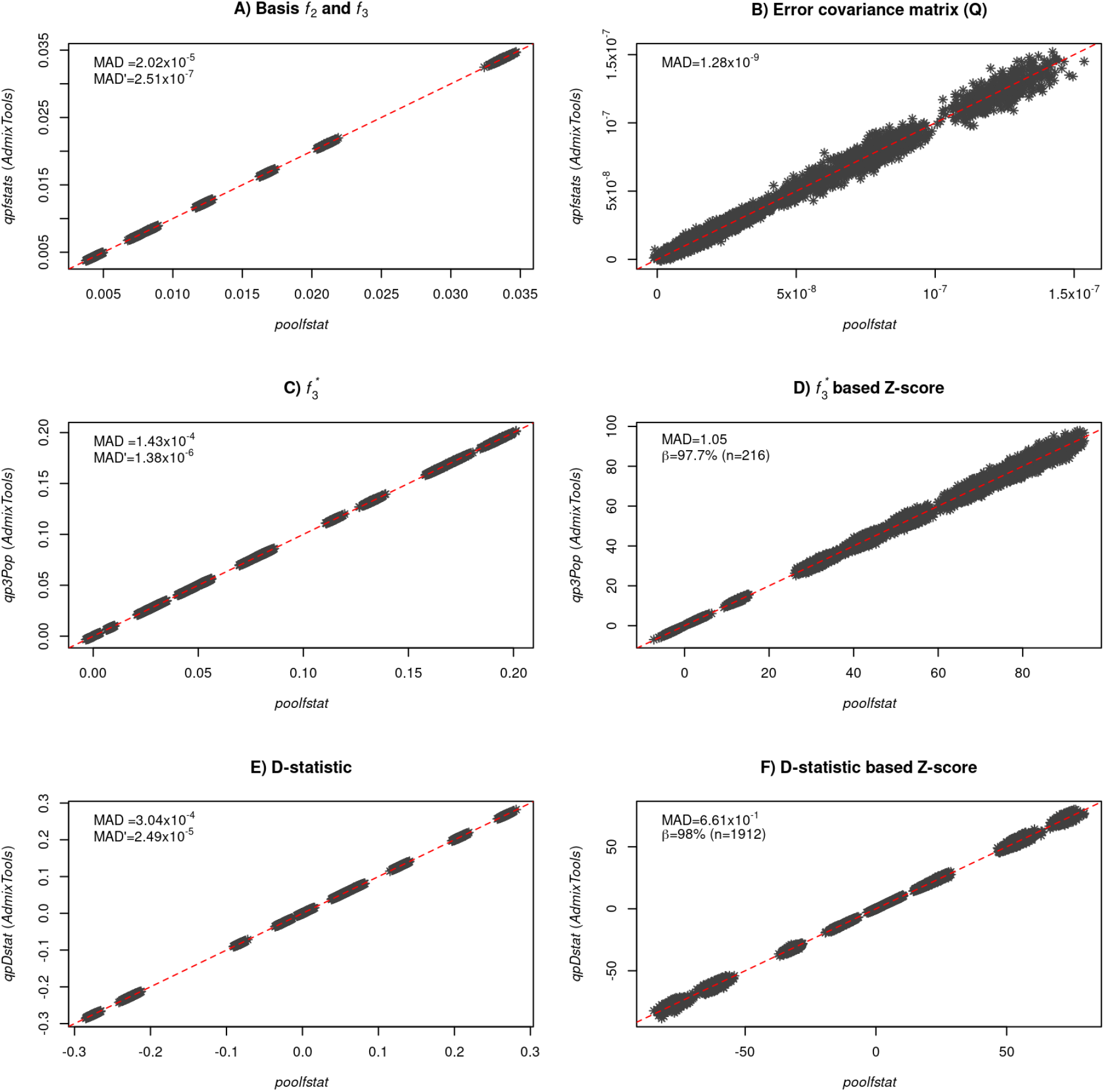
Comparison of poolfstat and AdmixTools estimates across 250 simulated allele count datasets (AC_*m*≥1%_). A) All estimates of the 15 basis *f* − statistics taking *P*1 as the reference population and corresponding to 5 *f*_2_ of the form (*P*1,*Px*) and the 10 *f*_3_ of the form (*P*1;*Px,Py*) (with *x* = 2, .., 6; *y* = 3, .., 6 and *y* > *x*). B) All Block-jackknife estimates of the covariance matrix ***Q*** of the 15 basis *f* − statistics (15 error variances and 105 error covariances). C) All estimates of the 60 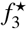 (scaled *f*_3_) and their associated Z-scores (D). E) All estimates of the 45 *D* − statistics (scaled *f*_4_) and their associated Z-scores (F). For each comparison, the Mean Absolute Difference (MAD) between the parameter estimates of the two programs are given on the upper left corner of the plots. In A), C) and E), `poolfstat` estimates correspond to block-jackknife means (i.e., they only include SNPs eligible for block-jackknife). The given MAD’ value is the MAD between `AdmixTools` and `poolfstat` estimates that include all SNPs (see documentation for the `compute.fstats` function). In D), a consistency score *β* is also given and was computed as the proportion of Z-scores < − 1.65 (i.e., significant three-population test of admixture at a 5% threshold) with both programs among the *n* = 216 ones significant in at least one of the two programs. Similarly, in F), the given consistency score *β* is computed as the proportion of absolute Z-scores < 1.96 (i.e., passing the four-population treeness at a 5% threshold) with both programs among the *n* = 1, 912 ones with an absolute Z-scores < 1.96 in at least one of the two programs)

#### 3.1.3 Performance of *f*_3_ and 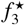 based tests of admixture and *f*_4_ and *D*-based tests of treeness for allele count and Pool-Seq data

We ran the `compute.fstats` function on all the simulated allele count and Pool-Seq datasets to estimate all *f* - and *D*-statistics. To further evaluate the impact of (improperly) treating read counts as allele counts when analyzing Pool-Seq data we also analyzed the simulated Pool-Seq datasets (focusing only on 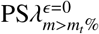 datasets, i.e., simulated without sequencing error) as if they were allele count data. Overall, 42 different configurations were thus investigated each originating from the 250 allele count datasets simulated under the demographic scenario represented in Figure 1, leading to a total of 42×250=10,500 analyses.

Tables 3 and S2 provide the estimated power (True Positive Rate, TPR) and False Positive Rate (FPR) of the *f*_3_- and 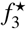-based test of admixture for each configurations. As P6 was the only admixed population, each TPR was estimated as the proportion of *f*_3_ (respectively 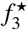) with an associated Z-score < −1.65 (95% significance threshold) for the (P6;P2,P3)population triplet (i.e., among 250 estimates). Conversely, the FPR was computed as the proportion of *f*_3_ (respectively 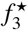) with an associated Z-score < −1.65 among all the 50 population triplets that do not involve P6 as a target (i.e., among 12,250=250×50 estimates). Consistent with Patterson *et al* (2012), the performance of *f*_3_- and 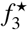-based test of admixture were virtually the same for all the configurations. When the same MAF threshold was applied, the performance of Pool-Seq data generated with no sequencing error were very close to that obtained with allele count data although the power tended to slightly decrease with decreasing sequencing coverage. Interestingly, increasing the MAF threshold from 1% to 5% increased the power by more than 10% and in all cases, no false positive signal of admixture was detected. Surprisingly, sequencing errors in Pool-Seq data also tended to increase the power from a negligible amount (less or close to 1%) at 5% MAF threshold to a quite substantial amount at 1% MAF threshold (decreasing with coverage and increasing with sequencing error rate). At the extreme, a power of 100% was even observed when *λ* ≤ 50 and *ϵ* ≥ 1‰. This trend was actually directly related to the proportion of false SNPs introduced by sequencing error (Figure S1B) that resulted in a downward bias of *f*_3_ and 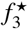 estimates, although the underlying tests remained robust as all the estimated FPR were null except for 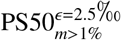 data (FPR=6.47%) which displayed the highest proportion of false SNPs (> 90%, Figure S1B). However, this observed apparent robustness of the three-population tests to false SNPs should be interpreted cautiously since it may rather result from the moderate to high expected *f*_3_ and 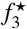 values in our simulated scenario for the population triplets that do not involve P6 as a target. Overall, applying a 5% MAF threshold on Pool-Seq data (even with *ϵ*=2.5‰) to remove false SNPs (see above) allowed recovering the performances similar to that obtained when analyzing datasets with no sequencing error. Finally, it is worth stressing that analyzing Pool-Seq data as allele counts, whatever the coverage or MAF threshold considered, lead to no power in detecting admixture event with *f*_3_ or 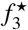 based tests due to a strong upward estimation bias.

**Table 3.**
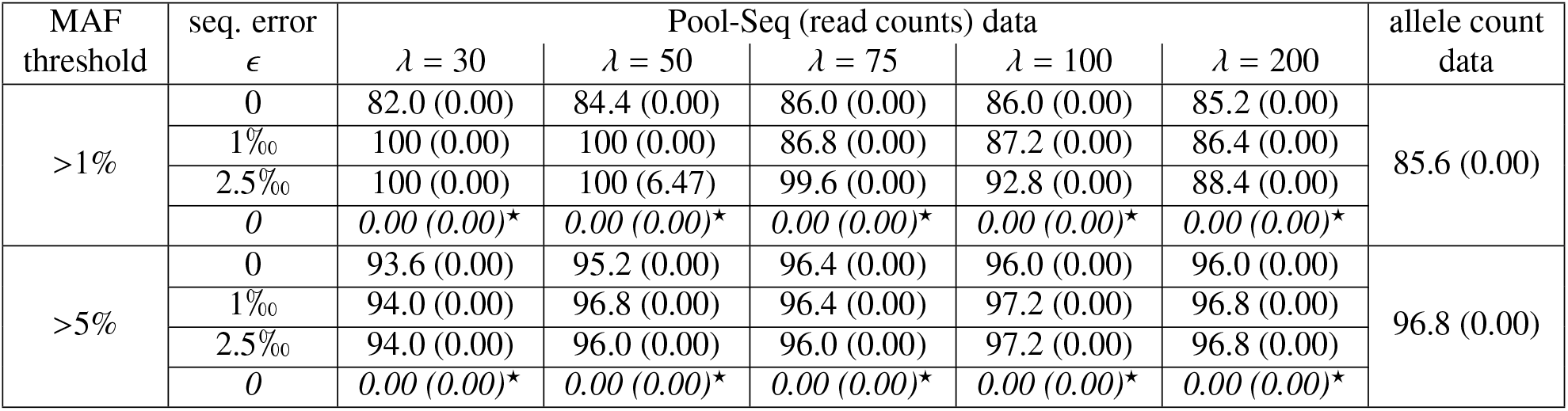
Comparison of the performance of *f*_3_-based tests of admixture for different types of data simulated under the Figure 1 scenario processing poolfstat analyses. For each MAF threshold (MAF> 1% or MAF> 5%), the table gives True and False (in parenthesis) Positive Rates (in %) for 21 different types of analyses relying on i) allele count data; ii) 15 different Pool-Seq read count data (five mean coverages *λ* and three sequencing error rates *ϵ*); and iii) Pool-Seq read count data simulated with *ϵ* = 0 treated as allele counts (corresponding results of this bad practice are highlighted in italics and ^⋆^). Each TPR was computed from the analysis of 250 independent datasets (generated from the data simulated under Figure 1 demographic scenario) as the proportion of *f*_3_ with an associated Z-score < − 1.65 (95% significance threshold) for the (P6;P2,P3) population triplet (n=250 estimates). The FPR was similarly computed as the proportion of *f*_3_ with an associated Z-score< −1.65 among all the 50 population triplets that do not involve P6 as target population (n=250×50=12,250 estimates).

Tables 4 and S3 similarly provide the estimated power (TPR) and FPR of the *f*_4_- and *D*-based tests of treeness for the 42 configurations investigated in the simulation study. Given the simulated scenario, eight of the 45 different population quadruplets (namely (P1,P2;P3,P4); (P1,P2;P3,P5); (P1,P2;P4,P5); (P1,P3;P4,P5); (P1,P6;P4,P5); (P2,P3;P4,P5); (P2,P6;P4,P5); and (P3,P6;P4,P5)) have a null expected *F*_4_ (and *D*) value. Note that this may easily be shown with the symbolic calculus derivation implemented in `graph.params2symbolic.fstats` (Table 2). For each configuration, the TPR of the treeness test was then estimated as the proportion of *f*_4_ (respectively *D*) with an associated absolute Z-score < 1.96 (95% significance threshold) for these eight population quadruplets ((P1,P2;P3,P4); (P1,P2;P3,P5); (P1,P2;P4,P5); (P1,P3;P4,P5); (P1,P6;P4,P5); (P2,P3;P4,P5); (P2,P6;P4,P5); (P3,P6;P4,P5)) over all the 250 different underlying analyses (i.e., among 2,000=250×8 estimates). Conversely, the FPR was estimated as the proportion of *f*_4_ (respectively *D*) with an associated absolute Z-score < 1.96 among all the 37 remaining population quadruplets (i.e., among 9,250=250×37 estimates). The power for both *F*_4_- and *D*-based tests were remarkably consistent across all the different configurations. In addition, the tests were all found almost perfectly calibrated since the estimated power were close to 95%, the probability of rejecting the null hypothesis at the chosen 95% significance threshold for Z-scores. Likewise, all FPR remained low (≤ 0.15%), although increasing with MAF thresholds (more than twice higher for a given type of data when increasing the MAF threshold from 1% to 5%). Overall, sequencing errors and coverage had no impact on the performance of the *f*_4_- and *D*-based test of treeness. As expected, analyzing read counts as allele count data did not affect the performance of these tests (see Discussion).

**Table 4.**
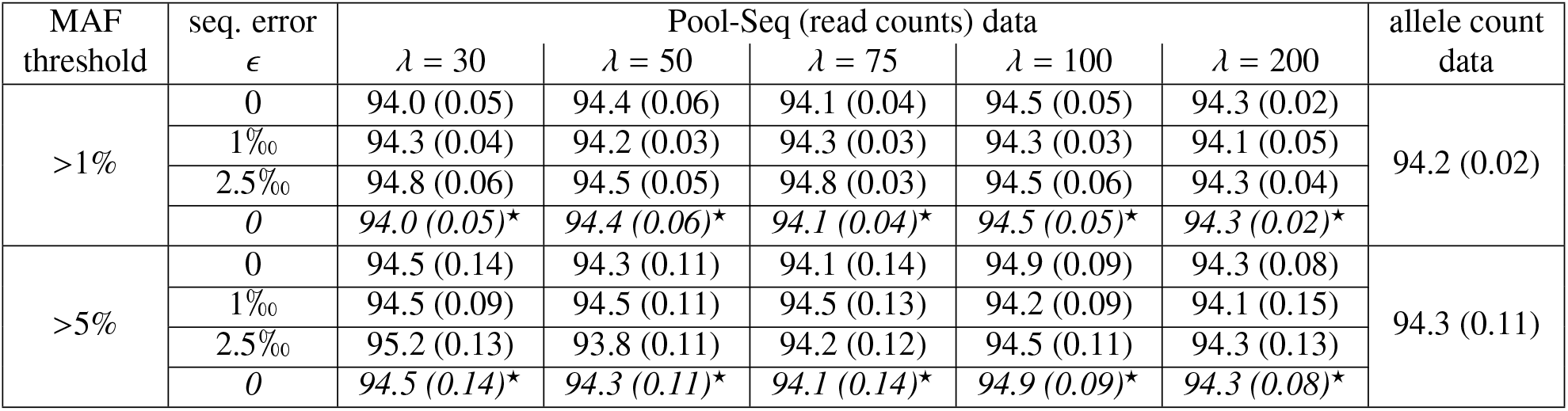
Comparison of the performance of *f*_4_-based test of treeness for different types of data simulated under the Figure 1 scenario processing poolfstat analyses. For each MAF threshold (MAF> 1% or MAF> 5%), the table gives True and False (in parenthesis) Positive Rates (in %) for 21 different types of analyses relying on i) allele count data; ii) 15 different Pool-Seq read count data (five mean coverages *λ* and three sequencing error rates *ϵ*); and iii) Pool-Seq read count data simulated with *ϵ* = 0 treated as allele counts (corresponding results of this bad practice are highlighted in italics and ^⋆^). Each TPR was computed from the analysis of 250 independent datasets (generated from the data simulated under Figure 1 demographic scenario) as the proportion of *f*_4_ with an associated absolute Z-score < 1.96 (95% significance threshold) among all the eight population quadruplets ((P1,P2;P3,P4); (P1,P2;P3,P5); (P1,P2;P4,P5); (P1,P3;P4,P5); (P1,P6;P4,P5); (P2,P3;P4,P5); (P2,P6;P4,P5); (P3,P6;P4,P5)) with a null expected *F*_4_ (n=250 × 8=2,000 estimates). The FPR was similarly computed as the proportion of *f*_4_ with an associated absolute Z-score < 1.96 among all the 37 remaining population quadruplets (n=250×37=9,250 estimates).

#### 3.1.4 Precision of the *F*_4_-ratio based estimation of the admixture rate *α*

Given the simulated scenario, two different ratios of *f*_4_ estimates could be used to estimate the admixture proportion *α* = 0.25 (Figure 1), namely 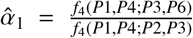 and 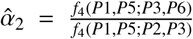 (Patterson *et al*, 2012). The `graph.params2symbolic.fstats` function (Table 2) may also prove useful to identify appropriate quadruplets (Supplementary Vignette V2). We used the `compute.f4ratio` function to obtain these two estimates from all the simulated datasets together with their 95% CI (defined as 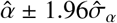 where 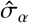 is the block-jackknife standard-error estimate). Tables 5 and S4 provide the mean of the estimated 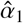 and 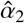 respectively over the 250 analyzed datasets for each of the 42 investigated configurations. As expected from the above evaluation of treeness tests, estimates of *α* were highly consistent among all the investigated configurations and similar for the two considered *f*_4_-ratio with a mean value varying between 0.245 and 0.248. Yet, a slight downward bias (always < 2%) could be noticed but the estimated 95% CIs were almost always optimal (or close to) since they contained the true simulated value (*α* = 0.25) from 90.0% to 95.2% of the time (Tables 5 and S4).

**Table 5.**
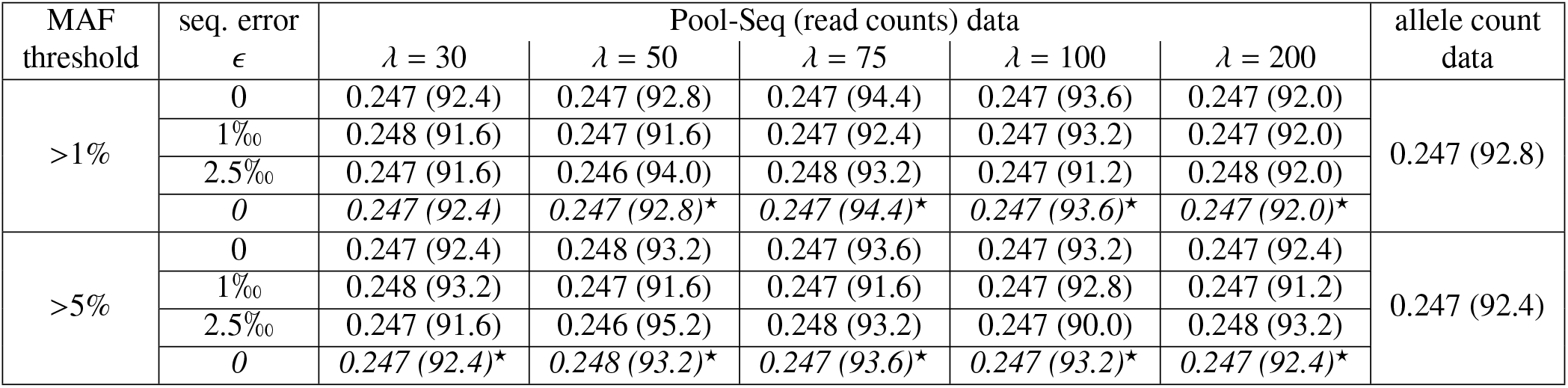
Comparison of *F*_4_-ratio based estimation of the simulated admixture proportion *α* in Figure 1 scenario for different types of data processing poolfstat analyses. For each MAF threshold (MAF> 1% or MAF> 5%), the table gives the mean of the estimated 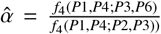 (across 250 independent simulated datasets) for 21 different types of analyses relying on i) allele count data; ii) 15 different Pool-Seq read count data (five mean coverages *λ* and three sequencing error rates *ϵ*); and iii) Pool-Seq read count data simulated with *ϵ* = 0 treated as allele counts (corresponding results of this bad practice are highlighted in italics and ^⋆^). The proportion (in %) of the 250 estimated 95% CIs that contain the true simulated value (*α* = 0.25) is given in parenthesis.

| MAF threshold | seq. error $\epsilon$ | Pool-Seq (read counts) data | | | | | allele count data |
| --- | --- | --- | --- | --- | --- | --- | --- |
| | | $\lambda = 30$ | $\lambda = 50$ | $\lambda = 75$ | $\lambda = 100$ | $\lambda = 200$ | |
| >1% | 0 | 0.247 (92.4) | 0.247 (92.8) | 0.247 (94.4) | 0.247 (93.6) | 0.247 (92.0) | 0.247 (92.8) |
|  | 1‰ | 0.248 (91.6) | 0.247 (91.6) | 0.247 (92.4) | 0.247 (93.2) | 0.247 (92.0) |  |
|  | 2.5‰ | 0.247 (91.6) | 0.246 (94.0) | 0.248 (93.2) | 0.247 (91.2) | 0.248 (92.0) |  |
|  | 0 | 0.247 (92.4) | 0.247 (92.8)* | 0.247 (94.4)* | 0.247 (93.6)* | 0.247 (92.0)* |  |
| >5% | 0 | 0.247 (92.4) | 0.248 (93.2) | 0.247 (93.6) | 0.247 (93.2) | 0.247 (92.4) | 0.247 (92.4) |
|  | 1‰ | 0.248 (93.2) | 0.247 (91.6) | 0.247 (91.6) | 0.247 (92.8) | 0.247 (91.2) |  |
|  | 2.5‰ | 0.247 (91.6) | 0.246 (95.2) | 0.248 (93.2) | 0.247 (90.0) | 0.248 (93.2) |  |
|  | 0 | 0.247 (92.4)* | 0.248 (93.2)* | 0.247 (93.6)* | 0.247 (93.2)* | 0.247 (92.4)* |  |

#### 3.1.5 Evaluation of graph fitting

We further estimated for all the simulated datasets branch lengths in drift units and admixture proportion *α* with their 95% CIs by fitting the simulated graph with `fit.graph`. As for the *f*_4_-ratio based estimation, estimates of *α* were virtually unbiased and consistent across all the 42 different investigated configurations (Figure S3). Nevertheless, the 95% CIs were always too narrow since they contained the actual value (*α* = 0.25) from only 40.8% to 74.4% of the time (Table S5) as expected from the *χ*^2^ approximation of the LRT underlying the computation of these CIs. Figures 4 and S4 plot the distributions of the estimated lengths for the ten branches of the simulated admixture graph branches (over the 250 estimates per configuration) when applying 5% and 1% MAF threshold respectively. The corresponding mean estimates and proportions of 95% CI’s including the true value are provided in Tables S6 to S15. Note that the branches *P*8 ↔ *R* and *P*9 ↔ *R* that are connected to the root *R* (Figure 1) can only be estimated jointly (as *τ*_*P*8↔*P*9_ = *τ*_*P*8↔*R*_ + *τ*_*P*9↔*R*_, *R* being arbitrarily set in its middle).

**Figure 4.**
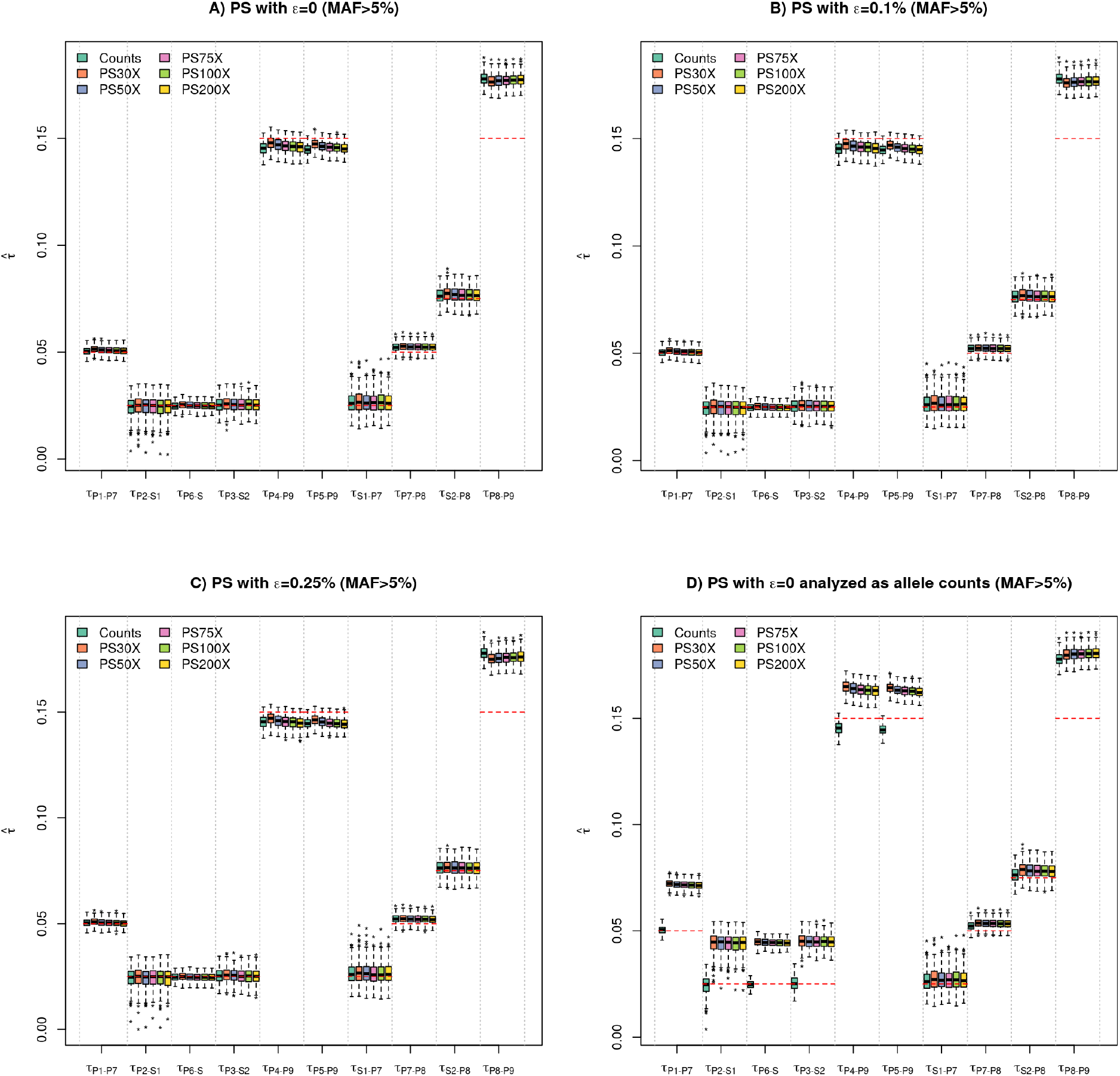
Distribution of the estimated drift-scaled lengths for all the branches in Figure 1 simulated scenario using admixture graph fitting (as implemented in the fit.graph function of poolfstat) for different types of data with a 5% threshold on the overall SNP MAF. Each box plot summarize the distribution of the 250 estimated lengths of each of the ten branches obtained from the analysis of either allele count dataset (“Counts”) or one of the five different simulated Pool-Seq read count datasets (“PS*λ*X”) with different mean coverages (*λ* = 30; 50; 75; 100; and200) as generated from the genotyping data simulated under the scenario depicted in Figure 1. Pool-Seq read count data were generated with no sequencing errors (*ϵ* = 0) in A) and D) and with a sequencing error rate of *ϵ* = 1‰ and *ϵ* = 2.5‰ in panel B) and C) respectively (Table S1). In D), the read count data were analyzed as allele counts which corresponds to a bad practice. Note that the two branches coming from the root are combined since the position of the root is not identifiable by the model (i.e., *τ*_*P*8↔*P*9_ = *τ*_*P*8↔*R*_ + *τ*_*P*9↔*R*_). Note that the box plots obtained from the analysis of count data are replicated in each panel for comparison purposes. For each branch, a red dotted line indicates the underlying simulated value. For Pool-Seq data, the overall MAF was estimated from read counts.

At the 5% MAF threshold, very similar performance were obtained for the allele count and the different Pool-Seq datasets whatever the simulated read coverage or sequencing error rates (Figures 4A, 4B, and 4C). Hence, mostly unbiased branch lengths were estimated for the four leaves (terminal branches) *τ*_*P*1↔*P*7_, *τ*_*P*2↔*S*1_, *τ*_*P*6↔*S*_ and *τ*_*P*3↔*S*2_. As previously observed with *α*, the estimated 95% CI’s remained too narrow particularly for *τ*_*P*2↔*S*1_ for which less than 50% of the CI’s contained the true value (Table S7) compared to more than 80% for *τ*_*P*1↔*P*7_ (Table S6). As expected from the the drift-scaling approximation, the estimated branch lengths tended to be slightly downwardly biased (ca. 2%) for the two other leaves (*τ*_*P*4↔*P*9_ and *τ*_*P*5↔*P*9_) but the estimated 95% CI displayed similar characteristics since from 48.0% to 87.6% contained the true values, the proportion increasing in Pool-Seq datasets when coverage and sequencing error decreased (Tables S10 and S11). Conversely, the internal branch lengths tended to be upwardly biased from a slight (ca. 2%) for *τ*_*S*1↔*P*7_, *τ*_*P*7↔*P*8_ and *τ*_*S*2↔*P*8_ (Tables S12 to S14), to a moderate amount (ca. 20%) for the root including branch *τ*_*P*8↔*P*9_, the true value being then always outside the estimated 95% CI’s (Table S15). Yet, when analyzing data with a lower MAF threshold of 1%, this bias almost completely vanished (Figure S4 and Table S15).

For the other branches, the estimates had similar characteristics (yet with a slightly decreased performance for the *τ*_*P*4↔*P*9_ and *τ*_*P*5↔*P*9_ leaves) for allele count data or Pool-Seq data simulated without sequencing error (Figure S4A). In agreement with previous observations, at the 1% MAF threshold, sequencing errors lead to strong downward bias at the lowest simulated coverages, i.e., when the percentage of false SNPs became non-negligible (Figures S4B and S4C). Finally, whatever the chosen MAF threshold, improperly analyzing read counts as allele count data always lead to a substantial upward bias of the lengths of all the leaves (Figure 4D). Notice however, that this had no or limited impact on the estimation of internal branch lengths.

#### 3.1.6 Evaluation of graph construction

To provide insights into the reliability of graph construction, we evaluated the performance of the `add.leaf` function in positioning the admixed population P6 on the underlying (((P1,P2),P3),(P4,P5)) tree (Figure 1) for the different types of simulated datasets. Table S16 gives the proportion of correctly inferred admixture graphs (i.e., corresponding to the simulated scenario) with a Δ_*BIC*_ > 6 support with all other tested graphs over the 250 analyzed datasets for each of the 42 investigated configurations. As the reference tree with rooted topology (((P1,P2),P3),(P4,P5)) consists of eight branches, P6 may be connected with either i) nine non-admixed edges (connection to either one of the eight branches or as an outgroup) or; ii) 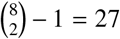 admixed edges from two-way admixture events. Except for the 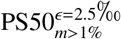 dataset (the one with the highest percentage of false SNPs), the correct admixture graph was always retrieved with a fairly high support (Δ_*BIC*_ > 15).

### 3.2 Analysis of real *Drosophila suzukii* Pool-Seq data

We here sketched the main findings from the analyses using `poolfstat` of a subset of the Pool-Seq data previously generated by Olazcuaga *et al* (2020) focusing on 14 population samples of the invasive species *D. suzukii*. For more details, we encourage readers to consult the Supplementary Vignette V2.

#### 3.2.1 Structuring of genetic diversity across the 14 populations

Overall, the estimated global *F*_*ST*_ across the 14 populations was 7.03% (95% CI; [6.90%; 7.32%]). Estimates of all the pairwise-population *F*_*ST*_ confirmed that populations tended to cluster according to their geographic area of origin (i.e., Asia, America and Hawaii; Figure 2A), with some geographically close populations showing low level of differentiation. For instance, in the American invasive area the US-Nca, US-Col and US-Nca populations all displayed pairwise *F*_*ST*_ significantly lower than 1%. Likewise, in the native area, the three populations CN-Bei, CN-Nin and CN-Lia originating from North-Western China were all found very closely related (all pairwise *F*_*ST*_ being lower or very close to 1%). Conversely, the Hawaiian sample (US-Haw) was found the most highly differentiated with all the other populations, all pairwise *F*_*ST*_ including US-Haw ranging from 11.7% (with US-Sok) to 17.0% (with US-Col) suggesting strong drift in this population as confirmed by its lowest estimated heterozygosity (Supplementary Vignette V2).

#### 3.2.2 *f*_3_-based tests of admixture suggest pervasive admixture in the invaded area

Out of the 14 sampled populations, two (CN-Lia and JP-Tok) from the Asian native area and four (US-Col, US-Nca, US-Wat and US-Wis) from the continental American invasive areas showed at least one significantly negative *f*_3_ at the 95% significance threshold (i.e., Z-score < −1.65). Table 6 summarizes for each of these 6 populations the number of significantly negative f3 together with the triplet with the lowest Z-score giving insights into the pair of populations that branch the closest to the two original sources (assuming a two-way admixture event). Except for CN-Lia, all the detected signals were significant at a far more stringent threshold (e.g., Z-score < −2.33 at 99% significance threshold). The *f*_3_ and 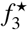 statistics gave almost exactly the same results (Supplementary Vignette V2).

**Table 6.** Results of the *f*_3_-based tests of admixture on populations from the *D. suzukii* invasive species. For all the population displaying at least one significant signal of admixture at the 95% significance threshold (f3 *Z* < − 1.65), the table gives the number of significant tests (out of the 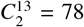 performed per population) and the triplet displaying the lowest Z-score (i.e., most significant test).

| Population | Origin | nb. of signif. tests ($f_3$ Z $< -1.65$) | triplet with the lowest $f_3$ Z-score |
| --- | --- | --- | --- |
| CN-Lia | Native | 1 | CN-Lia;CN-Shi,JP-Sap (Z=-1.66) |
| JP-Tok | Native | 11 | JP-Tok;CN-Nin,JP-Sap (Z=-7.11) |
| US-Col | Invasive (AM) | 2 | US-Col;BR-Pal,US-Wis (Z=-3.31) |
| US-Nca | Invasive (AM) | 6 | US-Nca;JP-Sap,US-Col (Z=-3.89) |
| US-Wat | Invasive (AM) | 13 | US-Wat;US-Sdi,US-Sok (Z=-23.6) |
| US-Wis | Invasive (AM) | 4 | US-Wis;JP-Sap,US-Col (Z=-5.02) |

In the native area, JP-Tok showed clear evidence of admixture with 11 significant tests that all involved JP-Sap (from Northern Japan) as a source proxy. The three lowest *f*_3_ values were obtained with three Chinese populations (CN-Nin, CN-Bei and CN-Shi in increasing order of *f*_3_). Assuming an admixture-graph like history, this suggests that the two populations branching the closest to the two sources of JP-Tok were JP-Sap and CN-Nin. The remaining Chinese population, CN-Lia showed some weak evidence for admixture with only one test barely significant at the 95% threshold for the triplet involving CN-Shi and JP-Sap as source proxies (Table 6).

Out of the seven invasive populations from continental America, the four populations US-Col, US-Wis, US-Nca and US-Wat showed strong evidence of admixture. Although it has up to now been considered as the closest to the first invading population of Continental America (based on historical records), the Western American US-Wat population displayed the strongest signals with 11 (strongly) significant tests. Interestingly, the three signals supported by the lowest (and hence more significant) Z-score all involved pairs of source population proxies originating from the continental American invasive area namely, in order of increasing Z-score (i.e., decreasing evidence), the (US-Sdi,US-Sok); (BR-Pal,US-Sok) and (US-Col,US-Sok) pairs. As the underlying *f*_3_ CI’s did not overlap with those of the other triplet configurations, these three pairs of populations may be considered as the closest (among the sampled populations) to the original US-Wat source populations. It is worth noting that the Western American US-Sok population was involved in nine of the 11 significant negative *f*_3_ statistic with US-Wat as a target. The three others populations, US-Col, US-Wis and US-Nca only had a moderate number of significant tests (compared to others). Such tests always involved at least one of the two other populations and overlapping *f*_3_ CI’s. This suggests complex patterns of recurrent admixture event among US-Col, US-Wis and US-Nca, a feature consistent with their low level of differentiation and close geographic origins.

#### 3.2.3 Exploring invasion scenarios with admixture graph construction and fitting

To provide further insights into the relationships of the surveyed populations and the probable scenarios of invasion of *D. suzukii* in the American area, we relied on admixture graph construction. Our purpose was not to build a comprehensive admixture graph for the 14 populations, which may be elusive given the close relationships of some populations and the pervasiveness of recent and presumably recurrent admixture events among the different populations, but rather to identify key regional event that occurred at early time of the invasion history of the species. From our extensive analyses (Supplementary Vignette V2), we were in fine able to build and estimate the parameters of two admixture graphs represented in Figure 2B and C. The first admixture graph described the somewhat complex and so far non-investigated relationships among the populations of the native area (including the early invasive population established in Hawaii since 1980) with a very good fit since the Z-score of the residuals for the worst fitted *f* -statistics was 1.06 (Figure 2B). In agreement with previous findings (and geographic proximity), the Hawaiian population was found more closely related to the Japanese population JP-Sap than to the other Chinese populations but it experienced a strong differentiation from their common ancestor (named JP in Figure 2B) with an estimated branch length of 0.255 drift units 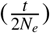. Yet, it was not possible from our data to definitively conclude that US-Haw originates from a Japanese population since we have no element to claim that the (ancestral) node population JP was located in Japan. To that end additional sampling of Japanese populations would be required. The inferred graph also confirmed above *f*_3_-based test results of an admixed origins of JP-Tok between a population closely related to JP-Sap (the main contributor) and a second source likely of Chinese origin although the same caution as for JP are needed regarding the geographic origins of this internal node populations. Similarly, CN-Lia was found admixed with a contributing source of Chinese ancestry related to CN-Shi largely predominant (estimated contribution 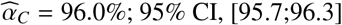), and a second (minor) contributing source of presumably Japanese origin (related to JP-Sap). This may explain why the corresponding *f*_3_-based test was only barely significant (Table 6). Interestingly, the graph topology also allowed estimating the Chinese ancestry of CN-Lia based on *F*_*4*_ -ratio resulting in consistent but larger 95% 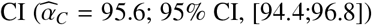 as expected from above simulation study. CN-Nin, the remaining population from the native area, could not be positioned with reasonable accuracy onto the admixture graph of Figure 2B, the resulting worst fitted *f* -statistics associated to the best fitting graph having a Z-score=3.43. However, both its genetic proximity with CN-Lia and the best fitting admixture graph resulting from its positioning onto the scaffold tree including US-Haw, JP-Sap, CN-Bei and CN-Shi suggested a small amount of Japanese introgression (see Supplementary Vignette V2 for more details).

The second admixture graph represented in Figure 2C allowed providing insights into the history of introduction of *D. suzukii* into the American continent. It related the three continental American population, US-Sok, US-Wat and BR-Pal to a scaffold including the four unadmixed populations US-Haw, JP-Sap, CN-Shi and CN-Bei with a good fit (the worst fitted *f* -statistics had a Z-score=-1.83). The underlying scenario suggested that continental American populations originated from at least two major and successive admixture events. The first admixture event lead to the internal node population named Am1 and occured in balanced proportions between two sources, a Japanese one closely related to JP-Sap and a Hawaiian one relatively distantly related to US-Haw (according to the estimated branch lengths). The US-Sok population was the sampled continental American population the closest to Am1 and may thus be assumed the most closely related to the first invading population (in agreement with *f*_3_-based test results). Yet US-Sok remained separated by about 0.0816 drift units from Am1 which may explain why no significantly negative *f*_3_ were found for triplets with US-Sok as a target.

The second major admixture events occurred between the internal node population named Am2 and a Chinese population closely related to the common ancestor of CN-Bei and CN-Shi, with CN-Shi contributing slightly more (58.5% against 41.5% for the other Am1 related ancestor). Interestingly, the closest Am2 representatives among the sampled populations were BR-Pal and US-Sdi (also in agreement with *f*_3_-based tests) suggesting a more Southern geographical origin for Am2. We found that some additional ancestry from a ghost population or recurrent admixture events (e.g., related to Hawaiian populations) may also have contributed to US-Sdi, but this lead to a poor fit (worst fitted *f* -statistics Z-score=-5.87 for the best fitting graph resulting from the positioning of US-Sdi onto the graph, see Supplementary Vignette V2). Therefore, US-Sdi is not represented in Figure 2C. Although geographically distant, the Brazilian population BR-Pal thus appeared as the best proxy for Am2 thereby suggesting a rapid spread of *D. suzukii* in South America from this population without any subsequent admixture events. Additional (and preferably ancient) sample from South-American populations would help refining this scenario. Finally, according to the inferred graph, US-Wat was found to originate from a recent admixture between a population very closely related to US-Sok (and thus Am1) and a population deriving from Am2 with similar contributions of both.

In agreement with *f*_3_-based admixture tests that suggested complex admixture histories among the closely related US-Col, US-Wis and US-Nca populations, no satisfactory admixture graph could be found when trying to position each of these onto the Figure 2C graph. Nevertheless, their resulting best fitted graphs all suggested a high contribution of the Am2 admixed source, a second contributing source being related to Japanese populations (Supplementary Vignette V2).

## 4 Discussion

### 4.1 A new version of poolfstat for *f-*statistics estimation and associated inference from both Pool-Seq and allele count data

The R package `poolfstat` was originally developed by Hivert *et al* (2018) to implement an unbiased estimator of *F*_*ST*_ for PoolSeq data and provide utilities to facilitate manipulation of such data. We here proposed a substantially improved version that implements unbiased estimators of *F*_2_, *F*_3_ and *F*_4_ parameters together with their scaled versions (i.e., pairwise *F*_*ST*_, 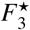 and *D* respectively). Although we primarily focused on the analysis of Pool-Seq data, we extended the package to analyze standard allele count (as obtained from individual genotyping or sequencing data) and to implement unbiased estimators equivalent to those available in the `AdmixTools` suite (Patterson *et al*, 2012) allowing us in turn to validate our estimation procedure. Recently, the `admixr` package was developed to interface most of the `AdmixTools` programs with R for the estimation of *f* -statistics (only from allele count data), with the noticeable exception of the admixture graph fitting program `qpGraph` (Petr *et al*, 2019). We implemented in `poolfstat` our own functions for fitting, building, visualizing and quality assessment of admixture graphs based on the estimated *f* -statistics. The underlying procedures shared strong similarities with those implemented in `qpGraph` (Patterson *et al*, 2012) resulting on the same fitting on some examples (e.g., Supplementary Vignette V1) or also `MixMapper` (Lipson *et al*, 2013, 2014) programs. As recognized by the developers themselves, the latter program specifically developed for admixture inference from allele count data was written in C++ and MATLAB making it ‘cumbersome to use’ for users, as ourselves, with no MATLAB license. Moreover, to facilitate local exploration of the admixture graphs space, we also implemented in `poolfstat` efficient semi-automated building utilities (`add.leaf` and `graph.builder` functions). It should be noticed that although it does not include functions for the estimation of *f* -statistics, the `admixturegraph` R package (Leppälä *et al*, 2017) also provides several alternative valuable utilities for the fitting (based on a slightly different approach), the manipulation, and the visualization of admixture graphs together with utilities for the plotting of the statistics with their confidence intervals or the symbolic derivation *f* -statistics (as `poolfstat`). Overall, our effort of developing with `poolfstat` a self-contained, efficient and user-friendly R package capable of performing the entire workflow for *f* -statistics based demographic inference from both standard allele count and Pool-Seq read count data will hopefully make such a powerful framework accessible to a wider range of researchers and biological models.

### 4.2 A unified definition of the *F* parameters in terms of probability of gene identities

To derive our unbiased estimators, we proposed to recapitulate and unify the different definitions of the *F* and *D* parameters in terms of probability of gene identity within population (*Q*_1_) or between pairs of populations (*Q*_2_) as summarized in equation 1. This formulation offers a complementary perspective to the original description of these parameters in terms of covariance of allele frequencies (Patterson *et al*, 2012). In practice, a little algebra shows that the unbiased estimators derived from these two alternative formulations are strictly equivalent (i.e., when comparing eq. 6 for allele count data with Appendix A in Patterson *et al*, 2012). Formally, the *Q*_1_ and *Q*_2_ probabilities can be viewed as expected identity (in state) of genes across independent replicates of the (stochastic) evolutionary process (Rousset, 2007) that may themselves be expressed as a function of other demo-genetic population parameters. Hence, the obtained expressions for *F*_2_, *F*_3_ and *F*_4_ in terms of *Q*_1_ and *Q*_2_ probabilities can be directly related to those by Peter (2016) in terms of coalescent times which allowed him to provide an in-depth exploration of their theoretical properties under a wide range of demographic models other than admixture graphs (see e.g., Figure 7 in Peter, 2016). More precisely, under an infinite-site mutation model with constant per-generation mutation rate *µ*, the probability that two genes are identical in state is 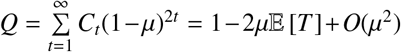, where *C*_*t*_ is the probability that the two genes coalesced *t* generations in the past and 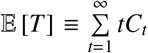 is the expected coalescence time of two genes (see Slatkin, 1991; Rousset, 2007, pp.58-59). Using 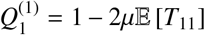 and 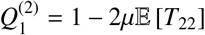 as the IIS probabilities within populations 1 and 2 respectively and *Q*_2_ = 1 − 2*µ*𝔼 [*T*_12_] as the IIS probability between 1 and 2 allows recovering equations 16 (after fixing a typo into it), 20c and 24 by Peter (2016) for *F*_2_, *F*_3_ and *F*_4_ respectively. Likewise, the estimators derived from (unbiased) estimators of *Q*_1_ and *Q*_2_ are equivalent to those expressed in terms of average pairwise differences between and within populations which are natural estimators for 2*µ*𝔼 [*T*] terms as proposed by Peter (2016, eq. 17) for *F*_2_ estimator based on allele count data (e.g., noting that 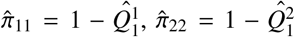 and 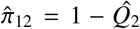 following his notations). For Pool-Seq data, replacing the latter estimators of nucleotide diversities by the unbiased estimators described in Ferretti *et al* (2013, eqs. 3 and 10) would also result in the same estimator for *F*_2_ (and other parameters) as those defined in our equation 6.

In practice, estimators are obtained by averaging over (a high) number of SNPs which amounts assuming that each represent an independent outcome of a common demographic processes that shaped the genome-wide patterns of genetic diversity. This generally allows to provide accurate estimations and LD between markers (i.e., violation of the marker independence assumption) can be accounted for with block-jackknife estimation of standard errors (Patterson *et al*, 2012). Importantly, as originally noticed by Patterson *et al* (2012), expressions of *F*_2_, *F*_3_ and *F*_4_ in terms of coalescent times (Peter, 2016) show explicitly that they both depend on the demography (via 𝔼 [*T*]) and the marker mutation rate (*µ*). In the scaled versions of *F*_2_ and *F*_3_ (*F*_*ST*_ and 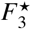 respectively), the parameter *µ* cancels out making them presumably more comparable across different datasets. It should however be noticed, that for demographic inference purposes, scaling of the *f* -statistics is not needed. Indeed, the three-population test of admixture is informed by the sign of *f*_3_ which is not affected by the denominator of 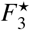. Similarly, the four-population test evaluates departure of *f*_4_ (i.e., the numerator of *D*) from a null value expected under the hypothesis of treeness. Patterson *et al* (2012) also showed both analytically and using simulations that *F*_3_ and *F*_4_ estimates remained mostly robust to various realistic SNP ascertainment scheme. It is finally worth stressing that admixture graph inference only requires additivity of *F*_2_ (Patterson *et al*, 2012), a feature not fulfilled by *F*_*ST*_ (or 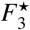 and *D*).

### 4.3 Estimation of *f* -statistics and inference from Pool-Seq data

Our simulation study showed that accurate estimates of *F* and *D* parameters could be obtained from Pool-Seq data from the unbiased estimators we developed, thereby extending our findings for the Pool-Seq *F*_*ST*_ estimator (Hivert *et al*, 2018). With no sequencing error, this remained true even at a read coverage as low as 30X which was here lower than our simulated haploid sample size of 50. Increasing the coverage only provided marginal gain. When introducing sequencing errors, the performance of the estimators tended to decrease for the lowest investigated read coverages (up to 50X) and MAF filtering threshold. This was however essentially due to the presence of spurious SNPs that were not completely filtered out when considering too loose criteria. As a result, simply increasing the threshold on the overall MAF (computed from read counts over all the pool samples) to 5% allowed to remove all the spurious SNPs and recover accurate estimates of the parameters at the lowest read coverages. In agreement with original observations made for allele count data (Patterson *et al*, 2012), all the *f* −statistics based analyses (i.e., three-population test of admixture, four-population test of treeness, *F*_4_-ratio estimation of admixture proportions or admixture graph fitting) remained remarkably robust to a MAF-based ascertainment process. From our simulation study, discarding lowly polymorphic SNPs was only found to increase the bias of the drift-scaled length estimates of internal branch in admixture graph. In practice, cost-effective designs consisting of sequencing pools of 30 to 50 individuals at a 50-100X coverage and applying MAF threshold of 5% to filter the called SNPs are expected to provide good performance for all the different *f* −statistics based inference methods we presented here.

For Pool-Seq data, all the above conclusions were nevertheless only valid for the analyses based on the unbiased estimator that accounts for the additional level of variation introduced by the sampling of the DNA of pooled individuals (non identifiable) at the sequencing step. We found that improperly analyzing Pool-Seq read counts as standard allele counts had high detrimental consequence on the estimation of all the *F* parameters that involved *Q*_1_ probabilities (within population probability of identity) in their definition, i.e., *F*_2_, *F*_*ST*_ (as previously observed by Hivert *et al*, 2018, see also Figure S5), *F*_3_ and 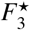 leading to a complete loss of power of the associated three-population test in our simulation. When processing admixture graph fitting, this also resulted in a strong upward bias in the estimation of branch lengths, including the external ones that were accurately estimated when relying on unbiased estimators. Loosely speaking, not accounting for the extra-variance introduced by the sampling of reads has the same effect of adding a (substantial) amount of extra drift explaining the two aforementioned observations. Although not investigated here (and of little interest since we should definitely rely on unbiased estimators), the amount of extra variance may be inversely proportional to the pool haploid sample size (i.e., bias may decrease with increasing pool sample size). Conversely, analyzing Pool-Seq read counts as standard allele counts did not affect the performance of the *f*_4_- (and *D*) based test of treeness or the estimation of admixture proportion from *F*_4_-ratio. This was expected from the properties of the underlying parameters that only depends on the *Q*_2_ probabilities across the different pairs of population involved in the quadruplet of interest (eq. 3) resulting in the same estimators (see eqns. 4 and 5) for both allele count and Pool-Seq data. More generally, analyzing Pool-Seq read count data with popular programs that were developed for standard allele count data such as those from the `AdmixTools` (Patterson *et al*, 2012) or `TreeMix` (Pickrell & Pritchard, 2012) suites should definitely be avoided and, if not, results should be carefully interpreted.

### 4.4 Insights into the history of the invasive species *D. suzukii* from Pool-Seq data analysis

To illustrate both the power and limitations of *f* −statistics based methods for historical and demographic inference as implemented in `poolfstat`, we analyzed Pool-Seq data available for 14 populations of the invasive species *D. suzukii* (Olazcuaga *et al*, 2020). These population samples were representative of both the Asian native area and the recently invaded American area. Most of them consisted of individuals originating from the same sites as those genotyped in Fraimout *et al* (2017) at microsatellite markers and analyzed under an ABC-RF framework. The results remained consistent between the two studies, both of them pointing to complex invasion pathways including multiple introductions leading to admixed origins of the continental American populations. The main source contributions were from Hawaii, where *D. suzukii* was described about 30 years earlier and the native area (China and Japan). However, some inferred scenarios appeared somewhat conflicting. First, for the Hawaiian population that played a key role in American invasion route, both `poolfstat` and Fraimout *et al* (2017) suggested a Japanese origin. However, we here found that the sample the closest to the source (internal node population JP in Figure 2) was JP-Sap (sampled in Sapporo) while Fraimout *et al* (2017) rather concluded it was JP-Tok (sampled in Tokyo) which was not found to be directly contributing to US-Haw in `poolfstat` analyses and was even found to be admixed by native populations from Japan and China. In the ABC-RF treatments by Fraimout *et al* (2017), all populations from the native area were assumed to be non-admixed and no “ghost” (i.e., unsampled) populations were included in the model whereas such populations are present in admixture graphs through internal nodes. Moreover, the samples from Hawaii and Tokyo both differed in their exact location and collection date (2013 and 2016 for Hawaii, 2014 and 2016 for Tokyo) between the two studies, which may further explain the observed discrepancies and more generally promotes the sequencing of additional samples in this area to better resolve the origin of the Hawaiian population(s).

Interestingly, `poolfstat` results challenged the initial view about the pioneering origin of the Californian population US-Wat in the invasion of continental America (as suggested by historical records) suggesting it rather originates from an admixture between two already established but unsampled continental American populations, one presumably Northern (related to Am1 and here represented by US-Sok) and the other presumably more Southern (related to Am2 and here represented by BR-Pal from Brazil and US-Sdi from South-California). This discrepancy between Fraimout *et al* (2017) and `poolfstat` results points to three key issues. First, a too strong reliance on the reported date of first observation of the species in the invasive area when formalizing the scenario to be compared in ABC modeling may actually mislead inference and this especially since *D. suzukii* was first observed at very close dates in the US-Wat, US-Sdi and US-Sok sampled locations (i.e., 2008, 2009 and 2009, respectively). As a matter of fact, Fraimout *et al* (2017) only considered scenarios in which US-Wat was the first population introduced in continental America. Second, in ABC, scenarios are defined by hand justifying the use of dates of first observation to minimize their number (Estoup & Guillemaud, 2010). The functions implemented in `poolfstat` circumvent this constraint by facilitating a quick and automatic exploration of the admixture graph space to identify key historical events relating the populations of interest. Third, our finding reinforces the concern that the formalization of invasion scenarios including the possibility of unsampled populations is crucial. This possibility is by construction included in admixture graph construction but is also possible in ABC modeling (e.g. Guillemaud *et al*, 2010). Similarly, Fraimout *et al* (2017) argued for an admixed origin of the Brazilian BR-Pal population (first observed in 2013) between undefined North-Western and North-Eastern American sources, while we here found that this population was the best proxy for the ancestral “ghost” American population Am2 (Figure 2C) which may be viewed as one of the main contributor of all the sampled North-American populations (but US-Sok). Again, this results underline advantages of not relying on historical dates as for `poolfstat` analyses, and promotes the sequencing of additional samples in South and North-Western America areas to more thoroughly decipher the invasion routes followed by continental American populations.

If Pool-Seq data analyzed with `poolfstat` allowed to refine historical and demographic scenarios in both the native and invasive areas, the *D. suzukii* Pool-Seq data analysis also illustrated some inherent constraints imposed to the modeled demographic history when fitting admixture graph. In particular, more complex histories involving recurrent admixture events turned out to be difficult or even impossible to fit unless a number of source key samples are included, as observed here for the North-Eastern American populations. In real-life applications involving a large number of invasive populations characterized by numerous and recurrent introduction events, summarizing precisely and with a good fit the history of all surveyed populations with a comprehensive admixture graph may remain elusive. However, as previously underlined (Patterson *et al*, 2012; Lipson & Reich, 2017; Lipson, 2020), in addition to providing robust formal tests of admixture or treeness, a decisive advantage of *f* −based inference methods is to allow straightforward assessment of the fitted admixture graph by carefully inspecting and reporting Z-score of the residuals of the fitted statistics, an option not available in other related methods such as `TreeMix` (Pickrell & Pritchard, 2012). Beyond modeling the history of populations as admixture graphs (via formal tests of admixture of treeness or graph fitting), Peter (2016) provided valuable theoretical insights to interpret the estimated *f* −statistics under alternative demographic models such as island, stepping-stone or serial founder models. This suggests in turn that these statistics should be informative to estimate the parameters of demographic scenarios more complex than admixture graphs (e.g., under an ABC framework as in Collin *et al*, 2021).

## Supporting information

Supplementary Vignette V1

Supplementary Vignette V2

## Acknowledgements

This work was supported by the French National Research Agency (ANR) for the projects SWING (ANR-16-CE02-0015-01) and GANDHI (ANR-20-CE02-0018).

## Data Accessibility

The vcf file generated for the *D. suzukii* Pool Seq data is publicly available from the Zenodo repository (http://doi.org/10.5281/zenodo.4709080). Further details are provided in Supplementary Vignettes V1 and V2.

## Supporting Information

Additional supporting information is available online and consists of three pdf files: i) vignette V1 for the R package `poolfstat` providing a detailed tutorial using Pool-Seq and allele count data simulated under the scenario depicted in Figure 1; ii) vignette V2 detailing the analysis of the real *Drosophila suzukii* Pool-Seq data using the R package `poolfstat`; and iii) a single pdf file with the sixteen supplementary Tables and the five supplementary Figures.

