## Supplementary Tables and Figures for "*f*-statistics estimation and admixture graph construction with Pool-Seq or allele count data using the R package *poolfstat*"

| Code | Count data type | MAF threshold | Coverage ($\lambda$) | Seq. error ($\epsilon$) | Nb. of SNPs Mean ± s.d. | Nb. of false SNPs Mean ± s.d. |
|---|---|---|---|---|---|---|
| $AC_{m>1\%}$ | Alleles | 1% | - | - | 471,919 ± 1,474 | - |
| $AC_{m>5\%}$ | Alleles | 5% | - | - | 240,369 ± 1,118 | - |
| $PS30^{\epsilon=0}_{m>1\%}$ | Reads | 1%$^{\star}$ | 30 | 0 | 406,742 ± 1,362 | - |
| $PS50^{\epsilon=0}_{m>1\%}$ | Reads | 1% | 50 | 0 | 442,426 ± 1,419 | - |
| $PS75^{\epsilon=0}_{m>1\%}$ | Reads | 1% | 75 | 0 | 449,663 ± 1,433 | - |
| $PS100^{\epsilon=0}_{m>1\%}$ | Reads | 1% | 100 | 0 | 454,056 ± 1,434 | - |
| $PS200^{\epsilon=0}_{m>1\%}$ | Reads | 1% | 200 | 0 | 462,429 ± 1,466 | - |
| $PS30^{\epsilon=0}_{m>5\%}$ | Reads | 5% | 30 | 0 | 246,560 ± 1,111 | - |
| $PS50^{\epsilon=0}_{m>5\%}$ | Reads | 5% | 50 | 0 | 245,296 ± 1,108 | - |
| $PS75^{\epsilon=0}_{m>5\%}$ | Reads | 5% | 75 | 0 | 244,706 ± 1,126 | - |
| $PS100^{\epsilon=0}_{m>5\%}$ | Reads | 5% | 100 | 0 | 244,432 ± 1,140 | - |
| $PS200^{\epsilon=0}_{m>5\%}$ | Reads | 5% | 200 | 0 | 243,993 ± 1,131 | - |
| $PS30^{\epsilon=1‰}_{m>1\%}$ | Reads | 1% | 30 | 1‰ | 613,255 ± 1,471 | 206,051 ± 443 |
| $PS50^{\epsilon=1‰}_{m>1\%}$ | Reads | 1% | 50 | 1‰ | 871,953 ± 1,605 | 432038 ± 654 |
| $PS75^{\epsilon=1‰}_{m>1\%}$ | Reads | 1% | 75 | 1‰ | 445,922 ± 1,417 | 3,866 ± 58 |
| $PS100^{\epsilon=1‰}_{m>1\%}$ | Reads | 1% | 100 | 1‰ | 440,012 ± 1,399 | 196 ± 14 |
| $PS200^{\epsilon=1‰}_{m>1\%}$ | Reads | 1% | 200 | 1‰ | 408,432 ± 1,300 | 0 ± 0 |
| $PS30^{\epsilon=1‰}_{m>5\%}$ | Reads | 5% | 30 | 1‰ | 246,442 ± 1,123 | 0 ± 0 |
| $PS50^{\epsilon=1‰}_{m>5\%}$ | Reads | 5% | 50 | 1‰ | 243,699 ± 1,113 | 0 ± 0 |
| $PS75^{\epsilon=1‰}_{m>5\%}$ | Reads | 5% | 75 | 1‰ | 240,422 ± 1,111 | 0 ± 0 |
| $PS100^{\epsilon=1‰}_{m>5\%}$ | Reads | 5% | 100 | 1‰ | 236,566 ± 1,089 | 0 ± 0 |
| $PS200^{\epsilon=1‰}_{m>5\%}$ | Reads | 5% | 200 | 1‰ | 215,471 ± 1,007 | 0 ± 0 |
| $PS30^{\epsilon=2.5‰}_{m>1\%}$ | Reads | 1% | 30 | 2.5‰ | 3,402,213 ± 2,147 | 2,999,100 ± 1,773 |
| $PS50^{\epsilon=2.5‰}_{m>1\%}$ | Reads | 1% | 50 | 2.5‰ | 6,803,455 ± 2,847 | 6,380,412 ± 2,512 |
| $PS75^{\epsilon=2.5‰}_{m>1\%}$ | Reads | 1% | 75 | 2.5‰ | 686,967 ± 1,464 | 282,076 ± 530 |
| $PS100^{\epsilon=2.5‰}_{m>1\%}$ | Reads | 1% | 100 | 2.5‰ | 417,951 ± 1,243 | 39,043 ± 189 |

| | | | | | | |
|---|---|---|---|---|---|---|
| $PS200^{\epsilon=2.5‰}_{m>1\%}$ | Reads | 1% | 200 | 2.5‰ | 251,968 ± 863 | 2 ± 1 |
| $PS30^{\epsilon=2.5‰}_{m>5\%}$ | Reads | 5% | 30 | 2.5‰ | 243,414 ± 1,088 | 0 ± 0 |
| $PS50^{\epsilon=2.5‰}_{m>5\%}$ | Reads | 5% | 50 | 2.5‰ | 234,158 ± 1,100 | 0 ± 0 |
| $PS75^{\epsilon=2.5‰}_{m>5\%}$ | Reads | 5% | 75 | 2.5‰ | 220,095 ± 1,050 | 0 ± 0 |
| $PS100^{\epsilon=2.5‰}_{m>5\%}$ | Reads | 5% | 100 | 2.5‰ | 203,702 ± 967 | 0 ± 0 |
| $PS200^{\epsilon=2.5‰}_{m>5\%}$ | Reads | 5% | 200 | 2.5‰ | 132,980 ± 669 | 0 ± 0 |

**Table S1. Description of the simulated datasets.** A total of 250 independent genotyping datasets were simulated according to the demographic scenario represented in Figure 1. For each simulated genotyping datasets, we generated two allele count datasets applying 1% or 5% threshold (column 3) on the overall MAF (computed from the allele counts); and simulated fifteen Pool-Seq read count datasets with five different mean read coverages $\lambda$ (column 4) and three different sequencing error rates $\epsilon$ (column 5) and applied 1% or 5% threshold (column 3) on the overall MAF (computed from read counts) for each of them (resulting in a total of 30 Pool-Seq read count dataset per simulated genotyping data). As highlighted by the asterisk ($\star$), for Pool-Seq data with $\lambda = 30$, the actual MAF threshold was slightly higher than indicated due the additional filtering criterion on the minimal read count (MRC>2, see Material and Methods). Indeed, when $\lambda = 30$, the overall coverage (that follows a Poisson distribution with parameter $6 \times 30 = 180$) is $\leq 200$ for on expected proportion of 93.5% of the SNPs resulting to a more stringent MAF threshold of $\frac{2}{6\times30} = 1.11\%$ when requiring MRC>2. The last two columns give the average number (and standard deviation) of SNPs and false SNPs (for read count data simulated with $\epsilon > 0$) computed over the 250 independent simulated datasets available for each of the 32 configurations.

| MAF threshold | seq. error $\epsilon$ | Pool-Seq (read counts) data | | | | | allele count data |
|---|---|---|---|---|---|---|---|
| | | $\lambda = 30$ | $\lambda = 50$ | $\lambda = 75$ | $\lambda = 100$ | $\lambda = 200$ | |
| >1% | 0 | 82.0 (0.00) | 84.4 (0.00) | 86.0 (0.00) | 86.0 (0.00) | 85.2 (0.00) | 85.6 (0.00) |
| | 1‰ | 100 (0.00) | 100 (0.00) | 86.8 (0.00) | 87.2 (0.00) | 86.4 (0.00) | |
| | 2.5‰ | 100 (0.00) | 100 (6.47) | 99.6 (0.00) | 92.8 (0.00) | 88.4 (0.00) | |
| | *0* | *0.00 (0.00)*⋆ | *0.00 (0.00)*⋆ | *0.00 (0.00)*⋆ | *0.00 (0.00)*⋆ | *0.00 (0.00)*⋆ | |
| >5% | 0 | 93.6 (0.00) | 95.2 (0.00) | 96.4 (0.00) | 96.0 (0.00) | 96.0 (0.00) | 96.8 (0.00) |
| | 1‰ | 94.0 (0.00) | 96.8 (0.00) | 96.4 (0.00) | 97.2 (0.00) | 96.8 (0.00) | |
| | 2.5‰ | 94.0 (0.00) | 96.0 (0.00) | 96.0 (0.00) | 97.2 (0.00) | 96.8 (0.00) | |
| | *0* | *0.00 (0.00)*⋆ | *0.00 (0.00)*⋆ | *0.00 (0.00)*⋆ | *0.00 (0.00)*⋆ | *0.00 (0.00)*⋆ | |

**Table S2. Comparison of the performance of $f_3^\star$-based tests of admixture for different types of data simulated under the Figure 1 scenario processing `poolfstat` analyses.** For each MAF threshold (MAF> 1% or MAF> 5%), the table gives True and False (in parenthesis) Positive Rates (in %) for 21 different types of analyses relying on i) allele count data; ii) 15 different Pool-Seq read count data (five mean coverages $\lambda$ and three sequencing error rates $\epsilon$); and iii) Pool-Seq read count data simulated with $\epsilon = 0$ treated as allele counts (corresponding results of this bad practice are highlighted in italics and $^\star$). Each TPR was computed from the analysis of 250 independent datasets (generated from the data simulated under Figure 1 demographic scenario) as the proportion of $f_3^\star$ with an associated Z-score $< -1.65$ (95% significance threshold) for the (P6;P2,P3) population triplet (n=250 estimates). The FPR was similarly computed as the proportion of $f_3^\star$ with an associated Z-score$< -1.65$ among all the 50 population triplets that do not involve P6 as target population (n=250×50=12,250 estimates).

| MAF threshold | seq. error $\epsilon$ | Pool-Seq (read counts) data | | | | | allele count data |
|---|---|---|---|---|---|---|---|
| | | $\lambda = 30$ | $\lambda = 50$ | $\lambda = 75$ | $\lambda = 100$ | $\lambda = 200$ | |
| >1% | 0 | 94.0 (0.05) | 94.4 (0.06) | 94.1 (0.04) | 94.5 (0.05) | 94.3 (0.02) | 94.2 (0.02) |
| | 1‰ | 94.3 (0.04) | 94.2 (0.03) | 94.3 (0.03) | 94.3 (0.03) | 94.2 (0.05) | |
| | 2.5‰ | 94.8 (0.06) | 94.5 (0.05) | 94.8 (0.03) | 94.5 (0.06) | 94.3 (0.04) | |
| | *0* | *94.0 (0.05)*$^{\star}$ | *94.4 (0.06)*$^{\star}$ | *94.1 (0.04)*$^{\star}$ | *94.5 (0.05)*$^{\star}$ | *94.3 (0.02)*$^{\star}$ | |
| >5% | 0 | 94.5 (0.14) | 94.3 (0.11) | 94.1 (0.14) | 94.8 (0.09) | 94.3 (0.08) | 94.3 (0.11) |
| | 1‰ | 94.5 (0.09) | 94.5 (0.11) | 94.5 (0.13) | 94.2 (0.09) | 94.0 (0.15) | |
| | 2.5‰ | 95.2 (0.13) | 93.8 (0.11) | 94.2 (0.12) | 94.5 (0.11) | 94.3 (0.13) | |
| | *0* | *94.5 (0.14)*$^{\star}$ | *94.3 (0.11)*$^{\star}$ | *94.1 (0.14)*$^{\star}$ | *94.9 (0.09)*$^{\star}$ | *94.3 (0.08)*$^{\star}$ | |

**Table S3. Comparison of the performance of $D$-based test of treeness for different types of data simulated under the Figure 1 scenario processing `poolfstat` analyses.** For each MAF threshold (MAF> 1% or MAF> 5%), the table gives True and False (in parenthesis) Positive Rates (in %) for 21 different types of analyses relying on i) allele count data; ii) 15 different Pool-Seq read count data (five mean coverages $\lambda$ and three sequencing error rates $\epsilon$); and iii) Pool-Seq read count data simulated with $\epsilon = 0$ treated as allele counts (corresponding results of this bad practice are highlighted in italics and $^{\star}$). Each TPR was computed from the analysis of 250 independent datasets (generated from the data simulated under Figure 1 demographic scenario) as the proportion of $D$-statistics with an associated absolute Z-score < 1.96 (95% significance threshold) among all the eight population quadruplets ((P1,P2;P3,P4); (P1,P2;P3,P5); (P1,P2;P4,P5); (P1,P3;P4,P5); (P1,P6;P4,P5); (P2,P3;P4,P5); (P2,P6;P4,P5); (P3,P6;P4,P5)) with a null expected $F_4$ (n=250×8=2,000 estimates). The FPR was similarly computed as the proportion of $D$-statistics with an associated absolute Z-score < 1.96 among all the 37 remaining population quadruplets (n=250×37=9,250 estimates).

| MAF threshold | seq. error $\epsilon$ | Pool-Seq (read counts) data | | | | | allele count data |
|---|---|---|---|---|---|---|---|
| | | $\lambda = 30$ | $\lambda = 50$ | $\lambda = 75$ | $\lambda = 100$ | $\lambda = 200$ | |
| >1% | 0 | 0.246 (94.0) | 0.246 (93.2) | 0.246 (92.0) | 0.245 (93.6) | 0.246 (93.2) | 0.246 (92.8) |
| | 1‰ | 0.246 (92.0) | 0.247 (96.0) | 0.246 (92.8) | 0.246 (93.2) | 0.245 (91.6) | |
| | 2.5‰ | 0.245 (91.2) | 0.245 (92.8) | 0.246 (92.0) | 0.246 (92.4) | 0.246 (94.0) | |
| | *0* | *0.246 (94.0)*$^{\star}$ | *0.246 (93.2)*$^{\star}$ | *0.246 (92.0)*$^{\star}$ | *0.245 (93.6)*$^{\star}$ | *0.246 (93.2)*$^{\star}$ | |
| >5% | 0 | 0.246 (93.2) | 0.246 (93.2) | 0.246 (92.4) | 0.245 (94.0) | 0.246 (93.6) | 0.246 (93.2) |
| | 1‰ | 0.246 (91.6) | 0.247 (94.8) | 0.246 (92.4) | 0.246 (93.2) | 0.246 (92.0) | |
| | 2.5‰ | 0.245 (91.2) | 0.245 (93.2) | 0.246 (91.2) | 0.246 (92.0) | 0.246 (92.8) | |
| | *0* | *0.246 (93.2)*$^{\star}$ | *0.246 (93.2)*$^{\star}$ | *0.246 (92.4)*$^{\star}$ | *0.245 (94.0)*$^{\star}$ | *0.246 (93.6)*$^{\star}$ | |

**Table S4. Comparison of $F_4$-ratio based estimation of the simulated admixture proportion $\alpha$ in Figure 1 scenario for different types of data processing `poolfstat` analyses.** For each MAF threshold (MAF> 1% or MAF> 5%), the table gives the mean of the estimated $\hat{\alpha} = \frac{f_4(P1,P5;P3,P6)}{f_4(P1,P5;P2,P3))}$ (across 250 independent simulated datasets) for 21 different types of analyses relying on i) allele count data; ii) 15 different Pool-Seq read count data (five mean coverages $\lambda$ and three sequencing error rates $\epsilon$); and iii) Pool-Seq read count data simulated with $\epsilon = 0$ treated as allele counts (corresponding results of this bad practice are highlighted in italics and $^{\star}$). The proportion (in %) of the 250 estimated 95% confidence intervals that contain the true simulated value ($\alpha = 0.25$) is given in parenthesis.

| MAF threshold | seq. error $\epsilon$ | Pool-Seq (read counts) data | | | | | allele count data |
|---|---|---|---|---|---|---|---|
| | | $\lambda = 30$ | $\lambda = 50$ | $\lambda = 75$ | $\lambda = 100$ | $\lambda = 200$ | |
| >1% | 0 | 0.247 (44.8) | 0.247 (44.0) | 0.247 (43.6) | 0.246 (42.0) | 0.247 (42.4) | 0.246 (42.8) |
| | 1‰ | 0.247 (41.6) | 0.247 (43.2) | 0.246 (44.4) | 0.247 (43.2) | 0.246 (44.4) | |
| | 2.5‰ | 0.251 (74.4) | 0.252 (56.8) | 0.247 (42.8) | 0.246 (40.4) | 0.247 (42.0) | |
| | *0* | *0.247 (44.8)*⋆ | *0.247 (44.0)*⋆ | *0.247 (43.6)*⋆ | *0.246 (42.0)*⋆ | *0.247 (42.4)*⋆ | |
| >5% | 0 | 0.247 (45.6) | 0.247 (43.2) | 0.247 (43.2) | 0.246 (41.6) | 0.247 (42.0) | 0.246 (42.8) |
| | 1‰ | 0.247 (42.8) | 0.247 (40.8) | 0.246 (44.0) | 0.247 (44.4) | 0.247 (42.8) | |
| | 2.5‰ | 0.246 (44.0) | 0.246 (43.6) | 0.247 (42.4) | 0.247 (43.2) | 0.247 (41.6) | |
| | *0* | *0.247 (45.2)*⋆ | *0.247 (43.2)*⋆ | *0.247 (43.2)*⋆ | *0.246 (41.6)*⋆ | *0.247 (41.6)*⋆ | |

**Table S5. Comparison of the estimation of the simulated admixture proportion $\alpha$ in Figure 1 scenario using admixture graph fitting (as implemented in the `fit.graph` function of `poolfstat`) for different types of data processing `poolfstat` analyses.** For each MAF threshold (MAF> 1% or MAF> 5%), the table gives the mean of the estimated $\hat{\alpha}$ (across 250 independent simulated datasets) for 21 different types of analyses relying on i) allele count data; ii) 15 different Pool-Seq read count data (five mean coverages $\lambda$ and three sequencing error rates $\epsilon$); and iii) Pool-Seq read count data simulated with $\epsilon = 0$ treated as allele counts (corresponding results of this bad practice are highlighted in italics and ⋆). The proportion (in %) of the 250 estimated 95% confidence intervals that contain the true simulated value ($\alpha = 0.25$) is also given in parenthesis.

| MAF threshold | seq. error $\epsilon$ | Pool-Seq (read counts) data | | | | | allele count data |
|---|---|---|---|---|---|---|---|
| | | $\lambda = 30$ | $\lambda = 50$ | $\lambda = 75$ | $\lambda = 100$ | $\lambda = 200$ | |
| >1% | 0 | $5.1{\times}10^{-2}$ (86.0) | $5.1{\times}10^{-2}$ (91.2) | $5.1{\times}10^{-2}$ (91.2) | $5.1{\times}10^{-2}$ (92.0) | $5.1{\times}10^{-2}$ (92.4) | $5.1{\times}10^{-2}$ (91.6) |
| | 1‰ | $4.4{\times}10^{-2}$ (0.40) | $4.1{\times}10^{-2}$ (0.00) | $5.1{\times}10^{-2}$ (94.4) | $5.1{\times}10^{-2}$ (94.0) | $5.1{\times}10^{-2}$ (95.6) | |
| | 2.5‰ | $6.0{\times}10^{-3}$ (0.00) | 1.1e-03 (0.00) | $4.3{\times}10^{-2}$ (0.80) | $4.9{\times}10^{-2}$ (90.0) | $5.0{\times}10^{-2}$ (97.2) | |
| | *0* | *$7.2{\times}10^{-2}$ (0.00)*⋆ | *$7.2{\times}10^{-2}$ (0.00)*⋆ | *$7.2{\times}10^{-2}$ (0.00)*⋆ | *$7.2{\times}10^{-2}$ (0.00)*⋆ | *$7.2{\times}10^{-2}$ (0.00)*⋆ | |
| >5% | 0 | $5.1{\times}10^{-2}$ (81.6) | $5.1{\times}10^{-2}$ (85.6) | $5.1{\times}10^{-2}$ (89.6) | $5.1{\times}10^{-2}$ (89.2) | $5.1{\times}10^{-2}$ (91.2) | $5.0{\times}10^{-2}$ (90.4) |
| | 1‰ | $5.1{\times}10^{-2}$ (84.0) | $5.1{\times}10^{-2}$ (88.8) | $5.1{\times}10^{-2}$ (89.6) | $5.1{\times}10^{-2}$ (91.6) | $5.0{\times}10^{-2}$ (92.0) | |
| | 2.5‰ | $5.1{\times}10^{-2}$ (87.2) | $5.0{\times}10^{-2}$ (90.0) | $5.0{\times}10^{-2}$ (92.4) | $5.0{\times}10^{-2}$ (90.0) | $5.0{\times}10^{-2}$ (90.8) | |
| | *0* | *$7.2{\times}10^{-2}$ (0.00)*⋆ | *$7.2{\times}10^{-2}$ (0.00)*⋆ | *$7.2{\times}10^{-2}$ (0.00)*⋆ | *$7.1{\times}10^{-2}$ (0.00)*⋆ | *$7.1{\times}10^{-2}$ (0.00)*⋆ | |

**Table S6. Comparison of the estimation of the simulated length (in drift units) for the P7↔P1 branch $\left(\tau_{P7\leftrightarrow P1} = 5 \times 10^{-2}\right)$ in Figure 1 scenario using admixture graph fitting (as implemented in the `fit.graph` function of `poolfstat`) for different types of data processing `poolfstat` analyses.** For each MAF threshold (MAF> 1% or MAF> 5%), the table gives the mean of the estimated $\hat{\tau}_{P7\leftrightarrow P1}$ (across 250 independent simulated datasets) for 21 different types of analyses relying on i) allele count data; ii) 15 different Pool-Seq read count data (five mean coverages $\lambda$ and three sequencing error rates $\epsilon$); and iii) Pool-Seq read count data simulated with $\epsilon = 0$ treated as allele counts (corresponding results of this bad practice are highlighted in italics and ⋆). The proportion (in %) of the 250 estimated 95% confidence intervals that contain the true simulated value is given in parenthesis.

| MAF threshold | seq. error $\epsilon$ | Pool-Seq (read counts) data | | | | | allele count data |
|---|---|---|---|---|---|---|---|
| | | $\lambda = 30$ | $\lambda = 50$ | $\lambda = 75$ | $\lambda = 100$ | $\lambda = 200$ | |
| >1% | 0 | $2.5\times10^{-2}$ (44.8) | $2.5\times10^{-2}$ (49.2) | $2.5\times10^{-2}$ (51.2) | $2.5\times10^{-2}$ (50.8) | $2.5\times10^{-2}$ (52.0) | $2.5\times10^{-2}$ (50.8) |
| | 1‰ | $2.0\times10^{-2}$ (32.4) | $1.9\times10^{-2}$ (12.0) | $2.4\times10^{-2}$ (48.8) | $2.5\times10^{-2}$ (47.6) | $2.4\times10^{-2}$ (51.6) | |
| | 2.5‰ | 0.00 (0.00) | 0.00 (0.00) | $2.0\times10^{-2}$ (25.6) | $2.4\times10^{-2}$ (49.6) | $2.4\times10^{-2}$ (48.8) | |
| | *0* | *$4.5\times10^{-2}$ (0.40)*$^\star$ | *$4.4\times10^{-2}$ (0.40)*$^\star$ | *$4.4\times10^{-2}$ (0.00)*$^\star$ | *$4.4\times10^{-2}$ (0.40)*$^\star$ | *$4.4\times10^{-2}$ (0.40)*$^\star$ | |
| >5% | 0 | $2.5\times10^{-2}$ (47.2) | $2.5\times10^{-2}$ (48.8) | $2.4\times10^{-2}$ (46.4) | $2.4\times10^{-2}$ (45.6) | $2.4\times10^{-2}$ (46.8) | $2.4\times10^{-2}$ (49.2) |
| | 1‰ | $2.5\times10^{-2}$ (44.0) | $2.5\times10^{-2}$ (46.4) | $2.4\times10^{-2}$ (47.2) | $2.4\times10^{-2}$ (47.2) | $2.4\times10^{-2}$ (48.4) | |
| | 2.5‰ | $2.4\times10^{-2}$ (44.8) | $2.4\times10^{-2}$ (47.6) | $2.4\times10^{-2}$ (46.0) | $2.4\times10^{-2}$ (46.8) | $2.4\times10^{-2}$ (46.4) | |
| | *0* | *$4.4\times10^{-2}$ (0.80)*$^\star$ | *$4.4\times10^{-2}$ (0.40)*$^\star$ | *$4.4\times10^{-2}$ (0.00)*$^\star$ | *$4.4\times10^{-2}$ (0.00)*$^\star$ | *$4.4\times10^{-2}$ (0.00)*$^\star$ | |

**Table S7. Comparison of the estimation of the simulated length (in drift units) for the P2↔S1 branch $\left(\tau_{P2\leftrightarrow S1} = 2.5\times10^{-2}\right)$ in Figure 1 scenario using admixture graph fitting (as implemented in the `fit.graph` function of `poolfstat`) for different types of data processing `poolfstat` analyses.** For each MAF threshold (MAF> 1% or MAF> 5%), the table gives the mean of the estimated $\hat{\tau}_{P2\leftrightarrow S1}$ (across 250 independent simulated datasets) for 21 different types of analyses relying on i) allele count data; ii) 15 different Pool-Seq read count data (five mean coverages $\lambda$ and three sequencing error rates $\epsilon$); and iii) Pool-Seq read count data simulated with $\epsilon = 0$ treated as allele counts (corresponding results of this bad practice are highlighted in italics and $^\star$). The proportion (in %) of the 250 estimated 95% confidence intervals that contain the true simulated value is given in parenthesis.

| MAF threshold | seq. error $\epsilon$ | Pool-Seq (read counts) data | | | | | allele count data |
|---|---|---|---|---|---|---|---|
| | | $\lambda = 30$ | $\lambda = 50$ | $\lambda = 75$ | $\lambda = 100$ | $\lambda = 200$ | |
| >1% | 0 | $2.5\times10^{-2}$ (65.6) | $2.5\times10^{-2}$ (70.8) | $2.5\times10^{-2}$ (70.8) | $2.5\times10^{-2}$ (68.8) | $2.5\times10^{-2}$ (68.8) | $2.5\times10^{-2}$ (70.0) |
| | 0.10% | $2.1\times10^{-2}$ (4.80) | $1.9\times10^{-2}$ (0.00) | $2.5\times10^{-2}$ (70.0) | $2.5\times10^{-2}$ (72.8) | $2.5\times10^{-2}$ (70.0) | |
| | 0.25% | 0.0 (0.00) | 0.0 (0.00) | $2.0\times10^{-2}$ (5.6) | $2.4\times10^{-2}$ (57.2) | $2.5\times10^{-2}$ (71.2) | |
| | *0* | *$4.5\times10^{-2}$ (0.00)*$^\star$ | *$4.5\times10^{-2}$ (0.00)*$^\star$ | *$4.5\times10^{-2}$ (0.00)*$^\star$ | *$4.5\times10^{-2}$ (0.00)*$^\star$ | *$4.5\times10^{-2}$ (0.00)*$^\star$ | |
| >5% | 0 | $2.5\times10^{-2}$ (62.0) | $2.5\times10^{-2}$ (62.4) | $2.5\times10^{-2}$ (62.0) | $2.5\times10^{-2}$ (62.4) | $2.5\times10^{-2}$ (62.8) | $2.5\times10^{-2}$ (60.8) |
| | 0.10% | $2.5\times10^{-2}$ (66.4) | $2.5\times10^{-2}$ (61.2) | $2.5\times10^{-2}$ (61.2) | $2.5\times10^{-2}$ (59.6) | $2.5\times10^{-2}$ (61.6) | |
| | 0.25% | $2.5\times10^{-2}$ (66.4) | $2.5\times10^{-2}$ (63.6) | $2.5\times10^{-2}$ (61.6) | $2.5\times10^{-2}$ (61.2) | $2.5\times10^{-2}$ (61.2) | |
| | *0* | *$4.5\times10^{-2}$ (0.00)*$^\star$ | *$4.5\times10^{-2}$ (0.00)*$^\star$ | *$4.5\times10^{-2}$ (0.00)*$^\star$ | *$4.4\times10^{-2}$ (0.00)*$^\star$ | *$4.4\times10^{-2}$ (0.00)*$^\star$ | |

**Table S8. Comparison of the estimation of the simulated length (in drift units) for the P6↔S branch $\left(\tau_{P6\leftrightarrow S} = 2.5\times10^{-2}\right)$ in Figure 1 scenario using admixture graph fitting (as implemented in the `fit.graph` function of `poolfstat`) for different types of data processing `poolfstat` analyses.** For each MAF threshold (MAF> 1% or MAF> 5%), the table gives the mean of the estimated $\hat{\tau}_{P6\leftrightarrow S}$ (across 250 independent simulated datasets) for 21 different types of analyses relying on i) allele count data; ii) 15 different Pool-Seq read count data (five mean coverages $\lambda$ and three sequencing error rates $\epsilon$); and iii) Pool-Seq read count data simulated with $\epsilon = 0$ treated as allele counts (corresponding results of this bad practice are highlighted in italics and $^\star$). The proportion (in %) of the 250 estimated 95% confidence intervals that contain the true simulated value is given in parenthesis.

| MAF threshold | seq. error $\epsilon$ | Pool-Seq (read counts) data | | | | | allele count data |
|---|---|---|---|---|---|---|---|
| | | $\lambda = 30$ | $\lambda = 50$ | $\lambda = 75$ | $\lambda = 100$ | $\lambda = 200$ | |
| >1% | 0 | $2.6\times10^{-2}$ (54.4) | $2.6\times10^{-2}$ (56.0) | $2.6\times10^{-2}$ (59.6) | $2.6\times10^{-2}$ (59.2) | $2.6\times10^{-2}$ (57.6) | $2.6\times10^{-2}$ (56.4) |
| | 1‰ | $2.1\times10^{-2}$ (28.8) | $1.9\times10^{-2}$ (10.0) | $2.6\times10^{-2}$ (57.6) | $2.6\times10^{-2}$ (61.6) | $2.6\times10^{-2}$ (59.6) | |
| | 2.5‰ | 0.00 (0.00) | 0.00 (0.00) | $2.1\times10^{-2}$ (25.2) | $2.5\times10^{-2}$ (55.2) | $2.5\times10^{-2}$ (58.8) | |
| | *0* | *$4.5\times10^{-2}$ (0.00)*$^\star$ | *$4.5\times10^{-2}$ (0.00)*$^\star$ | *$4.5\times10^{-2}$ (0.00)*$^\star$ | *$4.5\times10^{-2}$ (0.00)*$^\star$ | *$4.5\times10^{-2}$ (0.00)*$^\star$ | |
| >5% | 0 | $2.6\times10^{-2}$ (52.0) | $2.6\times10^{-2}$ (52.8) | $2.5\times10^{-2}$ (54.0) | $2.6\times10^{-2}$ (56.4) | $2.5\times10^{-2}$ (54.0) | $2.5\times10^{-2}$ (54.8) |
| | 1‰ | $2.6\times10^{-2}$ (53.2) | $2.5\times10^{-2}$ (51.2) | $2.5\times10^{-2}$ (54.0) | $2.5\times10^{-2}$ (54.4) | $2.5\times10^{-2}$ (54.4) | |
| | 2.5‰ | $2.6\times10^{-2}$ (56.8) | $2.5\times10^{-2}$ (54.4) | $2.5\times10^{-2}$ (54.0) | $2.5\times10^{-2}$ (50.8) | $2.5\times10^{-2}$ (53.6) | |
| | *0* | *$4.5\times10^{-2}$ (0.00)*$^\star$ | *$4.5\times10^{-2}$ (0.00)*$^\star$ | *$4.5\times10^{-2}$ (0.00)*$^\star$ | *$4.5\times10^{-2}$ (0.00)*$^\star$ | *$4.5\times10^{-2}$ (0.00)*$^\star$ | |

**Table S9. Comparison of the estimation of the simulated length (in drift units) for the P3↔S2 branch $\left(\tau_{P3\leftrightarrow S2} = 2.5\times10^{-2}\right)$ in Figure 1 scenario using admixture graph fitting (as implemented in the `fit.graph` function of `poolfstat`) for different types of data processing `poolfstat` analyses.** For each MAF threshold (MAF> 1% or MAF> 5%), the table gives the mean of the estimated $\hat{\tau}_{P3\leftrightarrow S2}$ (across 250 independent simulated datasets) for 21 different types of analyses relying on i) allele count data; ii) 15 different Pool-Seq read count data (five mean coverages $\lambda$ and three sequencing error rates $\epsilon$); and iii) Pool-Seq read count data simulated with $\epsilon = 0$ treated as allele counts (corresponding results of this bad practice are highlighted in italics and $^\star$). The proportion (in %) of the 250 estimated 95% confidence intervals that contain the true simulated value is given in parenthesis.

| MAF threshold | seq. error $\epsilon$ | Pool-Seq (read counts) data | | | | | allele count data |
|---|---|---|---|---|---|---|---|
| | | $\lambda = 30$ | $\lambda = 50$ | $\lambda = 75$ | $\lambda = 100$ | $\lambda = 200$ | |
| >1% | 0 | 0.143 (20.0) | 0.141 (6.00) | 0.141 (5.60) | 0.141 (4.80) | 0.141 (4.00) | 0.140 (3.20) |
| | 1‰ | 0.126 (0.00) | 0.120 (0.00) | 0.140 (2.40) | 0.140 (3.20) | 0.140 (2.00) | |
| | 2.5‰ | $4.2\times10^{-2}$ (0.00) | $3.0\times10^{-2}$ (0.00) | 0.125 (0.00) | 0.137 (0.00) | 0.139 (0.00) | |
| | *0* | *0.160 (2.80)*⋆ | *0.158 (10.0)*⋆ | *0.158 (9.20)*⋆ | *0.158 (10.0)*⋆ | *0.158 (12.4)*⋆ | |
| >5% | 0 | 0.148 (85.6) | 0.147 (77.6) | 0.146 (74.0) | 0.146 (70.4) | 0.146 (65.6) | 0.145 (60.0) |
| | 1‰ | 0.147 (80.4) | 0.146 (73.6) | 0.146 (69.2) | 0.146 (67.2) | 0.145 (62.4) | |
| | 2.5‰ | 0.147 (77.6) | 0.146 (67.6) | 0.145 (62.4) | 0.145 (59.6) | 0.145 (56.4) | |
| | *0* | *0.165 (0.00)*⋆ | *0.164 (0.00)*⋆ | *0.163 (0.00)*⋆ | *0.163 (0.80)*⋆ | *0.163 (0.80)*⋆ | |

**Table S10. Comparison of the estimation of the simulated length (in drift units) for the P4↔P9 branch ($\tau_{P4\leftrightarrow P9} = 0.150$) in Figure 1 scenario using admixture graph fitting (as implemented in the `fit.graph` function of `poolfstat`) for different types of data processing `poolfstat` analyses.** For each MAF threshold (MAF> 1% or MAF> 5%), the table gives the mean of the estimated $\hat{\tau}_{P4\leftrightarrow P9}$ (across 250 independent simulated datasets) for 21 different types of analyses relying on i) allele count data; ii) 15 different Pool-Seq read count data (five mean coverages $\lambda$ and three sequencing error rates $\epsilon$); and iii) Pool-Seq read count data simulated with $\epsilon = 0$ treated as allele counts (corresponding results of this bad practice are highlighted in italics and ⋆). The proportion (in %) of the 250 estimated 95% confidence intervals that contain the true simulated value is given in parenthesis.

| MAF threshold | seq. error $\epsilon$ | Pool-Seq (read counts) data | | | | | allele count data |
|---|---|---|---|---|---|---|---|
| | | $\lambda = 30$ | $\lambda = 50$ | $\lambda = 75$ | $\lambda = 100$ | $\lambda = 200$ | |
| >1% | 0 | 0.142 (12.0) | 0.141 (2.80) | 0.140 (1.60) | 0.140 (1.20) | 0.140 (0.80) | 0.140 (0.40) |
| | 1‰ | 0.126 (0.00) | 0.120 (0.00) | 0.140 (0.40) | 0.140 (0.40) | 0.139 (0.40) | |
| | 2.5‰ | $4.2\times10^{-2}$ (0.00) | $3.0\times10^{-2}$ (0.00) | 0.125 (0.00) | 0.137 (0.00) | 0.139 (0.00) | |
| | *0* | *0.159 (1.60)*⋆ | *0.158 (7.20)*⋆ | *0.158 (8.00)*⋆ | *0.158 (9.60)*⋆ | *0.157 (11.2)*⋆ | |
| >5% | 0 | 0.147 (87.6) | 0.146 (77.6) | 0.146 (70.0) | 0.146 (64.4) | 0.145 (59.2) | 0.145 (53.2) |
| | 1‰ | 0.147 (82.0) | 0.146 (72.0) | 0.145 (60.4) | 0.145 (58.4) | 0.145 (54.0) | |
| | 2.5‰ | 0.146 (75.6) | 0.145 (61.6) | 0.145 (53.6) | 0.145 (52.8) | 0.144 (48.0) | |
| | *0* | *0.164 (0.00)*⋆ | *0.163 (0.00)*⋆ | *0.163 (0.00)*⋆ | *0.163 (0.00)*⋆ | *0.162 (0.00)*⋆ | |

**Table S11. Comparison of the estimation of the simulated length (in drift units) for the P5↔P9 branch ($\tau_{P5\leftrightarrow P9} = 0.150$) in Figure 1 scenario using admixture graph fitting (as implemented in the `fit.graph` function of `poolfstat`) for different types of data processing `poolfstat` analyses.** For each MAF threshold (MAF> 1% or MAF> 5%), the table gives the mean of the estimated $\hat{\tau}_{P5\leftrightarrow P9}$ (across 250 independent simulated datasets) for 21 different types of analyses relying on i) allele count data; ii) 15 different Pool-Seq read count data (five mean coverages $\lambda$ and three sequencing error rates $\epsilon$); and iii) Pool-Seq read count data simulated with $\epsilon = 0$ treated as allele counts (corresponding results of this bad practice are highlighted in italics and ⋆). The proportion (in %) of the 250 estimated 95% confidence intervals that contain the true simulated value is given in parenthesis.

| MAF threshold | seq. error $\epsilon$ | Pool-Seq (read counts) data | | | | | allele count data |
|---|---|---|---|---|---|---|---|
| | | $\lambda = 30$ | $\lambda = 50$ | $\lambda = 75$ | $\lambda = 100$ | $\lambda = 200$ | |
| >1% | 0 | $2.7\times10^{-2}$ (48.0) | $2.6\times10^{-2}$ (54.0) | $2.6\times10^{-2}$ (50.0) | $2.6\times10^{-2}$ (50.4) | $2.6\times10^{-2}$ (50.0) | $2.6\times10^{-2}$ (52.8) |
| | 1‰ | $2.4\times10^{-2}$ (42.0) | $2.3\times10^{-2}$ (36.4) | $2.6\times10^{-2}$ (48.8) | $2.6\times10^{-2}$ (51.6) | $2.6\times10^{-2}$ (50.8) | |
| | 2.5‰ | $5.8\times10^{-3}$ (0.00) | $8.9\times10^{-4}$ (0.00) | $2.3\times10^{-2}$ (51.2) | $2.6\times10^{-2}$ (51.2) | $2.6\times10^{-2}$ (49.2) | |
| | *0* | *$2.7\times10^{-2}$ (46.4)*$^\star$ | *$2.7\times10^{-2}$ (54.0)*$^\star$ | *$2.7\times10^{-2}$ (50.4)*$^\star$ | *$2.7\times10^{-2}$ (50.0)*$^\star$ | *$2.7\times10^{-2}$ (52.0)*$^\star$ | |
| >5% | 0 | $2.7\times10^{-2}$ (44.0) | $2.7\times10^{-2}$ (49.2) | $2.7\times10^{-2}$ (44.8) | $2.7\times10^{-2}$ (48.0) | $2.7\times10^{-2}$ (46.8) | $2.7\times10^{-2}$ (50.8) |
| | 1‰ | $2.7\times10^{-2}$ (46.4) | $2.7\times10^{-2}$ (52.8) | $2.7\times10^{-2}$ (47.6) | $2.7\times10^{-2}$ (50.0) | $2.7\times10^{-2}$ (46.8) | |
| | 2.5‰ | $2.7\times10^{-2}$ (46.4) | $2.7\times10^{-2}$ (44.4) | $2.6\times10^{-2}$ (47.2) | $2.6\times10^{-2}$ (48.8) | $2.6\times10^{-2}$ (46.8) | |
| | *0* | *$2.8\times10^{-2}$ (45.2)*$^\star$ | *$2.7\times10^{-2}$ (49.2)*$^\star$ | *$2.7\times10^{-2}$ (46.0)*$^\star$ | *$2.7\times10^{-2}$ (48.8)*$^\star$ | *$2.7\times10^{-2}$ (46.8)*$^\star$ | |

**Table S12. Comparison of the estimation of the simulated length (in drift units) for the S1↔P7 branch $\left(\tau_{S1\leftrightarrow P7} = 2.5 \times 10^{-2}\right)$ in Figure 1 scenario using admixture graph fitting (as implemented in the `fit.graph` function of `poolfstat`) for different types of data processing `poolfstat` analyses.** For each MAF threshold (MAF> 1% or MAF> 5%), the table gives the mean of the estimated $\hat{\tau}_{S1\leftrightarrow P7}$ (across 250 independent simulated datasets) for 21 different types of analyses relying on i) allele count data; ii) 15 different Pool-Seq read count data (five mean coverages $\lambda$ and three sequencing error rates $\epsilon$); and iii) Pool-Seq read count data simulated with $\epsilon = 0$ treated as allele counts (corresponding results of this bad practice are highlighted in italics and $^\star$). The proportion (in %) of the 250 estimated 95% confidence intervals that contain the true simulated value is given in parenthesis.

| MAF threshold | seq. error $\epsilon$ | Pool-Seq (read counts) data | | | | | allele count data |
|---|---|---|---|---|---|---|---|
| | | $\lambda = 30$ | $\lambda = 50$ | $\lambda = 75$ | $\lambda = 100$ | $\lambda = 200$ | |
| >1% | 0 | $5.0\times10^{-2}$ (95.2) | $4.9\times10^{-2}$ (93.2) | $4.9\times10^{-2}$ (94.0) | $4.9\times10^{-2}$ (94.4) | $4.9\times10^{-2}$ (94.4) | $4.9\times10^{-2}$ (94.4) |
| | 1‰ | $4.4\times10^{-2}$ (7.60) | $4.3\times10^{-2}$ (0.80) | $4.9\times10^{-2}$ (92.8) | $4.9\times10^{-2}$ (94.8) | $4.9\times10^{-2}$ (93.2) | |
| | 2.5‰ | $1.9\times10^{-2}$ (0.00) | $1.5\times10^{-2}$ (0.00) | $4.4\times10^{-2}$ (3.60) | $4.8\times10^{-2}$ (81.6) | $4.9\times10^{-2}$ (92.0) | |
| | *0* | *$5.1\times10^{-2}$ (93.2)*$^\star$ | *$5.0\times10^{-2}$ (94.8)*$^\star$ | *$5.0\times10^{-2}$ (94.4)*$^\star$ | *$5.0\times10^{-2}$ (94.4)*$^\star$ | *$5.0\times10^{-2}$ (95.2)*$^\star$ | |
| >5% | 0 | $5.3\times10^{-2}$ (67.6) | $5.2\times10^{-2}$ (72.4) | $5.2\times10^{-2}$ (72.8) | $5.2\times10^{-2}$ (72.0) | $5.2\times10^{-2}$ (74.0) | $5.2\times10^{-2}$ (74.8) |
| | 1‰ | $5.2\times10^{-2}$ (72.8) | $5.2\times10^{-2}$ (73.6) | $5.2\times10^{-2}$ (73.6) | $5.2\times10^{-2}$ (73.6) | $5.2\times10^{-2}$ (76.0) | |
| | 2.5‰ | $5.2\times10^{-2}$ (72.8) | $5.2\times10^{-2}$ (76.4) | $5.2\times10^{-2}$ (76.4) | $5.2\times10^{-2}$ (76.4) | $5.2\times10^{-2}$ (80.0) | |
| | *0* | *$5.4\times10^{-2}$ (50.4)*$^\star$ | *$5.3\times10^{-2}$ (54.0)*$^\star$ | *$5.3\times10^{-2}$ (55.2)*$^\star$ | *$5.3\times10^{-2}$ (56.0)*$^\star$ | *$5.3\times10^{-2}$ (58.0)*$^\star$ | |

**Table S13. Comparison of the estimation of the simulated length (in drift units) for the P7↔P8 branch $\left(\tau_{P7\leftrightarrow P8} = 5.0 \times 10^{-2}\right)$ in Figure 1 scenario using admixture graph fitting (as implemented in the `fit.graph` function of `poolfstat`) for different types of data processing `poolfstat` analyses.** For each MAF threshold (MAF> 1% or MAF> 5%), the table gives the mean of the estimated $\hat{\tau}_{P7\leftrightarrow P8}$ (across 250 independent simulated datasets) for 21 different types of analyses relying on i) allele count data; ii) 15 different Pool-Seq read count data (five mean coverages $\lambda$ and three sequencing error rates $\epsilon$); and iii) Pool-Seq read count data simulated with $\epsilon = 0$ treated as allele counts (corresponding results of this bad practice are highlighted in italics and $^\star$). The proportion (in %) of the 250 estimated 95% confidence intervals that contain the true simulated value is given in parenthesis.

| MAF threshold | seq. error $\epsilon$ | Pool-Seq (read counts) data | | | | | allele count data |
|---|---|---|---|---|---|---|---|
| | | $\lambda = 30$ | $\lambda = 50$ | $\lambda = 75$ | $\lambda = 100$ | $\lambda = 200$ | |
| >1% | 0 | $7.4\times10^{-2}$ (71.6) | $7.4\times10^{-2}$ (68.0) | $7.4\times10^{-2}$ (69.6) | $7.4\times10^{-2}$ (68.8) | $7.4\times10^{-2}$ (69.2) | $7.4\times10^{-2}$ (67.6) |
| | 1‰ | $6.6\times10^{-2}$ (5.60) | $6.4\times10^{-2}$ (0.00) | $7.3\times10^{-2}$ (66.0) | $7.3\times10^{-2}$ (68.0) | $7.3\times10^{-2}$ (66.8) | |
| | 2.5‰ | $2.5\times10^{-2}$ (0.00) | $1.8\times10^{-2}$ (0.00) | $6.6\times10^{-2}$ (4.80) | $7.2\times10^{-2}$ (54.0) | $7.3\times10^{-2}$ (62.0) | |
| | *0* | *$7.6\times10^{-2}$ (76.0)*$^\star$ | *$7.5\times10^{-2}$ (77.2)*$^\star$ | *$7.5\times10^{-2}$ (77.6)*$^\star$ | *$7.5\times10^{-2}$ (76.0)*$^\star$ | *$7.5\times10^{-2}$ (76.4)*$^\star$ | |
| >5% | 0 | $7.7\times10^{-2}$ (65.6) | $7.7\times10^{-2}$ (67.6) | $7.7\times10^{-2}$ (69.6) | $7.7\times10^{-2}$ (70.4) | $7.7\times10^{-2}$ (71.6) | $7.7\times10^{-2}$ (71.6) |
| | 1‰ | $7.7\times10^{-2}$ (67.6) | $7.7\times10^{-2}$ (69.2) | $7.7\times10^{-2}$ (72.8) | $7.7\times10^{-2}$ (71.6) | $7.6\times10^{-2}$ (72.0) | |
| | 2.5‰ | $7.7\times10^{-2}$ (70.8) | $7.6\times10^{-2}$ (69.6) | $7.6\times10^{-2}$ (73.6) | $7.6\times10^{-2}$ (70.0) | $7.6\times10^{-2}$ (74.4) | |
| | *0* | *$7.9\times10^{-2}$ (52.4)*$^\star$ | *$7.9\times10^{-2}$ (54.4)*$^\star$ | *$7.8\times10^{-2}$ (58.4)*$^\star$ | *$7.8\times10^{-2}$ (60.8)*$^\star$ | *$7.8\times10^{-2}$ (60.8)*$^\star$ | |

**Table S14. Comparison of the estimation of the simulated length (in drift units) for the S2↔P8 branch $\left(\tau_{S2\leftrightarrow P8} = 7.5 \times 10^{-2}\right)$ in Figure 1 scenario using admixture graph fitting (as implemented in the `fit.graph` function of `poolfstat`) for different types of data processing `poolfstat` analyses.** For each MAF threshold (MAF> 1% or MAF> 5%), the table gives the mean of the estimated $\hat{\tau}_{S2\leftrightarrow P8}$ (across 250 independent simulated datasets) for 21 different types of analyses relying on i) allele count data; ii) 15 different Pool-Seq read count data (five mean coverages $\lambda$ and three sequencing error rates $\epsilon$); and iii) Pool-Seq read count data simulated with $\epsilon = 0$ treated as allele counts (corresponding results of this bad practice are highlighted in italics and $^\star$). The proportion (in %) of the 250 estimated 95% confidence intervals that contain the true simulated value is given in parenthesis.

| MAF threshold | seq. error $\epsilon$ | Pool-Seq (read counts) data | | | | | allele count data |
|---|---|---|---|---|---|---|---|
| | | $\lambda = 30$ | $\lambda = 50$ | $\lambda = 75$ | $\lambda = 100$ | $\lambda = 200$ | |
| >1% | 0 | 0.155 (59.2) | 0.153 (81.2) | 0.153 (84.4) | 0.153 (84.0) | 0.153 (87.2) | 0.152 (90.0) |
| | 1‰ | 0.138 (0.40) | 0.132 (0.00) | 0.152 (88.4) | 0.152 (88.4) | 0.152 (92.0) | |
| | 2.5‰ | $5.62{\times}10^{-2}$ (0.00) | $4.49{\times}10^{-2}$ (0.00) | 0.137 (0.00) | 0.149 (94.8) | 0.151 (96.4) | |
| | *0* | *0.158 (19.2)*⋆ | *0.156 (43.6)*⋆ | *0.156 (45.6)*⋆ | *0.156 (47.6)*⋆ | *0.156 (54.8)*⋆ | |
| >5% | 0 | 0.177 (0.00) | 0.177 (0.00) | 0.177 (0.00) | 0.177 (0.00) | 0.177 (0.00) | 0.178 (0.00) |
| | 1‰ | 0.176 (0.00) | 0.176 (0.00) | 0.177 (0.00) | 0.177 (0.00) | 0.177 (0.00) | |
| | 2.5‰ | 0.175 (0.00) | 0.176 (0.00) | 0.176 (0.00) | 0.176 (0.00) | 0.176 (0.00) | |
| | *0* | *0.18 (0.00)*⋆ | *0.18 (0.00)*⋆ | *0.18 (0.00)*⋆ | *0.181 (0.00)*⋆ | *0.181 (0.00)*⋆ | |

**Table S15. Comparison of the estimation of the simulated length (in drift units) for the P8↔P9 branch ($\tau_{P8\leftrightarrow P9} = 0.150$) that combines the two branches from the root *R* in Figure 1 scenario using admixture graph fitting (as implemented in the `fit.graph` function of `poolfstat`) for different types of data processing `poolfstat` analyses.** For each MAF threshold (MAF> 1% or MAF> 5%), the table gives the mean of the estimated $\hat{\tau}_{P8\leftrightarrow P9}$ (across 250 independent simulated datasets) for 21 different types of analyses relying on i) allele count data; ii) 15 different Pool-Seq read count data (five mean coverages $\lambda$ and three sequencing error rates $\epsilon$); and iii) Pool-Seq read count data simulated with $\epsilon = 0$ treated as allele counts (corresponding results of this bad practice are highlighted in italics and ⋆). The proportion (in %) of the 250 estimated 95% confidence intervals that contain the true simulated value is given in parenthesis.

| MAF threshold | seq. error $\epsilon$ | Pool-Seq (read counts) data | | | | | allele count data |
|---|---|---|---|---|---|---|---|
| | | $\lambda = 30$ | $\lambda = 50$ | $\lambda = 75$ | $\lambda = 100$ | $\lambda = 200$ | |
| >1% | 0 | 100 (23.5) | 100 (28.0) | 100 (29.7) | 100 (28.8) | 100 (33.8) | 100 (33.2) |
| | 1‰ | 100 (28.5) | 100 (31.4) | 100 (31.7) | 100 (31.9) | 100 (33.9) | |
| | 2.5‰ | 100 (29.9) | 42.4 (0.00) | 100 (31.2) | 100 (26.9) | 100 (26.4) | |
| | *0* | *100 (23.5)*⋆ | *100 (28.0)*⋆ | *100 (29.7)*⋆ | *100 (28.8)*⋆ | *100 (33.8)*⋆ | |
| >5% | 0 | 100 (17.3) | 100 (19.6) | 100 (24.5) | 100 (19.7) | 100 (23.3) | 100 (23.1) |
| | 1‰ | 100 (22.7) | 100 (24.8) | 100 (23.2) | 100 (22.4) | 100 (23.7) | |
| | 2.5‰ | 100 (19.8) | 100 (20.5) | 100 (23.2) | 100 (18.7) | 100 (19.3) | |
| | *0* | *100 (17.3)*⋆ | *100 (19.6)*⋆ | *100 (24.5)*⋆ | *100 (19.7)*⋆ | *100 (23.3)*⋆ | |

**Table S16. Performance of the `add.leaf` function in positioning the simulated population P6 on the underlying (((P1,P2),P3),(P4,P5)) tree (Figure 1) for different types of simulated data processing `poolfstat` analyses.** For each MAF threshold (MAF> 1% or MAF> 5%), the table gives the proportion of correctly inferred admixture graph (i.e., positioning of the P6 population as deriving from an admixture events between two populations directly ancestral to P2 and P3 with a $\Delta_{BIC} > 6$ support) across 250 independent simulated datasets for 21 different types of analyses relying on i) allele count data; ii) 15 different Pool-Seq read count data (five mean coverages $\lambda$ and three sequencing error rates $\epsilon$); and iii) Pool-Seq read count data simulated with $\epsilon = 0$ treated as allele counts (corresponding results of this bad practice are highlighted in italics and ⋆). Note that a total of 36 different positioning of P6 on the (((P1,P2),P3),(P4,P5)) rooted tree are evaluated for each call of `add.leaf` function. Indeed as the reference tree consists of eight branches, P6 may be connected with i) 9 non-admixed edges (connection to either one of the 8 branches or as an outgroup) or, ii) 27 admixed edges from two-way admixture events (27=$C_2^8$-1, an admixture event between the two branches from the root being not identifiable). The lowest $\Delta_{BIC}$ between the true graph and the 35 other possible graphs over the 250 different datasets is given in parenthesis for each type of analyses.

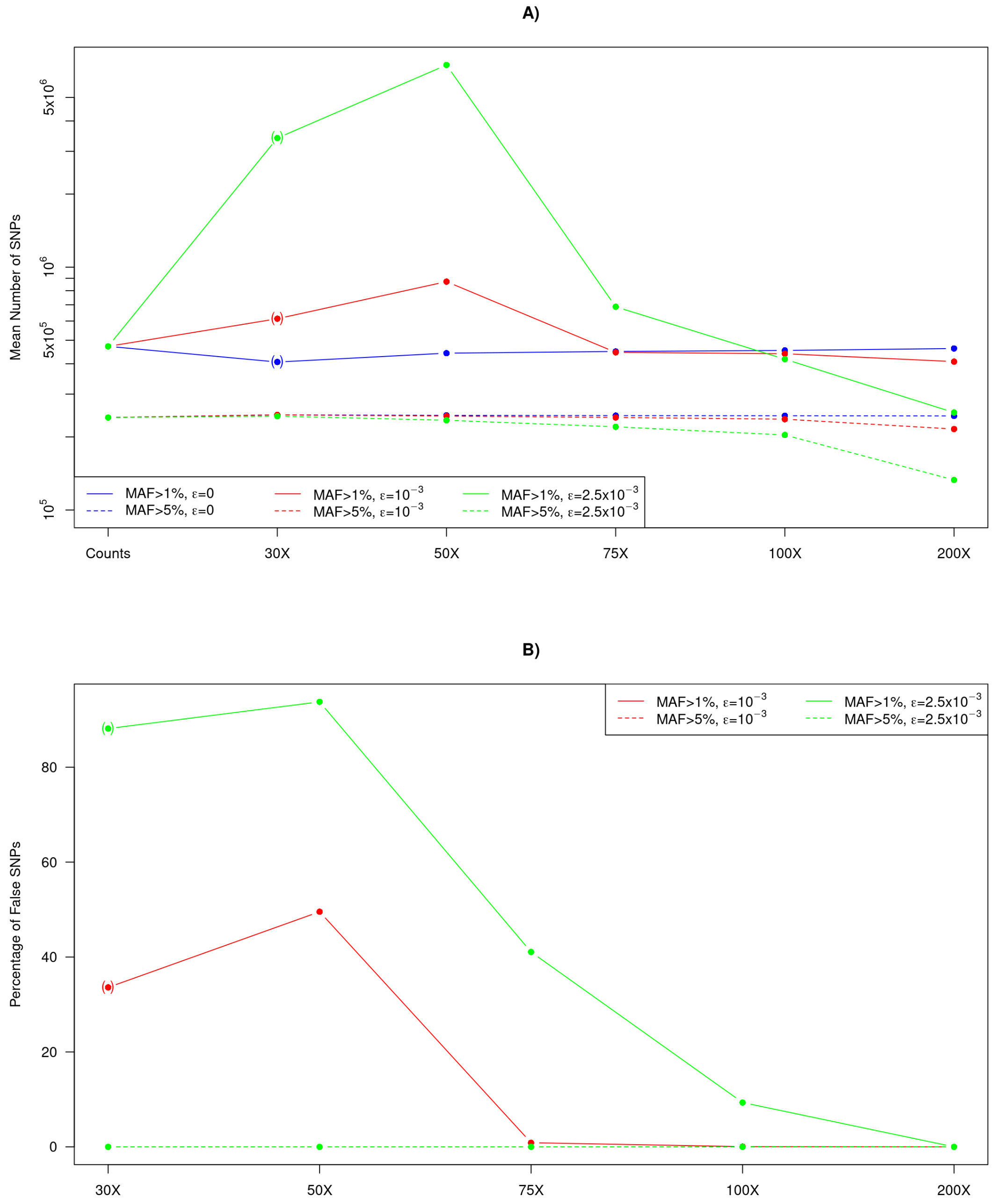


**Figure S1. Average number of SNPs (A) and percentage of false SNPs (B) in the simulated datasets as a function of read coverage and MAF filtering thresholds (see Table S1).** As highlighted with parentheses around the points in A and B, for Pool-Seq data with $\lambda = 30$, the actual MAF threshold is slightly higher than indicated due the additional filtering criterion imposed on the minimal read count (MRC>2; see Material and Methods). This is because for $\lambda = 30$, the overall coverage (that follows a Poisson distribution with parameter $6 \times 30 = 180$) is $\leq 200$ for on expected proportion of 93.5% of the SNPs resulting to a more stringent MAF threshold of $\frac{2}{6 \times 30} = 1.11\%$ when requiring MRC>2. This feature also explains the observed increased in the percentage of false observed from 30X to 50X coverage for the red (MAF>1% and $\epsilon = 10^{-3}$) and green (MAF>1% and $\epsilon = 2.5 \times 10^{-3}$) straight lines in B). Note also that the red (MAF>5% and $\epsilon = 10^{-3}$) and green (MAF>5% and $\epsilon = 2.5 \times 10^{-3}$) dashed lines are perfectly superposed in B) since no false SNP are included in all the corresponding configurations.

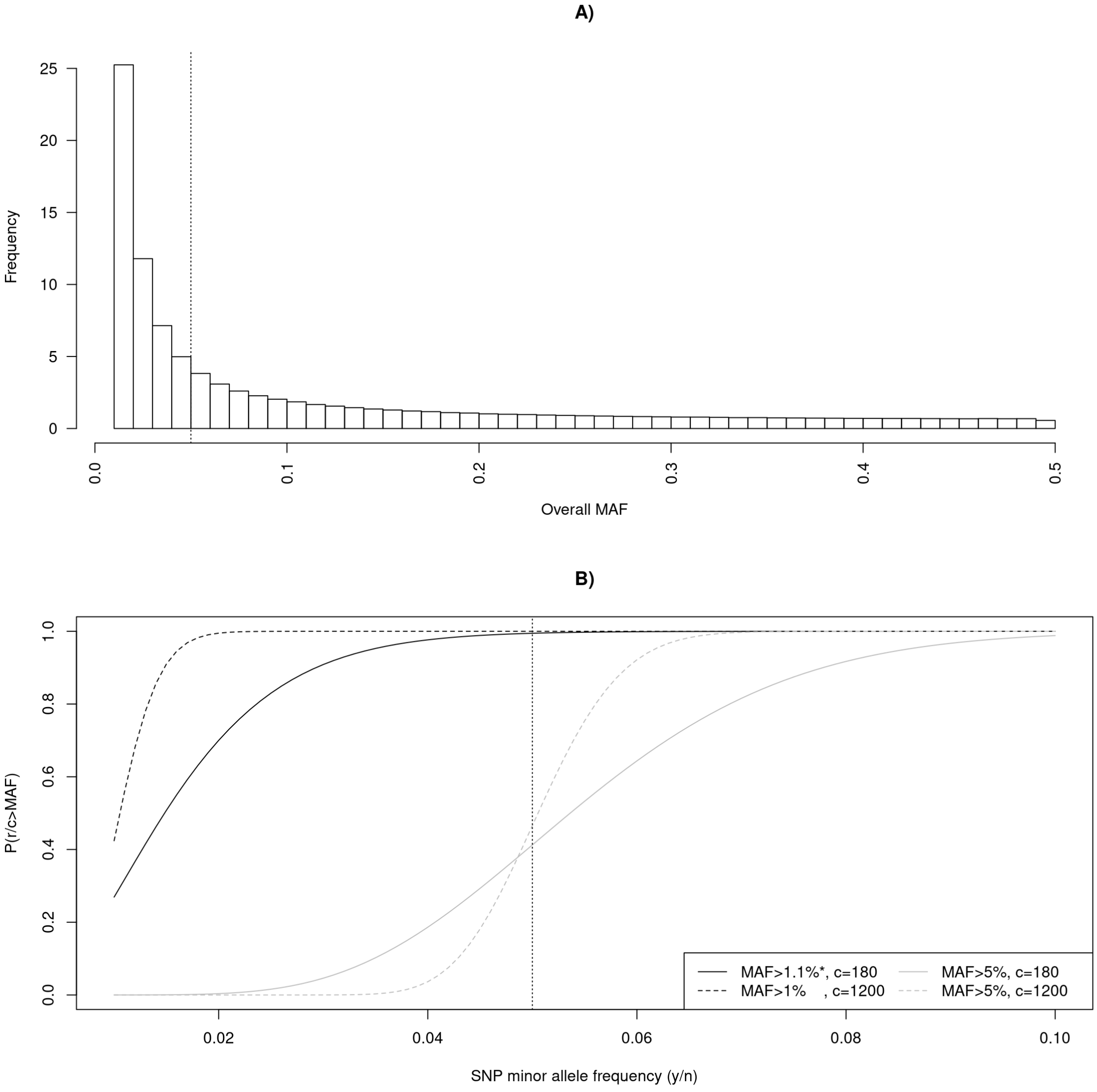


**Figure S2. Allele Frequency Spectrum of the simulated genotyping data (A) and read binomial sampling properties (B)**. A) Combined distribution of the overall SNP minor allele frequencies over all the 250 simulated genotyping datasets. B) Expected proportion of SNPs passing MAF filtering steps after binomial sampling of the minor read count $r$ (i.e., with $\frac{r}{c}$ >MAF, where $c$ is the overall read coverage) as a function of the SNP allele frequency $\frac{y}{n}$ (where $y$ is the minor allele count and $n$ is the haploid number of individuals) for two different coverage ($c = 30 \times 6 = 180$ and $c = 200 \times 6 = 1200$ corresponding to the two extremes of the simulated mean read coverage) and MAF thresholds (1% and 5%). The 5% MAF value is represented by a vertical dotted line in the two graphs.

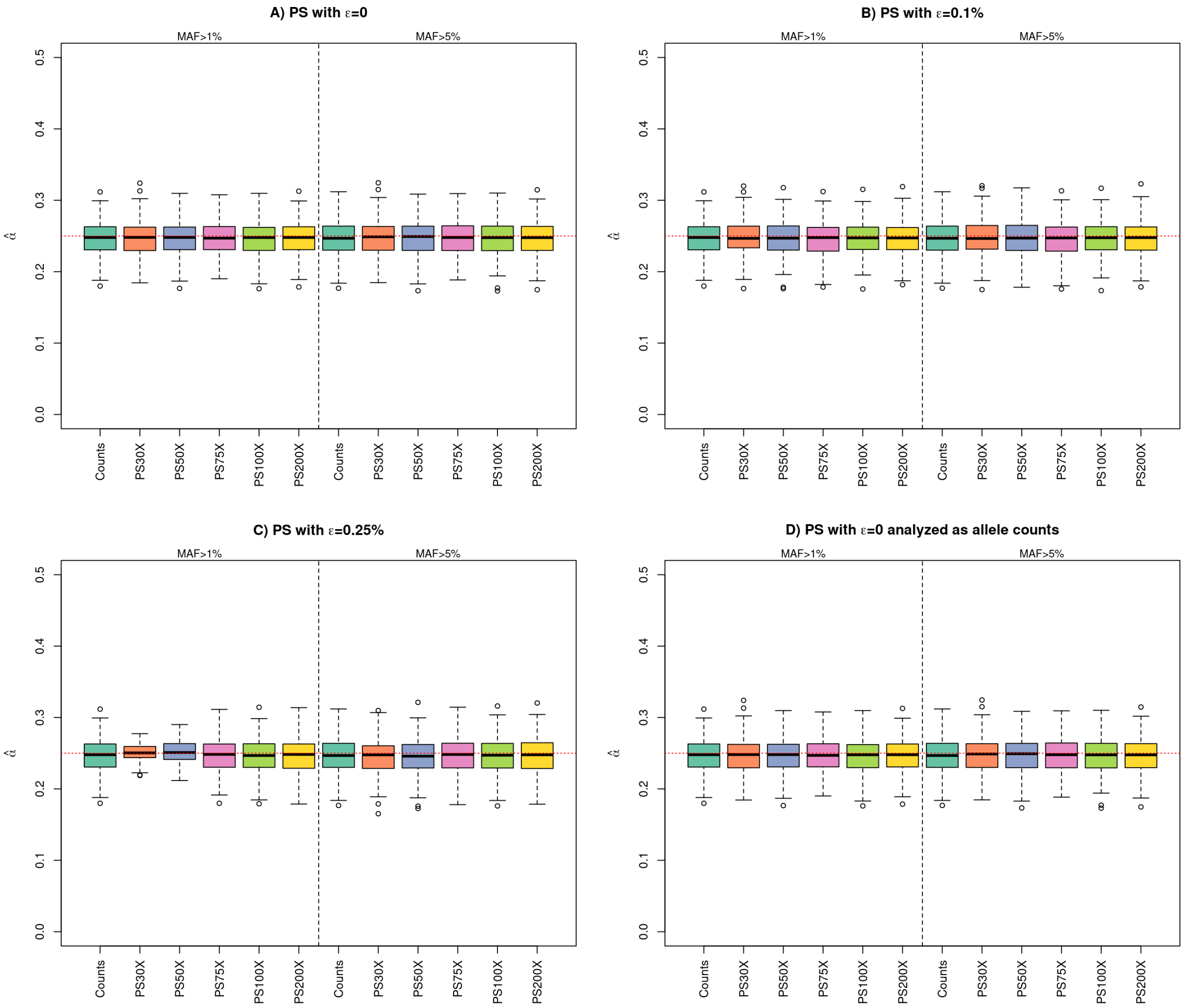


**Figure S3. Distribution of the estimated admixture proportion $\alpha = 0.25$ in the Figure 1 simulated scenario using admixture graph fitting (as implemented in the `fit.graph` function of `poolfstat`) for different types of data processing `poolfstat` analyses.** Each box plot summarizes the distribution of the 250 estimated values obtained from the analysis of either allele count dataset (the leftmost box named "Counts" of each group) or one of the five different simulated Pool-Seq read count datasets ("PS$\lambda$X") with different mean coverages ($\lambda = 30; 50; 75; 100;$ and 200) as generated from the genotyping data simulated under the scenario depicted in Figure 1. Pool-Seq read count data were generated with no sequencing errors ($\epsilon = 0$) in A) and D) and with a sequencing error rate of $\epsilon = 1‰$ and $\epsilon = 2.5‰$ in panel B) and C), respectively (Table S1). In D), the read count data were analyzed as allele counts which corresponds to a bad practice. In each of the four panels, analyses performed after discarding SNPs with an overall MAF (estimated from read counts in Pool-Seq data) ≤ 1% and ≤ 5%) are grouped in the left-hand and right-hand sides, respectively. Note that the two box plots obtained from the analysis of count data (MAF>1% and MAF>5%) are replicated in each panel for comparison purposes. The red dotted line indicates the simulated value of $\alpha$.

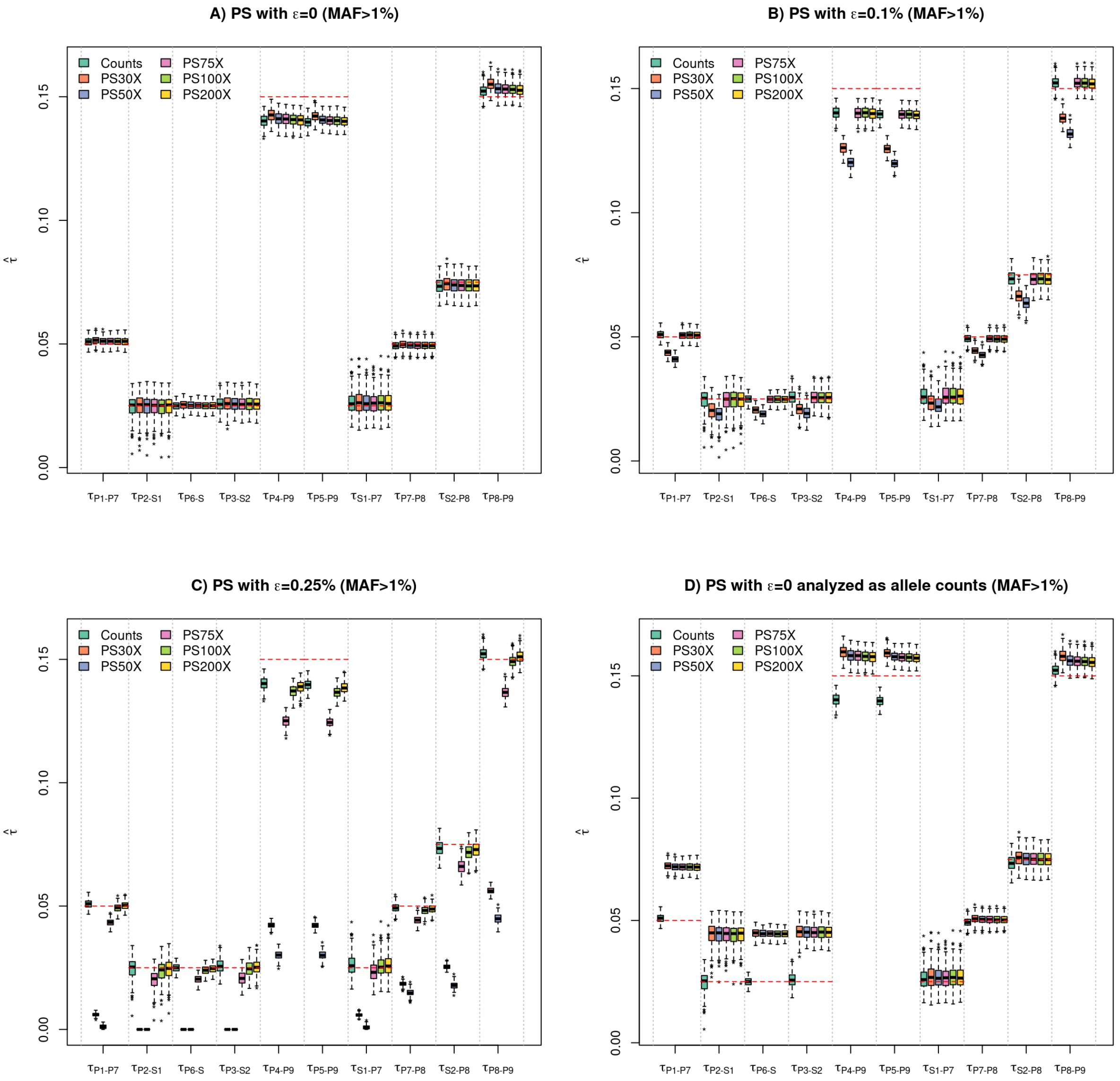


**Figure S4. Distribution of the estimated drift-scaled lengths for all the branches in Figure 1 simulated scenario using admixture graph fitting (as implemented in the `fit.graph` function of `poolfstat`) for different types of data with a 1% threshold on the overall SNP MAF.** Each box plot summarize the distribution of the 250 estimated lengths of each of the ten branches obtained from the analysis of either allele count dataset ("Counts") or one of the five different simulated Pool-Seq read count datasets ("PS$\lambda$X") with different mean coverages ($\lambda = 30; 50; 75; 100;$ and 200) as generated from the genotyping data simulated under the scenario depicted in Figure 1. Pool-Seq read count data were generated with no sequencing errors ($\epsilon = 0$) in A) and D) and with a sequencing error rate of $\epsilon = 1$‰ and $\epsilon = 2.5$‰ in panel B) and C), respectively (Table S1). In D), the read count data were analyzed as allele counts which corresponds to a bad practice. The two branches coming from the root are combined since the position of the root is not identifiable by the model (i.e., $\tau_{P8\leftrightarrow P9} = \tau_{P8\leftrightarrow R} + \tau_{P9\leftrightarrow R}$). The box plots obtained from the analysis of count data are replicated in each panel for comparison purposes. For each branch, a red dotted line indicates the underlying simulated value. For Pool-Seq data, the overall MAF was estimated from read counts.

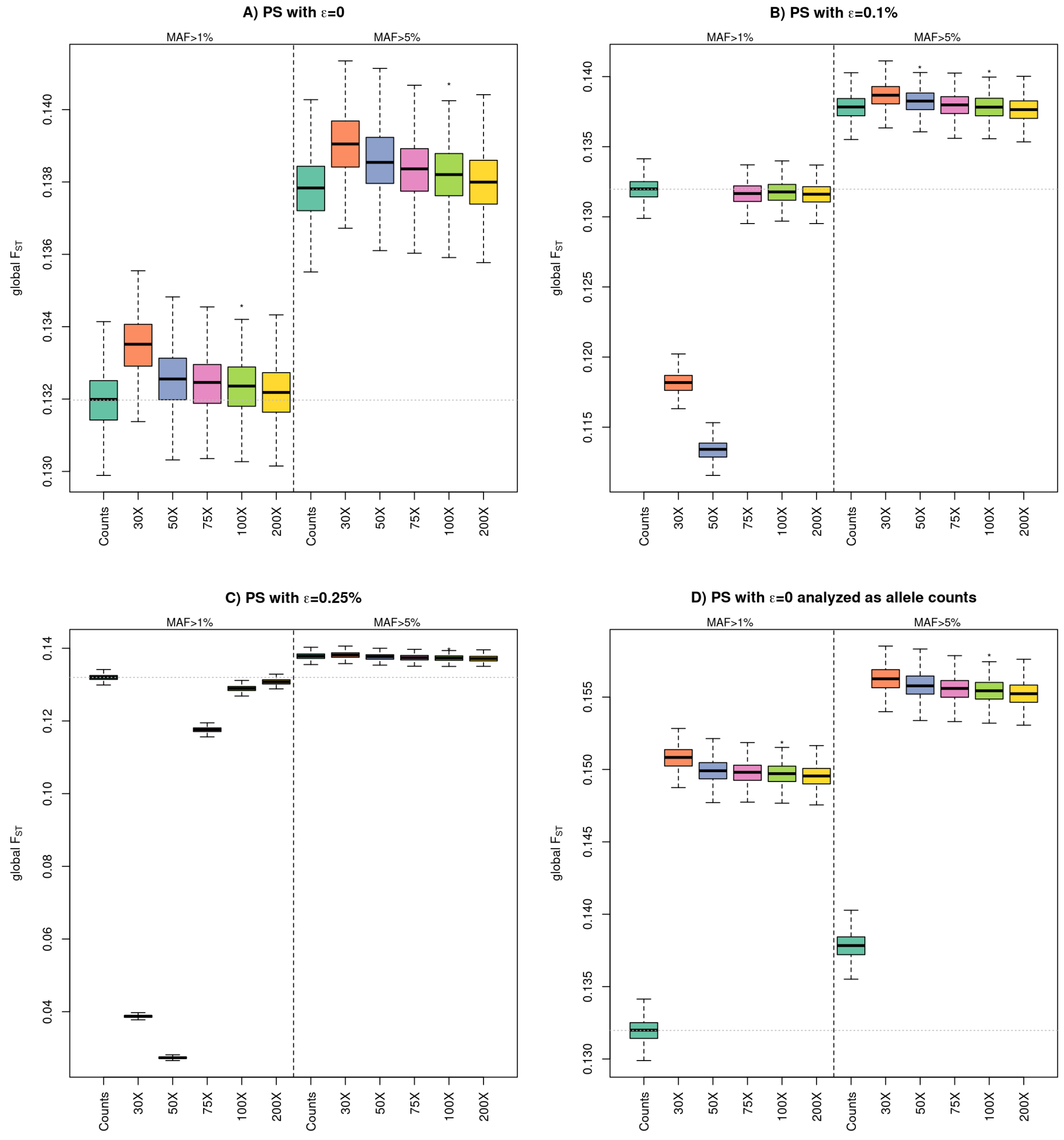


**Figure S5. Distribution of the estimated global $F_{ST}$ for the data simulated under the Figure 1 simulated scenario (using the `computeFST` function of `poolfstat` run with default option) for different types of data.** Each box plot summarizes the distribution of the 250 estimated values obtained from the analysis of either allele count dataset ("Counts") or one of the five different simulated Pool-Seq read count datasets ("PS$\lambda$X") with different mean coverages ($\lambda = 30; 50; 75; 100;$ and 200) as generated from the genotyping data simulated under the scenario depicted in Figure 1. Pool-Seq read count data were generated with no sequencing errors ($\epsilon = 0$) in A) and D) and with a sequencing error rate of $\epsilon = 1$‰ and $\epsilon = 2.5$‰ in panel B) and C) respectively (Table S1). In D), the read count data were analyzed as allele counts which corresponds to a bad practice. In each of the four panels, analyses performed after discarding SNPs with on the overall MAF (estimated from read counts in Pool-Seq data) ≤ 1% (respectively ≤ 5%) are grouped in the left-hand (respectively right-hand) side. The two box plots obtained from the analysis of count data (MAF>1% and MAF>5%) are replicated in each panel for comparison purposes and the mean estimated values for the allele count data with MAF>1% is indicated by a dotted line.
