## Supplementary Vignette V1 for "*f*-statistics estimation and admixture graph construction with Pool-Seq or allele count data using the R package *poolfstat*"

### Vignette for the package *poolfstat* (version 2+)

Mathieu Gautier

2021-05-26

**4 Estimating and visualizing $f$-statistics ($f_2$, $f_3$, $f_3^\star$, $f_4$ and D)** **10**

This vignette describes how the R package *poolfstat* can be used to compute various f- and D-statistics (estimation of $F_{ST}$, Patterson's $F_2$, $F_3$, $F_3^{\star}$, $F_4$ and $D$ parameters[1]) in population genomics studies from allele count or Pool-Seq read count data. The package also includes functions to fit and construct admixture graphs to infer the demographic history of populations based on the estimated $f$-statistics together with their visualization. This document is conceived as a hands-on tutorial providing users with an overview of the package functions with working example analyzing the Pool-Seq and allele count simulated data sets described in section 1 and publicly available for download from the Zenodo repository[2]. Details and (numerous) reference for the all the underlying methods are available in Gautier *et al.* (2021).

The *poolfstat* package is currently available for most platforms (Linux, MS Windows and MacOSX) from the CRAN repository (http://cran.r-project.org/) and may be installed using a standard procedure. Once the package has been successfully installed on your system, it can be loaded by:

```
library(poolfstat)
```

### 1 Preamble: presentation of the working example data set

Genetic data were simulated using the coalescent simulator *msprime* (Kelleher *et al.* 2016) for a total of 150 diploid individuals belonging to 6 different populations (n=25 per populations) that were historically related by the admixture graph represented in Figure 1. Each genome consisted of 20 independent chromosomes of $L = 100$ Mb assuming a scaled chromosome-wide recombination rate of $\rho = 4LN_e r = 4,000$[3]. Similarly, the scaled chromosome-wide mutation $\theta = 4LN_e\mu = 4,000$[4]. More precisely, the following *msprime* command was used:

```
mspms 300 20 -t 4000 -I 6 50 50 50 50 50 50 0 -es 0.0125 6 0.25 -ej 0.0125 6 2 -ej 0.0125
7 3 -ej 0.025 2 1 -ej 0.05 3 1 -ej 0.075 5 4 -ej 0.1 4 1 -r 4000 100000000 -p 8
```

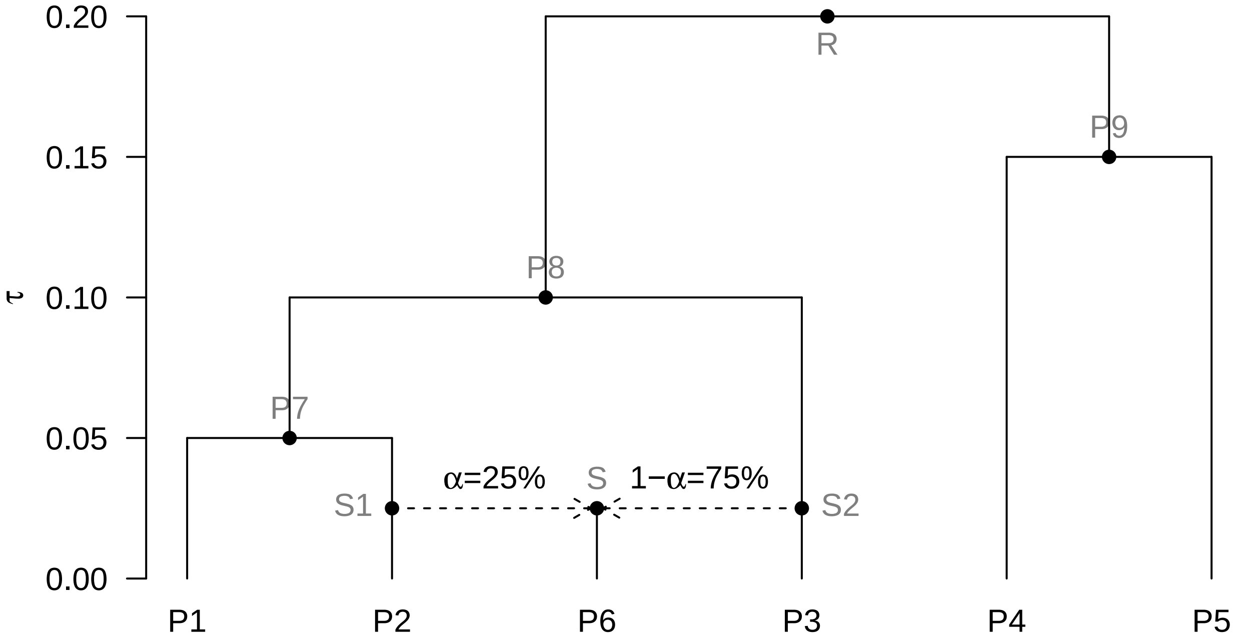

Figure 1: Simulated scenario relating the 6 populations of the working example. Names of the internal node populations for which no data is available are written in grey.

[1]Following Patterson *et al.* (2012), we use $F$ to refer to the parameter and $f$ to the statistics estimated from the data

[2]See http://doi.org/10.5281/zenodo.4709728

[3]For instance, $\rho = 4,000$ if the per-base and per-generation recombination rate $r = 10^{-8}$ (i.e., it the cM per Mb ratio is equal to 1) in a population of constant diploid effective size $N_e = 10^3$

[4]For instance, a nucleotide diversity of $\theta = 4,000$ is expected at mutation-drift equilibrium in a population of constant diploid effective size $N_e = 10^3$ if the per-base mutation rate is $\mu = 10^{-8}$

The simulation output was further parsed to remove all variants displaying a Minor Allele Frequency (MAF)[5] lower than 1% which lead to a total of 472,410 remaining SNPs (from 23,246 to 24,237 per chromosome) with position information stored in the file *snp6p.snpinfo.gz*[2]. From the resulting genotyping data, allele counts for both the ancestral and the derived alleles (which was taken as reference) were easily obtained by simple counting[6]. A Pool-Seq data set with no sequencing error was subsequently simulated from the allele count data as described in Gautier *et al.* (2021). Briefly, for each SNP $i$ in each population $j$, we sampled a read count $r_{ij}$ for the reference allele from a Binomial distribution following $r_{ij} \sim Bin\left(\frac{y_{ij}}{n_j}; c_{ij}\right)$ where $y_{ij}$ is the derived allele count for SNP $i$ in population $j$, $n_j$ is the (haploid) sample size of population $j$ (here $n_j = 50$ for all $j$) and $c_{ij}$ is the overall read coverage at SNP $i$ position[7]. Varying overall read coverages across pools and SNPs were simulated by sampling the $c_{ij}$'s from a Poisson distribution with mean $\lambda = 30$, i.e., assuming a read coverage of 30X for the different pools[8]. Two files representative of "real-life" data format were finally generated to store the simulate allele count and read count data:

- a file named *sim6p.genotreemix.gz*[2] containing allele count data for each SNP and each population in the same format as the one used in the population program *Treemix* (Pickrell and Pritchard 2012)
- a file named *sim6p.poolseq30X.vcf.gz*[2] containing read count data for each SNP and each population in a *vcf* format similar to the one obtained with the software *VarScan* (Koboldt *et al.* 2012)

### 2 Reading and manipulating input data

The package *poolfstat* includes several utilities to parse allele count or Pool-Seq read count input data stored in various standard formats. It is worth stressing that **distinguishing standard allele count data[9] from Pool-Seq read count data is critical** to rely on the appropriate $f$-statistics estimator (Hivert *et al.* 2018; Gautier *et al.* 2021). Hence, in the *poolfstat* package, two different S4 object classes are defined to store the data:

- the *countdata* S4 class for allele count data with elements (slots) detailed in the documentation page that can be accessed with the *help* function (or *?* operator) as follows:

```
help(countdata)
```

* the *pooldata* S4 class for (PoolSeq) read count data (PoolSeq data) with elements (slots) detailed in the documentation page that can be accessed with the *help* function (or *?* operator) as follows:

```
help(pooldata)
```

These classes characterize the type and origin of the data and are automatically detected by the *computeFST* (section 3.1.1), *compute.pairwiseFST* (section 3.2) and *compute.fstats* (section 4) functions that implement the different unbiased estimators thereby ensuring that the appropriate estimation procedure is used.

#### 2.1 Generating a countdata object for allele count data

A *countdata* object can be generated from allele count data stored in two different input file format:

- The input file format required by the popular *Treemix* program (Pickrell and Pritchard 2012) using the function *genotreemix2countdata*
- The input file format required by the program *BayPass* (Gautier 2015) for allele count data using the function *genobaypass2countdata*

The following example shows how to generate a *countdata* object (here named *sim6p.allelecount*) from the *sim6p.genotreemix.gz* example file that contains the allele count data (in a *Treemix* file format) for the simulated example data (section 1):

[5] MAF was estimated over all the 300 haploid individuals
[6] The simulated data being haploid this implicitly assumes Hardy-Weinberg equilibrium for the different populations
[7] Note that the read count for the alternate allele is simply $c_{ij} - r_{ij}$
[8] Such a coverage is actually in the lower limit of what is usually recommended for Pool-Seq experiment
[9] i.e., obtained from individual genotyping data

```
sim6p.allelecount<-genotreemix2countdata(genotreemix.file = "sim6p.genotreemix.gz")
```

Information about marker positions (chromosome or scaffold of origin of the markers and position on the chromosome) may be provided using the *snp.pos* argument of the *genotreemix2countdata* and *genobaypass2countdata* functions. For instance, to include the mapping information stored in the *snp6p.snpinfo.gz* example file (section 1), one may use the following commands[10]:

```
positions<-read.table("sim6p.snpinfo.gz",header=TRUE,row.names=1,stringsAsFactors = FALSE)
sim6p.allelecount<-genotreemix2countdata(genotreemix.file = "sim6p.genotreemix.gz",
                                         snp.pos=positions,verbose=FALSE)
sim6p.allelecount #display a summary of the object
```

```
 * * * Countdata Object * * *
 * Number of SNPs   =  472410
 * Number of Pops   =  6
 * Pop Names        :
 P1; P2; P3; P4; P5; P6
 * * * * * * * * * * * * * *
```

**Notice**

For operations requiring marker position information i.e., block-jackknife estimation of standard error (Appendix A.1) and estimation of multi-locus $F_{ST}$ on sliding windows over the genome (section 3.1.1), markers are always assumed to be ordered in the genome as their relative position in the input files. If no map information is provided, a default position is given to all the SNPs and they are all assumed to map to the same chromosome. In this case, some windows or blocks may then extend over two consecutive contigs[a].

[a]Providing the reference assembly used to order the markers is not too fragmented, the block-jackknife estimates of standard errors may only be marginally affected since the blocks are defined by a number of consecutive markers (and not by a physical length)

Additional arguments may allow filtering the data for low marker polymorphism levels (*min.maf*) or genotyping call rate (*min.indgeno.per.pop*). More details are available in the documentation pages of the *genotreemix2countdata* and *genobaypass2countdata* functions accessible with the commands `?genotreemix2pooldata` and `?genobaypasstreemix2pooldata`.

#### 2.2 Generating a pooldata object for Pool-Seq read count data

A **pooldata** object can be generated from Pool-Seq read count data stored in either of the four following formats:

- *vcf* file generated by most SNP calling software commonly used to analyze Pool-Seq data including *VarScan* (Koboldt *et al.* 2012), *bcftools/SAMtools* (Li *et al.* 2009), *GATK* (McKenna *et al.* 2010) or *FreeBayes* (Garrison and Marth 2012) using the *vcf2pooldata* function[11]
- *rsync* files generated by the *PoPoolation* software (Kofler *et al.* 2011) using the *popsync2pooldata* function
- The two input files (pool read count and pool haploid sizes) required by the program *BayPass* (Gautier 2015) to analyze Pool-Seq data using the *genobaypass2pooldata* function
- The input file required by the *SelEstim* program (Vitalis *et al.* 2014) to analyze Pool-Seq data using the *genoselestim2pooldata* function

[10]If the number of markers in the *snp.pos* object is not consistent with that of allele count file or the given matrix is not 2-column, default value for marker positions are given and a warning message is printed in the console

[11]The format of the *vcf* file is automatically detected from the genotype format that includes i) both an AD and RD fields for *VarScan vcf* files; or ii) only an AD field (with comma-separated read counts for the different allele) other than *VarScan vcf*. Parsing of *vcf* files has been substantially improved since *poolfstat* version *1.2* with computationally intensive text manipulation now implemented in *C++* routines inspired by those of the *vcfR* package (Knaus and Grünwald 2017)

The following example shows how to generate a *pooldata* object for the simulated Pool-Seq data contained in the *sim6p.poolseq30X.vcf.gz* vcf file (section 1).

```
sim6p.readcount30X <-vcf2pooldata(vcf.file="sim6p.poolseq30X.vcf.gz",poolsizes=rep(50,6))
```

```
Reading Header lines
Parsing allele counts
VarScan like format detected for allele count data: the AD field contains allele depth
for the alternate allele and RD field for the reference allele
(N.B., positions with more than one alternate allele will be ignored)
472410  lines processed in 0 h  0 m  3 s : 472410 SNPs found
Data consists of 472410 SNPs for 6 Pools
```

```
sim6p.readcount30X  #display a summary of the resulting pooldata object
```

```
 * * * PoolData Object * * *
 * Number of SNPs   =  472410
 * Number of Pools  =  6
 * Pool Names       :
 Pool1; Pool2; Pool3; Pool4; Pool5; Pool6
 * * * * * * * * * * * * * *
```

Additional arguments may allow filtering the data according to the read coverage of the pool (e.g., *min.maf*, *min.rc*, *min.cov.per.pool* or *max.cov.per.pool*). More details are available in the in the documentation pages of these different functions accessible with the commands `?vcf2pooldata`, `?popsync2pooldata`, `?genobaypass2pooldata` and `?genoselestim2pooldata` respectively.

#### 2.3 Manipulating *countdata* and *pooldata* objects

The function *pooldata.subset* (respectively *countdata.subset*) allows subsetting *pooldata* (resp. *countdata*) to retrieve just parts (e.g., some SNPs and/or population samples) of the original data sets according to various criteria (see `?pooldata.subset` or `?countdata.subset` for more details). In the example below, a *pooldata* object containing only information for populations *P2*, *P3* and *P6* for SNP with a MAF>0.05 is created from the *sim6p.readcount* object previously created:

```
sim6p.readcount30X.subset<-pooldata.subset(sim6p.readcount30X,pool.index=c(2,3,6),
                                           min.maf=0.05,verbose=FALSE)
sim6p.readcount30X.subset #display a summary of the resulting pooldata object
```

```
 * * * PoolData Object * * *
 * Number of SNPs   =  241280
 * Number of Pools  =  3
 * Pool Names       :
 Pool2; Pool3; Pool6
 * * * * * * * * * * * * * *
```

Note that indexes of the retained SNPs from the original data set may be obtained by setting the *return.snp.idx* argument to TRUE in the functions *pooldata.subset* (or *countdata.subset*). In this case the row of the matrix stored in the *snpinfo* slot of the *pooldata* (or *countdata*) output object are named *"rs"snp.idx* (where *snp.idx* is the SNP index for the original object) which makes it straightforward to obtain the indexes of the selected SNP as shown below:

```
sim6p.readcount30X.subset<-pooldata.subset(sim6p.readcount30X,pool.index=c(2,3,6),
                                           min.maf=0.05,return.snp.idx=TRUE,verbose=FALSE)
selected.snps.idx <- as.numeric(sub("rs","",rownames))
head(selected.snps.idx)
```

```
[1] 1 3 4 5 6 8
```

For *pooldata* objects, the functions *pooldata2genobaypass* and *pooldata2selestim* may also be used to generate input files for the aforementioned *BayPass* and *SelEstim* programs, including the generation of sub-samples of the original data (see `?pooldata2genobaypass` and `?pooldata2selestim` for more details).

### 3 Estimating $F_{ST}$

The $F_{ST}$ parameter is commonly used to quantify the structuring of the genetic diversity among populations (see e.g. Hivert *et al.* 2018 and references therein). It may be defined as:

$$F_{ST} \equiv \frac{Q_1 - Q_2}{1 - Q_2}$$

where $Q_1$ is the Identity In State (IIS) probability for genes sampled within populations (or pools), and $Q_2$ is the IIS probability for genes sampled between populations (or pools).

The *computeFST* and *compute.pairwiseFST* (for all pairs of populations) functions implement two different $F_{ST}$ estimators relying on:

- a decomposition of the total variance of allele or read count frequencies in an analysis-of-variance framework (Weir and Cockerham 1984) which is the default procedure of the functions (as specified by the argument *method="Anova"*). The implemented estimators are derived in Weir (1996) (eq. 5.2) (see also Akey *et al.* 2002) for allele count data (i.e., *countdata* objects, see 2.1); and in Hivert *et al.* (2018) (eq. 9) for (Pool-Seq) read count data (i.e., *pooldata* objects, see 2.2).
- unbiased estimators $\widehat{Q_1}$ and $\widehat{Q_2}$ of the IIS probabilities $Q_1$ and $Q_2$ (as specified by the argument *method="Identity"*). For allele count data (i.e., *countdata* objects, see 2.1) this estimator actually correspond to the one used by Karlsson *et al.* (2007). For Pool-Seq read count data (i.e., *pooldata* objects, see 2.2), the $\widehat{Q_1}$ of the $\widehat{Q_2}$ estimators are described in equations A39 and A43 respectively in Hivert *et al.* (2018) Supplementary Materials.

Note that multi-locus estimates (i.e., genome-wide estimates or sliding windows estimates) are derived as the sum of locus-specific numerators over the sum of locus-specific denominators of the different quantities (see, e.g., Hivert *et al.* 2018 and references therein).

#### 3.1 Estimating genome-wide $F_{ST}$ across all the populations

##### 3.1.1 The *computeFST* function

The *computeFST* function automatically uses the appropriate estimator given the input object class (either allele count for *countdata* objects or Pool-Seq read count data for *pooldata* objects). For instance with the simulated example data, we obtain the following estimates of $F_{ST}$ with:

- allele count data:

```
sim6p.allelecount.fst<-computeFST(sim6p.allelecount)
sim6p.allelecount.fst$FST      #genome-wide Fst over all populations
```

```
[1] 0.132319
```

- Pool-Seq read count data:

```
sim6p.readcount30X.fst<-computeFST(sim6p.readcount30X)
sim6p.readcount30X.fst$FST      #genome-wide Fst over all populations
```

```
[1] 0.1324199
```

Note that by default, the method *Anova* is implemented in the *computeFST* function which may be changed with the *method* argument of the function (see section 3).

###### 3.1.2 Block-Jackknife estimation of $\hat{F}_{ST}$ standard-error and confidence intervals:

Standard-error of the $F_{ST}$ estimates can be estimated using a block-jackknife sampling approach (see Appendix A.1) by specifying the number of consecutive SNPs defining a block with the argument *nsnp.per.bjack.block* (by default *nsnp.per.bjack.block=0*, i.e., no block-jackknife is carried out) as illustrated below for:

- allele count data:

```
sim6p.allelecount.fst<-computeFST(sim6p.allelecount,nsnp.per.bjack.block = 1000)
```

```
Starting Block-Jackknife sampling
462 Jackknife blocks identified with 462000 SNPs (out of 472410 ).
 SNPs map to 20 different chrom/scaffolds
Average (min-max) Block Sizes: 4.232 ( 3.515 - 4.975 ) Mb
```

The resulting genome-wide $F_{ST}$ estimated as the mean over block-jackknife sample is stored in the *mean.fst* element of the output list. Note that *mean.fst* may slightly differ from the default genome-wide estimate of $F_{ST}$ (stored in the *FST* element of the output list, as above with no block-jackknife sampling is carried out) as it is only computed from the SNPs eligible for block-jackknife (see Appendix A.1):

```
sim6p.allelecount.fst$FST      #genome-wide Fst over all populations
```

```
[1] 0.132319
```

```
sim6p.allelecount.fst$mean.fst #block-jacknife estimate of s.e.
```

```
[1] 0.1324889
```

The standard-error of the genome-wide $F_{ST}$ estimate is stored in the *se.fst* element of the output list and may be used to construct 95% confidence intervals (CI) intervals of the estimated values:

```
sim6p.allelecount.fst$se.fst   #s.e. of the genome-wide Fst estimate
```

```
[1] 0.0007603234
```

```
#95% c.i. of the estimated genome-wide Fst
sim6p.allelecount.fst$mean.fst+c(-1.96,1.96)*sim6p.allelecount.fst$se.fst
```

```
[1] 0.1309987 0.1339791
```

- Pool-Seq read count data:

```
sim6p.readcount30X.fst<-computeFST(sim6p.readcount30X,nsnp.per.bjack.block = 1000,
                                    verbose=FALSE)
sim6p.readcount30X.fst$FST      #genome-wide Fst over all populations
```

```
[1] 0.1324199
```

```
sim6p.readcount30X.fst$mean.fst #block-jacknife estimate of s.e.
```

```
[1] 0.1326089
```

```
sim6p.readcount30X.fst$se.fst   #s.e. of the genome-wide Fst estimate
```

```
[1] 0.0007620463
```

```
#95% c.i. of the estimated genome-wide Fst
sim6p.readcount30X.fst$mean.fst+c(-1.96,1.96)*sim6p.readcount30X.fst$se.fst
```

```
[1] 0.1311153 0.1341025
```

###### 3.1.3 Computing multi-locus $F_{ST}$ to scan the genome over sliding-windows of SNPs

The *sliding.window.size* argument allows computing multi-locus $F_{ST}$ for sliding windows over the different chromosomes (or scaffolds/contigs), e.g., to carry out genome-scans for adaptive differentiation. Each sliding window includes a number of consecutive SNPs specified by the *sliding.window.size* argument. This is illustrated below for the Pool-Seq read count example data (similar results would be obtained with allele count data):

```
sim6p.readcount30X.fst<-computeFST(sim6p.readcount30X,sliding.window.size=50)
```

```
Start sliding-window scan
20 chromosomes scanned (with more than 50 SNPs)

Average (min-max) Window Sizes 207.4 ( 90.1 - 453.2 ) kb
```

```
plot(sim6p.readcount30X.fst$sliding.windows.fst$CumulatedPosition/1e6,
     sim6p.readcount30X.fst$sliding.windows.fst$MultiLocusFst,
     xlab="Cumulated Position (in Mb)",ylab="Muli-locus Fst",
     col=as.numeric(sim6p.readcount30X.fst$sliding.windows.fst$Chr),pch=16)
abline(h=sim6p.readcount30X.fst$FST,lty=2)
```

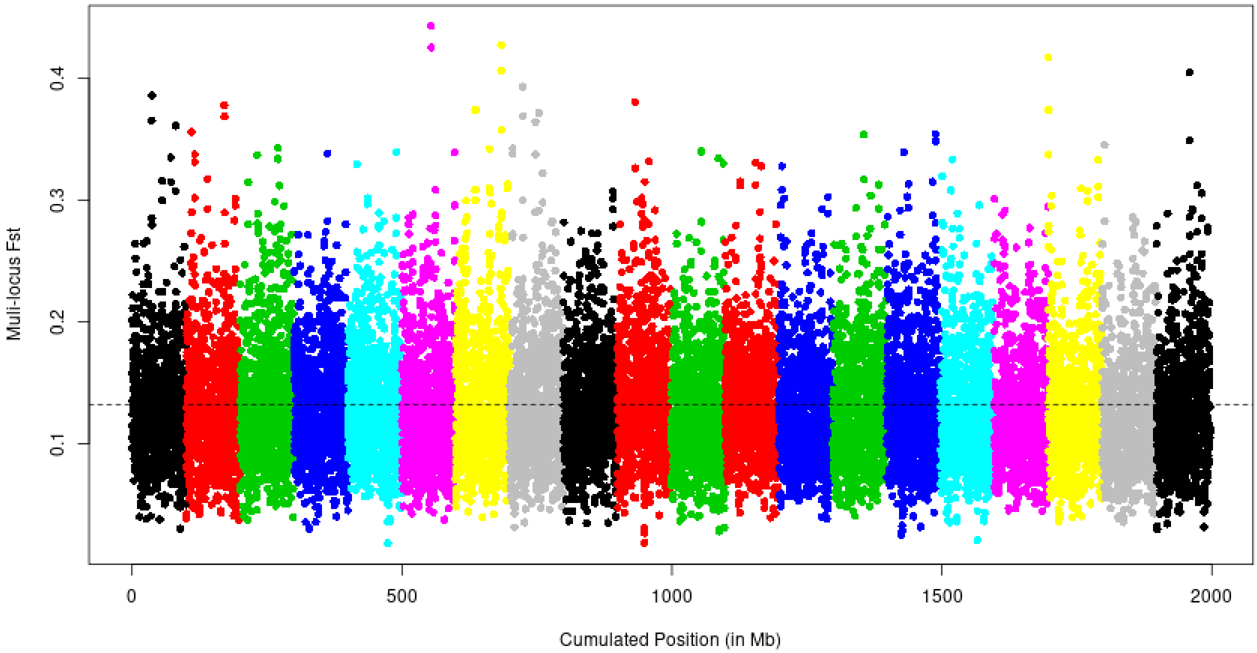

Figure 2: Manhattan plot of the multi-locus $F_{ST}$ computed over sliding-windows of 50 SNPs on the Pool-Seq example data. The dashed line indicates the estimated overall genome-wide $F_{ST}$. The 20 simulated chromosomes are represented by alternate colors.

As expected (since the data set was simulated under neutrality), no clear outlier signal of adaptive differentiation (like e.g., a tower of overly differentiated windows) shows up (Figure 2).

#### 3.2 Estimating and visualizing pairwise-population $F_{ST}$

##### 3.2.1 The *compute.pairwiseFST* and the *heatmap* functions

The *compute.pairwiseFST* function allows to estimate the genome-wide $F_{ST}$ for all the $\frac{npop(npop-1)}{2}$ pairs of populations from data stored in either a *countdata* or a *pooldata* object. As for the *computeFST* function (section 3), the *compute.pairwiseFST* function automatically uses the appropriate estimation procedure for the type of the input data (either allele count for *countdata* objects or Pool-Seq read count data for *pooldata* objects). The function returns an S4 object of class *pairwisefst* whose elements (slots) are detailed in the documentation page that can be accessed with the following command (or with the *?* operator):

```
help(pairwisefst)
```

The pairwise-population $F_{ST}$ may then be visualized using the generic *heatmap* function directly applied on the obtained *pairwisefst* object as illustrated below for Pool-Seq example results (similar results are obtained with the allele count data):

```
sim6p.pairwisefst<-compute.pairwiseFST(sim6p.readcount30X,verbose=FALSE)
```

```
Overall Analysis Time: 0 h 0 m 3 s
```

```
heatmap(sim6p.pairwisefst)
```

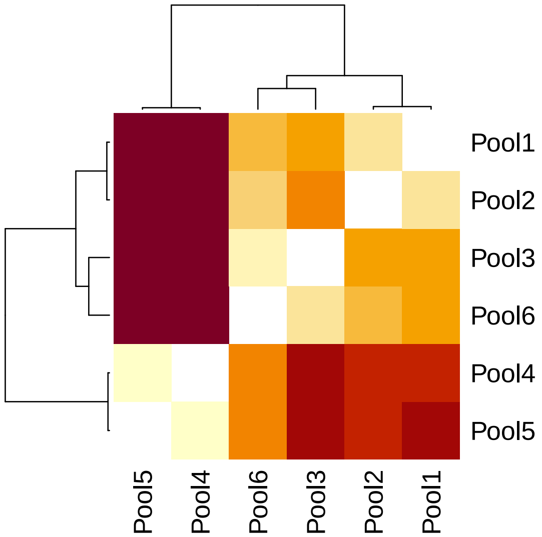

Figure 3: Heatmap representing the pairwise-population $F_{ST}$ matrix of the six populations of the 30X Pool-Seq example data set

The resulting heat map (Figure 3) is consistent with the simulated scenario (Figure 1). Note that the population *P3* is the closest to the admixed population *P6* (leading to their early clustering in the binary tree representation) as expected from the high contribution of the *P3* ancestor ($1-\alpha=75\%$) to the admixed ancestor of *P6* and the short timing of admixture ($\tau=\frac{t}{2N_e}=0.0125$).

##### 3.2.2 Block-Jackknife estimation of $\hat{F}_{ST}$ standard-error and visualisation of confidence intervals

As with the *computeFST* function, standard-error of the pairwise-population $F_{ST}$ estimates may be estimated using a block-jackknife sampling approach (see Appendix A.1) by specifying the number of consecutive SNPs forming each block with the argument *nsnp.per.bjack.block* (by default *nsnp.per.bjack.block=0*, i.e., no block-jackknife is carried out). The resulting estimated standard-errors may directly be used to derive confidence intervals (see above) that can also be plotted with the *plot_fstats* function (or directly using the *plot* command that calls *plot_fstats* for *pairwisefst* objects). This is illustrated below with the allele count example data (similar results are obtained with the Pool-Seq read count data):

```
sim6p.pairwisefst<-compute.pairwiseFST(sim6p.allelecount,
                                        nsnp.per.bjack.block = 1000,verbose=FALSE)
```

```
Overall Analysis Time: 0 h 0 m 7 s
```

```
#Estimated pairwise Fst are stored in the slot values:
#5 first estimated pairwise
head(sim6p.pairwisefst@values)
```

```
      Fst Estimate Fst bjack mean Fst bjack s.e. Q2 Estimate Q2 bjack mean Q2 bjack s.e.   Nsnp
P1;P2   0.04946710     0.04936227   0.0007453263   0.8323244     0.8323799  0.0005895541 472410
P1;P3   0.09478083     0.09499692   0.0010947422   0.8243978     0.8244107  0.0005931841 472410
P1;P4   0.17816331     0.17843286   0.0015418662   0.8095896     0.8096142  0.0006251644 472410
P1;P5   0.17716282     0.17716008   0.0015559078   0.8094495     0.8096229  0.0006214764 472410
P1;P6   0.07039455     0.07055328   0.0009233910   0.8265288     0.8265487  0.0005869646 472410
P2;P3   0.09543866     0.09560477   0.0011336466   0.8236410     0.8236723  0.0005878641 472410
```

```
plot(sim6p.pairwisefst)
```

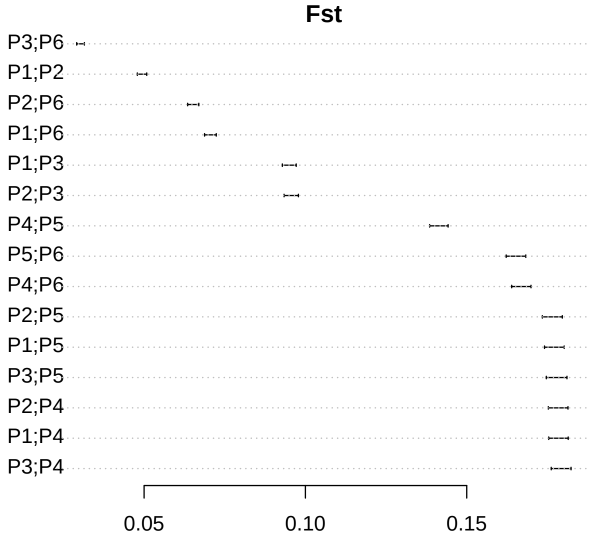

Figure 4: Estimated pairwise-population $F_{ST}$ with their 95% confidence intervals for allele count example data set

The resulting estimated pairwise-population $F_{ST}$ displayed in Figure 4 are consistent with the simulated scenario (Figure 1). The lowest level of differentiation is observed for the *P3* and *P6* population pair as expected from the high contribution of the *P3* ancestor ($1-\alpha=75\%$) to the admixed ancestor of *P6* and the short timing of admixture ($\tau=\frac{t}{2N_e}=0.0125$).

### 4 Estimating and visualizing $f$-statistics ($f_2$, $f_3$, $f_3^\star$, $f_4$ and D)

The $f_2$, $f_3$ and $f_4$ statistics were introduced in a seminal paper by Reich and co-workers (Reich *et al.* 2009) retracing the history of Indian human population and forms the core components of a general framework for demographic history inference detailed in Patterson *et al.* (2012; see also Lipson *et al.* 2013; Peter 2016; Lipson 2020). These statistics measure (expected) covariance in allele frequencies among sets of two ($F_2$), three ($F_3$) or four ($F_4$) populations and are formally defined as follows (denoting $p_i$ the SNP reference allele frequency in population $i$):

- $\mathrm{F}_2(A;B) \equiv \mathbb{E}\left[(p_A-p_B)^2\right]$
- $\mathrm{F}_3(A;B,C) \equiv \mathbb{E}\left[(p_A-p_B)(p_A-p_C)\right] \equiv \frac{1}{2}\left(\mathrm{F}_2(A;B)+\mathrm{F}_2(A;C)-\mathrm{F}_2(B;C)\right)$
- $\mathrm{F}_4(A,B;C,D) \equiv \mathbb{E}\left[(p_A-p_B)(p_C-p_D)\right] \equiv \frac{1}{2}\left(\mathrm{F}_2(A;D)+\mathrm{F}_2(B;C)-\mathrm{F}_2(A;C)-\mathrm{F}_2(B;D)\right)$

The definitions of the F parameters are not depending on the reference allele choice since $\left((1-p_A)-(1-p_B)\right)^2=(p_B-p_A)^2=(p_A-p_B)^2$. As a consequence, $F_2$ and all the other $F$ parameters may also be defined in terms of IIS within and between pairs of population as $\mathrm{F}_2(A;B)=Q_1-Q_2$ (see section 3) which allows deriving unbiased estimators for both Pool-Seq read count and standard allele count data (Gautier *et al.* 2021).

**Notice**

With $I$ populations, there are $\binom{I}{2} = \frac{1}{2}I(I-1)$ possible $F_2$ (i.e., 15 with $I = 6$ populations); $3\binom{I}{3} = \frac{1}{2}I(I-1)(I-2)$ possible $F_3$ (i.e., 60 with $I = 6$ populations); and $3\binom{I}{4} = \frac{1}{8}I(I-1)(I-2)(I-3)$ possible $F_4$ (i.e., 45 with $I = 6$ populations). However, due to their underlying linear dependency (see the above definitions), these $\frac{1}{8}I(I-1)(I^2-I+2)$ F-statistics form a vector space of dimension $\frac{1}{2}I(I-1)$ the basis of which may be specified by the set of all the $\binom{I}{2}$ possible $F_2$ statistics or, given a reference population $i$ (randomly chosen among the I ones) the set of all the $I-1$ $F_2$ statistics of the form $F_2(i;j)$ (with $j \neq i$) and the $\binom{I-1}{2}$ $F_3$ statistics of the form $F_3(i;j,k)$ (with $j \neq i$; $k \neq i$ and $j \neq k$) (Patterson *et al.* 2012; Lipson 2020). The resulting basis is informative about population history and may be used to fit admixture graph (see section 5).

Moreover, although $f_2$ statistics are difficult to interpret or to compare across pairs of populations (see the notice below), formal tests of population admixture ("3-populations" test) and tests of treeness of population quadruplets ("4-populations" test) can directly be performed using $f_3$ (see e.g., section 4.1.2) and $f_4$ (see e.g., section 4.1.3) statistics respectively (Patterson *et al.* 2012). Indeed, if $f_3(A;B,C) < 0$, we can conclude that population A originates from a population that is admixed between two source populations related to populations B and C respectively (although the signal may vanish if population A has drifted too much drifted since admixture or the admixture rates are too close to 0 or 1). Conversely, if $f_4(A,B;C,D) = 0$, the populations A, B, C and D are related by a bifurcating tree with the unrooted topology (A,B;C,D) although some may be admixed (if the paths connecting the (A,B) and (C,D) population pairs are not overlapping, see section 4.1.3 for an example). Finally, under certain circumstances, proportions of ancestry that contributed to a given admixed population can be estimated with ratios of $f_4$ statistics (see e.g., 4.3) for set of carefully related populations (Patterson *et al.* 2012).

**Notice**

The parameters $F_2$, $F_3$ and $F_4$ are not scaled with respect to the distribution of marker information content (i.e., heterozygosities). As a consequence, their resulting estimates may strongly depend on the chosen set of genetic markers (Patterson *et al.* 2012). The well-known $F_{ST}$ parameter and the two parameters $\mathrm{F}_3^\star$ and $D$ introduced by Patterson *et al.* (2012) correspond to scaled versions of $F_2$, $F_3$ and $F_4$ expected to be less sensitive to the SNP ascertainment and thus more comparable across data sets. As shown in Gautier *et al.* (2021), these can be defined in terms of IIS probabilities as:

- $$\mathrm{F_{ST}}(A;B) \equiv \frac{\mathrm{F}_2(A;B)}{1-Q_2^{A,B}} = \frac{Q_1^A + Q_1^B - 2Q_2^{A,B}}{2\left(1-Q_2^{A,B}\right)}$$
- $$\mathrm{F}_3^\star(A;B,C) \equiv \frac{\mathrm{F}_3(A;B,C)}{1-Q_1^A} = \frac{Q_1^A + Q_2^{B,C} - Q_2^{A,B} - Q_2^{A,C}}{2\left(1-Q_1^A\right)}$$
- $$\mathrm{D}(A,B;C,D) \equiv \frac{\mathrm{F}_4(A,B;C,D)}{\left(1-Q_2^{A,B}\right)\left(1-Q_2^{C,D}\right)} = \frac{Q_2^{A,C} + Q_2^{B,D} - Q_2^{A,D} - Q_2^{B,C}}{2\left(1-Q_2^{A,B}\right)\left(1-Q_2^{C,D}\right)}$$

Three-population and Four-population tests naturally extend to $F_3^\star$ and $D$ statistics respectively. An advantage of $D$ over $F_4$ is that it is constrained to the $[-1,1]$ interval and may thus be interpreted as the magnitude of deviation to treeness of the tested quadruplet (Patterson *et al.* 2012).

#### 4.1 The *compute.fstats* function and *fstats* objects

The *compute.fstats* function implements unbiased estimators of the parameters $F_2$ (and $F_{ST}$), $F_3$ (and $F_3^\star$), $F_4$ and $D$ defined above for allele count data (stored in *countdata* objects, see section 2.1) or Pool-Seq read count data (stored in *pooldata* objects, see section 2.2) as described in Gautier *et al.* (2021)[12]. The function also allows estimating within-population heterozygosities (defined as $1 - Q_1$) which is needed to scale branch lengths of admixture graphs in drift units (see section 5) or for rooting neighbor-joining trees of unadmixed populations (see section 6.1). Block-jackknife estimates of standard errors of the different estimators (needed for "Three-populations" and "Four-populations" tests, see below; and to fit admixture graph, see section 5) and their covariance (needed to fit admixture graph, see section 5) may also be performed.

As for the *computeFST* (section 3.1.1) and *compute.pairwiseFST* (section 3.2), the *compute.fstats* functions automatically detects which estimator to implement according to the class of the input object (either *countdata* or *pooldata*). The function estimates by default all the within population heterozygosities, the $F_2$ (and its scaled version $F_{ST}$[13]), $F_3$ (and its scaled version $F_3^\star$) $F_4$ statistics (and the scaled version $F_{ST}$ )[14]. Computation of $D$ statistics (i.e., scaled $F_4$) is not carried out by default (as specified with the *computeDstat* argument set to *FALSE* by default) since this may add some non negligible additional computation time for data with a large number of populations due to the extra computation of the $F_4$ scaling factor[15] although in the example below (with only 6 populations) the difference in running time is negligible.

The results are then stored in an object of class *fstats* whose elements (slots) are detailed in the documentation page accessible with the following command (or the *?* operator):

```
help(fstats)
```

The underlying f-statistics may then be easily accessible or visualized with the *plot_fstats* function (or directly using the *plot* command that calls *plot_fstats* for *fstats* objects) as illustrated below for the allele count (*sim6p.allelecount*) and Pool-Seq read count (*sim6p.readcount30X*) example data[16].

```
##Estimation of f-statistics on count data
sim6p.allelecount.fstats<-compute.fstats(sim6p.allelecount,nsnp.per.bjack.block = 1000,
                                         computeDstat = TRUE)
```

```
Estimating Q1
Estimating Q2
Estimating within-population heterozygosities
Estimating F2
Estimating F3
Estimating F4
 Computing Dstat
Starting Block-Jackknife sampling
462 Jackknife blocks identified with 462000 SNPs (out of 472410 ).
 SNPs map to 20 different chrom/scaffolds
Average (min-max) Block Sizes: 4.232 ( 3.515 - 4.975 ) Mb
   computing Q1 averages per blocks
   computing Q2 averages per blocks
   computing F2 averages per blocks
Starting computation of estimators s.e.
   within-pop heterozygosity s.e. estimation done
   F2 s.e. estimation done
   F3 and F3* s.e. estimation done
   estimating F4 and Dstat s.e. (may be long since require denominator averages per blocks)
   F4 and D s.e. estimation done

Overall Analysis Time: 0 h 0 m 3 s
```

[12]Although not defined in the same way, estimator for allele count data are strictly equivalent to those by Patterson *et al.* (2012)

[13]The estimator is then actually exactly the same as the one implemented in the *compute.pairwiseFST* or *computeFST* functions when the argument *method="Identity"*

[14]The *compute.fstats* function is optimized in such a way that the computational cost for the estimation of pairwise $F_{ST}$, $F_3$, $F_3^\star$ and $F_4$ from the $F_2$-statistics and $Q_2$ estimates is negligible

[15]For instance on a tested allele count real data set consisting of 640,000 SNPs genotyped on 24 populations, *compute.fstats* ran in 2 m 56 s (6 m 50 if *nsnp.per.bjack.block = 5000*) with *computeDstat=TRUE* and only 7 s (16 s) if *computeDstat=FALSE*

[16]Note that estimates of the different statistics are highly similar between the allele count and the Pool-Seq read count data.

```
sim6p.allelecount.fstats
```

```
 * * * fstats Object * * *
Example of useful visualization functions are plot.fstats
```

```
##Estimation of f-statistics on Pool-Seq data (without computation of Dstat)
sim6p.readcount30X.fstats<-compute.fstats(sim6p.readcount30X,nsnp.per.bjack.block = 1000,
                                          verbose=FALSE)
##Estimation of f-statistics on Pool-Seq data (with computation of Dstat)
sim6p.readcount30X.fstats<-compute.fstats(sim6p.readcount30X,nsnp.per.bjack.block = 1000,
                                         computeDstat = TRUE,verbose=FALSE)
```

###### 4.1.1 $f_2$ and $F_{ST}$ estimates (***f2.values*** and ***fst.values*** slots of the ***fstat*** object)

```
# count data (3 first f2)
head(sim6p.allelecount.fstats@f2.values,3)
```

```
         Estimate  bjack mean   bjack s.e.
P1,P2 0.008294424 0.008274108 0.0001298376
P1,P3 0.016643720 0.016680443 0.0002108642
P1,P4 0.033924154 0.033971079 0.0003456714
```

```
# 30X Pool-Seq data (3 first f2)
head(sim6p.readcount30X.fstats@f2.values,3)
```

```
              Estimate  bjack mean   bjack s.e.
Pool1,Pool2 0.00829429 0.008275487 0.0001351787
Pool1,Pool3 0.01665818 0.016696323 0.0002159221
Pool1,Pool4 0.03398383 0.034045968 0.0003525136
```

```
# count data (3 first Fst)
head(sim6p.allelecount.fstats@fst.values,3)
```

```
        Estimate bjack mean   bjack s.e.
P1,P2 0.04946710 0.04936227 0.0007453263
P1,P3 0.09478083 0.09499692 0.0010947422
P1,P4 0.17816331 0.17843286 0.0015418662
```

```
# 30X Pool-Seq data (3 first Fst)
head(sim6p.readcount30X.fstats@fst.values,3)
```

```
              Estimate bjack mean  bjack s.e.
Pool1,Pool2 0.04947320 0.04937253 0.000774369
Pool1,Pool3 0.09487048 0.09508626 0.001119595
Pool1,Pool4 0.17839940 0.17872795 0.001572758
```

Notice that the pairwise $F_{ST}$ estimates are the same as those obtained with the *compute.pairwiseFST* function (see section 3.2) run with *method=Identity* (which are here actually equal with estimates obtained by the default *Anova* method because there is no variation in the total allele counts for all SNPs).

**Notice**

By construction $F_2(A;B) = F_2(B,A)$ (and $F_{ST}(A;B) = F_{ST}(B,A)$). If $i_P$ is the index of population $P$ in the *popnames* or *poolnames* slots of the *countdata* or *pooldata* objects (i.e., the column order in the corresponding allele or read count data matrices) used to obtain the *fstats* object, the $F_2(A,B)$ (resp. $F_{ST}(B,A)$) configurations reported in the slot $f2.values$ (resp. $fst.values$) satisfy $i_A < i_B$.

###### 4.1.2 $f_3$ and $f_3^\star$ estimates (*f3.values* and *f3star.values* slots of the *fstat* object) and 3-Population tests:

```
# count data (3 first f3)
head(sim6p.allelecount.fstats@f3.values,3)
```

```
            Estimate  bjack mean   bjack s.e.  Z-score
P1;P2,P3 0.004053338 0.004048392 0.0001176972 34.39668
P1;P2,P4 0.004089298 0.004087474 0.0001326605 30.81155
P1;P2,P5 0.004155216 0.004149524 0.0001341405 30.93416
```

```
# 30X Pool-Seq data (3 first f3)
head(sim6p.readcount30X.fstats@f3.values,3)
```

```
                     Estimate  bjack mean   bjack s.e.  Z-score
Pool1;Pool2,Pool3 0.004057570 0.004053794 0.0001247034 32.50750
Pool1;Pool2,Pool4 0.004114142 0.004114670 0.0001365234 30.13893
Pool1;Pool2,Pool5 0.004157049 0.004154896 0.0001411765 29.43051
```

```
# count data (3 first f3*)
head(sim6p.allelecount.fstats@f3star.values,3)
```

```
           Estimate bjack mean   bjack s.e.  Z-score
P1;P2,P3 0.02552286 0.02549608 0.0007467871 34.14102
P1;P2,P4 0.02574929 0.02574221 0.0008480235 30.35554
P1;P2,P5 0.02616436 0.02613299 0.0008581050 30.45430
```

```
# 30X Pool-Seq data (3 first f3*)
head(sim6p.readcount30X.fstats@f3star.values,3)
```

```
                    Estimate bjack mean   bjack s.e.  Z-score
Pool1;Pool2,Pool3 0.02555404 0.02553297 0.0007888059 32.36915
Pool1;Pool2,Pool4 0.02591032 0.02591640 0.0008722134 29.71337
Pool1;Pool2,Pool5 0.02618054 0.02616977 0.0009009973 29.04533
```

**Notice**

By construction $F_3(A;B,C) = F_3(A;C,B)$ (and $F_3^\star(A;B,C) = F_3^\star(A;C,B)$). If $i_P$ is the index of population $P$ in the *popnames* or *poolnames* slots of the *countdata* or *pooldata* objects (i.e., the column order in the corresponding allele or read count data matrices) used to obtain the *fstats* object, the $F_3(A;B,C)$ (and $F_3^\star(A;B,C)$) configurations reported in the slot $f3.values$ satisfy $i_B < i_C$.

As shown in the above example, activation of block-jackknife estimation of standard errors (i.e., argument *nsnp.per.bjack.block*>0) results in the computation of Z-scores (i.e., ratio of the block-jackknife estimated mean and standard-error) which quantifies the deviation of the estimated $f_3$-statistics from 0 (in units of s.e.). This gives a simple decision criterion for three-population tests of admixture (i.e., negative $f_3$ or negative $f_3^\star$). For instance a Z-score $< -1.65$ provides evidence for admixture (i.e., significantly negative $f_3$) at the 95% significance threshold:

```
# count data (F3-based 3-pop test)
tst.sel<-sim6p.allelecount.fstats@f3.values$`Z-score`< -1.65
sim6p.allelecount.fstats@f3.values[tst.sel,]
```

```
              Estimate    bjack mean   bjack s.e.   Z-score
P6;P2,P3 -0.0002671143 -0.0002623528 8.638581e-05 -3.036989
```

```
# 30X Pool-Seq data (F3-based 3-pop test)
tst.sel<-sim6p.readcount30X.fstats@f3.values$`Z-score`< -1.65
sim6p.readcount30X.fstats@f3.values[tst.sel,]
```

```
                       Estimate    bjack mean   bjack s.e.   Z-score
```

```
Pool6;Pool2,Pool3 -0.0002417509 -0.0002304611 8.851744e-05 -2.603567
```

```r
# count data (F3*-based 3-pop test)
tst.sel<-sim6p.allelecount.fstats@f3star.values$`Z-score`< -1.65
sim6p.allelecount.fstats@f3star.values[tst.sel,]
```

```
              Estimate   bjack mean   bjack s.e.   Z-score
P6;P2,P3 -0.001631657 -0.001603205 0.0005274779 -3.039379
```

```r
# 30X Pool-Seq data (F3*-based 3-pop test)
tst.sel<-sim6p.readcount30X.fstats@f3star.values$`Z-score`< -1.65
sim6p.readcount30X.fstats@f3star.values[tst.sel,]
```

```
                      Estimate   bjack mean   bjack s.e.   Z-score
Pool6;Pool2,Pool3 -0.001475866 -0.001407435 0.0005402774 -2.605024
```

In agreement with the simulated scenario (Figure 1), both the allele count and Pool-Seq read count data support an admixed origin for population *P6* with ancestral sources related to *P2* and *P3*.

###### 4.1.3 $f_4$ and $D$ estimates (*f4.values* and *Dstat.values* slots of the *fstat* object) and 4-Population tests:

```r
# count data (3 first f4)
head(sim6p.allelecount.fstats@f4.values,3)
```

```
                Estimate   bjack mean   bjack s.e.   Z-score
P1,P2;P3,P4 3.595944e-05 3.908225e-05 1.154193e-04 0.3386111
P1,P2;P3,P5 1.018776e-04 1.011325e-04 1.197917e-04 0.8442359
P1,P2;P3,P6 5.594310e-04 5.622762e-04 6.138658e-05 9.1595941
```

```r
# 30X Pool-Seq data (3 first f4)
head(sim6p.readcount30X.fstats@f4.values,3)
```

```
                            Estimate   bjack mean   bjack s.e.   Z-score
Pool1,Pool2;Pool3,Pool4 5.657166e-05 6.087576e-05 1.194004e-04 0.5098456
Pool1,Pool2;Pool3,Pool5 9.947887e-05 1.011014e-04 1.253135e-04 0.8067878
Pool1,Pool2;Pool3,Pool6 5.653400e-04 5.621051e-04 6.409453e-05 8.7699380
```

```r
# count data (3 first D)
head(sim6p.allelecount.fstats@Dstat.values,3)
```

```
                Estimate   bjack mean  bjack s.e.   Z-score
P1,P2;P3,P4 0.0006651386 0.0007230774 0.002135688 0.3385688
P1,P2;P3,P5 0.0018826436 0.0018705234 0.002216171 0.8440338
P1,P2;P3,P6 0.0107508025 0.0108152267 0.001182842 9.1434253
```

```r
# 30X Pool-Seq data (3 first D)
head(sim6p.readcount30X.fstats@Dstat.values,3)
```

```
                           Estimate  bjack mean  bjack s.e.   Z-score
Pool1,Pool2;Pool3,Pool4 0.001045743 0.001125524 0.002208046 0.5097373
Pool1,Pool2;Pool3,Pool5 0.001837431 0.001868847 0.002316956 0.8065961
Pool1,Pool2;Pool3,Pool6 0.010853673 0.010799873 0.001235136 8.7438717
```

**Notice**

When comparing two pairs of populations $(A, B)$ and $(C, D)$, the $f_4$ statistics for the 8 quadruplets (A,B;C,D); (B,A;C,D); (A,B;D,C); (B,A;D,C); (C,D;A,B); (C,D;B,A); (D,C;A,B) and (D,C;B,A) have the same absolute value by definition of the $F_4$ parameter:

$$\begin{aligned} F_4(A,B;C,D) &= F_4(B,A;D,C) = F_4(C,D;A,B) = F_4(D,C;A,B) \\ -F_4(A,B;C,D) &= F_4(B,A;C,D) = F_4(C,D;B,A) = F_4(D,C;B,A) = F_4(A,B;D,C) \end{aligned}$$

and similarly

$$\begin{aligned} D(A,B;C,D) &= D(B,A;D,C) = D(C,D;A,B) = D(D,C;A,B) \\ -D(A,B;C,D) &= D(B,A;C,D) = D(C,D;B,A) = D(D,C;B,A) = D(A,B;D,C) \end{aligned}$$

If $i_P$ is the index of population $P$ in the *popnames* or *poolnames* slots of the *countdata* or *pooldata* objects (i.e., the column order in the corresponding allele or read count data matrices) used to obtain the *fstats* object, the $F_4(A,B;C,D)$ (and $D(A,B;C,D)$) configurations reported in the slot $f4.values$ (and $Dstat.values$) satisfy $i_A < i_B$; $i_A < i_C$ and $i_C < i_D$.

As for $f_3$ and $f_3^\star$ (section 4.1.2), activating block-jackknife estimation of standard errors (i.e., the argument *nsnp.per.bjack.block*>0) results in the computation of Z-scores (i.e., ratio of the block-jackknife estimated mean and standard-error) which quantifies the deviation of the estimated $f_4$-statistics from 0 (in units of s.e.). This gives a simple decision criterion for four-population tests of treeness (i.e., non null $F_4$ or $D$). For instance a Z-score lower than 1.96 in absolute value provides no evidence against the null-hypothesis of treeness for the tested population configuration at the 95% significance threshold:

```
# count data
tst.sel<-abs(sim6p.allelecount.fstats@f4.values$`Z-score`)<1.96
sim6p.allelecount.fstats@f4.values[tst.sel,]
```

```
                 Estimate    bjack mean   bjack s.e.    Z-score
P1,P2;P3,P4  3.595944e-05  3.908225e-05 1.154193e-04  0.3386111
P1,P2;P3,P5  1.018776e-04  1.011325e-04 1.197917e-04  0.8442359
P1,P2;P4,P5  6.591816e-05  6.205022e-05 9.313675e-05  0.6662270
P1,P3;P4,P5  1.309202e-05  1.464589e-05 1.268061e-04  0.1154983
P1,P6;P4,P5  4.031413e-05  4.458874e-05 1.058348e-04  0.4213050
P2,P3;P4,P5 -5.282615e-05 -4.740433e-05 1.302898e-04 -0.3638376
P2,P6;P4,P5 -2.560403e-05 -1.746147e-05 1.085089e-04 -0.1609221
P3,P6;P4,P5  2.722212e-05  2.994286e-05 7.650662e-05  0.3913760
```

```
# 30X Pool-Seq data
tst.sel<-abs(sim6p.readcount30X.fstats@f4.values$`Z-score`)<1.96
as.data.frame(sim6p.readcount30X.fstats@f4.values)[tst.sel,]
```

```
                             Estimate    bjack mean   bjack s.e.     Z-score
Pool1,Pool2;Pool3,Pool4  5.657166e-05  6.087576e-05 1.194004e-04  0.50984558
Pool1,Pool2;Pool3,Pool5  9.947887e-05  1.011014e-04 1.253135e-04  0.80678780
Pool1,Pool2;Pool4,Pool5  4.290721e-05  4.022566e-05 9.502690e-05  0.42330817
Pool1,Pool3;Pool4,Pool5  3.769747e-07 -3.302134e-06 1.277902e-04 -0.02584028
Pool1,Pool6;Pool4,Pool5  4.073843e-05  4.734750e-05 1.053345e-04  0.44949661
Pool2,Pool3;Pool4,Pool5 -4.253023e-05 -4.352780e-05 1.336022e-04 -0.32580156
Pool2,Pool6;Pool4,Pool5 -2.168779e-06  7.121841e-06 1.102031e-04  0.06462467
Pool3,Pool6;Pool4,Pool5  4.036145e-05  5.064964e-05 8.000836e-05  0.63305432
```

In other words, both the allele count and Pool-Seq read count data only provide no evidence against the null hypothesis of treeness at the 95% threshold for quadruplets involving *i)* non-admixed populations (*P1*, *P2*, *P3*, *P4* and *P5*) for configurations consistent with the simulated scenario; and *ii)* the admixed population *P6* for configurations of the form *(P6,X;P4,P5)* where *P4* and *P5* are the two outgroup populations and X=*P1*,

*P2* or *P3*. This is actually expected since for these latter quadruplets, the path connecting *P4* and *P5* is not overlapping with either of the paths connecting *P6* to *P1*, *P2* or *P3* in the simulated graph (Figure 1).

###### 4.1.4 Population heterozygostity estimates (*heterozygosities* slot of the *fstat* object)

```
# count data  (3 first populations)
head(sim6p.allelecount.fstats@heterozygosities,3)
```

```
    Estimate bjack mean   bjack s.e.
P1 0.1588121  0.1587849 0.0005972556
P2 0.1599502  0.1599070 0.0006057583
P3 0.1591048  0.1590328 0.0005900926
```

```
# 30X Pool-Seq data (3 first populations)
head(sim6p.readcount30X.fstats@heterozygosities,3)
```

```
       Estimate bjack mean   bjack s.e.
Pool1 0.1587839  0.1587670 0.0006012812
Pool2 0.1599319  0.1599083 0.0006096762
Pool3 0.1590771  0.1590230 0.0006024449
```

##### 4.2 The *plot_fstats* function for visualization of $f_2$ (and pairwise $F_{ST}$), $f_3$ (and $f_3^\star$) and $f_4$ (and $D$) estimates and their confidence intervals

The *plot_fstats* function (that may be called directly using *plot* on *fstats* objects) allows plotting all or only some (using the *pop.sel*, *pop.f3.target* or *value.range* arguments) of the estimated $f_2$, $f_{ST}$, $f_3$, $f_3^\star$, $f_4$ or $D$ statistics from a *fstats* object. In addition, for $f_3$, $f_3^\star$, $f_4$ and $D$ statistics, the *highlight.signif* argument allows highlighting in red significant (as defined with the *ci.perc* argument) three-population or four-population tests. Some example plots are shown in Figures 5, 6 and 7 which were generated with the following codes:

###### 4.2.1 Example of $f_2$ statistics plot (Figure 5)

```
layout(matrix(1:2,2,1,byrow=T))
plot(sim6p.allelecount.fstats,main="F2 (Allele Count)")
plot(sim6p.readcount30X.fstats,main="F2 (30X Pool-Seq)")
```

Similar plots may be obtained for the scaled $f_2$ (i.e., pairwise $f_{ST}$) by specifying *stat.name="Fst"* (see also the *compute.pairwiseFST* functions described in section 3.2).

###### 4.2.2 Example of $f_3$ statistics plot (Figure 6)

```
layout(matrix(1:4,2,2,byrow=T))
plot(sim6p.allelecount.fstats,stat.name="F3",main="F3 (Allele Count)")
plot(sim6p.readcount30X.fstats,stat.name="F3",main="F3 (30X Pool-Seq)")
plot(sim6p.readcount30X.fstats,stat.name="F3",pop.f3.target=c("Pool6","Pool1"),
     main="30X Pool-Seq (only F3 with P6 or P1 as target pops)")
plot(sim6p.readcount30X.fstats,stat.name="F3",value.range=c(NA,5e-3),
     main="30X Pool-Seq (only F3 < 5e-3)")
```

Similar plots may be obtained for the scaled $f_3$ (i.e., $f_3^\star$) by specifying *stat.name="F3star"*.

###### 4.2.3 Example of $f_4$ and $D$ statistics plot (Figure 7)

```
layout(matrix(1:6,3,2,byrow=T))
plot(sim6p.allelecount.fstats,stat.name="Dstat",main="D (Allele Count)")
plot(sim6p.readcount30X.fstats,stat.name="Dstat",main="D (30X Pool-Seq)")
plot(sim6p.allelecount.fstats,stat.name="F4",main="F4 (Allele Count)")
```

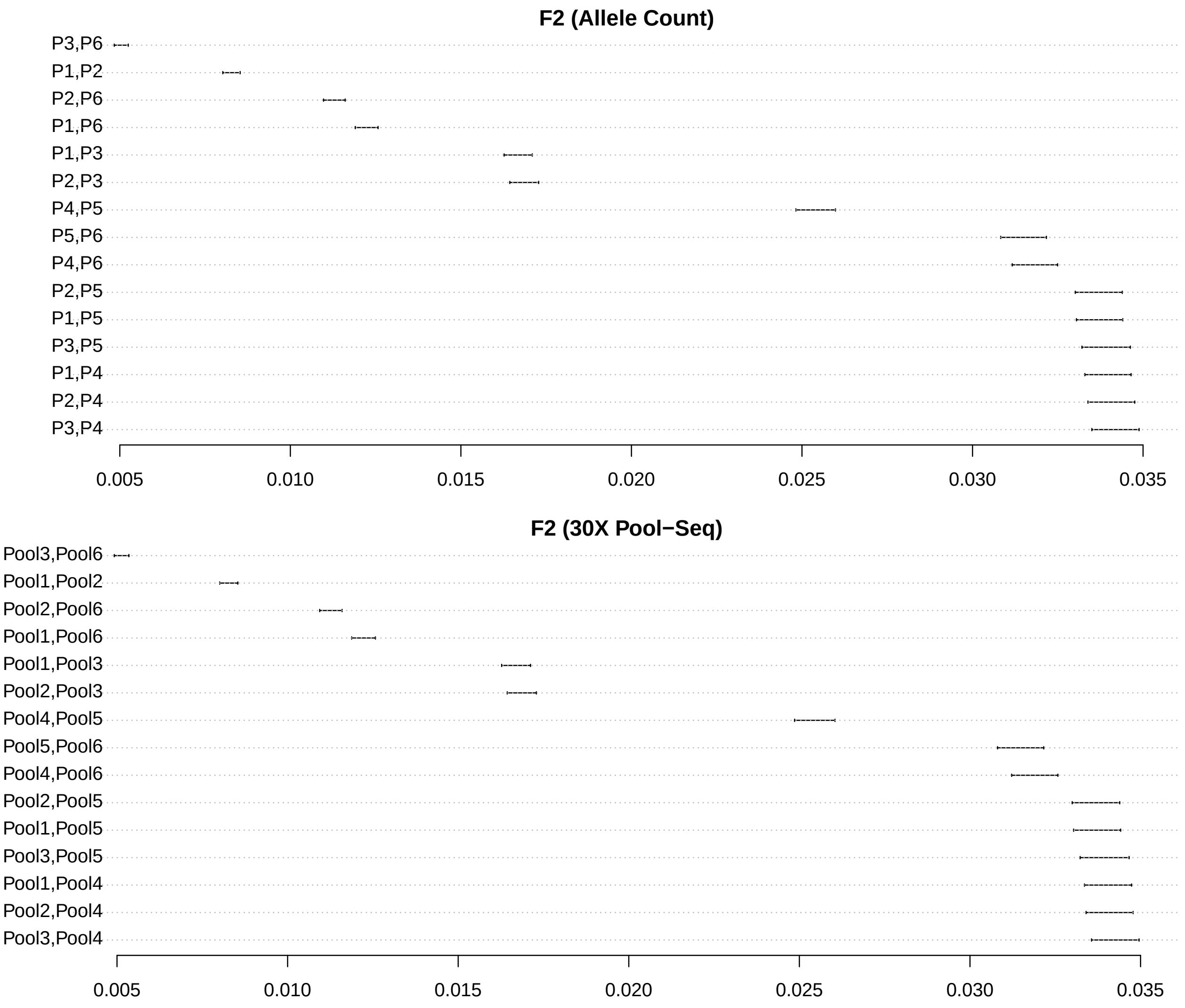

Figure 5: Estimated $f_2$ statistics with their 95% confidence intervals for the allele count and 30X Pool-Seq data sets

```
plot(sim6p.readcount30X.fstats,stat.name="F4",main="F4 (30X Pool-Seq)")
plot(sim6p.readcount30X.fstats,stat.name="F4",pop.sel=c("Pool1","Pool2"),
     main="30X Pool-Seq (only F4 with both P1 and P2)")
plot(sim6p.readcount30X.fstats,stat.name="F4",value.range=c(-2e-3,2e-3),
     main="30X Pool-Seq (only -2e-3 < F4 < 2e-3)")
```

#### 4.3 Estimating admixture proportions with $f_4$-ratios

Given an admixture graph (assumed to be correct), ratios of $f_4$-statistics (Patterson *et al.* 2012) may provide estimates of the relative contributions of the ancestral sources of a (two-way) admixed populations (*P6* in our example) if outgroups (*P1* and *P4* or *P5* in our example) for the two source population proxies (e.g., *P2* and *P3* in our example) have been sampled. For instance, the proportion $\alpha$ of $P2$-related ancestry in population $P6$ (Figure 1) is equal to:

$$\alpha = \frac{F_4\left(P1, P4; P3, P6\right)}{F_4\left(P1, P4; P2, P3\right)} = \frac{F_4\left(P1, P5; P3, P6\right)}{F_4\left(P1, P5; P2, P3\right)}$$

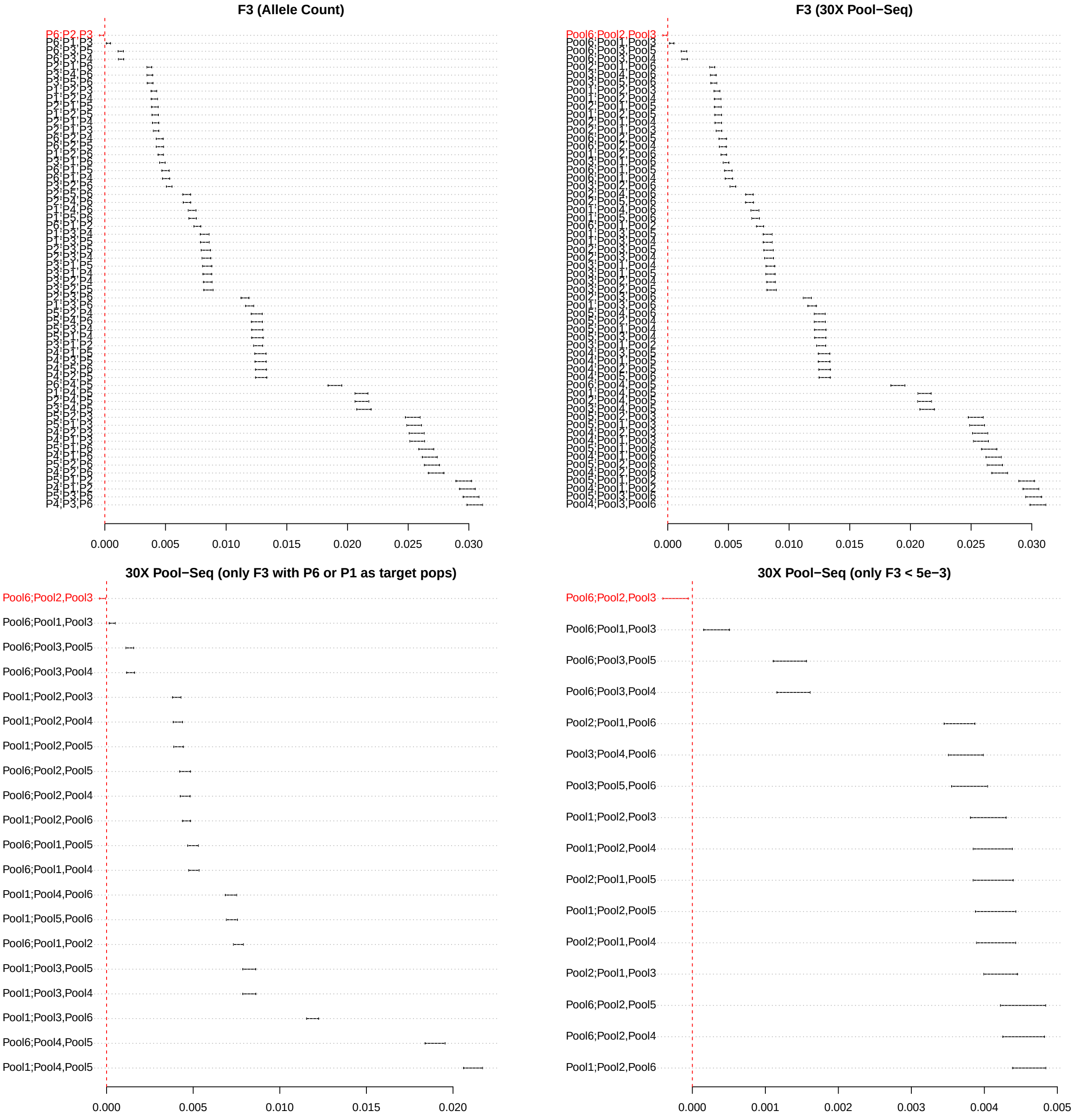

Figure 6: Estimated $f_3$ statistics with their 95% confidence intervals for the allele count and 30X Pool-Seq data sets

The *compute.f4ratio* function implements $f_4$-ratio based estimators of admixture proportion from an *fstats* object. It requires to specify both the numerator (*num.quadruplet* argument) and the denominator (*den.quadruplet*) $F_4$ quadruplets[17]. The following example illustrates how to use the function to the admixture proportion $\alpha$ ($\alpha_{\text{simulated}} = 0.25$) from the P2-related source that contributed to the P6 ancestral population (Figure 1):

[17]In practice, the required $f_4$ may not be directly available from the *f4.values* slot of the *fstats* since, as mentioned in section 4.1.3, only one out of the 8 equivalent quadruplet configurations is reported (the chosen one only depending on the ordering of the populations in the *countdata* or *pooldata* objects used for estimation). However, the function automatically computes the $f_4$ for the numerator and denominator configurations specified by the user from the estimator available in the input *fstats* object

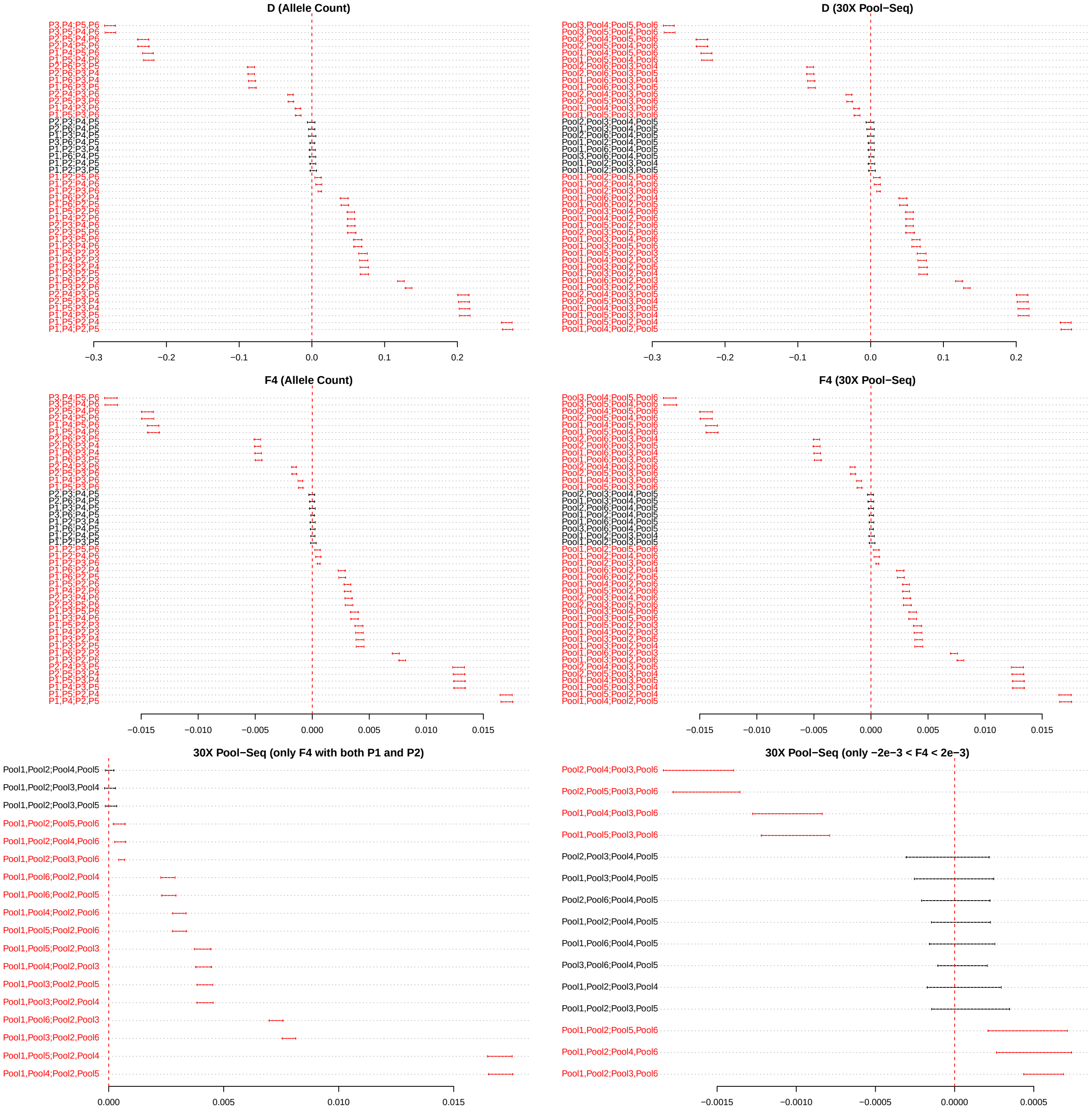

Figure 7: Estimated $f_4$ and $D$ statistics with their 95% confidence intervals for the allele count and 30X Pool-Seq data sets

```
# count data (two possible estimates)
compute.f4ratio(sim6p.allelecount.fstats,num.quadruplet = "P1,P4;P3,P6",
                                         den.quadruplet="P1,P4;P2,P3")
```

```
[1] -0.2497209
```

```
compute.f4ratio(sim6p.allelecount.fstats,num.quadruplet = "P1,P5;P3,P6",
                                         den.quadruplet="P1,P5;P2,P3")
```

```
[1] -0.2462923
```

```r
# 30X Pool-Seq data (two possible estimates)
compute.f4ratio(sim6p.readcount30X.fstats,num.quadruplet = "Pool1,Pool4;Pool3,Pool6",
                                          den.quadruplet="Pool1,Pool4;Pool2,Pool3")
```

```
[1] -0.25285
```

```r
compute.f4ratio(sim6p.readcount30X.fstats,num.quadruplet = "Pool1,Pool5;Pool3,Pool6",
                                          den.quadruplet="Pool1,Pool5;Pool2,Pool3")
```

```
[1] -0.24559
```

Standard errors of the estimated admixture proportions are automatically computed if the input *fstats* object was obtained by running the *compute.fstats* function with *return.F4.blockjackknife.samples = TRUE* (and *nsnp.per.bjack.block>0*) to activate block-jackknife estimation of standrd errors) as illustrated below for the Pool-Seq read count data:

```r
sim6p.readcount30X.fstats<-compute.fstats(sim6p.readcount30X,nsnp.per.bjack.block = 1000,
                                    verbose=FALSE,return.F4.blockjackknife.samples = TRUE)
alpha.est=compute.f4ratio(sim6p.readcount30X.fstats,num.quadruplet = "Pool1,Pool4;Pool3,Pool6",
                                                    den.quadruplet="Pool1,Pool4;Pool2,Pool3")
#95% c.i. of alpha derived from the se
alpha.est[2]+c(-1.96,1.96)*alpha.est[3]
```

```
[1] -0.3013463 -0.2100506
```

```r
#prediction error in units of s.e.
as.numeric(abs(0.25-alpha.est[2])/alpha.est[3])
```

```
[1] 21.71337
```

Note that the simulated value ($\alpha = 0.25$) is within the confidence interval and actually less than 0.25 standard error higher than the $F_4$-ratio estimate.

### 5 Using f-statistics to estimate parameters of admixture graphs

The f-statistics can be used to estimate the parameters (branch lengths and/or admixture proportions) of trees or admixture graphs (i.e., trees including admixture edges) that summarize the demographic history of the surveyed populations. The approach implemented in *poolfstat* to fit admixture graphs from f-statistics is directly inspired (and actually highly similar) to the one used in the *qpGraph* software originally described by Patterson *et al.* (2012; see also Lipson 2020). The core functions used for admixture graph fitting consist of:

- the *generate.graph.params* function to define graph parameters specifying the candidate graph to fit and the underlying f-statistics provided as an *fstats* object (section 4)
- the *fit.graph* function to estimate graph parameters using an optimization algorithm
- the *compare.fitted.fstats* function to assess model fit by comparing estimated and fitted f-statistics (Patterson *et al.* 2012; Lipson 2020)
- the *add.leave* function to evaluate all the possible admixture graphs (or trees) resulting from the addition of a new leave to an existing graph (connected with either non-admixed or admixed edges).

#### 5.1 Generating a *graph.params* object with the *generate.graph.params* function

##### 5.1.1 Specifying the structure of the admixture graph in a *graph.params* object

Admixture graph specification including the structure of the graph (i.e., topology consisting of edges and, if any admixture proportions) are defined in an object of class *graph.params* detailed in the documentation page accessible with the following command (or the *?* operator):

```r
help(graph.params)
```

The *graph.params* objects may be constructed with the *generate.graph.params* function from a user-defined (character) matrix specifying the structure of the admixture graph (or tree if no admixture edges are included) and consisting of three columns defining for each edge (whether admixed or not) i) the child node; ii) the parent node; iii) the admixture proportion (blank for non-admixed edges). As a result, in the input matrix, each admixture event is specified by two rows for the two admixture edges corresponding to the same admixed child node and two different source nodes as the parent node (i.e., source populations). Their third column elements contain the two underlying admixture proportions coded as *a* and *(1-a)* (the parentheses are mandatory and an error message is printed if absent) where *a* is the name of the admixture proportion (names of admixture proportions should not include space). The example below shows the construction of a *graph.params* object specifying the admixture graph for the scenario used to simulate the example data (Figure 1):

```
sim.graph<-rbind(c("P1","P7",""),c("P2","s1",""),c("P3","s2",""),c("P6","S",""),
                c("S","s1","a"),c("S","s2","(1-a)"),c("s2","P8",""),c("s1","P7",""),
                c("P4","P9",""),c("P5","P9",""),c("P7","P8",""),
                c("P8","R",""),c("P9","R",""))
sim.graph.params<-generate.graph.params(sim.graph)
sim.graph.params
```

```
 * * * graph.params Object * * *
Example of useful functions are:
  plot() to visualize the graph (interface for grViz() from the DiagrammeR package)
  fit.graph() to estimate graph parameter values
 * * * * * * * * * * * * * * * *
```

The root is automatically identified by the *generate.graph.params* function (as the node only present in the parent node column) and several checks are made within the function. It is however recommended to check the graph by plotting using the *plot* function. The following code shows how to plot the graph stored in the example *sim.graph.params* object that was generated above (Figure 8):

```
plot(sim.graph.params)
```

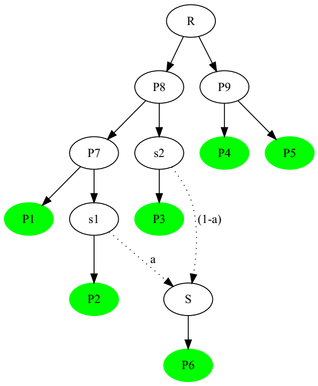

Figure 8: Plot of the admixture graph specifying the simulated scenario

**Notice**

The *plot* function applied to *graph.params* objects internally calls the *grViz* function from the *DiagrammeR* package (Iannone 2020) which actually generates an object of class *htmlwidget* that will print itself into HTML in a browser. Also, if run within the *Rstudio* IDE, the graph will be plotted in the View pane[a]. Hence, graph plots may not be easily exported from R into a pdf (or other) device although the following trick may be useful to that end:

```
require(webshot,htmlwidgets,imager)
tmp<-plot(sim.graph.params)            #plot the graph
saveWidget(tmp,"tmp.html") ; webshot("tmp.html","tmp.png")
load.image("tmp.png") %>% autocrop()  %>% plot(axes=F)
```

To insert a graph plot into an external document or edit it (e.g., to change node or edge colors, node names, etc.), one may also directly rely on its *dot* coded definition[b] stored in the *dot.graph* slot of the *graph.params* (or *fitted.graph*, see section 5.2) object outside R. A graph *dot* file may also be generated by specifying an output file name prefix with the *outfileprefix* argument of the *generate.graph.params* function[c].

[a]Regularly clearing the View items using the broom icon is recommended

[b]From the open source graph visualization software *graphviz* (https://graphviz.org/). For instance, *dot* files can be converted into *png* file using the command *dot -Tpng inputgraph.dot* in a Linux terminal. Several online user-friendly implementations also allow very convenient manipulation of *dot* files from a web-browser, see e.g., https://dreampuf.github.io/GraphvizOnline

[c]Alternatively, one may also use the command *writeLines(x@dot.graph,con=outfile)* where *x* is the *graph.params* object and *outfile* is the desired name of the dot output file (e.g., *"out.dot"*)

The *graph.params* object produced by the *generate.graph.params* function includes a symbolic representation of the graph incidence matrix (slot *graph.matrix*) which consists of a $n_l$ leaves by $n_e$ edges matrix containing the edges weight for the paths from each leave to the root[18]. Examples for the *sim.graph.params* object that specifies the simulation scenario are given below:

```
#names of the edges (automatically given)
sim.graph.params@edges.names
```

```
 [1] "P7<->P1" "s1<->P2" "s2<->P3" "S<->P6"  "P8<->s2" "P7<->s1" "P9<->P4" "P9<->P5" "P8<->P7" "R<->P8"  "R<->P9"
```

```
#names of the admixture proportions (automatically given)
sim.graph.params@adm.params.names
```

```
[1] "a"
```

```
#graph incidence matrix
sim.graph.params@graph.matrix
```

```
   P7<->P1 s1<->P2 s2<->P3 S<->P6 P8<->s2 P7<->s1 P9<->P4 P9<->P5 P8<->P7 R<->P8 R<->P9
P1 "1"     "0"     "0"     "0"    "0"     "0"     "0"     "0"     "1"     "1"    "0"
P2 "0"     "1"     "0"     "0"    "0"     "1"     "0"     "0"     "1"     "1"    "0"
P3 "0"     "0"     "1"     "0"    "1"     "0"     "0"     "0"     "0"     "1"    "0"
P6 "0"     "0"     "0"     "1"    "1-a"   "a"     "0"     "0"     "a"     "1"    "0"
P4 "0"     "0"     "0"     "0"    "0"     "0"     "1"     "0"     "0"     "0"    "1"
P5 "0"     "0"     "0"     "0"    "0"     "0"     "0"     "1"     "0"     "0"    "1"
```

[18]In the symbolic representation, the names for the graph edges and admixture proportions correspond to those stored in the *edges.names* and *adm.params.names* slots of the *graph.params* object respectively

**Notice**

As mentioned by PICKRELL and PRITCHARD (2012), PATTERSON *et al.* (2012) and LIPSON (2020), the three branch lengths surrounding an admixture event (e.g., edges s1 $\leftrightarrow$ S, s2 $\leftrightarrow$ S and S $\leftrightarrow$ P6 connecting *s1* to *S*; *s2* to *S*; and *S* to *P6* respectively for the admixture event in Figure 8) are not identifiable and can only be estimated jointly in a single compound parameter (e.g., $\zeta = a^2 \times e_{\mathrm{s1}\leftrightarrow\mathrm{S}} + (1-a)^2 \times e_{\mathrm{s2}\leftrightarrow\mathrm{S}} + e_{\mathrm{S}\leftrightarrow\mathrm{P6}}$ in Figure 8) unless samples from the source populations (*s1*, *s2* and/or *S*) or samples from different populations deriving from the same admixed source are available (LIPSON 2020). Following PATTERSON *et al.* (2012) and LIPSON (2020), this identifiability issue is solved by nullifying the length of admixture edges (i.e., setting $e_{\mathrm{s1}\leftrightarrow\mathrm{S}} = e_{\mathrm{s2}\leftrightarrow\mathrm{S}} = 0$ in the above example) which may lead to some overestimation of the divergence (branch length) of the admixed population (here *P6*) from its direct admixed ancestor (here S) if the two source populations (here *s1* and *s2*) have experienced strong divergence since their separation from the graph connecting the other populations (which is not the case in the simulated example in which both $e_{\mathrm{s1}\leftrightarrow\mathrm{S}}$ and $e_{\mathrm{s2->S}}$ are equal to 0). This differs from the choice made by PICKRELL and PRITCHARD (2012) in the *Treemix* model consisting of setting $e_{\mathrm{S}\leftrightarrow\mathrm{P6}} = 0$ and $e_{\mathrm{s2}\leftrightarrow\mathrm{S}} = 0$ (if $a > 0.5$) or $e_{\mathrm{s1}\leftrightarrow\mathrm{S}} = 0$ (if $a < 0.5$).

The graph incidence matrix plays a pivotal role for graph fitting since the *fit.graph* function (see section \ref{graphparams2} use it to build the model equations. A (simplified) symbolic representation of these model equations together with expression for the parameters $F_2$, $F_3$ and $F_4$ can also be generated from a *graph.params* object with the *graph.params2symbolic.fstats* function (see section 7.1).

##### 5.1.2 Generating a *graph.params* object with f-statistics estimates for admixture graph fitting

The f-statistics estimates ($f_2$ and $f_3$) need to be included within the *graph.params* object to allow further fitting of the admixture graph (i.e., estimation of its edge lengths and, if any, admixture proportions) with the *fit.graph* function (see section 5.2). This can be done by providing the *generate.graph.params* function with an *fstats* object including all the $f_2$ and $f_3$ statistics involving the graph leaves (which is usually the case) and block-jackknife estimates of their standard errors and error covariance matrix (i.e., the *compute.fstats* function used to generate the *fstats* object must have been run with *nsnp.per.bjack.block*>0), otherwise an error message is returned[19]. If $n_l$ the number of leaves of the admixture graph and $A$ a "reference" population among the $n_l$ ones[20], the *generate.graph.params* function selects the estimates (block-jackknife means) for the $n_l - 1$ $f_2$ statistics of the form $f_2(A;B)$ (with $B$ a leave population other than $A$) and the $\binom{n_l-1}{2}$ $f_3$ statistics of the form $F_3(A;B,C)$ (with $B$ and $C$ two leave populations other than $A$) which form the basis of the f-statistics vector (see section 4). The basis f-statistics are stored in the *f2.target* and *f3.target* slots of the resulting *graph.params* object (with their corresponding names available in the *f2.target.pops* and *f3.target.pops* slots respectively). In addition, as required to further fit the admixture graph (see section 5.2), the $\frac{n_l(n_l-1)}{2}$ by $\frac{n_l(n_l-1)}{2}$ covariance matrix of the basis f-statistics is stored in the *f.Qmat* slot of the *graph.params* object. Optionally, if available in the *fstats* object, estimates of the leaves heterozygoties (needed to scale fitted branch lengths in drift units, see 5.2.1) are stored in the *Het* slot of the *graph.params* object. The following code shows how to generate a *graph.params* object for the example (allele count) data:

```
sim.graph.params<-generate.graph.params(sim.graph,fstats = sim6p.allelecount.fstats)
```

```
Total Number of Parameters: 11 (10 edges lengths + 1 adm. coeff.)
Total Number of Statistics: 15 (5 F2 and 10 F3)
```

As shown above, the functions returns the number of parameters $n_{\mathrm{par}} = 2n_l + 2n_a - 3$ of the admixture graph where $n_a$ and $n_l$ are the number of admixture events $n_a$ and the number of leaves $n_l$, respectively (LIPSON 2020). Note that the plotting properties of the *graph.params* object remain the same whether *fstats* information is included or not (i.e., the *plot* function may be used as above to generate the representation

[19]Conversely, the *fstats* object may include f-statistics involving populations other than the graph leaves, the *generate.graph.params* function selecting only the f-statistics relevant for the fitting of the input graph

[20]by default, $A$ is the first population in the vector of leaves but, although of limited interest, another reference population may be specified with the *popref* argument

displayed in Figure 8).

#### 5.2 Fitting a graph with the *fit.graph* function

The *fit.graph* functions provides estimate of the parameters (i.e., edge length and admixture proportions) of an admixture graph stored in a *graph.params* object as detailed in GAUTIER *et al.* (2021) and directly inspired by PATTERSON *et al.* (2012). Briefly, let $\widehat{\mathbf{f}}$ represent the vector of length $\frac{n_l(n_l-1)}{2}$ (where $n_l$ is the number of graph leaves) of the estimated $f_2$ and $F_3$ basis f-statistics[21] and $\mathbf{g}(\mathbf{e};\mathbf{a})=\mathbf{X}(\mathbf{a})\times\mathbf{e}$ the vector of these expected basis f-statistics values given the vector of graph edges lengths $\mathbf{e}$ and the incidence matrix $\mathbf{X}(\mathbf{a})$ that depends on the structure of the graph and the admixture rates $\mathbf{a}$[22]. Let further $\mathbf{Q}$ represent the $\frac{n_l(n_l-1)}{2}$ by $\frac{n_l(n_l-1)}{2}$ covariance matrix of the basis F-statistics estimates estimated by block-jackknife and stored in the slot *f.Qmat* of the input *graph.params* object (see 5.1.2)[23]. The function attempts to find the graph parameter values ($\hat{\mathbf{e}}$ and $\hat{\mathbf{a}}$) that minimize a cost (score of the model) defined as $S(\mathbf{e};\mathbf{a})=\left(\widehat{\mathbf{f}}-\mathbf{g}(\mathbf{e};\mathbf{a})\right)'\mathbf{Q}^{-1}\left(\widehat{\mathbf{f}}-\mathbf{g}(\mathbf{e};\mathbf{a})\right)$. As mentioned by PATTERSON *et al.* (2012), given admixture rates $\mathbf{a}$, $S(\mathbf{e};\mathbf{a})$ is actually quadratic in the edge lengths $\mathbf{e}$ allow the *fit.graph* function to rely on the Lawson-Hanson non-negative linear least squares algorithm implemented in the *nnls* function (*nnls* package) to estimate the vector $\hat{\mathbf{e}}$ that minimizes $S(\mathbf{e};\mathbf{a})$ (subject to the constraint of positive edge lengths). Full minimization of $S(\mathbf{e};\mathbf{a})$ is thus reduced to the identification of the admixture rates $\mathbf{a}$ which is performed using the L-BFGS-B method[24] implemented in the *optim* function (*stats* package). The *eps.admix.prop* argument allows specifying the lower and upper bound of the admixture rates to *eps.admix.prop* and *1-eps.admix.prop* respectively. In addition, assuming $\widehat{\mathbf{f}}\sim\mathrm{N}\left(g(\hat{\mathbf{e}};\hat{\mathbf{a}}),\mathbf{Q}\right)$[25], $S(\hat{\mathbf{e}};\hat{\mathbf{a}})=-2log(L)-K$ where $L$ is the likelihood of the fitted graph and $K=n\log(2\pi)+\log(|\mathbf{Q}|)$ allowing to almost directly derive a $BIC$ (Bayesian Information Criterion) of the fitted graph from the optimized score $S(\hat{\mathbf{e}};\hat{\mathbf{a}})$[26]. The BIC may be used for comparison of different admixture graphs (see section 5.3) providing they were all fitted based on the same vector of f-statistics (i.e., they include the same set of populations). Indeed, when comparing two graphs $\mathcal{G}_1$ and $\mathcal{G}_2$ with $BIC$ equal to $BIC_1$ and $BIC_2$, $\Delta_{12}=BIC_2-BIC_1\simeq 2\log\left(BF_{12}\right)$ where $BF_{12}$ is the Bayes Factor associated to $\mathcal{G}_1$ and $\mathcal{G}_2$ graph comparison (eq. 9, KASS and RAFTERY 1995). The (slightly) modified Jeffreys' rule proposed by KASS and RAFTERY (1995) might further be used to evaluate to which extent the data support $\mathcal{G}_1$ or $\mathcal{G}_2$ with e.g., $\Delta_{12}>6$ (respectively $\Delta_{12}>10$) providing "strong" (respectively "very strong") evidence in favor of $G_1$[27].

The *fit.graph* function returns an object of class *fitted.graph* detailed in the documentation page accessible with the following command (or the *?* operator):

```
help(fitted.graph)
```

The following code shows how to fit the example graph stored in the *graph.params* object *sim.graph.params* generated above and some features of the resulting *fitted.graph* object:

```
sim.fittedgraph<-fit.graph(sim.graph.params)
```

```
Starting estimation of admixture rates (LBFGS score optimisation)
    Initial score= 316.0469
Estimation ended in 0 m  0 s
    Final Score: 1.004106
    BIC: 276.6086
```

[21] stored in the *f2.target* and *f3.target* slots of the input *graph.params* object, see section 5.1.2

[22] If there is no admixture in the graph, $\mathbf{X}(\mathbf{a})$ only contains only 0 or 1

[23] The argument *Q.lambda* of the *fit.graph* function may be used to add a small constant (e.g., $10^{-4}$) to all to the diagonal elements of $\mathbf{Q}$ (i.e., the variance of the basis f-statistics estimates) as proposed by PATTERSON *et al.* (2012; see also LIPSON 2020) to avoid numerical problems. Note that *Qmat.diag.adjust* is always set 0 in the final estimate of the score $S$ or the $BIC$

[24] Limited-memory Broyden-Fletcher-Goldfarb-Shanno algorithm with box constraints

[25] i.e., the observed vector of f-statistics is multivariate normal distributed around the expected $g(e;a)$ vector specified by the admixture graph and the covariance structure empirically estimated

[26] $\mathrm{BIC}=S+n_{\mathrm{par}}\log\left(\frac{1}{2}n_l(n_l-1)\right)-\frac{1}{2}n_l(n_l-1)\log(2\pi)-\log(|\ \mathbf{Q}\ |)$

[27] The two thresholds of 6 and 10 on $\Delta_{\mathrm{BIC}}$ corresponds to thresholds of 13 and 21 deciban (units of $10\log_{10}$ scale) on BF. The original Jeffreys' rule considered $BF_{12}$ thresholds of 10, 15 and 20 decibans (corresponding to $\Delta_{\mathrm{BIC}}$ thresholds of 4.6, 6.9 and 9.2) as "strong," "very strong" and "decisive" evidence in favor of model 1.

```r
#Estimated edge lengths
sim.fittedgraph@edges.length
```

```
    P7<->P1     s1<->P2     s2<->P3      S<->P6     P8<->s2     P7<->s1     P9<->P4     P9<->P5     P8<->P7      R<->P8      R<->P9
0.004079551 0.001970516 0.002148837 0.002111818 0.006334944 0.002228311 0.012858490 0.012529775 0.004160468 0.006445943 0.006445943
```

```r
#Estimated admixture proportion
sim.fittedgraph@admix.prop
```

```
[1] 0.2478947
```

```r
#Final Score
sim.fittedgraph@score
```

```
[1] 1.004106
```

```r
#BIC
sim.fittedgraph@bic
```

```
[1] 276.6086
```

The *fitted.graph* object also stores the output results from the *optim* optimization function in the slot *optim.results*. For instance, in the example below, convergence was reached without any issue (*convergence=0*)[28]:

```r
#optim function results (list)
sim.fittedgraph@optim.results
```

```
$par
[1] 0.2478947

$value
[1] 1.004106

$counts
function gradient
       8        8

$convergence
[1] 0

$message
[1] "CONVERGENCE: REL_REDUCTION_OF_F <= FACTR*EPSMCH"
```

As for *graph.params* objects (see section 5.1.1), the *plot* function may be directly applied to *fitted.graph* objects to plot the admixture graph with estimated parameter values[29]. The *admix.fact* and *edge.fact* argument of the *fit.graph* function allow to apply a multiplying factor to the printed branch lengths and admixture proportions (by default *admix.fact*=100, i.e., admixture proportions are printed in %; and *edge.fact*=1000, i.e. edge lengths are printed in ‰). Figure 9 shows the fitted example graph and was generated with the following command:

```r
plot(sim.fittedgraph)
```

[28]In case of convergence problem (i.e., *convergence* not equal to 0), a message detailing execution error in the optimization algorithm is stored in the *optim.results* component named *message*. For more details, see the documentation for the *optim* function with the command *?optim* providing the package *optim* is loaded

[29]the graph is also coded in *dot* format with a character vector stored in the *dot.graph* slot of the resulting *fitted.graph* object. As for *generate.graph.params*, a *dot* file may also be printed out by specifying an output file name prefix with the *outfileprefix* argument of the *fit.graph* function (or using the command *writeLines(x@dot.graph,con=outfile)* where *x* is the *fitted.graph* object and *outfile* is the desired named of the dot output file). See 5.1.1 for more details on how to externally export and customize *dot* files

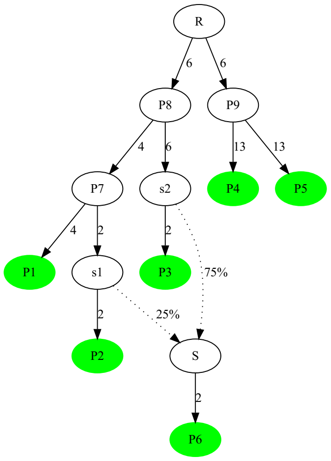

Figure 9: Plot of the admixture graph specifying the simulated scenario with fitted parameter values (x1000 for edge lengths)

> **Notice**
> The two edges from the root node of the graph are not identifiable and only their joint lengths can be estimated. The root position is arbitrarily set in the mid position (i.e., the two root edges have the same length by construction as shown in Figure 9).

##### 5.2.1 Scaling of branch lengths in drift units

The estimated edge lengths are in the same scale as the other f-statistics which limits their interpretation since they strongly depend on the SNP ascertainment process (see section 4). Lipson *et al.* (2013) showed however that the lengths may be approximately scaled in genetic drift units (i.e., in units of $\tau = \frac{t}{2N_e}$ where $t$ is a number of generations and $N_e$ is the diploid effective population along the branch) using estimates of overall marker heterozygosities within (i.e., $1 - Q_1$) or across (i.e., $1 - Q_2$) population (Gautier *et al.* 2021). Briefly, for a given edge $\mathrm{P} \leftrightarrow \mathrm{C}$ relating a child node $C$ to its parent node $P$ with an (unscaled) estimated branch length $\hat{e}_{\mathrm{P}\leftrightarrow\mathrm{C}}$, $\hat{\tau}_{\mathrm{P}\leftrightarrow\mathrm{C}} = 2\frac{\hat{e}_{\mathrm{P}\leftrightarrow\mathrm{C}}}{\widehat{h_P}}$ where $\hat{\tau}_{\mathrm{P}\leftrightarrow\mathrm{C}}$ is the estimated branch length scaled in drift units and $\widehat{h_P}$ is the estimated heterozygosity in the (parent) node $P$. The parent node heterozygosities can be estimated from leaves to root by using the property $h_P = h_C + 2e_{\mathrm{P}\leftrightarrow\mathrm{C}} = 0$ that relate the child $C$ and parent $P$ node heterozygosities ($h_C$ and $h_P$ respectively) and $e_{\mathrm{P}\leftrightarrow\mathrm{C}}$ (Lipson *et al.* 2013). When the *drift.scaling* argument is set to *TRUE*[30], the *fit.graph* function returns the edge lengths scaled in drift units in a slot named *edges.length.scaled* together with the estimated node heterozygosities (*nodes.het* slot) as shown below with the example data:

```
sim.fittedgraph.scaled<-fit.graph(sim.graph.params,drift.scaling = TRUE,verbose=FALSE)
```

Note that the obtained results are the same as above with no drift scaling since the latter is a post-processing step independent of admixture graph parameters estimation

```
#Estimated edge lengths
sim.fittedgraph.scaled@edges.length.scaled
```

```
   P7<->P1     s1<->P2     s2<->P3      S<->P6     P8<->s2     P7<->s1     P9<->P4     P9<->P5     P8<->P7      R<->P8      R<->P9
0.04964054 0.02405296 0.02631275 0.02516072 0.07378969 0.02711439 0.14321071 0.13954967 0.04846131 0.07700936 0.07700936
```

When plotting the resulting *fitted.graph* objects, branch lengths are displayed in drift scaled units as shown in the Figure 10 below obtained with the following command:

[30]providing the input *graph.params* object includes estimates of within-population heterozygosities (see section 5.1.2)

```r
plot(sim.fittedgraph.scaled)
```

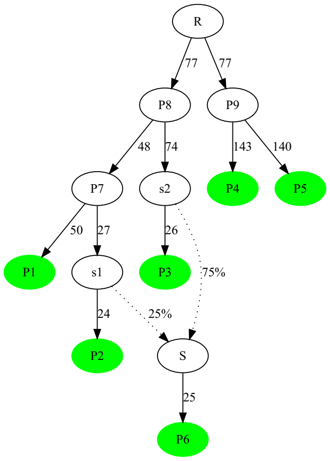

Figure 10: Plot of the admixture graph specifying the simulated scenario with fitted edge lengths scaled in drift units (x1,000)

The estimated branch lengths are close to the simulated ones (Figure 1) as represented below (see GAUTIER *et al.* 2021 for a more in-depth exploration of the accuracy of the estimates):

```
             P7<->P1    s1<->P2    s2<->P3     S<->P6    P8<->s2    P7<->s1   P9<->P4   P9<->P5    P8<->P7 R<->P8+R<->P9
Estimated 0.04964054 0.02405296 0.02631275 0.02516072 0.07378969 0.02711439 0.1432107 0.1395497 0.04846131     0.1540187
Simulated 0.05000000 0.02500000 0.02500000 0.02500000 0.07500000 0.02500000 0.1500000 0.1500000 0.05000000     0.1500000
```

##### 5.2.2 Estimating 95% confidence intervals of the estimated parameters values

Calling the *fit.graph* function with the *compute.ci* argument set to *TRUE* allows deriving (rough) 95% confidence intervals for the admixture graph parameter estimates. The procedure considers each parameter in turn (the other parameters being set to their estimated values) and consists of interpreting the score difference $S_\nu - S^\star$ (where $S^\star$ is the fitted graph score associated with estimated parameter value $\nu^\star$ and $S_\nu$ is the score associated with a parameter value $\nu \neq \nu^\star$) as a likelihood-ratio test statistics following a $\chi^2$ distribution with one degree of freedom (see GAUTIER *et al.* 2021 for details). The lower and upper boundaries $\nu_{\min}$ and $\nu_{\max}$ of the 95% CI (such $S_\nu - S^\star < 3.84$ for all $\nu_{\min} < \nu < \nu_{\max}$) are then simply computed using a bisection method (with a $10^{-4}$ precision threshold).

```r
sim.fittedgraph.with.ci<-fit.graph(sim.graph.params,compute.ci=TRUE,
                                   drift.scaling = TRUE,verbose = FALSE)
#95% CI for the admixture proportion

```

```
   95% Inf.  95% Sup.
a 0.2354879 0.2604726
```

```r
#95% CI for edge length (including drift scaled as drift.scaling=TRUE)

```

```
           95% Inf.    95% Sup. 95% Inf. (drift scaled) 95% Sup. (drift scaled)
P7<->P1 0.003888322 0.004322695              0.04731364              0.05259916
s1<->P2 0.001785780 0.002153260              0.02179800              0.02628362
s2<->P3 0.001947384 0.002392453              0.02384593              0.02929586
S<->P6  0.001913835 0.002294537              0.02280190              0.02733768
P8<->s2 0.006037994 0.006638187              0.07033080              0.07732187
P7<->s1 0.002019406 0.002471907              0.02457242              0.03007851
P9<->P4 0.012406433 0.013340492              0.13817596              0.14857899
P9<->P5 0.012040331 0.013011938              0.13409851              0.14491974
```

```
P8<->P7 0.003835432 0.004464375                0.04467527                0.05200122
R<->P8  0.006219328 0.006686937                0.07430200                0.07988849
R<->P9  0.006219328 0.006686937                0.07430200                0.07988849
```

The simulated values are all within the estimated 95% CI of the estimated value except for the long $e_{P9\leftrightarrow P4}$ and $e_{P9\leftrightarrow P5}$ branches that are slightly higher than the upper boundary. Note also that if the 95% CI for the admixture proportion $a$ contains the simulated value of 0.25, it is narrower than the one that may be derived from the block-jackknife estimate of the $F_4$-ratio standard error (see section 4.3 and Gautier *et al.* 2021 for a more thorough evaluation of the estimated 95% CI).

##### 5.2.3 Assessing the fit of the graph with the *compare.fitted.fstats* function

As outlined by Patterson *et al.* (2012) and Lipson (2020), a straightforward but highly informative approach to assess the fit of the graph is to compare the f-statistics derived from the fitted admixture graph parameters to the estimated ones, i.e., to evaluate to which extent the fitted F-statistics lie within the confidence intervals of the estimated ones. This may be summarized by computing a Z-score of the residuals for each f-statistics as $Z = \frac{f-g}{\sigma_g^2}$ where $f$ and $g$ stand for the fitted and estimated values respectively and $\sigma_g^2$ is the standard error of $g$. This information is available for the basis f-statistics in the *fitted.outstats* slot of the *fitted.graph* object generated by the *fit.graph* functions, as illustrated below:

```
#Fitted basis F-stats
sim.fittedgraph.scaled@fitted.outstats
```

```
         Stat. value Fitted Value     Z-score
P1,P2    0.008274108  0.008278377  0.03287705
P1,P3    0.016680443  0.016723801  0.20561876
P1,P4    0.033971079  0.033990395  0.05587989
P1,P5    0.033727224  0.033661681 -0.18890298
P1,P6    0.012237561  0.012265163  0.16197672
P1;P2,P3 0.004048392  0.004079551  0.26473869
P1;P2,P4 0.004087474  0.004079551 -0.05972586
P1;P2,P5 0.004149524  0.004079551 -0.52164304
P1;P2,P6 0.004610668  0.004631937  0.19288766
P1;P3,P4 0.008230291  0.008240019  0.05210483
P1;P3,P5 0.008244937  0.008240019 -0.02672913
P1;P3,P6 0.011937638  0.011973206  0.20637614
P1;P4,P5 0.021148713  0.021131906 -0.06164584
P1;P4,P6 0.007189623  0.007208661  0.11673856
P1;P5,P6 0.007234212  0.007208661 -0.16250690
```

However, the fit should be evaluated for all the f-statistics (not only those forming the f-statistics vector-space basis used to fit the admixture graph) with the *compare.fitted.fstats* function. This may in turn provide insights into those populations (or graph edges) leading to poor fit (Lipson 2020). As shown below, the function requires the original *fstats* object (that may contain f-statistics for additional populations not represented in the admixture graph) and the *fitted.graph* object. It then produces a matrix with information on all the fitted stats and the *n.worst.stats* (by default *n.worst.stats*=5) f-statistics, i.e. with the lowest absolute Z-score, are directly printed in the console:

```
sim.fitted.fstats.comp<-compare.fitted.fstats(sim6p.allelecount.fstats,sim.fittedgraph)
```

```
5 Worst fit for:
               Estimated       Fitted   Z-score
P1,P2;P3,P5 1.011325e-04 0.000000e+00 -0.8442359
P1,P2;P5,P6 4.611437e-04 5.523863e-04  0.7459512
P1,P2;P4,P5 6.205022e-05 3.469447e-18 -0.6662270
P2;P1,P5    4.124584e-03 4.198826e-03  0.5401737
P1;P2,P5    4.149524e-03 4.079551e-03 -0.5216430
```

```
#Information on the last five fitted F-statistics
tail(sim.fitted.fstats.comp)
```

```
                   Estimated         Fitted    Z-score
P2,P6;P3,P4 -4.787097e-03 -4.764545e-03  0.16513037
P2,P6;P3,P5 -4.804558e-03 -4.764545e-03  0.27257790
P2,P6;P4,P5 -1.746147e-05  0.000000e+00  0.16092207
P3,P4;P5,P6 -1.765179e-02 -1.765643e-02 -0.01671976
P3,P5;P4,P6 -1.762185e-02 -1.765643e-02 -0.12418834
P3,P6;P4,P5  2.994286e-05  6.938894e-18 -0.39137603
```

As shown above, no outlying fitted f-statistics (e.g., with $|Z| > 2$) is observed on the example providing strong support for the fitted admixture graph.

##### 5.3 Adding a new leaf to an existing graph

The *add.leaf* function allows to perform iterative calls to the *fit.graph* function in order to evaluate all possible connections of a given leaf (population) to an existing graph with non-admixed and/or admixed edges. Three input arguments are required:

- a graph specified within a *graph.params* object (obtained with the *generate.graph.params* function, see section 5.1.1) or a *fitted.graph* object (obtained with the *fit.graph*, *add.leaf* or *graph.builder* functions, see sections 5.2 and 6.2 below allowing more convenient exploration of the admixture graph space via recursive calls)
- the name of the leaf to add (*leaf.to.add* argument)
- an *fstats* object (see 4) containing *a minima* estimates of all the $f_2$ and $f_3$ statistics (and their standard errors) involving the leaves of the input graph and the leaf to add

By default the function tests all the possible positions of the candidate leaf (*leaf.to.add*) to the graph with non-admixed (including a new rooting with the candidate leaf as an outgroup) or admixed edges. If $n_e$ is the number of non-admixed edges in the original graph, the number of tested graphs equals $n_e + 1$ for non-admixed candidate edges[31] and $\frac{1}{2}n_e(n_e - 1)$ for admixed candidate edges[32]. Optional arguments may allow disabling the evaluation of non-admixed (by setting the *only.test.non.admixed.edges* argument to *TRUE*) or admixed (by setting the *only.test.admixed.edges* argument to *TRUE*) candidate edges.

The object returned by the *add.leaf* function is a list consisting of:

- an element named *n.graphs* corresponding to the number of tested graphs
- an element named *fitted.graphs.list* which consists of a list of *fitted.graph* objects (indexed from 1 to n.graphs and in the same order as the list "graphs") containing the *fit.graph* function results for each candidate graph
- an element named *best.fitted.graph* which is the *fitted.graph* object associated to the candidate graph with minimal *BIC* (see 5.2) among all the *n.graphs* graphs within *fitted.graphs.list*.
- an element named *bic* which is a vector consisting of the *n.graphs BIC* (indexed from 1 to *n.graphs* and in the same order as the *fitted.graphs.list* list).

Use of the *add.leaf* function is illustrated below on the example data by evaluating the connection of the *P6* population on the graph (actually tree) connecting the five other populations (*P1*, *P2*, *P3*, *P4* and *P5*) specified according to the simulated topology in a *graph.params* object (5.1.1) named *sim5p.tree.params* and plotted in Figure 11:

```
sim5p.tree<-sim.graph<-rbind(c("P1","P7",""),c("P2","P7",""),c("P3","P8",""),
                             c("P7","P8",""),c("P4","P9",""),c("P5","P9",""),
                             c("P8","R",""),c("P9","R",""))
sim5p.tree.params<-generate.graph.params(sim5p.tree)
#Note: fstats object is not necessary at this stage
plot(sim5p.tree.params)
```

[31] The newly added node is named "N-"*leaf.to.add*

[32] The three added nodes are named "S-"*leaf.to.add*, "S1-"*leaf.to.add* and "S2-"*leaf.to.add* and the admixture proportions are named with a letter (A to Z depending on the number of admixed nodes already present in the graph)

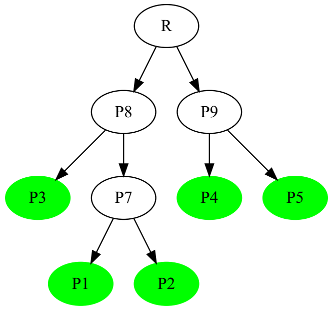

Figure 11: Plot of the five-population graph on which to add the $P6$ population with the *add.leaf* function

All the possible positions of the *P6* population on the *sim5p.tree* graph (here using the f-statistics estimated on the simulated allele count data, see section 4), are tested as follows:

```
add.P6<-add.leaf(sim5p.tree.params,leaf.to.add="P6",
                 fstats=sim6p.allelecount.fstats,
                 verbose=FALSE,drift.scaling=TRUE)
```

Note that the *verbose* option set to *FALSE* allows disabling the printing of the progress and timing of each analysis (which may be useful in practice) and the *drift.scaling* option set to *TRUE* allows passing it to each *fit.graph* call to obtained estimates of branch lengths in drift units (see section 5.2).

The graph with the lowest *BIC* among all the tested graphs may then be plotted by calling the *plot* function on the corresponding *fitted.graph* object stored in the *best.fitted.graph* element of the *add.P6* output list with the following command that generates Figure 12:

```
plot(add.P6$best.fitted.graph)
```

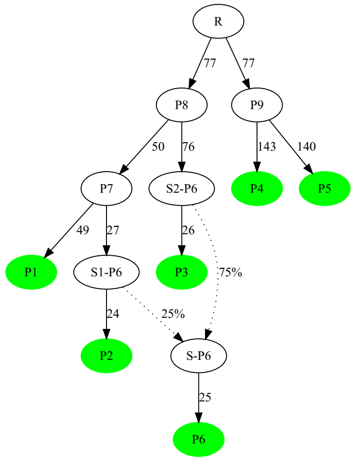

Figure 12: Plot of the graph with the lowest BIC among all the possible graphs connecting $P6$ to the five-population tree tested by the *add.leaf*. The fitted edge lengths are in drift units (x1000) since *drift.scaling* argument was set to *TRUE* when calling *add.leaf*.

The best fitting graph based on *BIC* criterion (stored in the *best.fitted.graph* slot of the output list) is in perfect agreement with the simulated scenario (Figure 1). It actually corresponds to the one directly fitted using the simulated scenario which was represented in Figure 10 above (only the names of the nodes involved in the admixture events differ since automatically given by the *add.leaf* function). In addition, comparisons of the *BIC* of the different graphs provide strong support in favor of this best fitting graph, the second lowest *BIC* being more than *86* units larger i.e., far above the threshold of 10 for very strong evidence (see section 5.2):

```
#D_BIC w.r.t. best fitted BIC
D_BIC=add.P6$bic-add.P6$best.fitted.graph@bic
#5 First lowest DeltaBIC (the first value of zero corresponding to the best fitted graph)
head(sort(D_BIC))
```

```
[1]   0.00000  86.31675  86.31675 279.48967 284.90577 284.90577
```

### 6 Building admixture graph from scratch

Lipson *et al.* (2013) proposed a two-step approach (implemented in the *MixMapper* software[33]) to build admixture graph when prior knowledge about history and relationships of investigated population is limited (which is usually the case). It consists of first building a scaffold tree of unadmixed populations and then adding the remaining populations successively on the graph. Such a supervised approach nevertheless requires to carefully assess at each step the graph fit and possibly try different ordering in the inclusion of populations (or removal of some populations). The *poolfstat* package provides functions to help building scaffold trees that may further be used as input tree for the *add.leaf* function previously described above (5.3) to implement the Lipson *et al.* (2013) two-step approach.

#### 6.1 Building scaffold trees of unadmixed populations

In the absence of admixture, the $f_2$ statistics among all pairs of populations are expected to be additive along the paths of the (binary) tree summarizing the history of the populations (Lipson *et al.* 2013). As a result, the (unrooted) tree topology and branch lengths connecting unadmixed populations may be inferred with a neighbor-joining algorithm to derive a scaffold tree for further admixture graph construction. Based on the estimated f-statistics stored in a *fstats* object, the functions described below allows to *i)* identify candidate sets of unadmixed populations among the genotyped ones (*find.tree.popset* function); *ii)* infer a neighbor-joining scaffold tree from a candidate set of unadmixed populations (*rooted.njtree.builder* function); and *iii)* to infer root position based on the consistency of within population heterozygosities between the two resulting partitions of rooted trees (see p1799 in Lipson *et al.* 2013).

##### 6.1.1 The *find.tree.popset* function to identify sets of candidate scaffold populations

The *find.tree.popset* function selects maximal sets of unadmixed populations from an *fstats* object[34]. The procedure involves *i)* discarding all the populations showing a significantly negative $f_3$ at a significance theshold specified with the *f3.zcore.threshold* argument (equal to $-1.65$ by default, i.e., 95% significance threshold, see section 4.1.2); and *ii)* keeping only sets of populations with all possible quadruplets passing the $f_4$-based test of treeness i.e., with an absolute $f_4$ Z-score lower than a threshold specified with the *f4.zcore.threshold* argument (equal 2 by default, i.e., 95% significance threshold, see section 4.1.3). The latter step is implemented via a greedy algorithm (that may be run in parallel by specifying a number of threads with the *nthreads* argument) consisting of trying to extend the size of the population sets from all sets of four populations after adding candidate populations one at a time. If the number of populations is large, this algorithm may take some times. Note that increasing (respectively decreasing) *f3.zcore.threshold* toward value closer to 0 may allow decreasing (respectively increasing) the number of initial candidate populations to be tested for inclusion in a set. Similarly, increasing (respectively decreasing) *f4.zcore.threshold* may allow increasing (respectively decreasing) the size of the sets. When applied to the example allele count and read count data, a single set of 5 unadmixed populations (*P1*, *P2*, *P3*, *P4* and *P5*) is retrieved as expected from the simulated scenario (Figure 1):

```
# count data
scaf.pops<-find.tree.popset(sim6p.allelecount.fstats,verbose=FALSE)
scaf.pops$pop.sets
```

[33] http://cb.csail.mit.edu/cb/mixmapper/

[34] providing it was Z-scores were estimated for $f_3$ and $f_4$ statistics, i.e., that block-jackknife estimates of s.e. were carried out, see section 4

```
        [,1] [,2] [,3] [,4] [,5]
PopSet1 "P1" "P2" "P3" "P4" "P5"
```

```
# 30X Pool-Seq data
scaf.pops<-find.tree.popset(sim6p.readcount30X.fstats,verbose=FALSE)
scaf.pops$pop.sets
```

```
        [,1]    [,2]    [,3]    [,4]    [,5]
PopSet1 "Pool1" "Pool2" "Pool3" "Pool4" "Pool5"
```

As previously mentioned (section 4), for a given set consisting of n populations, a total of $3\binom{n}{4} = \frac{1}{8}n(n-1)(n-2)(n-3)$ quadruplets can be formed. In other words, a given set of four populations A, B, C and D is actually represented by only three quadruplets representative of the three possible unrooted tree topologies[35] i) (A,B;C,D); ii) (A,C;B,D); and iii) (A,D;B,C). Among these, only a single quadruplet is expected to pass the treeness test (i.e., if the correct unrooted tree topology is (A,C;B,D), then the absolute value of the Z-scores associated to f4(A,B;C,D) and f4(A,D;B,C) or their equivalent will be high). For each of identified sets of presumably unadmixed populations, the list of the $\binom{n}{4}$ quadruplets passing the treeness test is given in the *passing.quadruplets* element of the output list as illustrated below:

```
# list of the 15 quadruplets passing the treeness test for the identified set
scaf.pops$passing.quaduplets
```

```
        [,1]                      [,2]                      [,3]
PopSet1 "Pool1,Pool3;Pool4,Pool5" "Pool2,Pool3;Pool4,Pool5" "Pool1,Pool2;Pool4,Pool5"
        [,4]                      [,5]
PopSet1 "Pool1,Pool2;Pool3,Pool4" "Pool1,Pool2;Pool3,Pool5"
```

In addition, for each of the identified sets, the range of variation of the passing quadruplets is given in the *Z_f4.range* element of the output list:

```
scaf.pops$Z_f4.range
```

```
        Min. |Zscore| Max. |Zscore|
PopSet1    0.02584028     0.8067878
```

When several sets are identified, this information may be useful to prioritize the sets of unadmixed populations (e.g., via a minimax criterion consisting of choosing the set of populations that has the lowest maximal absolute Z-score for its underlying quadruplets that pass the treeness test).

###### 6.1.2 The *rooted.njtree.builder* to building (and root) a tree of candidate scaffold populations

The *rooted.njtree.builder* allows first building a Neighbor-Joining[36] of a set of presumably unadmixed populations (as obtained e.g., from the *find.tree.popset* functions) given as a vector of population names in the *pop.sel* argument based on the matrix of their pairwise $f_2$ stored in the provided *fstats* object (*fstats* argument). The resulting (unrooted) tree is then rooted by relying on the property that root $R$ heterozygosity $h_R = 1 - Q_2^{A,B}$ estimated from all the possible pairs of populations $A$ and $B$ that satisfies the property of being only connected through R in the tree (i.e., $A$ and $B$ each belong to one of the two tree partitions defined by the $R$) should be consistent (Lipson *et al.* 2013). In other words, the most likely rooted tree among the $(2n_l - 3)$ possible ones should be the one displaying the narrower range of variation of the $h_R$ estimates. Note that the root position is always placed in the mid-position of the candidate branch.

The object returned by the *rooted.njtree.builder* function is a list consisting of:

- an element named *n.rooted.trees* corresponding to the number of possible rooted binary trees that were evaluated
- an element named *fitted.rooted.trees.list* which consists of a list of *fitted.graph* objects (indexed from 1 to *n.rooted.trees*).

[35] for each of these quadruplets, seven other equivalent combinations formed by permuting populations within each pair can be defined as mentioned in the notice p16

[36] relying on the *nj* function from the popular package *ape* (Paradis *et al.* 2004)

- an element named *best.rooted.tree* which corresponds to the *fitted.graph* object associated with the most likely rooted tree (among all the *fitted.rooted.trees.list* ones) identified as the one displaying the minimal standard deviation over the $h_R$ estimates
- an element named *root.het.est.var* consisting of a matrix of *n.rooted.trees* rows and 4 columns with i) the average estimated root heterozygosity $h_R$ across all the pairs of leaves that are relevant for estimation (see above); ii) the size of the range of variation; iii) the standard deviation of the $h_R$ estimates, and iv) the number of population pairs relevant for estimation
- if *n.edges*>3, an element named *nj.tree.eval* that gives for each evaluated rooted tree, the five $f-$statistics configuration displaying the worst fit, i.e., with the five highest absolute Z-score for the predicted value (obtained by internally calling the *compare.fitted.fstats* function). Note that these are independent of the rooting (so cannot be used to infer the root position).

Use of the *rooted.njtree.builder* function is illustrated below to build the scaffold tree (using Pool-Seq read count data) based on the set of unadmixed populations (identified with *find.tree.popset*) and plotted with the *plot* function applied to the *fitted.graph* object of the resulting list (*best.rooted.tree* element) to generate Figure 13:

```
scaf.tree<-rooted.njtree.builder(fstats=sim6p.readcount30X.fstats,
                                 pop.sel=scaf.pops$pop.sets[1,],plot.nj=FALSE)
```

```
Score of the NJ tree: 0.6884542
Construction of all the 7 possible rooted tree from the NJ tree
(stored as graph in the rooted.graph object of the output list)
```

```
plot(scaf.tree$best.rooted.tree)
```

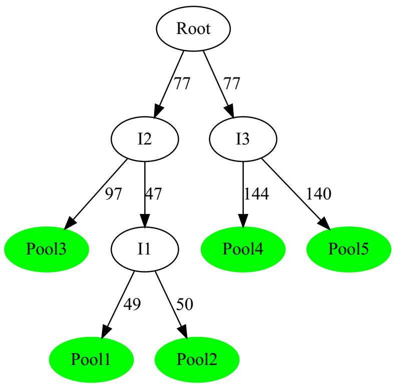

Figure 13: Plot of the rooted scaffold tree of unadmixed populations inferred by *rooted.njtree.builder*. The fitted edge lengths are in a drift scale (x1000).

The fit of the Neighbor-Joining tree can be checked by inspecting the *nj.tree.eval* element:

```
scaf.tree$nj.tree.eval
```

```
                            Estimated      Fitted   Z-score
Pool1,Pool2;Pool3,Pool5 1.011014e-04 0.000000000 -0.8067878
Pool1,Pool2;Pool3,Pool4 6.087576e-05 0.000000000 -0.5098456
Pool2;Pool1,Pool5       4.120591e-03 0.004181198  0.4330143
Pool1;Pool2,Pool5       4.154896e-03 0.004094289 -0.4293005
Pool1,Pool2;Pool4,Pool5 4.022566e-05 0.000000000 -0.4233082
```

The following range of variation of the $h_R$ estimates were obtained for the different possible root position:

```
scaf.tree$root.het.est.var
```

```
           Mean        Range           sd ncomps
Tree1 0.1814754 0.0234940763 0.0115371318      4
```

```
Tree2 0.1810025 0.0228772762 0.0113327886      4
Tree3 0.1857808 0.0155159278 0.0076153034      6
Tree4 0.1907099 0.0007923822 0.0002825614      6
Tree5 0.1880369 0.0113894828 0.0055518382      4
Tree6 0.1878868 0.0111334494 0.0054505348      4
Tree7 0.1833542 0.0152408946 0.0085429545      4
```

The "best" inferred rooted scaffold tree (i.e., the fourth one with both lowest range of $h_R$ variation) is consistent with the simulated scenario. It may further be used as a reference graph to construct the complete admixture graph after adding *Pool6* population using the *add.leaf* function (see 5.3). The obtained graph plotted in Figure 14 with the commands below is very close to the one previously inferred with allele count data (Figure 12) and very strongly supported by the data:

```
add.pool6<-add.leaf(scaf.tree$best.rooted.tree,leaf.to.add="Pool6",
                    fstats=sim6p.readcount30X.fstats,verbose=FALSE,drift.scaling=TRUE)
plot(add.pool6$best.fitted.graph)
```

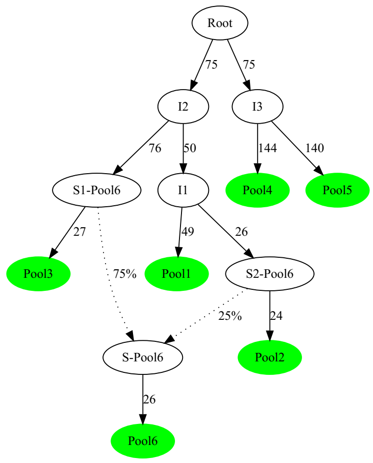

Figure 14: Plot of the graph with the lowest BIC among all the possible graphs connecting *Pool*6 to the scaffold tree of unadmixed population tested by the *add.leaf*. The fitted edge lengths are in drift units (x1000) since *drift.scaling* argument was set to *TRUE* when calling *add.leaf*.

```
#D_BIC w.r.t. best fitted BIC
D_BIC=add.pool6$bic-add.pool6$best.fitted.graph@bic
#5 First lowest DeltaBIC (the first value of zero corresponding to the best fitted graph)
head(sort(D_BIC))
```

```
[1]   0.00000  78.75611  78.75611 254.27229 259.68839 259.68839
```

**Notice**

In practice, the *rooted.njtree.builder* function should be used with caution since both the Neighbor-Joining tree construction and the heterozygosity-based rooting of the tree may be sensitive to long-branch attraction (most particularly if some highly diverged populations are included). The inferred topology may even violate treeness test for some of the quadruplets (see e.g., the empirical example detailed in the Supplementary Vignette V2 by Gautier *et al.* 2021).

#### 6.2 Extending an initial tree (or graph) with the *graph.builder* function

The *graph.builder* function implements an heuristic to carry out a larger exploration of the space of possible graphs (but usually still not exhaustive) obtained from the joint addition of several populations (leaves) given as an input vector (*leaves.to.add* argument) to an existing graph (as generated using the *rooted.njtree.builder* function described above or included in a *graph.params* or *fitted.graph* object) or a list of graphs. The algorithm consists of adding the leaves in the order of the input vector to each of the graphs stored in a heap via successive calls to the *add.leaf* function (section 5.3). More precisely, the heap first consists of the initial input graph (or list of graph) and at each iteration, the function *add.leaf* is used to evaluate all the possible connections of each candidate leaf (with non-admixed or admixed edges) to all the graphs of the heap. For each of the latter, the newly fitted graphs displaying a *BIC* less than *heap.dbic* units (set to 6 by default) away from the best fitting graph (i.e., the one with the lowest *BIC*) included in the new heap. Once all the graphs have been evaluated, if the heap contains more than *max.heap.size* (set to 25 graphs by default) graphs, only the *max.heap.size* graphs with the lowest BIC are kept in the heap. Finally, after testing the latest candidate leaf, only the graphs with a *BIC* less than *heap.dbic* units away from the graph with lowest *BIC* in the heap are retained in the final list of graphs.

The object returned by the function is a list consisting of:

- an element named *n.graphs* corresponding to the final number of graphs
- an element named *fitted.graphs.list* which consists of a list of *fitted.graph* objects (indexed from 1 to *n.graphs*) containing the *fit.graph* function results for each graph
- an element named *best.fitted.graph* which is the *fitted.graph* object associated to the graph with the lowest *BIC* among all the *n.graphs* graphs included in *fitted.graphs.list*.
- an element named *bic* which is a vector containing the *BIC* of the *n.graphs BIC* (indexed from 1 to *n.graphs* and in the same order as *fitted.graphs.list*).

Use of the *graph.builder* function is illustrated below on the PoolSeq example data by starting from an initial rooted tree constructed with the *rooted.njtree.builder* for the three populations *Pool1*, *Pool3*, *Pool4* and *Pool5*. This tree is extended by successively adding the two remaining populations *Pool2* and *Pool6*:

```
#build an initial 3 population trees with "Pool1","Pool3","Pool4" and "Pool5"
init.tree<-rooted.njtree.builder(fstats=sim6p.readcount30X.fstats,
                                 pop.sel=c("Pool1","Pool3","Pool4","Pool5"),plot.nj=FALSE)
```

```
Score of the NJ tree: 0.0006790273
Construction of all the 5 possible rooted tree from the NJ tree
(stored as graph in the rooted.graph object of the output list)
```

```
#adding the three remaining pops
final.graphs<-graph.builder(x=init.tree$best.rooted.tree,leaves.to.add=c("Pool2","Pool6"),
                            fstats=sim6p.readcount30X.fstats)
```

```
####################
 Adding:  Pool2
21 graphs evaluated in 0 h 0 m 1 s
6 graphs stored in the heap
####################
 Adding:  Pool6
261 graphs evaluated in 0 h 0 m 5 s
7 graphs stored in the heap

Final Number of graphs: 7
(min. BIC= 275.8408 )

Overall Analysis Time: 0 h 0 m 6 s ( 282 graphs evaluated)
```

```
#D_BIC w.r.t. to the "true" graph as identified previously (object add.pool6$best.fitted.graph)
D_BIC=final.graphs$bic-add.pool6$best.fitted.graph@bic
#5 First lowest DeltaBIC (the first value of zero corresponding to the best fitted graph)
head(sort(D_BIC))
```

```
[1] 0.000000 4.667688 4.923348 4.923348 4.923348 5.416100
```

Among the 7 final graphs, the one with the lowest *BIC* is exactly the same as the one plotted in Figure 14 and corresponds to the simulated scenario. It should however be noticed that other alternative graphs are also identified with a good support. Moreover, starting with other population trees (e.g., a three population tree consisting of *Pool1*, *Pool2* and *Pool5*) could result in several graphs with the same support (i.e., $\Delta_{BIC} = 0$) but with a different positioning of the root (not shown). In practice, it may be important to start with scaffold trees that are as large as possible and representative of the structuring of diversity of the represented populations (i.e., not too unbalanced with respect to the leaves to be added). Some prior knowledge about the relationships of some of the populations may also be helpful to that respect. As examplified in the Supplementary Vignette V2 of Gautier *et al.* (2021), it is also highly recommended to test different orders of inclusion (possibly all) of the leaves (as specified in the vector *leaves.to.add*).

### 7 Other utilities

#### 7.1 Symbolic representation of the $F$ parameters, admixture graph equations and the scaled covariance matrix $\mathbf{\Omega}$ with *graph.params2symbolic.fstats*

Given a graph topology relating the populations stored in a *graph.params* object, the *graph.params2symbolic.fstats* functions provide symbolic representation of the model equations used to fit the underlying admixture graph and all the $F_2$, $F_3$ and $F_4$ parameters together with the scaled covariance matrix of population allele frequencies called $\mathbf{\Omega}$ after Gautier (2015). Such representation may be useful for a closer examination of graph properties (or education purposes). The output objects consists of a list with the following elements:

- a character matrix named *model.matrix* consisting of the matrix $\mathbf{M}$ relating the parameters underlying the basis f-statistics and graph edge lengths in the model equations defined as $\mathbf{f} = \mathbf{Mb}$ where $\mathbf{f}$ is the vector of the basis f-statistics (row names of the *model.matrix* $\mathbf{M}$) and $\mathbf{b}$ is the vector of graph edges (column names of *model.matrix* $\mathbf{M}$).
- a character matrix named *omega* consisting of the the scaled covariance matrix of population allele frequencies $\mathbf{\Omega}$ (see e.g., Gautier 2015).
- a character vector *F2.equations* consisting of the symbolic representations of all the $\frac{1}{2}n_l(n_l-1)$ $F_2$ parameters (with edge and admixture proportion parameter names as defined in the *graph.params* object)
- a character vector *F3.equations* consisting of the symbolic representations of all the $\frac{1}{2}n_l(n_l-1)(n_l-2)$ $F_3$ parameters (with edge and admixture proportion parameter names as defined in the *graph.params* object)
- a character vector *F4.equations* consisting of the symbolic representations of all the $\frac{1}{8}n_l(n_l-1)(n_l-2)(n_l-3)$ $F_4$ parameters (with edge and admixture proportion parameter names as defined in the *graph.params* object)

These different equations can also be printed in an output text file (with name specified by the *outfile* argument). The following example shows results obtained using the *graph.params* object *sim.graph.params* generated in 5.1.1 (Figure 8) that specifies the simulation scenario (Figure 1):

```
sim.fstats.sym<-graph.params2symbolic.fstats(sim.graph.params,outfile = "Fstats_equations")
```

```
#Model equations matrix
sim.fstats.sym$model.matrix
```

```
              P7<->P1 s1<->P2 s2<->P3 S<->P6 P8<->s2      P7<->s1 P9<->P4 P9<->P5 P8<->P7      R<->P8 R<->P9
F2(P1,P2)     "1"     "1"     "0"     "0"    "0"          "1"     "0"     "0"     "0"          "0"    "0"
F2(P1,P3)     "1"     "0"     "1"     "0"    "1"          "0"     "0"     "0"     "1"          "0"    "0"
F2(P1,P4)     "1"     "0"     "0"     "0"    "0"          "0"     "1"     "0"     "1"          "1"    "1"
F2(P1,P5)     "1"     "0"     "0"     "0"    "0"          "0"     "0"     "1"     "1"          "1"    "1"
F2(P1,P6)     "1"     "0"     "0"     "1"    "a^2-2*a+1"  "a^2"   "0"     "0"     "a^2-2*a+1"  "0"    "0"
F3(P1;P2,P3)  "1"     "0"     "0"     "0"    "0"          "0"     "0"     "0"     "0"          "0"    "0"
F3(P1;P2,P4)  "1"     "0"     "0"     "0"    "0"          "0"     "0"     "0"     "0"          "0"    "0"
F3(P1;P2,P5)  "1"     "0"     "0"     "0"    "0"          "0"     "0"     "0"     "0"          "0"    "0"
F3(P1;P2,P6)  "1"     "0"     "0"     "0"    "0"          "a"     "0"     "0"     "0"          "0"    "0"
F3(P1;P3,P4)  "1"     "0"     "0"     "0"    "0"          "0"     "0"     "0"     "1"          "0"    "0"
F3(P1;P3,P5)  "1"     "0"     "0"     "0"    "0"          "0"     "0"     "0"     "1"          "0"    "0"
F3(P1;P3,P6)  "1"     "0"     "0"     "0"    "1-a"        "0"     "0"     "0"     "1-a"        "0"    "0"
F3(P1;P4,P5)  "1"     "0"     "0"     "0"    "0"          "0"     "0"     "0"     "1"          "1"    "1"
```

```
F3(P1;P4,P6) "1"     "0"     "0"     "0"    "0"         "0"     "0"     "0"     "1-a"       "0"    "0"
F3(P1;P5,P6) "1"     "0"     "0"     "0"    "0"         "0"     "0"     "0"     "1-a"       "0"    "0"
```

```
#scaled covariance matrix of allele frequencies (Omega)
sim.fstats.sym$omega
```

```
   P1                         P2                               P3
P1 "P7<->P1+P8<->P7+R<->P8" "P8<->P7+R<->P8"                 "R<->P8"
P2 "P8<->P7+R<->P8"         "s1<->P2+P7<->s1+P8<->P7+R<->P8" "R<->P8"
P3 "R<->P8"                 "R<->P8"                         "s2<->P3+P8<->s2+R<->P8"
P6 "P8<->P7*a+R<->P8"       "(P7<->s1+P8<->P7)*a+R<->P8"     "P8<->s2+R<->P8-P8<->s2*a"
P4 "0"                      "0"                              "0"
P5 "0"                      "0"                              "0"
   P6                                                                    P4              P5
P1 "P8<->P7*a+R<->P8"                                                    "0"             "0"
P2 "(P7<->s1+P8<->P7)*a+R<->P8"                                          "0"             "0"
P3 "P8<->s2+R<->P8-P8<->s2*a"                                            "0"             "0"
P6 "S<->P6+(P8<->s2+P7<->s1+P8<->P7)*a^2-2*P8<->s2*a+P8<->s2+R<->P8" "0"             "0"
P4 "0"                                                                   "P9<->P4+R<->P9" "R<->P9"
P5 "0"                                                                   "R<->P9"         "P9<->P5+R<->P9"
```

```
#F2 statistics (first five)
head(sim.fstats.sym$F2.equations)
```

```
[1] "F2(P1,P2) = P7<->P1+s1<->P2+P7<->s1"
[2] "F2(P1,P3) = P7<->P1+P8<->P7+s2<->P3+P8<->s2"
[3] "F2(P1,P6) = P7<->P1+(a^2-2*a+1)*P8<->P7+S<->P6+(a^2-2*a+1)*P8<->s2+a^2*P7<->s1"
[4] "F2(P1,P4) = P7<->P1+P8<->P7+R<->P8+P9<->P4+R<->P9"
[5] "F2(P1,P5) = P7<->P1+P8<->P7+R<->P8+P9<->P5+R<->P9"
[6] "F2(P2,P3) = s1<->P2+P7<->s1+P8<->P7+s2<->P3+P8<->s2"
```

```
#F3 statistics (first five)
head(sim.fstats.sym$F3.equations)
```

```
[1] "F3(P1;P2,P3) = P7<->P1"
[2] "F3(P1;P2,P6) = P7<->P1+a*P7<->s1"
[3] "F3(P1;P2,P4) = P7<->P1"
[4] "F3(P1;P2,P5) = P7<->P1"
[5] "F3(P1;P3,P6) = P7<->P1+(1-a)*P8<->P7+(1-a)*P8<->s2"
[6] "F3(P1;P3,P4) = P7<->P1+P8<->P7"
```

```
#F4 statistics (first five)
head(sim.fstats.sym$F4.equations)
```

```
[1] "F4(P1,P2;P3,P6) = P7<->s1*a"
[2] "F4(P1,P2;P3,P4) = 0"
[3] "F4(P1,P2;P3,P5) = 0"
[4] "F4(P1,P2;P6,P4) = -P7<->s1*a"
[5] "F4(P1,P2;P6,P5) = -P7<->s1*a"
[6] "F4(P1,P2;P4,P5) = 0"
```

#### 7.2 Generating files for the *qpGraph* software with *graph.params2qpGraphFiles*

The *graph.params2qpGraphFiles* function allows generating the files required by the *qpGraph* software (Patterson *et al.* 2012) from a *graph.params* object that includes estimates of f-statistics (see section 5.1.2). If *f* is the prefix character specified with the *outfileprefix* argument of the function (by default *f=out*), these are:

- a file named "*f*.graph" that specifies the graph in *qpGraph* format
- a file named "*f*.fstats" with estimates of F-statistics (and their covariance) included in the input *graph.params* object
- a parameter file named "*f*.parqpGraph" to run *qpGraph* (this file may be edited by hand if other options are needed).

The *qpGraph* software (*v7365* and above to allow f-statistics estimates to be provided as input) may then be run on a terminal using the following options:

```
qpGraph -p f.parqpGraph -g f.graph -o out.ggg -d out.dot.input
```

The "*f*.fstats" f-statistics file must be in the same directory or its PATH should be explicitly specified by editing the "*f*.parqpGraph" parameter file. The following example runs *qpGraph* (providing appropriate install of the software) on the *sim.graph.params* object generated in 5.2 (see Figure 9 representing the fitted graph obtained with the *fit.graph* function):

```
graph.params2qpGraphFiles(sim.graph.params,outfileprefix = "sim.graph")
```

```
Fstats input file for qpGraph written in sim.graph.fstats
Graph input file for qpGraph written in sim.graph.graph
Parameter File for qpGraph with some default parameters written in sim.graph.parqpGraph
```

```
#running qpGraph (installed locally) outside R
system("qpGraph -p sim.graph.parqpGraph -g sim.graph.graph -o sim.graph.g -d sim.graph.dot")
#plotting the dot file generated by qpGraph with grViz (as done internally by poolfstat)
require(DiagrammeR)
grViz("sim.graph.dot")
```

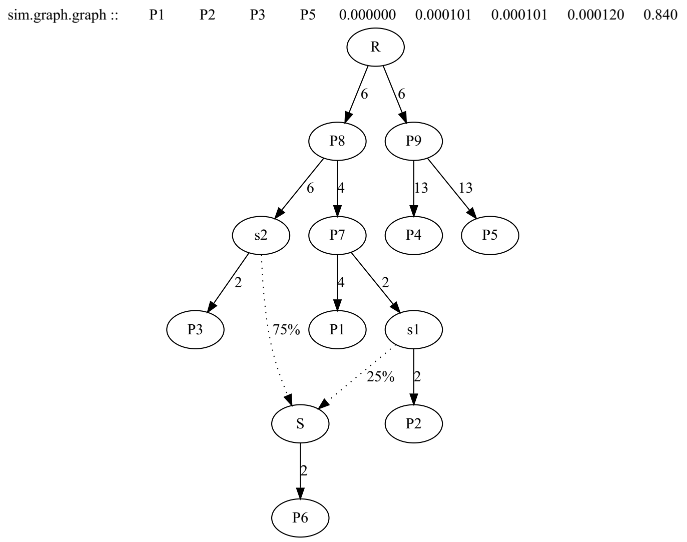

Figure 15: Fitting results obtained by qpGraph on the same data as the one used to generate Figure 9.

Comparison of Figures 15 and 9 shows that the same results are obtained with the two fitting methods (note that edge lengths are not scaled in drift units on the two figures).

### A Apprendix

#### A.1 Block-Jackknife estimation of standard errors

Standard-error of genome-wide estimates of $F_{ST}$ and other $f$-statistics can estimated using a block-jackknife sampling approach (see Patterson *et al.* 2012 and references therein). The algorithm implemented in *poolfstat* consists of dividing the genome into contiguous chunks of a pre-defined number of SNPs (specified by the argument *nsnp.per.bjack.block* of the *computeFST*, *compute.pairwiseFST* or *compute.fstats* functions, see sections 3.1.1, 3.2 and 4 respectively) and then removing each block in turn to quantify the variability of the estimates. If $n_b$ blocks are available and $\widehat{f_i}$ is the estimate of the statistics when removing all the SNPs belonging to block $i$, the standard error $\widehat{\sigma_f}$ of the estimator $\widehat{f}$ of the statistics of interest is computed as:

$$\widehat{\sigma_f} = \sqrt{\frac{n_b - 1}{n_b} \sum_{i=1}^{i=n_b} \left( \widehat{f_i} - \widehat{\mu_f} \right)^2}$$

where $\widehat{\mu_f} = \frac{1}{nb} \sum_{i=1}^{i=nb} \widehat{f_i}$ which may be slightly different from the estimator obtained with all the markers since the latter may include SNPs that are not eligible for block-jackknife sampling (e.g., those at the chromosome/scaffolds boundaries or those belonging to chromosome/scaffolds with less than *nsnp.per.bjack.block* SNPs). Finally, block-jackknife sampling may also be used to obtain estimates of the covariance between the estimates $\widehat{f^a}$ and $\widehat{f^b}$ as (using similar notations):

$$\widehat{\mathbb{C}\text{ov}} \left( \widehat{f^a}; \widehat{f^b} \right) = \frac{n_b - 1}{n_b} \sum_{i=1}^{i=n_b} \left( \widehat{f_i^a} - \widehat{\mu_{f^a}} \right) \left( \widehat{f_i^b} - \widehat{\mu_{f^b}} \right)$$
