## Supplementary Vignette V2 for "*f*-statistics estimation and admixture graph construction with Pool-Seq or allele count data using the R package *poolfstat*"

### Analysis of a D. suzukii Pool-Seq data set with the R package *poolfstat*

2021-05-27

#### Contents

**1 Data preparation and SNP filtering** **2**

**2 Estimation of global $F_{ST}$ and all $f-$ and D- statistics** **4**

**3 Overview of the within-population genetic diversity from estimates of heterozygosities** **6**

**4 Overview of the structuring of genetic diversity across populations from pairwise-population $F_{ST}$ estimates** **6**

**5 Insights from $f_3$-based admixture tests** **8**

**6 Exploring invasion scenarios with admixture graph construction** **12**

**References** **34**

This vignette details the analysis with the R package *poolfstat*[1] of the Pool-Seq data previously generated by Olazcuaga *et al.* (2020) for 14 population samples of the invasive species *Drosophila suzukii* (Figure 1 and Table 1). The objective of this analysis was to provide an illustration of the main *poolfstat* package functionalities on real data while providing insights into the history of both native populations and the recent invasion of this species into America (Fraimout *et al.* 2017). As illustrated in Figure 1, the pool samples represented 6 populations from the Asian native area, one from Hawaii (see Figure 1 legend) and 7 populations from the American continent. Out of the 14 population samples, all but CN-Bei originate from the same site as in Fraimout *et al.* (2017) who inferred the routes of invasion on a worldwide scale under an Approximate Bayesian Computation Random Forest (ABC-RF) approach using a data set consisting of 685 individuals belonging to 23 populations (from 15 to 44 individuals per sample) and genotyped at 25 autosomal microsatellite loci. As detailed in Table 1, for nine (CN-Lia, CN-Nin, CN-Shi, JP-Sap, BR-Pal, US-Col, US-Sdi, US-Sok and US-Wat) of the 14 population samples, all or most (30 out 50 of and 40 out 50 for CN-Lia and CN-Nin, respectively) of the individuals included in the sequenced pool were in common among the two studies. This thus makes the outcome of the Pool-Seq data analysis based on *poolfstat* directly comparable with the analysis performed by Fraimout *et al.* (2017).

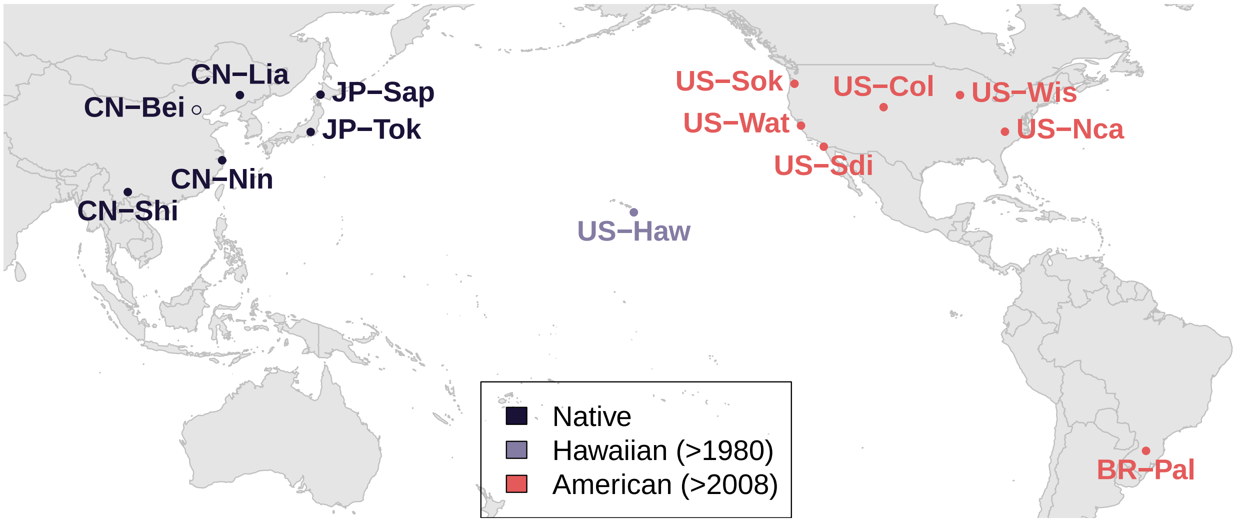

Figure 1: Origin of the 14 population samples of *D. suzukii*. Population names are colored according to their area of origin, i.e., whether they originate from the Asian native, Hawaii or continental America where the species was first observed in 2008. As done in previous studies, we distinguished the Hawaiian population from the American invasive area because the presence of *D. suzukii* was first described in 1980 in Hawaii. Hence, this Pacific Island population may be considered as intermediate between the Asian native and most recently invaded areas (approximately 300 generations later assuming 10 generations per year). The 13 population samples originating from the same site as in Fraimout et al. (2017) are indicated by solid points.

### 1 Data preparation and SNP filtering

The read alignment (*bam*) files from Olazcuaga *et al.* (2020) obtained after mapping Pool-Seq data for each of the 14 population samples onto the latest near-chromosome scale assembly of the *Drosophila suzukii* (Paris *et al.* 2020) were combined into an *mpileup* file using *SAMtools* 1.9 with options *-q 20 -Q20* (Li *et al.* 2009). Variant calling was then performed on the resulting file using VarScan mpileup2snp v2.3.4 (Koboldt *et al.* 2012) run with options *–min-coverage 10 –min-avg-qual 25 –min-var-freq 0.005 –p-value 0.5 –output-vcf 1* ensuring very loose criteria to identify SNPs further SNP filtering being delayed to the parsing of the vcf file. In particular we set the p-value threshold to 0.5 leading to the identification

[1]All the analysis were carried on a laptop computer equipped with an intel Xeon Quad-Core 3.0 Ghz processor and 32 Gb RAM

| Sample name | Status | Sample Size | Sampling site | Sampling date | SRA ID |
|---|---|---|---|---|---|
| CN-Bei (-) | Native | 50 (0) | Beijing, China | 2014 | SRR10260017 |
| CN-Lia (CN-Lia) | Native | 50 (50) | Liaoyuan, China | 2014 | SRR10260016 |
| CN-Nin (CN-Nin) | Native | 50 (30) | Ningbo, China | 2014/2016 | SRR10260027 |
| CN-Shi (CN-Shi) | Native | 50 (40) | Shiping county, China | 2014/2016 | SRR10260024 |
| JP-Sap (JP-Sap) | Native | 50 (50) | Sapporo, Japan | 2014 | SRR10260023 |
| JP-Tok (JP-Tok) | Native | 50 (0) | Tokyo, Japan | 2016 | SRR10260022 |
| BR-Pal (BR-PA) | Invasive (AM) | 50 (50) | Porto Alegre, Brazil | 2014 | SRR10260033 |
| US-Col (US-Col) | Invasive (AM) | 50 (50) | Fort Collins, USA | 2015 | SRR10260032 |
| US-Haw (US-Haw) | Invasive (PA) | 50 (0) | Hawaii (Hilo), USA | 2016 | SRR10260031 |
| US-Nca (US-NC) | Invasive (AM) | 100 (0) | Raleigh, USA | 2016 | SRR10260030 |
| US-Sdi (US-SD) | Invasive (AM) | 50 (50) | San-Diego, USA | 2014 | SRR10260029 |
| US-Sok (US-Sok) | Invasive (AM) | 75 (75) | Dayton, USA | 2014 | SRR10260028 |
| US-Wat (US-Wat) | Invasive (AM) | 50 (50) | Watsonville, USA | 2014 | SRR10260026 |
| US-Wis (US-Wis) | Invasive (AM) | 75 (0) | Barneveld, USA | 2016 | SRR10260025 |

Table 1: Pool sample description (adapted from Table S2 in Olazcuaga et al., 2020). Population names and the number of individuals included in the pools in common with Fraimout et al. (2017) are given in parenthesis in the first and third columns, respectively. Only the CN-Bei population sample is absent from the microsatellite data set of Fraimout et al. (2017). As for the population origins (column status), we distinguished i) the Asian native area (6 population samples); ii) the Pacific islands (PA) invasive area (the Hawaiian sample); and iii) the contiental American invasive area (7 population samples).

of SNP with only one read supporting the non-reference allele in a single pool whatever the coverage[2]. Positions mapping to non-autosomal contigs (Paris *et al.* 2020) were subsequently discarded from the vcf file to obtain the file named *dsu.auto.vcf.gz* which can be downloaded from the public Zenodo repository (http://doi.org/10.5281/zenodo.4709080) together with a file named *dsu.auto.info* that contains names and haploid sample sizes of each pool. As shown below, we then used the *vcf2pooldata* function to parse this vcf file relying on default options except for the overall MAF threshold (computed from read counts) that was set to 5% (*min.maf* argument) and the minimal read coverage for each pool that was set to 50 (*min.cov.per.pool* argument). The resulting *pooldata* object was further filtered with the *pooldata.subset* function to discard i) all positions with a coverage higher than the 99% quantile coverage in at least one pool (*cov.qthres.per.pool* argument); and ii) discard all SNPs with MAF<5% over all the populations from the native area to ensure that mostly ancestral SNPs were included in the data as implicitly assumed by f-statistics based approaches (see the main text of the manuscript and associated references).

```r
#loading information file containing:
#            i)  pool names (in the same order as in the vcf file)
#            ii) haploid pool size
dsu.info=read.table("dsu.auto.info",stringsAsFactors = F)
#parsing of the vcf file for autosomal contigs
dsu.dat=vcf2pooldata(vcf.file="dsu.auto.vcf.gz",
                     poolsizes =dsu.info[,2],poolnames = dsu.info[,1],
                     min.maf = 0.05,min.cov.per.pool = 50,remove.indels = T,
                     nlines.per.readblock = 1e7)
```

```
Reading Header lines
Parsing allele counts
VarScan like format detected for allele count data: the AD field contains allele depth
for the alternate allele and RD field for the reference allele
(N.B., positions with more than one alternate allele will be ignored)
1e+07  lines processed in 0 h  5 m  54 s : 773642 SNPs found
```

[2]The Fisher exact test use for variant calling in *VarScan* gives $p = \frac{C_r^c C_0^c}{C_r^{2c}} = \frac{C_r^c}{C_r^{2c}}$ where $c$ is the coverage and $r$ is the number of reads supporting the non-reference allele. Hence, $p = 0.5$ if $r = 1$

```
2e+07  lines processed in 0 h  10 m  50 s : 1546200 SNPs found
3e+07  lines processed in 0 h  15 m  27 s : 2136045 SNPs found
30922056  lines processed in 0 h  15 m  50 s : 2161633 SNPs found
Data consists of 2161633 SNPs for 14 Pools
```

```
#Further filtering of SNPs keeping only those with MAF>0.05 on the native populations
#and coverage not exceeding the 99% pool-specific quantile of coverage in a least one pop
pop.nat.names=c("CN-Bei","CN-Lia","CN-Nin","CN-Shi","JP-Tok","JP-Sap")
pop.nat.idx=which(dsu.dat@poolnames %in% pop.nat.names)
tmp.f.nat=rowSums(dsu.dat@refallele.readcount[,pop.nat.idx])/
          rowSums(dsu.dat@readcoverage[,pop.nat.idx])
maf.native=0.5-abs(0.5-tmp.f.nat)
dsu.dat=pooldata.subset(dsu.dat,snp.index = which(maf.native>0.05),
                        cov.qthres.per.pool = c(0,0.99))
```

```
Data consists of 1588569 SNPs for 14 Pools
```

The final data sets thus consists of 1,588,569 SNPs with sequencing coverage statistics among the 14 different pools detailed in Table 2.

| | Mean Coverage | Median Coverage | Coverage Range |
|---|---|---|---|
| BR-Pal | 76.72 | 76 | 50-135 |
| CN-Bei | 95.93 | 95 | 50-160 |
| CN-Lia | 69.68 | 69 | 50-127 |
| CN-Nin | 66.7 | 65 | 50-122 |
| CN-Shi | 68.41 | 67 | 50-138 |
| JP-Sap | 84.53 | 84 | 50-174 |
| JP-Tok | 68.05 | 67 | 50-116 |
| US-Col | 79.35 | 79 | 50-144 |
| US-Haw | 95.41 | 95 | 50-160 |
| US-Nca | 74.24 | 73 | 50-121 |
| US-Sdi | 89.64 | 89 | 50-143 |
| US-Sok | 65.29 | 64 | 50-130 |
| US-Wat | 73.34 | 72 | 50-126 |
| US-Wis | 76.7 | 76 | 50-143 |

Table 2: Summary of the sequencing coverage of the 14 Pool-Seq samples over the 1,588,569 selected SNPs

#### 2 Estimation of global $F_{ST}$ and all $f-$ and D- statistics

To characterize the overall structuring of genetic diversity, we estimated the $F_{ST}$ over all the 14 populations using the estimator previously described by Hivert *et al.* (2018) and implemented in the *computeFST* function. We estimated the standard error of the resulting estimate using a block-jackknife approach specifying blocks of 10,000 SNPs. As shown below (information printed by the *computeFST* function), this resulted in 145 blocks of 700 kb on average. Note that the number of SNPs eligible for black-jackknife (and thus used for estimation) were reduced to 1,450,000 SNPs (i.e., 90% out of the 1,588,569) due to the removal of SNPs mapping to contigs represented by less than 10,000 SNPs (i.e., small contigs which may actually be less reliable as corresponding to the most fragmented part of the assembly) and to a lesser extent SNPs mapping to contig boundaries.

```
dsu.global.fst=computeFST(dsu.dat,nsnp.per.bjack.block = 10000)
```

```
Starting Block-Jackknife sampling
145 Jackknife blocks identified with 1450000 SNPs (out of 1588569 ).
 SNPs map to 15 different chrom/scaffolds
Average (min-max) Block Sizes: 0.698 ( 0.414 - 2.306 ) Mb
```

```r
#Global Fst estimate
dsu.global.fst$mean.fst
```

```
[1] 0.07107105
```

```r
#95% CI of the Fst estimate (based on the block-jackknife s.e.)
dsu.global.fst$mean.fst+c(-1.96,1.96)*dsu.global.fst$se.fst
```

```
[1] 0.06897769 0.07316441
```

We next computed all the $f-$ and $D-$ statistics together with their block-jackknife standard error (with blocks of 15,000 SNPs as for the global $F_{ST}$) using the *compute.fstats* function. These included:

- $f_2$ for all the 91 pairs of populations together with their scaled version that correspond to the pairwise $F_{ST}$ estimator based on the IIS probabilities (i.e., similar to the one implemented by the *computeFST* or *compute.pairwiseFST* functions when specifying *method="Idendity"*)
- $f_3$ (and their scaled version $f_3^\star$) for all the 1,092 possible triplets of populations
- $f_4$ (and their scaled version $D$) for all the 3,003 possible quadruplets of populations

```r
dsu.fstats=compute.fstats(dsu.dat,computeDstat = TRUE,
                          return.F4.blockjackknife.samples = TRUE,
                          nsnp.per.bjack.block = 10000)
```

```
Estimating Q1
Estimating Q2
Estimating within-population heterozygosities
Estimating F2
Estimating F3
Estimating F4
 Computing Dstat
Starting Block-Jackknife sampling
145 Jackknife blocks identified with 1450000 SNPs (out of 1588569 ).
 SNPs map to 15 different chrom/scaffolds
Average (min-max) Block Sizes: 0.698 ( 0.414 - 2.306 ) Mb
   computing Q1 averages per blocks
   computing Q2 averages per blocks
   computing F2 averages per blocks
Starting computation of estimators s.e.
   within-pop heterozygosity s.e. estimation done
   F2 s.e. estimation done
   F3 and F3* s.e. estimation done
   estimating F4 and Dstat s.e. (may be long since require denominator averages per blocks)
   F4 and D s.e. estimation done

Overall Analysis Time: 0 h 1 m 37 s
```

Note that the *return.F4.blockjackknife.samples* argument was set to *TRUE* to allow for estimation of some admixture rates using $f_4$ ratios (see below). In addition, as shown below most of the computation time is spent on estimating D-statistics which are actually not always needed (e.g., treeness tests may be based on $f_4$) for most future analysis steps. In addition, due to internal optimization of the code block-jackknife estimation of the standard errors is performed at very limited computational costs (except for D-statistics):

```r
#computing all fstats without D-statistics
#with verbose=FALSE and hence use of sys.time to estimate computation time
tb=Sys.time()
tmp<-compute.fstats(dsu.dat,computeDstat = FALSE,
                    return.F4.blockjackknife.samples = TRUE,
```

```
                   nsnp.per.bjack.block = 10000,verbose=FALSE)
cat("Analysis (without D-statistics estimation) took",
    round(difftime(Sys.time(),tb,units="secs"),1),"s")
```

```
Analysis (without D-statistics estimation) took 12.6 s
```

```
#computing all fstats without D-statistics
# and without blockjacknife estimates of s.e. (little interest)
tb=Sys.time()
tmp<-compute.fstats(dsu.dat,computeDstat = FALSE,verbose=FALSE)
cat("Analysis (without D-statistics and block-jackknife s.e. estimation)
    took",round(difftime(Sys.time(),tb,units="secs"),1),"s")
```

```
Analysis (without D-statistics and block-jackknife s.e. estimation)
    took 6.3 s
```

### 3 Overview of the within-population genetic diversity from estimates of heterozygosities

Estimates of within-population heterozygosities (as $1-\hat{Q}_1$) provide a rough assessment of the genetic diversity of the different population. It should here be recalled that the SNP ascertainment process (see section 1) is expected to favor SNP polymorphic in the native area. The estimates and their 95% CI can be plotted as follows:

```
plot(dsu.fstats@heterozygosities$`bjack mean`,ylab="Heterozygosity",xaxt="n",xlab="",
     pch="",las=3,ylim=c(0.21,0.29))
axis(1,1:dsu.dat@npools,rownames(dsu.fstats@heterozygosities),las=3)
#plotting CI from bloc jackknife estimate of s.e.
ci=cbind(dsu.fstats@heterozygosities$`bjack mean` - 1.96*dsu.fstats@heterozygosities$`bjack s.e.`,
         dsu.fstats@heterozygosities$`bjack mean` + 1.96*dsu.fstats@heterozygosities$`bjack s.e.`)
tmp<-sapply(1:dsu.dat@npools,f<-function(z){
       abline(v=z,lty=3,col="grey")
       arrows(z,ci[z,1],z,ci[z,2],angle = 90,code=3,length=0.05,lwd=1.5,
              col=tmp.col[rownames(dsu.fstats@heterozygosities)[z]])
 })
legend("bottomleft",c("Native","Hawaiian (>1980)","American (>2008)"),
       fill=c("#1A1237","#847CA3","#E45A5A"))
```

As shown in Figure 2, US-Haw was clearly depleted in diversity as previously observed based on microsatellite data (Fraimout *et al.* 2017) which is also consistent with its island origin. The populations from the native area tended to display more diversity than those of the American invasive area with the noticeable exception of CN-Shi (that displayed significantly lower heterozygosity than the other populations from the native area) and US-Wat (that displayed the highest level of diversity among the populations from the invasive American area, similar to those of the native area). Note that filtering of SNPs during data preparation (see 1) may have resulted in upwardly biasing the relative amount of diversity in the native area.

### 4 Overview of the structuring of genetic diversity across populations from pairwise-population $F_{ST}$ estimates

Figure 3 plots estimates of $F_{ST}$ with their 95% CI for all the populations using the plot function applied directly onto the *dsu.fstat fstats* object computed above.

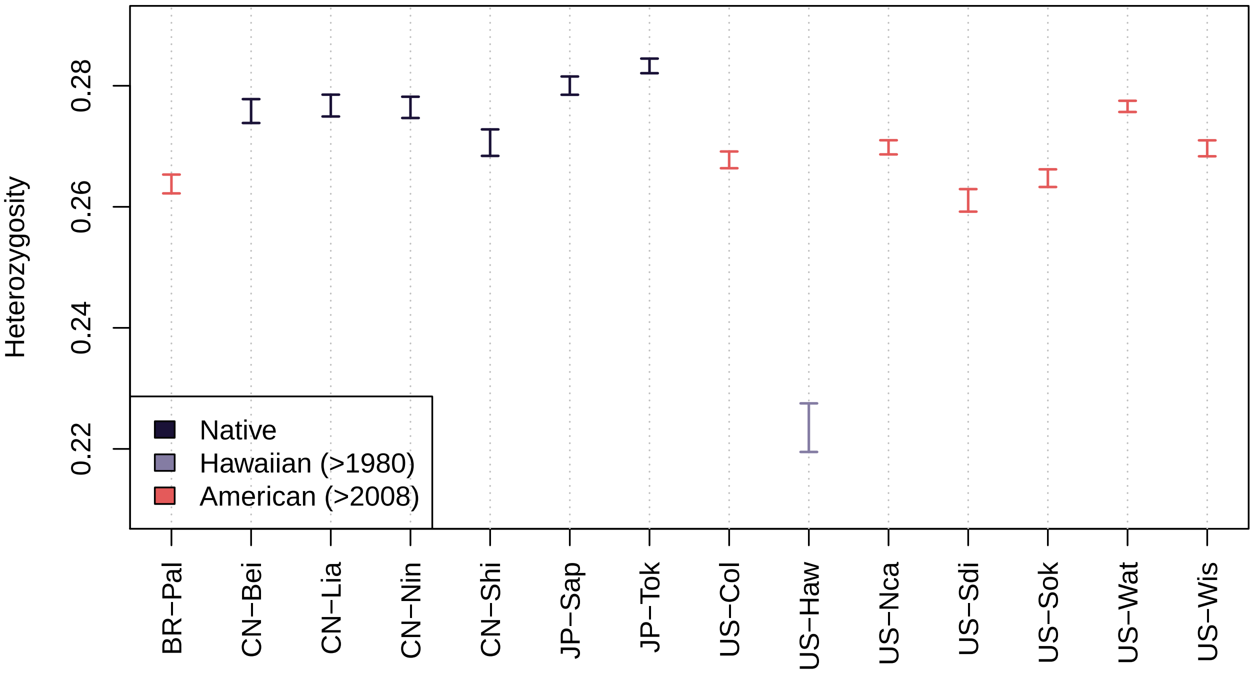

Figure 2: Estimates of within-population heterozygosities with their 95% CI from the ascertained SNPs (MAF>0.05 in the native area).

```
layout(matrix(1:2,1,2))
plot(dsu.fstats,stat.name="Fst",main="Fst<=5%",cex.main=1.5,value.range=c(0,0.05))
abline(v=0.01,lty=2,col="blue")
#plot(dsu.fstats,stat.name="Fst",main="5%<Fst<10%",cex.main=1.5,value.range=c(0.05,0.1))
plot(dsu.fstats,stat.name="Fst",main="Fst>5%",cex.main=1.5,value.range=c(0.05,NA))
```

An alternative (and complementary) view, based on the *compute.pairwiseFST* function (estimating by default the $F_{ST}$ estimator derived under an analysis-of-variance framework) and representing the matrix of pairwise $F_{ST}$ stored in the resulting *pairwisefst* object as an heatmap (using the *heatmap* function), is provided in Figure 4.

```
dsu.pairwise.fst=compute.pairwiseFST(dsu.dat)
```

```
Computation of the 91 pairwise Fst

Overall Analysis Time: 0 h 0 m 38 s
```

```
heatmap(dsu.pairwise.fst)
```

Note that the two different estimates of pairwise-population $F_{ST}$ remain very similar (Figure 5).

```
plot(dsu.pairwise.fst@values$`Fst Estimate`,dsu.fstats@fst.values$`bjack mean`,
     xlab="compute.pairwiseFST  (method=Anova)",ylab="compute.fstats")
abline(a=0,b=1,lty=2)
```

As expected from their recent history and as already observed with other analysis (Olazcuaga *et al.* 2020), populations tended to cluster according to their area of origin (Figure 4) with some closely geographically related populations showing very low level of differentiation. For instance, in continental America, US-Nca, US-Col and US-Nca all displayed pairwise $F_{ST} < 0.01$ with each others (Figure 3) and in the Asian native area, the three populations CN-Bei, CN-Nin and CN-Lia originating for North-West China were all found

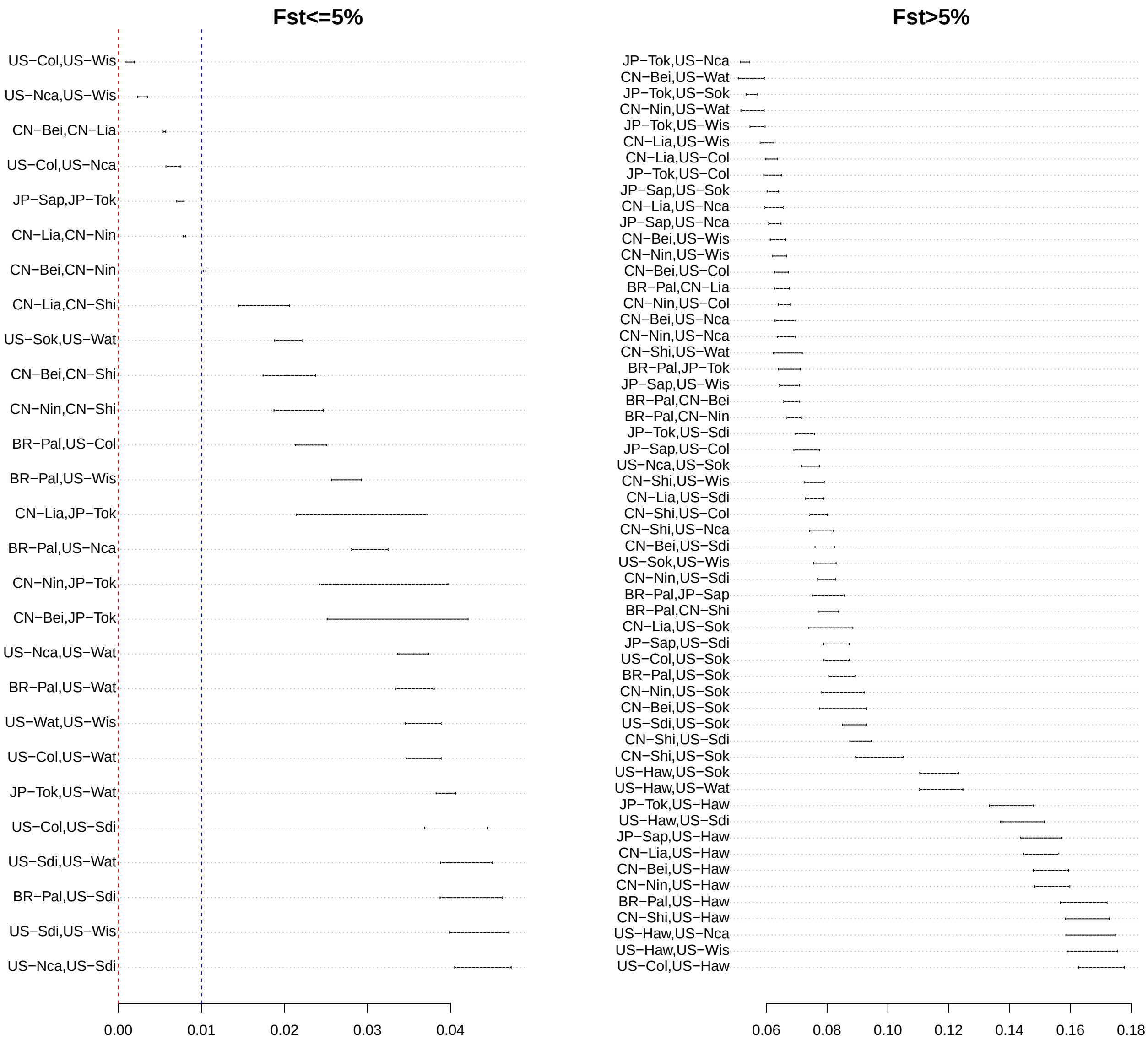

Figure 3: Estimates of pairwise $F_{ST}$ for the population pairs of the *D. suzukii* data set. For ease of reading, plots have been arranged according to the level of differentiation into small (<0.05) and moderate to high level of differentiation (>0.05). In the leftmost plot, the blue dotted line represent the 0.01 $F_{ST}$ threshold (i.e., very weak levels of differentiation)

closely related. Conversely, the Hawaiian sample (US-Haw) was found the most highly differentiated with all the other populations, the pairwise $F_{ST}$ ranging from 11.7% (with US-Sok) to 17.0% (with US-Col). Note that this explains why US-Haw behave as an outgroup in the heatmap of Figure 4, thereby illustrating that such representation may be misleading if interpreted in terms of demographic history (Hawaii obviously not being the area of origin of the species).

### 5 Insights from $f_3$-based admixture tests

Six of the 14 populations showed at least one significantly negative $f_3$ or $f_3^{\star}$ statistics (both statistics being almost exactly similar) at the 95% threshold of significance (i.e., with an associated Z-score $< -1.65$). Table 3 summarizes for each of these 6 target populations, the number of significantly negative $f_3$ (and $f_3^{\star}$) together with the triplet displaying the lowest Z-score (out of the $\frac{13\times12}{2} = 78$ tests per population), as obtained with

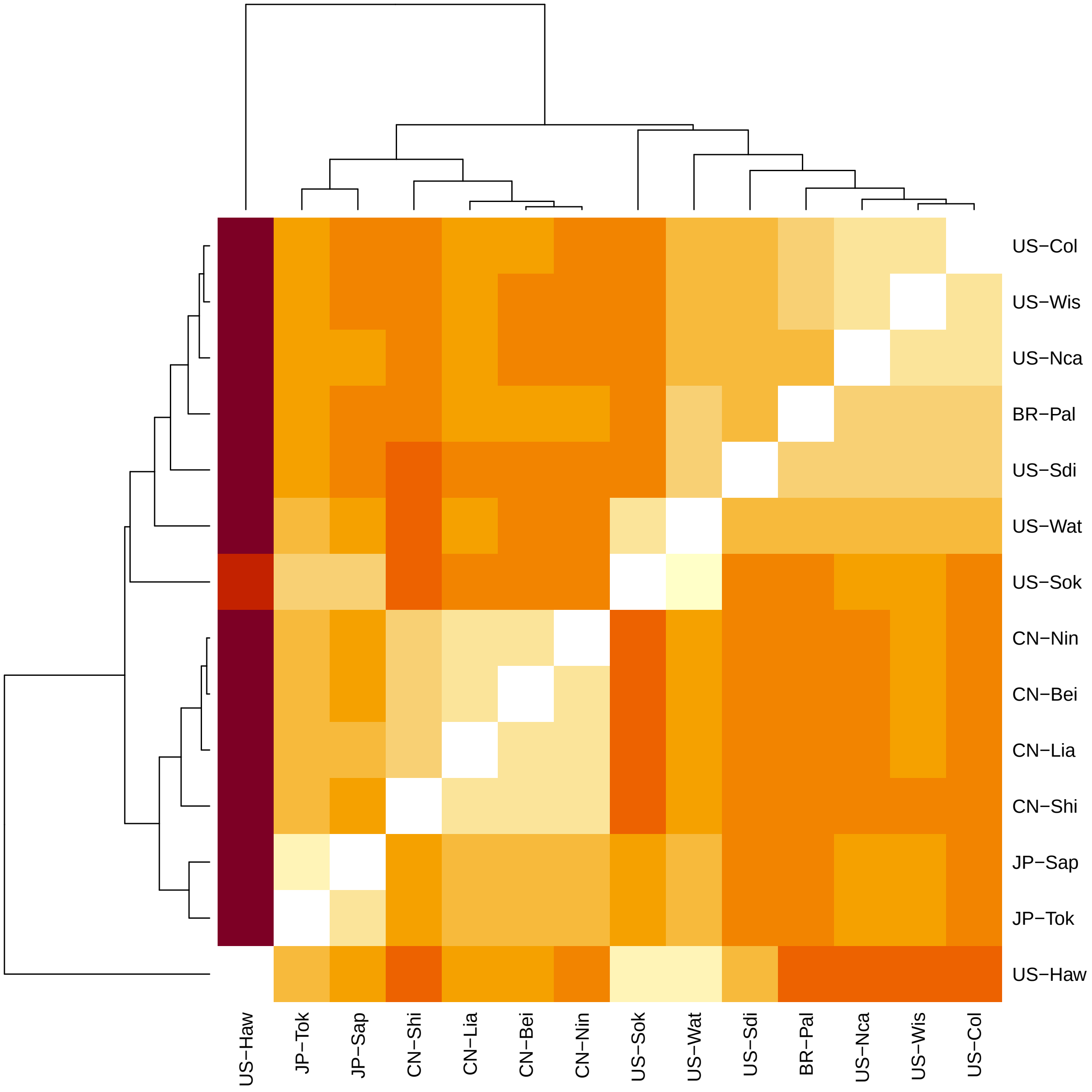

Figure 4: Heatmap representation of the matrix of pairwise-population $F_{ST}$ of 14 *D. suzukii* populations.

the following script:

```
f3.signif=dsu.fstats@f3.values$`Z-score`< -1.65
f3s.signif=dsu.fstats@f3star.values$`Z-score`< -1.65
nf3.signif.per.pop=table(dsu.fstats@comparisons$F3[f3.signif,1])
nf3s.signif.per.pop=table(dsu.fstats@comparisons$F3star[f3s.signif,1])
##custom function to retrieve the triplet with lowest Z-score for a given target pop.
tmp.f<-function(x,stat="F3"){
  if(stat=="F3"){
    x.f3=dsu.fstats@f3.values[dsu.fstats@comparisons$F3[,1]==x,]
  }else{
    x.f3=dsu.fstats@f3star.values[dsu.fstats@comparisons$F3star[,1]==x,]
```

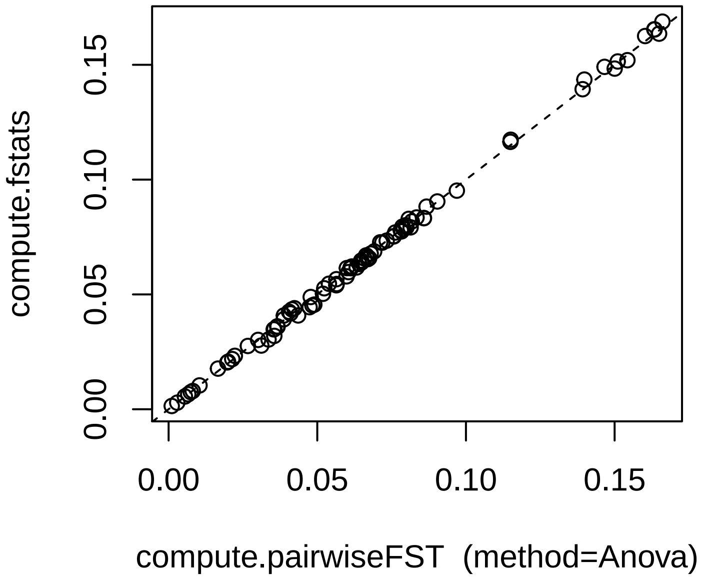

Figure 5: Comparison of the pairwise-population estimates $F_{ST}$ computed by the compute.pairwiseFst (method="Anova") and compute.fstats (equivalent to method="Identity" in compute.pairwiseFst). Note that for the latter, the estimates were taken as the block-jacknife means.

```
  }
  idx=which.min(x.f3$`Z-score`)
  return(paste0(rownames(x.f3)[idx]," (Z=",
                round(x.f3$`Z-score`[idx],2),")"))
}
###
pop.signif=unique(c(names(nf3.signif.per.pop),names(nf3s.signif.per.pop)))
f3.mostsignif=sapply(pop.signif,tmp.f)
f3s.mostsignif=sapply(pop.signif,tmp.f,stat="F3s")
```

| | Pop. Origin | #f3<0 | #f3*<0 | most. sign. triplet (f3) | Most sign. triplet (f3*) |
|---|---|---|---|---|---|
| CN-Lia | Native | 1 | 1 | CN-Lia;CN-Shi,JP-Sap (Z=-1.66) | CN-Lia;CN-Shi,JP-Sap (Z=-1.65) |
| JP-Tok | Native | 11 | 11 | JP-Tok;CN-Nin,JP-Sap (Z=-7.11) | JP-Tok;CN-Nin,JP-Sap (Z=-7.09) |
| US-Col | Invasive (AM) | 2 | 2 | US-Col;BR-Pal,US-Wis (Z=-3.31) | US-Col;BR-Pal,US-Wis (Z=-3.32) |
| US-Nca | Invasive (AM) | 6 | 6 | US-Nca;JP-Sap,US-Col (Z=-3.89) | US-Nca;JP-Sap,US-Col (Z=-3.9) |
| US-Wat | Invasive (AM) | 13 | 13 | US-Wat;US-Sdi,US-Sok (Z=-23.64) | US-Wat;US-Sdi,US-Sok (Z=-23.81) |
| US-Wis | Invasive (AM) | 4 | 4 | US-Wis;JP-Sap,US-Col (Z=-5.02) | US-Wis;JP-Sap,US-Col (Z=-5.05) |

Table 3: Summary of the f3 and f3* based tests for all the populations displaying at least one significant test at the 95% threshold (Z< -1.65)

Figure 6 plots the lowest $f_3$ (including the significantly negative ones highlighted in red) for the six populations displaying at least one significant three-population test.

```
layout(matrix(1:6,3,2))
for(i in pop.signif){
plot(dsu.fstats,stat.name="F3",main=i,cex.main=1.5,value.range=c(NA,1e-3),
     pop.f3.target=i,ci.perc=90)
  #ci.perc was here set to 90% to obtain the same significant
  #tests (highlighted in red) as with Z<-1.65 (since the test is one-sided)
}
```

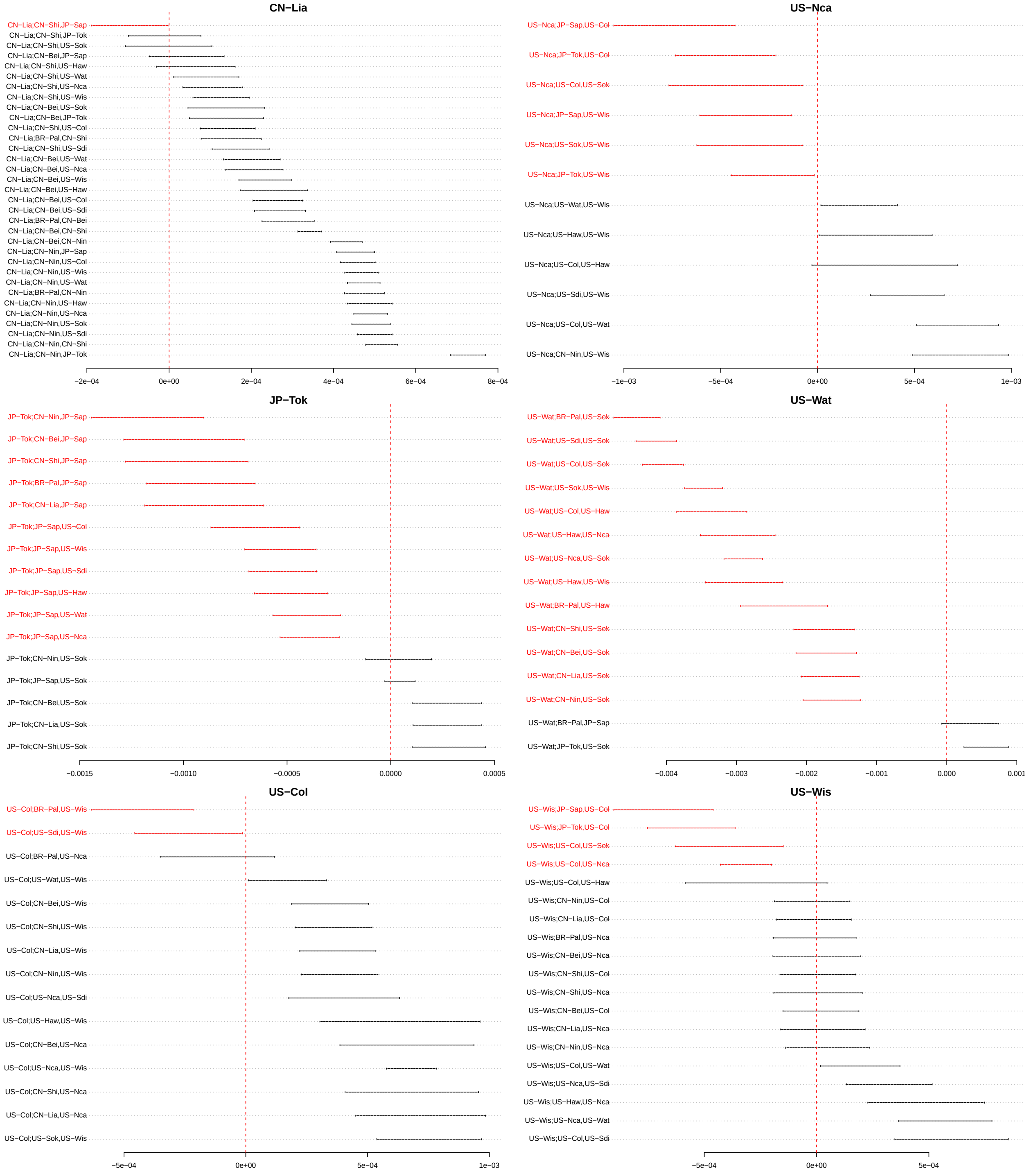

Figure 6: Estimates of the lowest $f_3$ statistics with their 95% Confidence Intervals for the 6 populations displaying at least one significant three-population test at the 95% threshold (shown in red).

In the native area, JP-Tok showed clear evidence of admixture with 11 significant tests that all involved JP-Sap as a source proxy. The three lowest $f_3$ were obtained with the three Chinese populations (CN-Nin, CN-Bei and CN-Shi in increasing order of $f_3$). Assuming an admixture-graph like history, this suggested that the two populations branching the closest to the two sources of JP-Tok were JP-Sap and CN-Nin. The remaining Chinese population, CN-Lia showed some little evidence of admixture with only one test barely significant at the 95% threshold for the triplet involving CN-Shi and JP-Sap as source proxies.

Out of the 7 populations from the American continent (excluding Hawaii), 4 showed evidence of admixture, namely US-Col, US-Wis, US Nca and US-Wat. The strongest evidence (13 significant tests) was found for US-Wat which has up to now been considered as the closest to the first invader of continental America based on historical records (Fraimout *et al.* 2017). Moreover, the three signals with the lowest Z-score all involved two source populations originating from the (continental) American invasive area. As their underlying $f_3$ CI did not overlap with the other tests, these three pairs of populations may be considered as the closest (among the sampled populations) to the original US-Wat source populations. Hence, the three most significant signals all involved US-Sok (Northern US) as a source proxy while the other source proxies were, in order of increasing Z-score (i.e., decreasing evidence as Z-score are here negative), BR-Pal, US-Sdi and US-Col respectively. Challenging the initial view, this thus suggests that US-Wat derived from an admixture between two populations already established in the continental America, one Northern (here represented by US-Sok) and the other Southern (here represented by BR-Pal, US-Sdi and to a lesser extent US-Col that may actually derived from US-Sdi considering historical records) as will be investigated below via admixture graph construction. For this Southern "ghost" population, we may further speculate that BR-Pal, although geographically distantly related, could be the closest proxy as a result of a rapid spread of *D. suzukii* in Southern America although the absence of additional samples in South America makes this hypothesis difficult to test. The three other American populations, US-Col, US-Wis and US-Nca, with at least one significantly negative $f_3$ displayed only a moderate number of significant tests (compared to other populations) and were all very closely related (section 4). For each of these, the significantly negative $f_3$ had highly overlapping CI and always involved one or the two other populations[3]. This thus suggests complex patterns of recurrent admixture events among these three populations (consistent with their low level of differentiation and close geographic origins).

### 6 Exploring invasion scenarios with admixture graph construction

To provide further insights into the relationships of the surveyed populations and the probable scenarios of invasion of the species in American and European area, we relied on admixture graph construction. The purpose was no here to build a comprehensive admixture graph, which may be elusive given the close relationships of the populations and the pervasiveness of recent admixture events among the different populations, but rather to identify key regional event that happened at early time of invasion. Importantly, we pay critical attention to the support of the different graphs to assess the validity of the proposed scenarios by making extensive use of the *compare.fitted.fstats* function.

#### 6.1 Scaffold trees built (naively) from scratch are unreliable

We first evaluated a naive approach consisting of building a scaffold tree of presumably unadmixed populations using the *find.tree.popset* and *rooted.njtree.builder* functions. Although two candidate sets of 5 populations could be identified, the bifurcating trees obtained from them provided a poor fit to the data, making them obviously unreliable for further admixture graph construction. More precisely, we inferred for each of these two sets the best rooted bifurcating tree using the *rooted.njtree.builder* function and evaluate its fit with the *compare.fitted.fstats* function (note that the fitted f-statistics are invariant to root positioning for such bifurcating trees). In addition, we also used the branch length estimates obtained with the *fit.graph* function (instead of those relying on neighbor-joining algorithm used by *rooted.njtree.builder*) of the best rooted tree but this lead to similar conclusions.

```
#Identification of candidate (unadmixed) scaffold populations from scratch
scaf.pop=find.tree.popset(dsu.fstats)
```

```
Number of sets: 24 of Npops= 4 each
```

[3]More precisely, the four significant tests with US-Wis as a target all involved US-Col as a source proxy and one also involved US-Nca (although with the lowest $f_3$); out of the six significant tests with US-Nca as a target, three involved US-Col as a source proxy and the three other involved US-Wis; and the two significant tests with US-Col as a proxy involved US-Wis as a source proxy

```
Number of sets: 2 of Npops= 5 each
```

```r
scaf.pop$pop.sets
```

```
        [,1]     [,2]     [,3]     [,4]     [,5]
PopSet1 "BR-Pal" "CN-Bei" "CN-Shi" "JP-Sap" "US-Sok"
PopSet2 "CN-Bei" "CN-Shi" "JP-Sap" "US-Sdi" "US-Sok"
```

```r
#construction of scaffold trees for each set
scaf.pop.trees=list()
for(i in 1:scaf.pop$n.sets){
  scaf.pop.trees[[i]]=list()
  tmp.njtree<-rooted.njtree.builder(pop.sel = scaf.pop$pop.sets[i,],
                                    fstats = dsu.fstats,verbose=FALSE)
  scaf.pop.trees[[i]][["njtree"]]<-tmp.njtree$best.rooted.tree
  tmp.graph=generate.graph.params(tmp.njtree$best.rooted.tree@graph,dsu.fstats,verbose=F)
  scaf.pop.trees[[i]][["fitgraph"]]<-fit.graph(tmp.graph,verbose=FALSE)
  rm(tmp.njtree,tmp.graph)
}
#Evaluation of scaffold trees using compare.fitted.fstats
for(i in 1:scaf.pop$n.sets){
  cat("\nScaffold set:",i,"\n")
  cat("\tRooted NJ tree\n")
  tmp.comp<-compare.fitted.fstats(dsu.fstats,
                                  fitted.graph = scaf.pop.trees[[i]][["njtree"]],
                                  n.worst.stats = 1)
  cat("\tRooted fit graph tree\n")
  tmp.comp<-compare.fitted.fstats(dsu.fstats,
                                  fitted.graph = scaf.pop.trees[[i]][["fitgraph"]],
                                  n.worst.stats = 1)
}
```

```
Scaffold set: 1
    Rooted NJ tree
1 Worst fit for:
                              Estimated        Fitted Z-score
BR-Pal,CN-Shi;JP-Sap,US-Sok -0.005313611 -1.734723e-18 31.46101
    Rooted fit graph tree
1 Worst fit for:
                              Estimated Fitted Z-score
BR-Pal,CN-Shi;JP-Sap,US-Sok -0.005313611      0 31.46101

Scaffold set: 2
    Rooted NJ tree
1 Worst fit for:
                             Estimated      Fitted  Z-score
CN-Shi,US-Sdi;JP-Sap,US-Sok 0.005093941 0.002936964 -11.15586
    Rooted fit graph tree
1 Worst fit for:
                            Estimated Fitted Z-score
CN-Shi,JP-Sap;US-Sdi,US-Sok 0.00433532      0 -6.78807
```

Hence, the bifurcating scaffold tree inferred for the first and second candidate sets of candidate scaffold populations lead to f-statistics outlying from the raw estimated f-statistics, by up to 31 and 6.8 standard errors (as measured by the printed Z-score) respectively when considering neighbor-joining estimates of branch lengths.

It should be noticed that for both sets, the resulting NJ tree lead to quadruplets that violate the treeness test, i.e., that resulted in a quadruplet configurations that disagree with the one that was retained by the *find.tree.popset* function. For instance, in the first set of candidate populations, the obtained NJ tree topology resulted in the BR-Pal,CN-Shi;JP-Sap,US-Sok quadruplet which did not pass the $f_4$-based treeness tests as the quadruplet configuration passing the treeness test for these four populations is BR-Pal,JP-Sap;CN-Shi,US-Sok:

```
tmp.sel=apply(dsu.fstats@comparisons$F4,1,
              ff<-function(x){sum(x %in% c("BR-Pal","CN-Shi","JP-Sap","US-Sok"))==4})
dsu.fstats@f4.values[tmp.sel,]
```

```
                                Estimate    bjack mean   bjack s.e.    Z-score
BR-Pal,CN-Shi;JP-Sap,US-Sok -0.0051982044 -0.0053136113 0.0001688951 -31.461012
BR-Pal,JP-Sap;CN-Shi,US-Sok  0.0005166386  0.0002464223 0.0009666008   0.254937
BR-Pal,US-Sok;CN-Shi,JP-Sap  0.0057148430  0.0055600336 0.0008822927   6.301802
```

#### 6.2 Investigating the relationships among the populations from the native area (and the Hawaiian population)

We thus adopted an alternative and more historically informed strategy to build the starting scaffold graph, relying on the known Asian origin of the species (which may safely be assumed to be correct). We first seek to describe the historical relationships among the 6 populations from the native area also including the Hawaiian population that was first observed in 1980, i.e., more than 30 years (ca. 300 generations for *D. suzukii*) earlier than the other populations of invasive areas (Fraimout *et al.* 2017).

##### 6.2.1 Building a scaffold tree of native and the Hawaiian populations

We thus used the same approach as in the above section 6.1 to build a scaffold tree but focusing only on the six native and the Hawaiian populations:

```
#Identification of candidate (unadmixed) scaffold populations
pop.am.continent=c("BR-Pal","US-Col","US-Nca","US-Sdi","US-Sok","US-Wat","US-Wis")
scaf.pop.nat=find.tree.popset(dsu.fstats,excluded.pops = pop.am.continent)
```

```
Number of sets: 2 of Npops= 4 each
```

```
scaf.pop.nat$pop.sets
```

```
        P1       P2       P3       P4
PopSet1 "CN-Bei" "CN-Nin" "CN-Shi" "US-Haw"
PopSet2 "CN-Bei" "CN-Shi" "JP-Sap" "US-Haw"
```

```
#f4 associated Z-score range
#(here the range min and max are the same since only four populations in the sets)
scaf.pop.nat$Z_f4.range
```

```
        Min. |Zscore| Max. |Zscore|
PopSet1      1.086422      1.086422
PopSet2      1.060292      1.060292
```

Here two sets of 4 candidate populations could then be proposed. We chose to focus on the PopSet 2 (CN-Bei,CN-Shi ; JP-Sap,US-Haw) because it displayed the smallest $f_4$ associated Z-score (|Z|=1.06) and CN-Bei was more distantly related to CN-Shi than CN-Nin from both a geographical (Figure 1) and genetical point of view (Figure 3). As this selected set only contained 4 populations, the (best) unrooted topology was directly available from the quadruplet passing the $f_4$ treeness test, i.e. ((CN-Bei,CN-Shi),(JP-Sap,US-Haw)). Rooting the tree based on heterozygosities of leaf populations (as implemented in the *rooted.njtree.builder* function) would actually be quite sensitive to long-branch attraction by the highly diverged US-Haw (Figure 3) resulting in positionning US-Haw as the outgroup. Yet, based on historical knowledge of the introduced status of this population, we here chose to set the root of the tree on the branch relating the common ancestor of the Chinese populations (CN-Bei and CN-Shi) and the common ancestor of JP-Sap and US-Haw. Such a positioning of the root is consistent with a (recent) Japanese origin of the Hawaiian population (Fraimout *et al.* 2017). We finally obtained the scaffold tree represented in Figure 7a fitted as follows:

```
#define (manually) the rooted scaffold tree
scaf.tree=rbind(c("CN","R",""),
                c("JP","R",""),
                c("CN-Bei","CN",""),
                c("CN-Shi","CN",""),
                c("JP-Sap","JP",""),
                c("US-Haw","JP",""))
#fit the scaffold tree (with edges lengths drift units and 95% CI estimated)
scaf.tree.params<-generate.graph.params(scaf.tree,fstats = dsu.fstats)
```

```
Total Number of Parameters: 5 (5 edges lengths + 0 adm. coeff.)
Total Number of Statistics: 6 (3 F2 and 3 F3)
```

```
scaf.tree.fit<-fit.graph(scaf.tree.params,drift.scaling = TRUE,compute.ci = TRUE)
```

```
Estimation started (direct algebraic solution)
Estimation ended in 0 m  0 s
    Final Score: 1.124219
    BIC: 97.95432
```

```
#plot the tree
plot(scaf.tree.fit)
```

```
#CI of edges lengths

```

```
                95% Inf.    95% Sup. 95% Inf. (drift scaled) 95% Sup. (drift scaled)
R<->CN      0.0025699799 0.003649279            0.022374993             0.031771684
R<->JP      0.0025699799 0.003649279            0.022374993             0.031771684
CN<->CN-Bei 0.0009098188 0.001161736            0.006498229             0.008297507
CN<->CN-Shi 0.0038994202 0.005559202            0.027850956             0.039705673
JP<->JP-Sap 0.0052402047 0.007662313            0.034940631             0.051090761
JP<->US-Haw 0.0365762281 0.039862096            0.243882932             0.265792437
```

##### 6.2.2 Positionning separately each remaining native populations on the scaffold tree

We then positioned the three other native populations (CN-Nin, CN-Lia and JP-Tok) on the scaffold tree as follows:

```
for(p in c("CN-Nin","CN-Lia","JP-Tok")){
  add.p=add.leaf(scaf.tree.fit,leaf.to.add = p,dsu.fstats,verbose=FALSE,
                 drift.scaling=TRUE,compute.ci=TRUE)
  add.p.delta.bic=format(sort(add.p$bic-min(add.p$bic))[2],digits = 3)
  add.p.comp.fit=compare.fitted.fstats(dsu.fstats,add.p$best.fitted.graph,
                                       n.worst.stats = 0)
  add.p.worst.fit=add.p.comp.fit[which.max(abs(add.p.comp.fit$`Z--score`)),]
  plot(add.p$best.fitted.graph)
}
```

As shown in Figure 7, all the three populations could unambiguously be placed onto the scaffold tree as admixed populations with a good fit of each resulting graphs. Indeed, for each set of five populations (four scaffold one and the added population), the $BIC$ of the best fitting graph was always far lower than all the other possible graphs ($\Delta_{BIC} > 10$). Note that for JP-Tok $\Delta_{BIC} = 0$ in Figure 7d because the graph with the second lowest BIC corresponds to the positioning of the admixed source S1-JP-Tok to the other

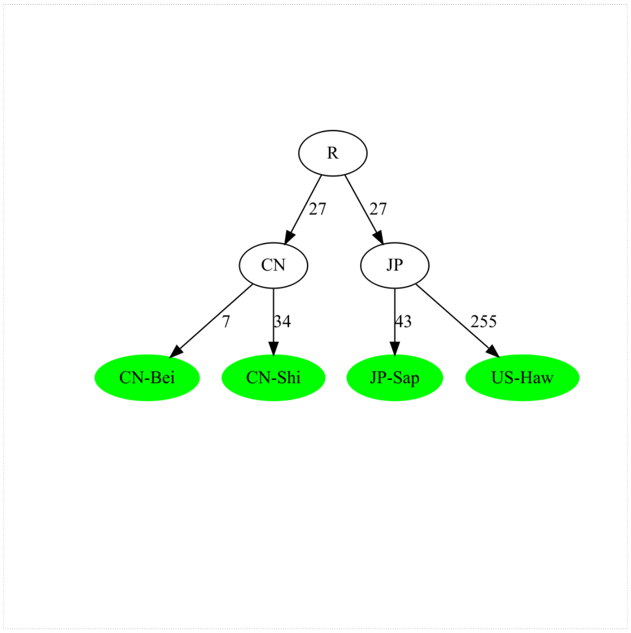

(a) Scaffold

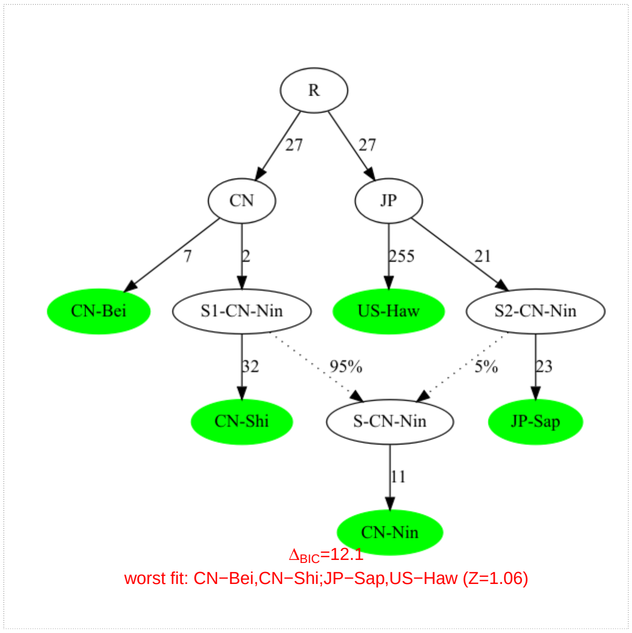

(b) Placing CN-Nin onto scaffold

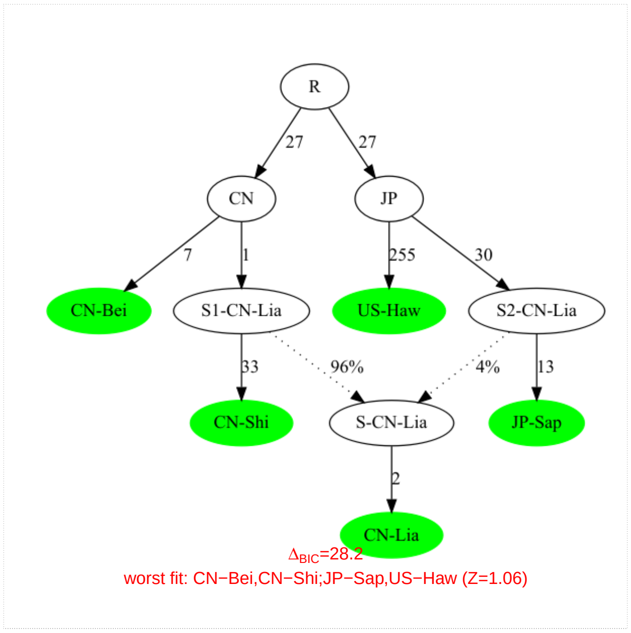

(c) Placing CN-Lia onto scaffold

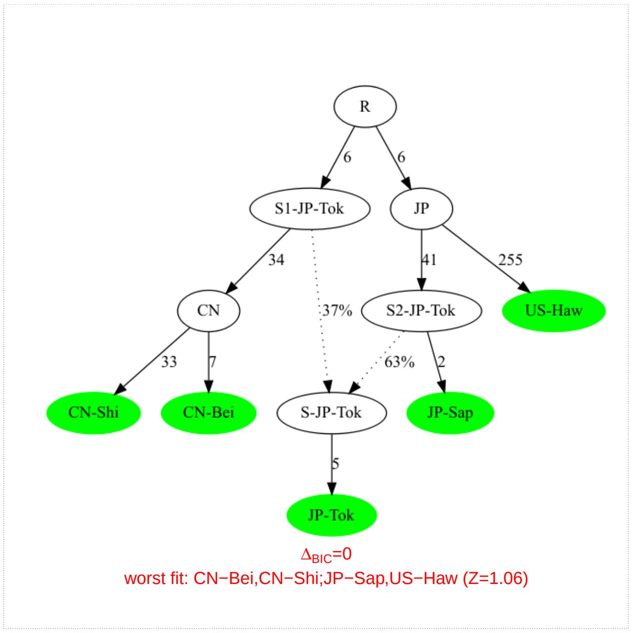

(d) Placing JP-Tok onto scaffold

Figure 7: Inference of admixture graph connecting the populations from the native area (extended to Hawaii). The scaffold tree is displayed in a) together with the best inferred admixture graph obtained by the (independent) positionning of the three other populations from the native area, CN-Nin (b); CN-Lia (c) and JP-Tok (d) as obtained with the add.leaf function. For each of the three latter graphs, the worst fitted f-statistics and its associated Z-score is given and the difference of BIC of the graph with the graphs displaying the second lowest BIC is provided ($\Delta_{BIC}$) as a measure of support. For all graphs, the fitted edge lengths are in drift units (x1000) since *drift.scaling* argument was set to *TRUE*.

alternative branch which is strictly equivalent to the graph displayed (as the two branches coming from R are not identifiable). These two equivalent graphs are highly supported since $\Delta_{BIC} = 20.4$ between either of them and the graph with the third lowest BIC).

##### 6.2.3 Searching for the most comprehensive admixture graph relating all native populations

We next relied on the *graph.builder* function to try to jointly include all the three remaining populations (CN-Lia, CN-Nin and JP-Tok) on the graph. As a first step, we considered simply adding two of these populations simultaneously, leading to 6 different runs of the function (3 pairs of populations and 2 orders of inclusion for each pair. Importantly, we carefully evaluated the fit of each of the best fitting graph identified using the *compare.fitted.fstats* function:

```
#Creating the six pairs of leaves to add for graph.builder
#a custom function to remove sets (generate by expand grid) with non unique pop
cleangrid<-function(x){n=ncol(x);x[apply(x,1,f<-function(y){length(unique(y))==n}),n:1]}
pops.to.include<-expand.grid(c("CN-Nin","CN-Lia","JP-Tok"),
                             c("CN-Nin","CN-Lia","JP-Tok")) %>% as.matrix() %>% cleangrid()
#exploring  and evaluating the graphs
for(i in 1:nrow(pops.to.include)){
  cat("###########################################\n")
  cat("Inclusion Order:",pops.to.include[i,],"\n")
  tmp.graph<-graph.builder(scaf.tree.fit,leaves.to.add = pops.to.include[i,],
                           fstats=dsu.fstats,verbose=F)
  #number of graphs within Dbic<6 (default)
  cat("\tNumber Of Graphs:",tmp.graph$n.graphs,"\n")
  #min BIC
  cat("\tMin BIC:",min(tmp.graph$bic),"\n")
  #worst fitted fstats (best fitting graph)
  cat("\tWorst fstats fit:\n")
  tmp.comp.fit=compare.fitted.fstats(dsu.fstats,tmp.graph$best.fitted.graph,
                                     n.worst.stats = 0)
  print(tmp.comp.fit[which.max(abs(tmp.comp.fit$`Z--score`)),])
  }
```

```
###########################################
Inclusion Order: CN-Nin CN-Lia
    Number Of Graphs: 2
    Min BIC: 332.6374
    Worst fstats fit:
                                Estimated Fitted Z-score
CN-Bei,CN-Lia;JP-Sap,US-Haw -0.0002110636      0 4.995489
###########################################
Inclusion Order: CN-Nin JP-Tok
    Number Of Graphs: 2
    Min BIC: 308.1103
    Worst fstats fit:
                                Estimated      Fitted  Z-score
CN-Shi,US-Haw;JP-Sap,JP-Tok -0.0005025053 -0.00067728 -1.530277
###########################################
Inclusion Order: CN-Lia CN-Nin
    Number Of Graphs: 2
    Min BIC: 316.4586
    Worst fstats fit:
                                Estimated       Fitted Z-score
CN-Bei,CN-Nin;JP-Sap,US-Haw -0.0001769363 3.469447e-18  3.62954
```

```
##########################################
Inclusion Order: CN-Lia JP-Tok
    Number Of Graphs: 6
    Min BIC: 299.4749
    Worst fstats fit:
                                Estimated        Fitted Z-score
CN-Bei,CN-Shi;JP-Sap,US-Haw -5.079243e-05 -3.469447e-18 1.060292
##########################################
Inclusion Order: JP-Tok CN-Nin
    Number Of Graphs: 2
    Min BIC: 368.2099
    Worst fstats fit:
                              Estimated       Fitted  Z-score
CN-Bei,US-Haw;CN-Shi,JP-Tok 0.005775447 6.938894e-18 -7.207447
##########################################
Inclusion Order: JP-Tok CN-Lia
    Number Of Graphs: 6
    Min BIC: 299.4749
    Worst fstats fit:
                                Estimated Fitted Z-score
CN-Bei,CN-Shi;JP-Sap,US-Haw -5.079243e-05      0 1.060292
```

Note the benefits of exploring all the possible orders of inclusion with *graph.builder* for a given set of populations. For instance, the best fitted graph obtained when including JP-Tok and CN-Nin displayed a highly lower BIC if CN-Nin was the first to be included. Conversely, in some cases, the same best fitting graph may be obtained with the different possible inclusion orders (e.g., see the results obtained with the CN-Lia and JP-Tok pair).

Interestingly, this exploration suggests that CN-Lia, CN-Nin and JP-Tok may not be included jointly to the scaffold tree to obtain a comprehensive admixture graph for the seven native populations. Indeed, and although these two populations are closely related, the graphs obtained when adding both CN-Nin and CN-Lia to the scaffold tree displayed poor fitted f-stats (some with associated $|Z|>3.95$). This was confirmed when trying to add the three populations simultaneously (and exploring all the 6 possible orders of inclusion) with *graph.builder* as shown below:

```
###a custom function to list all possible orders of inclusion (permutations)
#adapted from https://www.r-bloggers.com/2019/06/learning-r-permutations-and-combinations-with-base-r/
permutations <- function(x) {
          n <- length(x)
          if (n == 1){out<-x}else{
                out <- NULL
                for (i in 1:n){out <- rbind(out, cbind(x[i], permutations(x[-i])))}
               }
          return(out)
}
##
pops.to.include<-permutations(c("CN-Nin","CN-Lia","JP-Tok"))
#exploring  and evaluating the graphs
for(i in 1:nrow(pops.to.include)){
  cat("##########################################\n")
  cat("Inclusion Order:",pops.to.include[i,],"\n")
  tmp.graph<-graph.builder(scaf.tree.fit,leaves.to.add = pops.to.include[i,],
                           fstats=dsu.fstats,verbose=F)
  #number of graphs within Dbic<6 (default)
  cat("\tNumber Of Graphs:",tmp.graph$n.graphs,"\n")
```

```r
  #min BIC
  cat("\tMin BIC:",min(tmp.graph$bic),"\n")
  #worst fitted fstats (best fitting graph)
  cat("\tWorst fstats fit:\n")
  tmp.comp.fit=compare.fitted.fstats(dsu.fstats,tmp.graph$best.fitted.graph,
                                     n.worst.stats = 0)
  print(tmp.comp.fit[which.max(abs(tmp.comp.fit$`Z--score`)),])
  }
```

```
###########################################
Inclusion Order: CN-Nin CN-Lia JP-Tok
    Number Of Graphs: 2
    Min BIC: 480.5329
    Worst fstats fit:
                                Estimated Fitted Z-score
CN-Bei,CN-Lia;JP-Sap,US-Haw -0.0002110636      0 4.995489
###########################################
Inclusion Order: CN-Nin JP-Tok CN-Lia
    Number Of Graphs: 5
    Min BIC: 480.5329
    Worst fstats fit:
                                Estimated        Fitted Z-score
CN-Bei,CN-Lia;JP-Sap,US-Haw -0.0002110636 -3.469447e-18 4.995489
###########################################
Inclusion Order: CN-Lia CN-Nin JP-Tok
    Number Of Graphs: 2
    Min BIC: 463.636
    Worst fstats fit:
                                Estimated        Fitted Z-score
CN-Lia,JP-Tok;CN-Nin,CN-Shi -0.0007377375 -0.0004944342 3.733745
###########################################
Inclusion Order: CN-Lia JP-Tok CN-Nin
    Number Of Graphs: 11
    Min BIC: 513.4301
    Worst fstats fit:
                              Estimated      Fitted  Z-score
CN-Lia,US-Haw;CN-Shi,JP-Tok 0.005572856 0.000100214 -7.206377
###########################################
Inclusion Order: JP-Tok CN-Nin CN-Lia
    Number Of Graphs: 8
    Min BIC: 518.0997
    Worst fstats fit:
                              Estimated       Fitted  Z-score
CN-Lia,JP-Tok;CN-Nin,US-Haw 0.004911067 -0.000220809 -7.205941
###########################################
Inclusion Order: JP-Tok CN-Lia CN-Nin
    Number Of Graphs: 8
    Min BIC: 518.0997
    Worst fstats fit:
                              Estimated       Fitted  Z-score
CN-Lia,JP-Tok;CN-Nin,US-Haw 0.004911067 -0.000220809 -7.205941
```

Overall, only six out of the seven populations from the native area (extended to Hawaii) could be related with an admixture graph with good fitting properties i.e., either by adding to the scaffold tree (consisting of

US-Haw, JP-Sap, CN-Bei and CN-Shi) the population Jp-Tok and either CN-Lia or CN-Nin. Note that the two resulting best fitting graph cannot be compared based on the BIC criterion as the two sets of populations differ. A closer inspection of the different graphs obtained (6 within $\Delta_{BIC} < 6$ with the best fitting graph for the CN-Lia, Jp-Tok included pairs and only one for the CN-Nin, Jp-Tok pair) made us chose as the most relevant graph the graph displayed in Figure 8. As a matter of fact, this graph had the lowest absolute Z-score for the worst fitted f-stats (that was actually the same as the one for the scaffold tree) and it was also quite consistent with the one obtained when adding each population individually to the scaffold tree (Figure 7c and 7d).

```
natgraph<-graph.builder(scaf.tree.fit,leaves.to.add = c("CN-Lia","JP-Tok"),dsu.fstats,
                         verbose=F,drift.scaling=T,compute.ci=TRUE)
plot(natgraph$best.fitted.graph)
```

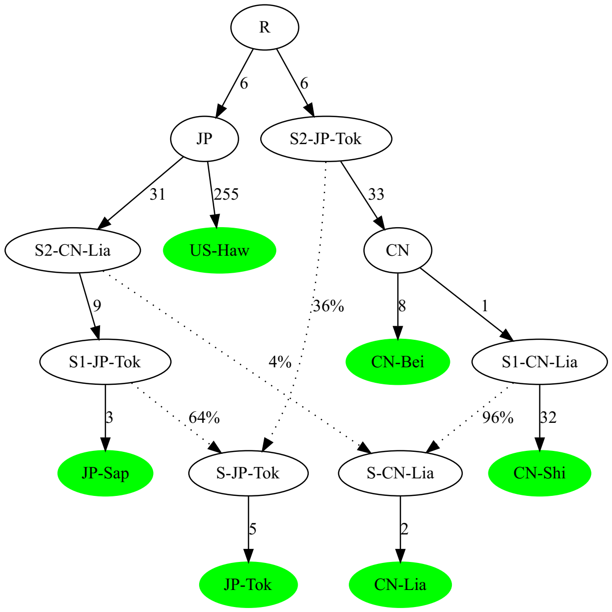

Figure 8: Best fitting admixture graph connecting six out of the seven populations representative of the native area (extended to Hawaii). The fitted edge lengths are in drift units (x1000) since *drift.scaling* argument was set to *TRUE*.

The difficulties encountered to jointly add CN-Lia and CN-Lin may be related to the fact that they are only very slightly differentiated (Figures 3 and 4) making it difficult to simplify their joint history with simple two-way admixture graphs (likely due to a high amount of gene flow between them). It should also be noticed that the CN-Nin sample may be slightly heterogeneous as it mixes individuals from very close locations but sampled at two different times (Table 1).

##### 6.2.4 Estimates of the parameters of the admixture graph for the native and Hawaiian populations and their confidence intervals

The CI's for branch lengths (on both unscaled and drift scaled units) of the admixture graph connecting the 6 populations Figure 8 together with the two admixture proportions are directly available from the *natgraph* object (as it was generated with options *drift.scaled=TRUE* and *compute.ci=TRUE*):

```
#branch lengths
natgraph$
```

```
                          95% Inf.     95% Sup. 95% Inf. (drift scaled) 95% Sup. (drift scaled)
S-JP-Tok<->JP-Tok     6.315686e-04 0.0007827838            0.0044362902             0.005498462
S2-CN-Lia<->S1-JP-Tok 8.258283e-04 0.0018089613            0.0058431246             0.012799255
```

```
R<->S2-JP-Tok         4.876279e-04 0.0009225970           0.0043358045           0.008203386
S1-JP-Tok<->JP-Sap    3.915283e-04 0.0005084688           0.0027874828           0.003620039
S2-JP-Tok<->CN        4.100608e-03 0.0054398001           0.0287396430           0.038125541
S-CN-Lia<->CN-Lia     2.491030e-04 0.0003931522           0.0017960161           0.002834601
CN<->S1-CN-Lia        7.342082e-05 0.0002078678           0.0005283615           0.001495888
JP<->S2-CN-Lia        4.088231e-03 0.0052797637           0.0272544120           0.035197830
S1-CN-Lia<->CN-Shi    3.821499e-03 0.0051975302           0.0273294661           0.037170156
R<->JP                4.876279e-04 0.0009225970           0.0043358045           0.008203386
CN<->CN-Bei           9.842052e-04 0.0011107899           0.0070826785           0.007993626
JP<->US-Haw           3.675243e-02 0.0397726523           0.2450120628           0.265146533
```

```
#admixture proportions
#The names were automatically given by graph.builder as A (and 1-A) for ancestry proportion
#contributing to the first (CN-Lia) and B for ancestry proportion contributing to
#the second (JP-Tok) included populations
natgraph$
```

```
   95% Inf.  95% Sup.
B 0.6312323 0.6576909
A 0.9566978 0.9633939
```

Alternatively, given the topology of the graph, it is also possible to estimate the proportion of Chinese ancestry in the CN-Lia population using ratios of $f_4$ estimates as:

```
#Prop of Chinese ancestry in CN-Lia (A parameter defined above)
#The ratio F4(JP-Sap,CN-Shi;CN-Bei,US-Haw)/F4(JP-Sap,CN-Lia;CN-Bei,US-Haw)
#estimates A as can be shown with symbolic computations of F4
natgraph.par<-generate.graph.params(natgraph$best.fitted.graph@graph,verbose=FALSE)
natgraph.symbolicfstats<-graph.params2symbolic.fstats(natgraph.par)
natgraph.symbolicfstats$F4.equations[
  grep("F4\\(JP-Sap,CN-Shi;CN-Bei,US-Haw\\)",natgraph.symbolicfstats$F4.equations)]
```

```
[1] "F4(JP-Sap,CN-Shi;CN-Bei,US-Haw) = -(S2-JP-Tok<->CN+R<->S2-JP-Tok+R<->JP)"
```

```
natgraph.symbolicfstats$F4.equations[
  grep("F4\\(JP-Sap,CN-Lia;CN-Bei,US-Haw\\)",natgraph.symbolicfstats$F4.equations)]
```

```
[1] "F4(JP-Sap,CN-Lia;CN-Bei,US-Haw) = -(R<->JP+S2-JP-Tok<->CN+R<->S2-JP-Tok)*A"
```

```
#estimation via ratio of F4
alpha_lia=compute.f4ratio(dsu.fstats,num.quadruplet = c("CN-Bei","US-Haw","CN-Lia","JP-Sap"),
                                     den.quadruplet = c("CN-Bei","US-Haw","CN-Shi","JP-Sap"))
alpha_lia[2]
```

```
bjack mean
 0.9559757
```

```
#95% CI
alpha_lia[2] + c(-1.96,1.96)*alpha_lia[3]
```

```
[1] 0.9435186 0.9684329
```

The resulting estimates of Chinese ancestry in the CN-Lia population are thus consistent. However, as expected from the simulation study (see the main text of the manuscript), the confidence intervals estimated with the graph fitting procedure are smaller (and presumably too optimistic) than the ones estimates with the $f_4$-ratio which may be considered as more reliable.

Interestingly, a similar proportions of Chinese ancestry could be obtained using $f_4$-ratio for CN-Nin than for CN-Lia (assuming CN-Nin and CN-Lia have a similar ancestry with respect to the other populations). It was also consistent with the one obtained in Figure 7b:

```
#Prop of Chinese ancestry in CN-Lia
alpha_nin=compute.f4ratio(dsu.fstats,num.quadruplet = c("CN-Bei","US-Haw","CN-Nin","JP-Sap"),
                                     den.quadruplet = c("CN-Bei","US-Haw","CN-Shi","JP-Sap"))
alpha_nin[2]
```

```
bjack mean
 0.9469928
```

```
#95% CI
alpha_nin[2] + c(-1.96,1.96)*alpha_nin[3]
```

```
[1] 0.9342755 0.9597100
```

Note that no such $f_4$-ratio based estimation of admixture proportions could be carried for the JP-Tok population as no outgroup was available for one of its ancestry (i.e., one source connects directly to the root).

#### 6.3 Investigating the origins of the populations from the American invasive area using admixture graphs

The $f_3$-based tests of admixture (section 5) showed that four (US-Col, US-Nca, US-Wat and US-Wis) of the seven American populations displayed admixture signals. In addition, all involved the most significant ones always involved a source related to a population from the invasive area suggesting that that were not directly connected to the first invader. We will thus first attempt to position the remaining American populations (BR-Pal, US-Sdi and US-Sok) with respect to the native and Hawaiian populations.

##### 6.3.1 Positioning BR-Pal, US-Sdi and US-Sok with respect to the native populations

We investigated the relationships of the three American populations, BR-Pal, US-Sdi and US-Sok, with the native populations by identifying with the *add.leaf* function the best positioning of each population onto either the admixture graph (Figure 8) or the scaffold tree (Figure 7a) obtained above using the following command (output not shown):

```
for(p in c("BR-Pal","US-Sdi","US-Sok")){
  #positioning onto the admixture graph of native pops
  add.p=add.leaf(natgraph$best.fitted.graph,leaf.to.add = p,dsu.fstats,verbose=FALSE,
                drift.scaling=TRUE,compute.ci=TRUE)
  plot(add.p$best.fitted.graph)
  add.p.comp=compare.fitted.fstats(dsu.fstats,add.p$best.fitted.graph,n.worst.stats = 1)
  #positioning onto the scaffold tree of native pops
  add.p=add.leaf(scaf.tree.fit,leaf.to.add = p,dsu.fstats,verbose=FALSE,
                drift.scaling=TRUE,compute.ci=TRUE)
  plot(add.p$best.fitted.graph)
  add.p.comp=compare.fitted.fstats(dsu.fstats,add.p$best.fitted.graph,n.worst.stats = 1)
}
```

As summarized in Figure 9, the three populations could be placed with a good resulting fit to each graph of native populations. The Z-score of the worst fitted f-statistics was always $< 2$ in absolute value except for the admixture resulting from the placing of US-Sok on the admixture graph of native populations (Z=-4.14; Figure 9E). In addition, for each population, the positioning onto the two graphs was concordant and suggested that they were all admixed although this was not detected by the $f_3$ based test of admixture (Table 3). This is likely due to a too strong subsequent divergence from their ancestral admixed source. All the three populations shared an Hawaiian (ancestral to US-Haw) source, the other source being of presumably Chinese origin for both US-Sdi and BR-Pal and of Japanese origin (ancestral to JP-Sap) for US-Sok. For BR-Pal, it should be noticed that the support for the positioning onto the scaffold tree (Figure 9B) remained loose

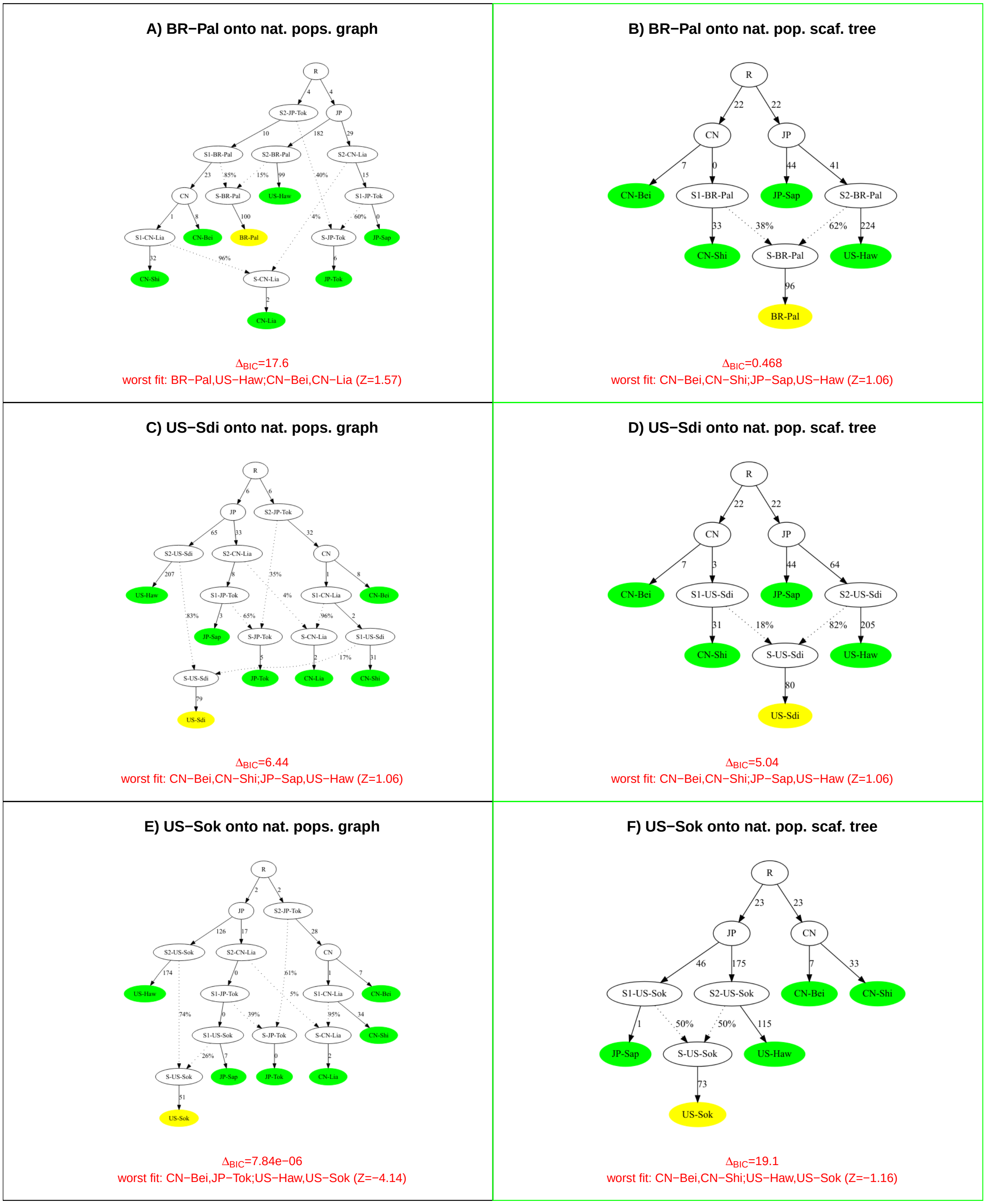

Figure 9: Positionning of the BR-Pal, US-Sdi and US-Sok populations from the American invasive areas onto the admixture graph connecting the six native and Hawaiian populations (Figure 8) and the scaffold tree of native (and Hawaiian) populations (Figure 7a). The best fitting graphs (as obtained with the function add.leaf) are displayed together with i) the worst fitted f-statistics and their associated Z-score; and ii) the difference of their BIC with respect to the graphs displaying the second lowest BIC ($\Delta_{BIC}$) as a measure of support. The target populations are highlighted in yellow. For all the graphs, the fitted edge lengths are in drift units (x1,000) since *drift.scaling* argument was set to *TRUE*.

($\Delta_{BIC} < 6$) and a closer inspection of the corresponding *add.leaf* results showed that three alternative graphs resulted in a BIC very similar to this best fitting graph:

```
addPal=add.leaf(scaf.tree.fit,leaf.to.add = "BR-Pal",dsu.fstats,verbose=FALSE,
                drift.scaling=T,compute.ci=T)
#Difference of BIC wrt the best fitting graph
addPal.dbic=addPal$bic-min(addPal$bic)
format(addPal.dbic,digits = 3)
```

```
 [1] "111.581" "111.581" "246.719" "246.647" "169.198" " 27.333" "111.581" "116.186" "115.099"
[10] "116.186" "  0.468" "116.186" "115.099" "116.186" "  0.468" "251.252" "116.186" "  0.468"
[19] "115.099" "  0.000" " 31.939"
```

As shown in Figure 10, these corresponded to the branching of S1-BR-Pal source on i) the CN<->CN-Bei branch (Figure 10D) with a null distance from CN which is actually the same as Figures 9B and 10A (the estimated admixture proportions being also the same); ii) the R<->CN branch (Figure 10B) or strictly equivalently the R<->JP branch (Figure 10C). The two latter did not lead to any null branches (which is usually preferable) and the branching of S1-BR-Pal was always closer to CN ($10\times10^{-3}$ and $11\times10^{-3}$ drift units respectively, recalling that the positioning of the root R is arbitrarily set in the middle of the JP<->CN unrooted branch) than JP in agreement with a Chinese origin. This is concordant with Figures 9A) in which the presence of JP-Tok and its presumably Chinese ancestral source (automatically named S2-JP-Tok) allows to rule out this latter positioning of S1-BR-Pal with a high support ($\Delta_{BIC} = 17.6$). Note also that this resulted in either cases in an higher estimated Chinese ancestry for BR-Pal.

For US-Sdi, the three same types of branching as for BR-Pal were within 5.04 BIC units of the best fitting graph (i.e., positioning of the S1-US-Sdi source with respect to CN in Figure 9D), i.e., the support for these alternative graphs was clearly lower according to the BIC criterion (and the absolute Z-score for the worst fitted f-stats was also higher) as shown with the following codes:

```
addSdi=add.leaf(scaf.tree.fit,leaf.to.add = "US-Sdi",dsu.fstats,verbose=FALSE,
                drift.scaling=T,compute.ci=T)
#Difference of BIC wrt the best fitting graph
addSdi.dbic=addSdi$bic-min(addSdi$bic)
format(addSdi.dbic,digits = 3)
```

```
 [1] "185.91" "185.91" "541.90" "541.30" "206.66" " 13.04" "185.91" "190.52" "187.77" "190.52"
[11] "  5.04" "190.52" "187.77" "190.52" "  5.04" "545.90" "190.52" "  5.04" "187.77" "  0.00"
[21] " 17.65"
```

```
#worst fitted fstats for the three graphs within 6 BIC units from the best one
sel.idx=which(addSdi.dbic<6)
sel.idx=sel.idx[order(addSdi.dbic[sel.idx])]
sel.idx=sel.idx[-1] #remove the best fitting graph
for(i in 1:length(sel.idx)){
tmp.comp=compare.fitted.fstats(dsu.fstats,addSdi$fitted.graphs.list[[sel.idx[i]]],
                               n.worst.stats = 1)
}
```

```
1 Worst fit for:
                               Estimated       Fitted  Z-score
CN-Bei,CN-Shi;US-Haw,US-Sdi 9.460063e-05 3.469447e-18 -2.410857
1 Worst fit for:
                               Estimated       Fitted  Z-score
CN-Bei,CN-Shi;US-Haw,US-Sdi 9.460063e-05 3.469447e-18 -2.410857
1 Worst fit for:
                               Estimated       Fitted  Z-score
CN-Bei,CN-Shi;US-Haw,US-Sdi 9.460063e-05 3.469447e-18 -2.410857
```

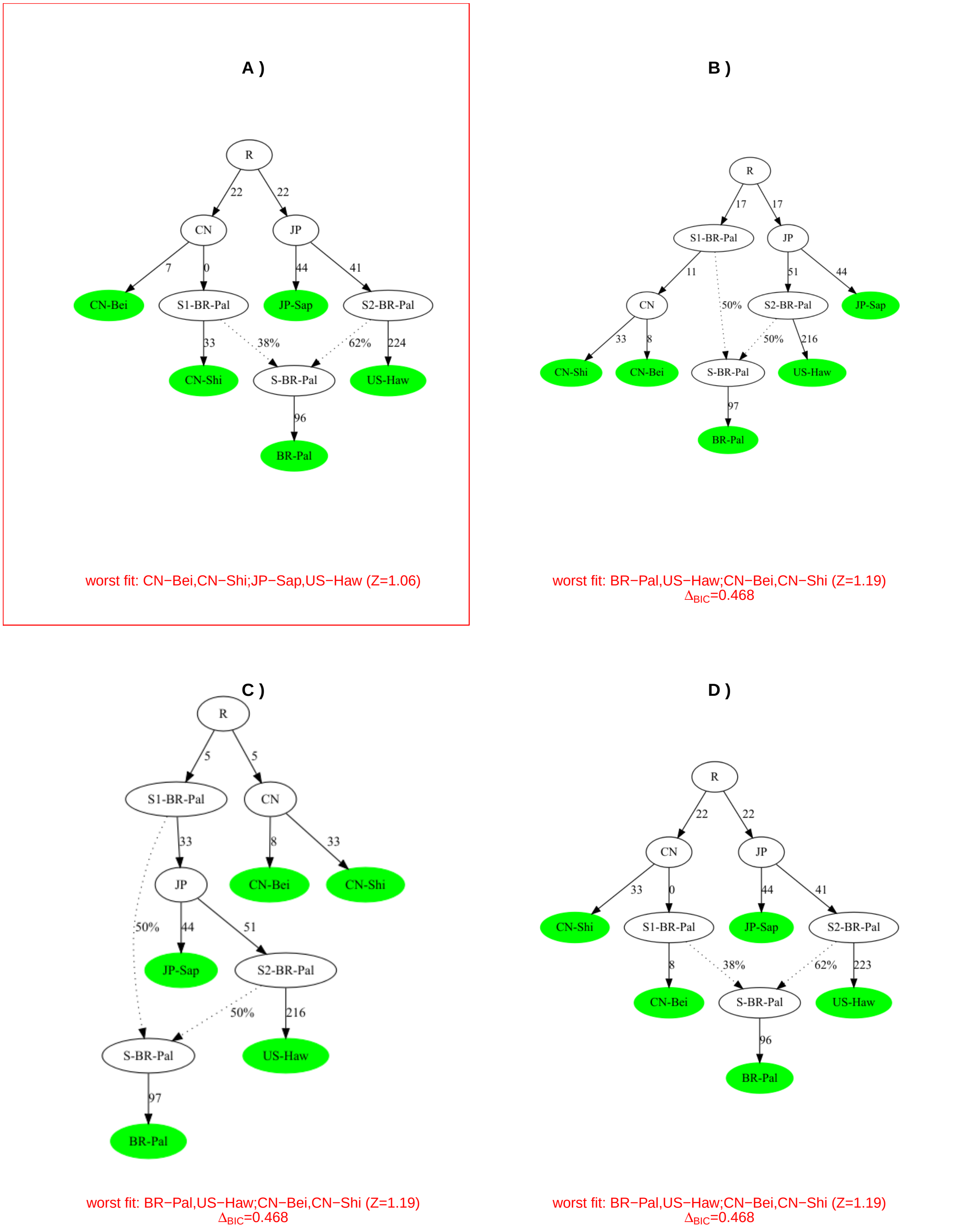

Figure 10: Admixture graphs resulting from the positioning of BR-Pal onto the scaffold tree of native and Hawaiian populations (Figure 7a) with BIC less than 6 units higher than the BIC with the best fitting graph (within red box and represented in Figure 9B). For each population, the graph (as obtained with the function add.leaf) is displayed together with i) the worst fitted f-statistics and its associated Z-score; and ii) the difference of BIC of the graph with the graphs displaying the best fitting graph ($\Delta_{BIC}$) as a measure of support. For all the graphs, the fitted edge lengths are in drift units (x1,000) since *drift.scaling* argument was set to *TRUE*.

In addition, the positions of US-Sdi on the extended graph and the scaffold graph of native populations were concordant (Figures 9C and 9D) making the position of the Chinese source of US-Sdi on the CN<->CN-Shi more likely. The distance of S1-US-Sdi from CN was also non null (as opposed to the distance separating S1-BR-Pal from CN in Figure 9B).

Finally the position of US-Sok on the scaffold tree of native populations was highly supported (Figure 9F). However, its positioning onto the extended graph of native populations was less clear (Figure 9E) with two other alternative graphs with almost the same BIC as the best fitted graph:

```
addSok=add.leaf(natgraph$best.fitted.graph,leaf.to.add = "US-Sok",dsu.fstats,verbose=FALSE,
                drift.scaling=T,compute.ci=T)
#Difference of BIC wrt the best fitting graph
addSok.dbic=addSok$bic-min(addSok$bic)
#worst fitted fstats for the three graphs within 6 BIC units from the best one
sel.idx=which(addSok.dbic<6)
sel.idx=sel.idx[order(addSok.dbic[sel.idx])]
sel.idx=sel.idx[-1] #remove the best fitting graph
#Delta_BIC (wrt the best fitted graph)
format(addSok.dbic[sel.idx],digits = 3)
```

```
[1] "7.84e-06" "9.73e-06"
```

```
for(i in 1:length(sel.idx)){
tmp.comp=compare.fitted.fstats(dsu.fstats,addSok$fitted.graphs.list[[sel.idx[i]]],
                               n.worst.stats = 1)
}
```

```
1 Worst fit for:
                               Estimated       Fitted   Z-score
CN-Bei,JP-Tok;US-Haw,US-Sok 0.002670761 0.0002218372 -4.139588
1 Worst fit for:
                               Estimated       Fitted   Z-score
CN-Bei,JP-Tok;US-Haw,US-Sok 0.002670761 0.0002218628 -4.139545
```

These actually only differed by the position of the Japanese source named S1-US-Sok in Figure 9E with respect to the Japanese sources of CN-Lia (S2-CN-Lia) and JP-Tok (S1-JP-Tok), the estimated distances between these three sources S1-US-Sok, S2-CN-Lia and S1-JP-Tok being null. Hence, simplifying the graph by combining the three source into a single one lead to a similar fit (but lower BIC as the number of parameters is reduced):

```
tst.SokGraph=rbind(c("S-JP-Tok","S1-JP-Tok","B"),c("S-JP-Tok","S2-JP-Tok","(1-B)"),
               c("JP-Tok","S-JP-Tok",""), c("S1-JP-Tok" ,"JP", ""),
               c("S2-JP-Tok","R",""),c("JP-Sap","S1-JP-Tok",""),
               c("CN","S2-JP-Tok",""),c("S-CN-Lia","S1-CN-Lia","A"),
               c("S-CN-Lia","S1-JP-Tok","(1-A)"),c("CN-Lia","S-CN-Lia",""),
               c("S1-CN-Lia","CN",""),c("CN-Shi","S1-CN-Lia",""),c("JP","R",""),
               c("CN-Bei","CN",""),c("US-Haw","S1-US-Sok",""),
               c("US-Sok","S-US-Sok",""),c("S-US-Sok","S1-JP-Tok","C"),
               c("S-US-Sok","S1-US-Sok","(1-C)"),c("S1-US-Sok","JP",""))
tst.SokGraph.par=generate.graph.params(tst.SokGraph,dsu.fstats)
```

```
Total Number of Parameters: 15 (12 edges lengths + 3 adm. coeff.)
Total Number of Statistics: 21 (6 F2 and 15 F3)
```

```
tst.SokGraph.fit=fit.graph(tst.SokGraph.par,drift.scaling = T,verbose=FALSE)
tst.SokGraph.fit@bic
```

```
[1] 435.3341
```

```
tmp.comp=compare.fitted.fstats(dsu.fstats,tst.SokGraph.fit,n.worst.stats = 1)
```

```
1 Worst fit for:
                              Estimated      Fitted  Z-score
CN-Bei,JP-Tok;US-Haw,US-Sok 0.002670761 0.000221864 -4.139543
```

Yet this simplified version of the Figure 9E graph should be interpreted with caution as i) the absolute Z-score for the worst fitted f-statistics remains high (Z=-4.14); ii) the estimated distance (in drift units) of JP-Tok from its source S-JP-Tok is null; and iii) it somewhat contradicts the original graph displayed in Figure 8 (e.g., in which the two Japanese sources of CN-Lia and JP-Tok are clearly differentiated). Overall this suggests more complex relationships among the Japanese sources of US-Sok and CN-Lia and JP-Tok from the native area that cannot be captured by the sample at hand (or even modeled as a simple admixture graph).

To simplify the modeling of the demographic history of the American invasive populations, we further used the scaffold tree of native populations to represent their ancestry. Hence, we discarded the two admixed native populations CN-Lia and JP-Tok. Indeed, no evidence for a direct contribution of CN-Lia or JP-Tok to the ancestry of BR-Pal, US-Sdi or US-Sok could be found from the above analyses.

##### 6.3.2 Searching for the most comprehensive admixture graph including American invasive populations

We relied on the *graph.builder* function to try to jointly include American invasive populations first focusing on BR-Pal, US-Sdi and US-Sok for the same reasons as mentioned above and evaluated all the possible inclusion order by pair (as previously when analyzing the native populations). We below simplified the output by only printing the analyses that resulted in a best fitted graph with a worst fitted f-statistics Z-score lower than 2 (in absolute value):

```
cleangrid<-function(x){n=ncol(x);x[apply(x,1,f<-function(y){length(unique(y))==n}),n:1]}
pops.to.include<-expand.grid(c("BR-Pal","US-Sdi","US-Sok"),
                             c("BR-Pal","US-Sdi","US-Sok")) %>% as.matrix() %>% cleangrid()
#exploring  and evaluating the graphs
for(i in 1:nrow(pops.to.include)){
 tmp.graph<-graph.builder(scaf.tree.fit,leaves.to.add = pops.to.include[i,],
                           fstats=dsu.fstats,verbose=F)
 tmp.comp.fit=compare.fitted.fstats(dsu.fstats,tmp.graph$best.fitted.graph,
                                     n.worst.stats = 0)
  if(max(abs(tmp.comp.fit$`Z--score`))<2){
  cat("###########################################\n")
  cat("Inclusion Order:",pops.to.include[i,],"\n")
  #number of graphs within Dbic<6 (default)
  cat("\tNumber Of Graphs:",tmp.graph$n.graphs,"\n")
  #min BIC
  cat("\tMin BIC:",min(tmp.graph$bic),"\n")
  #worst fitted fstats (best fitting graph)
  cat("\tWorst fstats fit:\n")
  print(tmp.comp.fit[which.max(abs(tmp.comp.fit$`Z--score`)),])
}
 }
```

```
###########################################
Inclusion Order: US-Sok BR-Pal
    Number Of Graphs: 4
    Min BIC: 271.6173
    Worst fstats fit:
                              Estimated        Fitted  Z-score
CN-Bei,CN-Shi;US-Haw,US-Sok 4.971676e-05 -1.040834e-17 -1.160098
```

As shown, out the six tested (ordered) pairs of included populations, including first US-Sok and BR-Pal resulted in the best fitted graph (worst fitted f-statistics associated Z-score equal to -1.16). As before, three alternative but equivalent graph, since they simply differ according to the positioning of the Chinese related source of BR-Pal, its distance from the CN node being always null (Figure 11).

```
addSokPal=graph.builder(scaf.tree.fit,leaves.to.add = c("US-Sok","BR-Pal"),dsu.fstats,
                        verbose=FALSE,drift.scaling=T,compute.ci=T)
#Difference of BIC wrt the best fitting graph
addSokPal.dbic=addSokPal$bic-min(addSokPal$bic)
format(addSokPal.dbic,digits = 3)
```

```
[1] "0.388" "0.388" "0.388" "0.000"
```

We thus simplified the graph by defining CN (the ancestor of the two Chinese populations CN-Bei and CN-Shi) as the Chinese source of BR-Pal (thereby removing one branch length parameter) and redefined some node names to clarify the corresponding inferred scenario. Indeed all graphs suggested that American invasive populations originated from two independent admixture events among the populations from the native area (including US-Haw):

- A first (older) admixture event between a Japanese and an Hawaiian related sources (called hereafter J-Am1 and H-Am1 respectively) that lead to the internal node population called Am1 which is directly ancestral to US-Sok
- A second (and more recent) admixture event between a source related to Am1 (called S-Am2 hereafter) and a Chinese closely related to the ancestor of CN-Shi and CN-Bei. We hereafter called this second admixed population, directly related to BP-Pal, Am2

The corresponding graph, equivalent to the ones on Figure 11, was defined (and fitted) as follows and plotted on Figure 12 (graph framed in red):

```
am.scaf.graph<-rbind(c('Am2','S-Am2','B'),c('Am2','CN','(1-B)'),c('BR-Pal','Am2',''),
                     c('S-Am2','Am1',''),c('US-Sok','S-Am2',''),c('CN-Shi','CN',''),
                     c('Am1','J-Am1','A'),c('Am1','H-Am1','(1-A)'),c('J-Am1','JP',''),
                     c('JP-Sap','J-Am1',''),c('US-Haw','H-Am1',''),c('H-Am1','JP',''),
                     c('CN','R',''),c('JP','R',''),c('CN-Bei','CN',''))
am.scaf.par=generate.graph.params(am.scaf.graph,dsu.fstats)
```

```
Total Number of Parameters: 12 (10 edges lengths + 2 adm. coeff.)
Total Number of Statistics: 15 (5 F2 and 10 F3)
```

```
am.scaf.fit=fit.graph(am.scaf.par,drift.scaling = T,verbose=FALSE)
am.scaf.fit@bic
```

```
[1] 269.2973
```

```
tmp.comp=compare.fitted.fstats(dsu.fstats,am.scaf.fit,n.worst.stats = 1)
```

```
1 Worst fit for:
                               Estimated Fitted Z-score
BR-Pal,US-Haw;CN-Bei,CN-Shi -5.09979e-05      0 1.187469
```

##### 6.3.3 Positioning US-Col, US-Nca, US-Sdi, US-Wat and US-Wis onto the scaffold graph extended with US-Sok and BR-Pal

We then explored the position of the five remaining American populations (US-Col, US-Nca, US-Sdi, US-Wat and US-Wis) on the scaffold graph *am.scaf* that included BR-Pal and US-Sok using the *add.leaf* function:

As shown on Figure 12, only US-Wat could be placed without ambiguity on the graph ($\Delta_{BIC} > 8$ and worst fitted f-statistic absolute Z-score $< 2$) and could be interpreted as a recently admixed populations between a source related to a (presumably) Northern American population Am1 and the admixed (presumably)

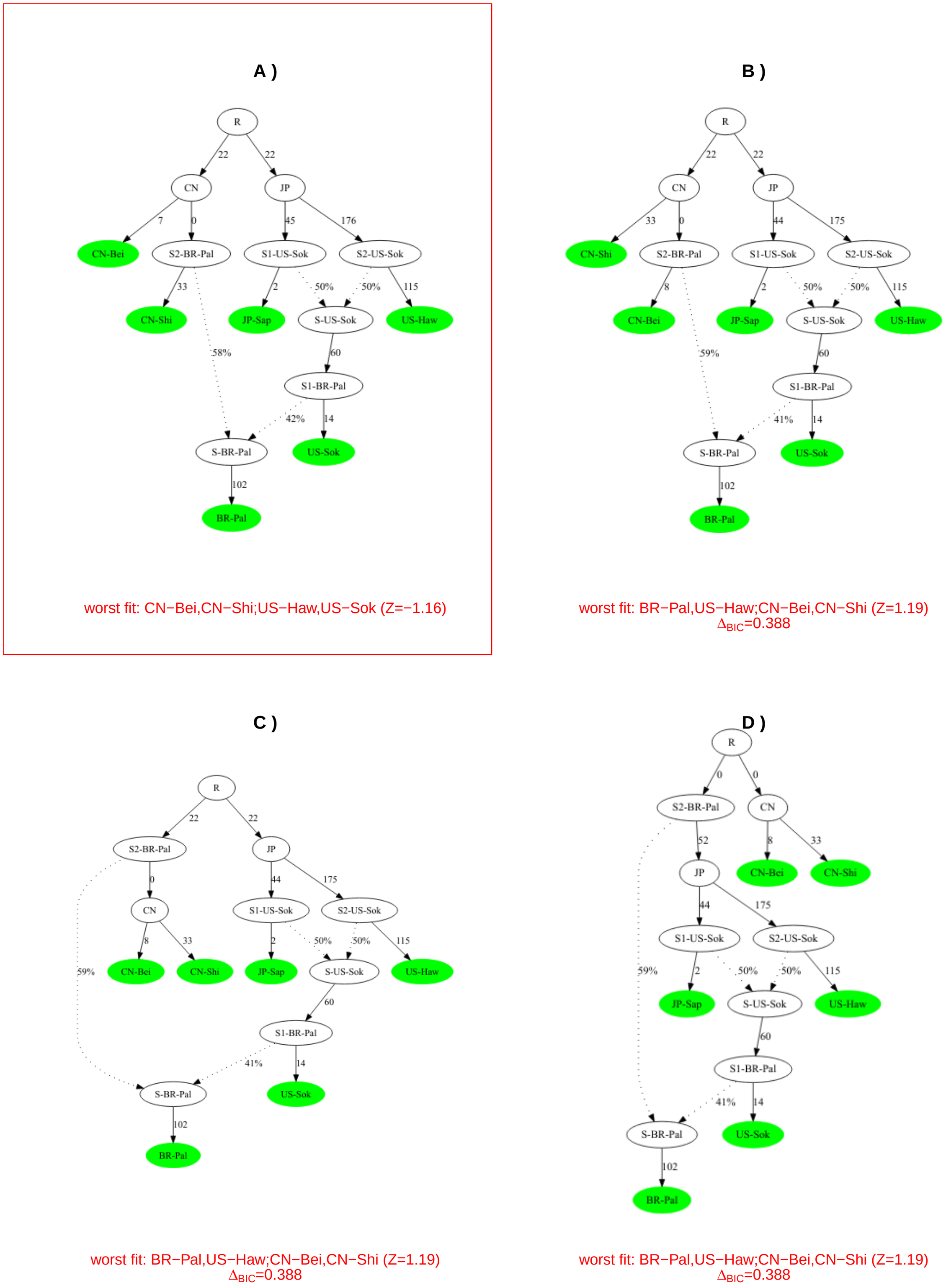

Figure 11: Admixture graphs resulting from the joint positioning of US-Sok and BR-Pal onto the scaffold tree of native populations (Figure 7a) with BIC less than 6 units higher than the BIC with the best fitting graph (within red box and represented in Figure 9B). For each population, the graph (as obtained with the function add.leaf) is displayed together with i) the worst fitted f-statistics and its associated Z-score; and ii) the difference of BIC of the graph with the graphs displaying the best fitting graph ($\Delta_{BIC}$) as a measure of support. For all the graphs, the fitted edge lengths are in drift units (x1,000) since *drift.scaling* argument was set to *TRUE*.

Southern Am2 population. The positioning of US-Sdi also resulted in a best fitting graph with a high support ($\Delta_{BIC} > 33$) and a rather good fit, the worst fitted f-statistic Z-score being equal -2.41. The resulting scenario suggested US-Sdi derived from a population admixed between a source deriving from Am2 (and actually closely related to BR-Pal) and a source related to an (another?) Hawaiian population distantly related to H-Am1 the Hawaiian source of Am1 (that also contributed to Am2).

This suggested that US-Sdi resulted from multiple admixture events between populations closely related the Am2 and Hawaii that are rather difficult to fit with simple admixture graph (worst fitted f-statistics associated Z-score slightly higher than 2). Accordingly, when jointly adding US-Wat and US-Sdi (whatever the order of inclusion) to the *am.scaf* graph, the best fitted graph was consistent with the ones obtained above when adding the US-Sdi and US-Wat, the Am2 related source being common for the two populations (the inferred distance between S1-US-Wat and S1-US-Sdi being null) and the Hawaiian related source of US-Sdi being closely related to H-Am1 (Figure 13A). The inferred distance were also more consistent with geographical and historical origins of the populations and the high contribution of the S1-US-Sdi/S1-Wat source was also consistent with the observed $f_3$ values (see section 5). Nevertheless, the fit remained poor as the worst fitted f-statistics were clearly off (absolute Z-score > 6) and a likely alternative scenario would be that the second US-Sdi source is a ghost population deriving from H-Am1 according to the Figure plotted in Figure 13B. However, as indicated by the error message generated by the *fit.graph* function, such a scenario is not fittable because the resulting system of equations is singular.

```
#Joint positioning of US-Wat and US-Sdi on the am.scaf graph
am.scafWatSdi=graph.builder(am.scaf.fit,leaves.to.add = c("US-Wat","US-Sdi"),dsu.fstats,
                            drift.scaling=TRUE,verbose = FALSE)
```

```
#An alternative (not fittable) admixture graph scenario
am.scafWatSdi.sc<-rbind(c('Am2','S-Am2','B'),c('Am2','CN','(1-B)'),
                        c('BR-Pal','Cal',''),c("Cal","Am2",""),
                        c("CN","R",""),c('S-Am2','Am1',''),
                        c('US-Sok','W',''),c('CN-Shi','CN',''),
                        c("W","S-Am2",""),c('Am1','J-Am1','A'),
                        c('Am1','H-Am1','(1-A)'),c('J-Am1','JP',''),
                        c('JP-Sap','J-Am1',''),c('US-Haw','H-Am1',''),
                        c('H-Am1','JP',''),c('JP','R',''),c('CN-Bei','CN',''),
                        c("US-Wat","SW",""),c("SW","W","D"),c("SW","Cal","(1-D)"),
                        c("US-Sdi","SSdi",""),c("SSdi","Cal","E"),
                        c("SSdi","S-SSdi","(1-E)"),c("S-SSdi","H-Am1",""))
am.scafWatSdi.sc.par=generate.graph.params(am.scafWatSdi.sc,dsu.fstats)
```

```
Total Number of Parameters: 19 (15 edges lengths + 4 adm. coeff.)
Total Number of Statistics: 28 (7 F2 and 21 F3)
```

```
#The graph can be plotted using plot(am.scafWatSdi.sc.par) but
#This scenario is not fittable as indicated by the error message from the fit.graph function
am.scafWatSdi.sc.fit=fit.graph(am.scafWatSdi.sc.par,drift.scaling = T)
```

```
The system is singular: the rank of the incidence matrix is lower than n.edges-1 (e.g., some branches a
It is not possible to properly fit this admixture graph.
```

In agreement with the analysis of $f_3$-based admixture tests results that suggested complex admixture histories among these closely related populations, no proper admixture graph could be found when trying to position US-Col, US-Wis or US-Nca onto the *am.scaf* scaffold graph (Figure 12). Nevertheless, for each of these populations, all the resulting best fitted graph suggested a high contribution of the Am2 admixed source (BR-Pal being its best proxy) that may be closely related to the common source of US-Sdi and US-Wat mentioned above, a second contributing source being related to Japanese populations.

Finally, no proper admixture graph could be obtained when trying to jointly include each possible pairs of the five populations:

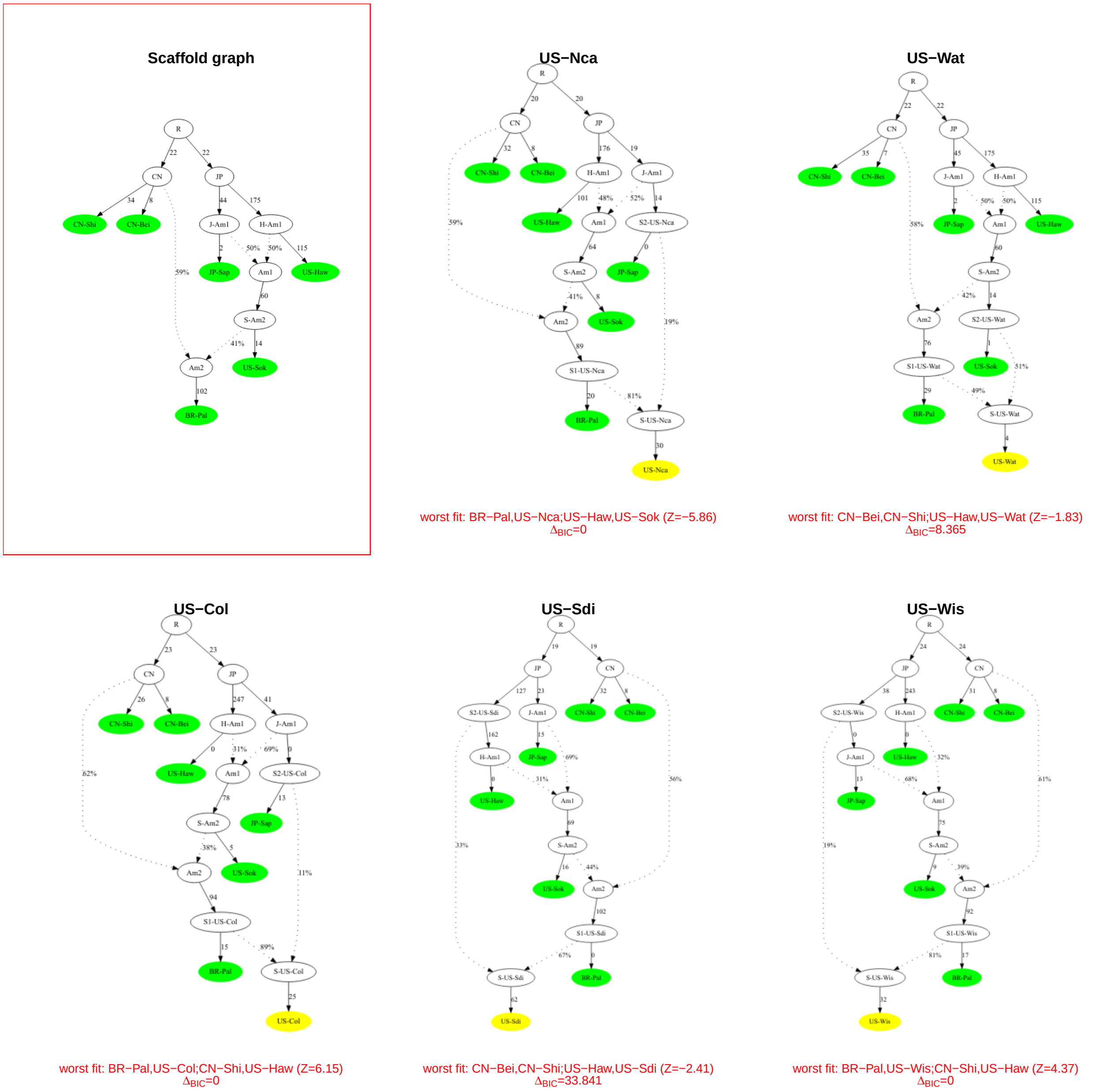

Figure 12: Admixture graphs resulting from the positioning of US-Col, US-Nca, US-Sdi, US-Wat and US-Wis onto the am.scaf scaffold graph that relates the scaffold tree of native populations and BR-Pal and US-Sok (red frame). The best fitting graphs (as obtained with the function add.leaf) are displayed together with i) the worst fitted f-statistics and their associated Z-score; and ii) the difference of their BIC with respect to the graphs displaying the second lowest BIC ($\Delta_{BIC}$) as a measure of support. The target populations are highlighted in yellow. For all the graphs, the fitted edge lengths are in drift units (x1,000) since *drift.scaling* argument was set to *TRUE*.

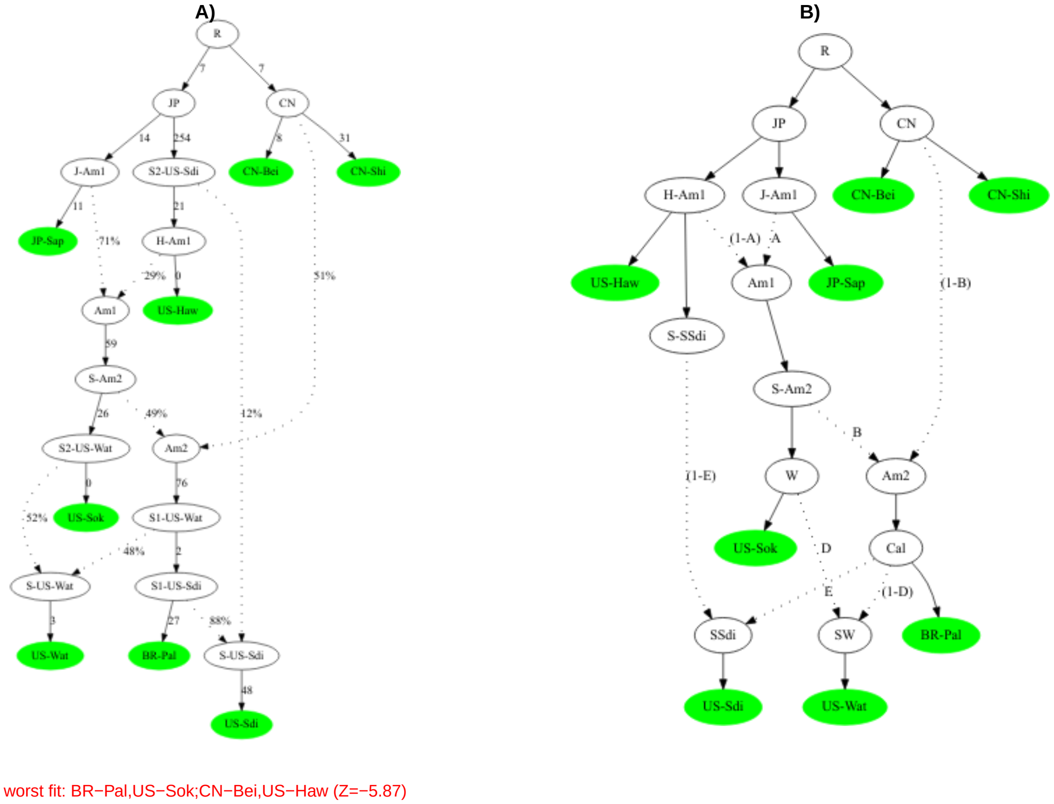

Figure 13: Connecting US-Wat and US-Sdi to the am.scaf scaffold graph. A) Best fitted admixture graph obtained with the graph.builder function (the fitted edge lengths are in drift units (x1000) since *drift.scaling* argument was set to *TRUE*). B) Possible (but not fittable) scenario connecting US-Wat and US-Sdi to the am.scaf scaffold graph.

```r
cleangrid<-function(x){n=ncol(x);x[apply(x,1,f<-function(y){length(unique(y))==n}),n:1]}
pops.to.include<-expand.grid(c("US-Col","US-Nca","US-Sdi","US-Wat","US-Wis"),
                             c("US-Col","US-Nca","US-Sdi","US-Wat","US-Wis")) %>%
                             as.matrix() %>% cleangrid()
#exploring  and evaluating the graphs
for(i in 1:nrow(pops.to.include)){
 tmp.graph<-graph.builder(am.scaf.fit,leaves.to.add = pops.to.include[i,],
                           fstats=dsu.fstats,verbose=F)
 tmp.comp.fit=compare.fitted.fstats(dsu.fstats,tmp.graph$best.fitted.graph,
                                    n.worst.stats = 0)
  if(max(abs(tmp.comp.fit$`Z--score`))<3){
  cat("##########################################\n")
  cat("Inclusion Order:",pops.to.include[i,],"\n")
  #number of graphs within Dbic<6 (default)
  cat("\tNumber Of Graphs:",tmp.graph$n.graphs,"\n")
  #min BIC
  cat("\tMin BIC:",min(tmp.graph$bic),"\n")
  #worst fitted fstats (best fitting graph)
  cat("\tWorst fstats fit:\n")
  print(tmp.comp.fit[which.max(abs(tmp.comp.fit$`Z--score`)),])
}
 }
```
